# Discovery and optimization of a guanylhydrazone-based small molecule to replace bFGF for cell culture applications

**DOI:** 10.1101/2025.01.30.635029

**Authors:** Mikhail Feofanov, Gerrit Martin Daubner, Andrea Saltalamacchia, Karsten Köhler, Christine Schulz, Clare Elizabeth Henry, Michael Josef Ziegler, Mohammed Benabderrahmane, Florence Andrée Suzanne Hiault, Tim-Michael Decker, Mei-Chun Shen, Jürgen Pahl, Sophie Lambertz, Hamid R. Noori

**Author notes:** These authors contributed equally (Mikhail Feofanov, Gerrit Martin Daubner).

## Abstract

Replacing growth factors with a synthetic alternative molecule is an attractive opportunity to increase consistency, scalability and cost-effectiveness of cell-based products. Herein, we describe the discovery of a chemical class of FGFR1 agonists mimicking the action of basic fibroblast growth factor (bFGF), an essential component of cell-culture media. The guanylhydrazone-based molecule, TCB-32, was identified via structure-based virtual screening on the orthosteric binding site of FGFR1. It was shown to significantly increase cell proliferation by activating the FGFR1 signalling pathway like bFGF and exhibited enhanced thermostability over bFGF by containing activity over the course of several days. After extensive structure-activity relationship studies it was possible to increase potency and efficacy leading to three highly potent agonists. This finding has the potential to remove current bottlenecks in large-scale cell production as required for applications like cultivated meat or cell therapy.

**GRAPHICAL ABSTRACT:** 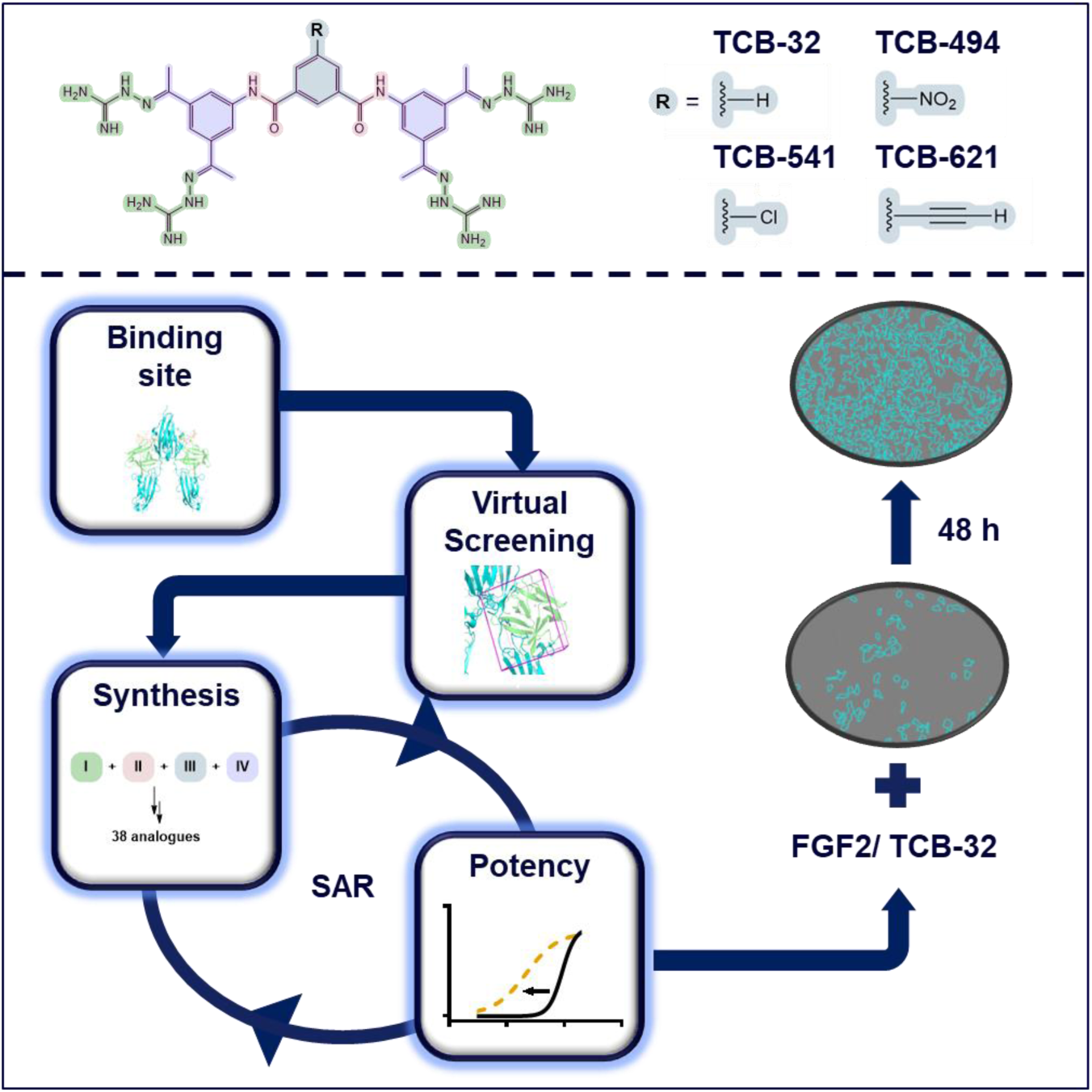

**HIGHLIGHTS:**

- Discovery of a highly potent small molecule to replace bFGF in serum-free medium
- Activation of proliferation through the FGFR1-signalling pathway
- Enhanced thermal stability over bFGF
- SAR led to three small molecules with decreased EC50, named TCB-494, TCB-541 and TCB-621

## INTRODUCTION

Human embryonic stem cells (hESCs) or human-induced pluripotent stem cells (hIPSCs) require precisely controlled cellular responses to maintain pluripotency or to induce differentiation. It is therefore essential to use a cell-culture medium that has a refined list of ingredients at known concentrations. This requirement is in stark contrast to medium supplemented with fetal bovine serum (FBS) routinely used for immortalised cells. While it has superior growth promoting features, it contains an unknown assembly of factors and is subject to huge batch-to-batch variations. Therefore, many efforts to decrease the dependency on FBS and to develop a serum-free media for stem cells were initiated. The first important advancement towards a serum-free medium was the publication of TeSR medium in 2006 that required basal medium and 18 additional components to maintain stem cells. ^1,2^ While this enabled more precise control over cellular responses compared to serum-containing medium, it was still challenging to ensure the quality and to determine the specific functions of so many components. Therefore, the development of Essential 8 from the same research laboratory five years later was a huge advancement, since it reduced the number of additional media components.^3^

The emergence of new technologies like cell therapy and cultivated meat applications is driving the demand for even more refined serum-free conditions, largely due to regulatory compliance and the requirements of large-scale cell production. The significance of improved serum-free media is highlighted by the challenges faced by the cell therapy industry, which is struggling to secure funding in 2024 mainly because of manufacturing and scalability problems.^4^ These issues result in excessively high production costs making it difficult for even the most promising treatments to recover their investment.^5^ One way to further optimise serum-free media is to replace the four protein-components in Essential 8 medium (bFGF, TGF-β1, Insulin, Transferrin), since proteins are expensive to produce, have high batch-to-batch variability, are difficult to handle and thus have limited scalability. Of special interest is bFGF, since it was identified to be the most expensive ^6^ and indispensable ^7^ component for serum-free media compositions.

One approach to replace bFGF is to engineer cells that ectopically express the bespoken protein as shown for immortalised bovine satellite cells.^8^ While this significantly reduces media costs, the process to generate this cell line described in the publication is complex and must be repeated for each new cell line. In addition, genetic-modified organism (GMO) face high regulatory hurdles for commercial applications, especially in the European Union.^9^ Another approach is to produce more stable variants of growth factors as shown for FGF2-G3. This variant retains full biological activity after 20 days at 37°C due to nine unique point mutations that were identified by minimization of the Gibbs free energy of the native state.^10^ This increase in stability permits the usage of lower amounts of protein to decrease costs, but it is still required to express and purify a protein in large-scale using a biological organism. The latter is also true for other approaches like peptide-based C19jun ^11^ or agonistic single-domain antibodies.^12^ It is still an unfulfilled hope of the industry that costs of protein production decrease once the high demand for growth factors eventually lead to better manufacturing processes and capacities.

Our approach was to identify a synthetic small molecule to replace bFGF in serum-free medium. Such a chemical is produced using an easily scalable manufacturing process that reduces costs and leads to more consistent quality compared to proteins purified from biological systems. To achieve our goal, we identified the central interactions at the bFGF-FGFR protein-protein interaction site and conducted a virtual structure-based screen to identify potential binders. As a result, we identified a guanylhydrazone-based small molecule named TCB-32 that had a strong proliferative effect in NIH 3T3 cells, activated the same FGFR1-signalling pathway as bFGF and showed increased thermostability. We could further optimise its activity using structure-activity relationship (SAR) studies and finally identified three molecules that showed increased potency and efficacy for cell proliferation.

## RESULTS

### Proliferation assay to screen small molecules

Virtual screening approaches for identification of novel small molecule binders become increasingly popular due to their ability to decrease infrastructure costs and potentially overcome the possible biases connected to traditional high throughput screening of large physical libraries^13^. We wanted to figure out if a ligand-based or structure-based computational approach is more suitable for finding a novel FGFR1 activating molecule. Therefore, we first tested those small molecules that were claimed to act as bFGF replacement in the literature. To develop an assay to measure their activity, we used the fibroblast mouse cell line NIH 3T3, that is widely accepted as gold standard for assessment of growth factors in cell culture media and was previously applied to assess growth factor activity in low serum-containing medium.^14^ To be able to identify a small molecule to replace bFGF, we needed to avoid serum-containing medium due to its undefined protein composition that could lead to a high background and mask effects. Therefore, we adapted the protocol from low serum-containing to serum-free conditions in a 384-well plate format. For this reason, cells were plated in Essential 8 medium without bFGF and TGF-β1, stimulated by FGF2-G3, grown for 48 hours and measured by CellTiter-Glo^®^ 2.0 reagent. The cells showed a standard growth curve that reached its maximum at 10 ng/ mL (≈ 0.6 nM) and plateaued at higher concentrations (**Fig. 2A**). We then used identical assay conditions without FGF2-G3 to test if small molecules can induce proliferation by their own action to a similar degree than the growth factor. However, the four literature-based molecules SUN 11602 ^15^, dimeric SSR128129E (**SI-1**) ^16^ , ID-8 and FK-506 ^17^ did all not induce proliferation (**Fig. S1**). This either means that they are not suitable to induce proliferation or are missing an essential cofactor in our assay conditions. Based on these results and the availability of crystal structure of FGF2-FGFR1 complex, we decided to address the challenges of discovering a novel FGFR1-activating molecule through virtual structure-based screening based on the orthosteric binding site of FGF2 on FGFR1.

### Analysis of FGFR1 binding site and virtual screening

The starting point for our development was the analysis of the interaction between FGF2 an FGFR1 **(Fig.1. A)** at the orthosteric binding site. Due to the size of the protein-protein interaction area of ca. 1344 Å², that is much larger than typical average contact area between a small molecule and protein (which is estimated to be between 300 to 1000 Å^2^ ^18^), it was necessary to discretize it and identify the relevant interactions . Protein-protein interaction hot spots were collected from mutational studies,^19–24^ crystal structure contacts,^25,26^ persisting interactions in the MD simulation, and strong interactions in the QM/MM decomposition **(Fig. 1B)**.

**Figure 1.**
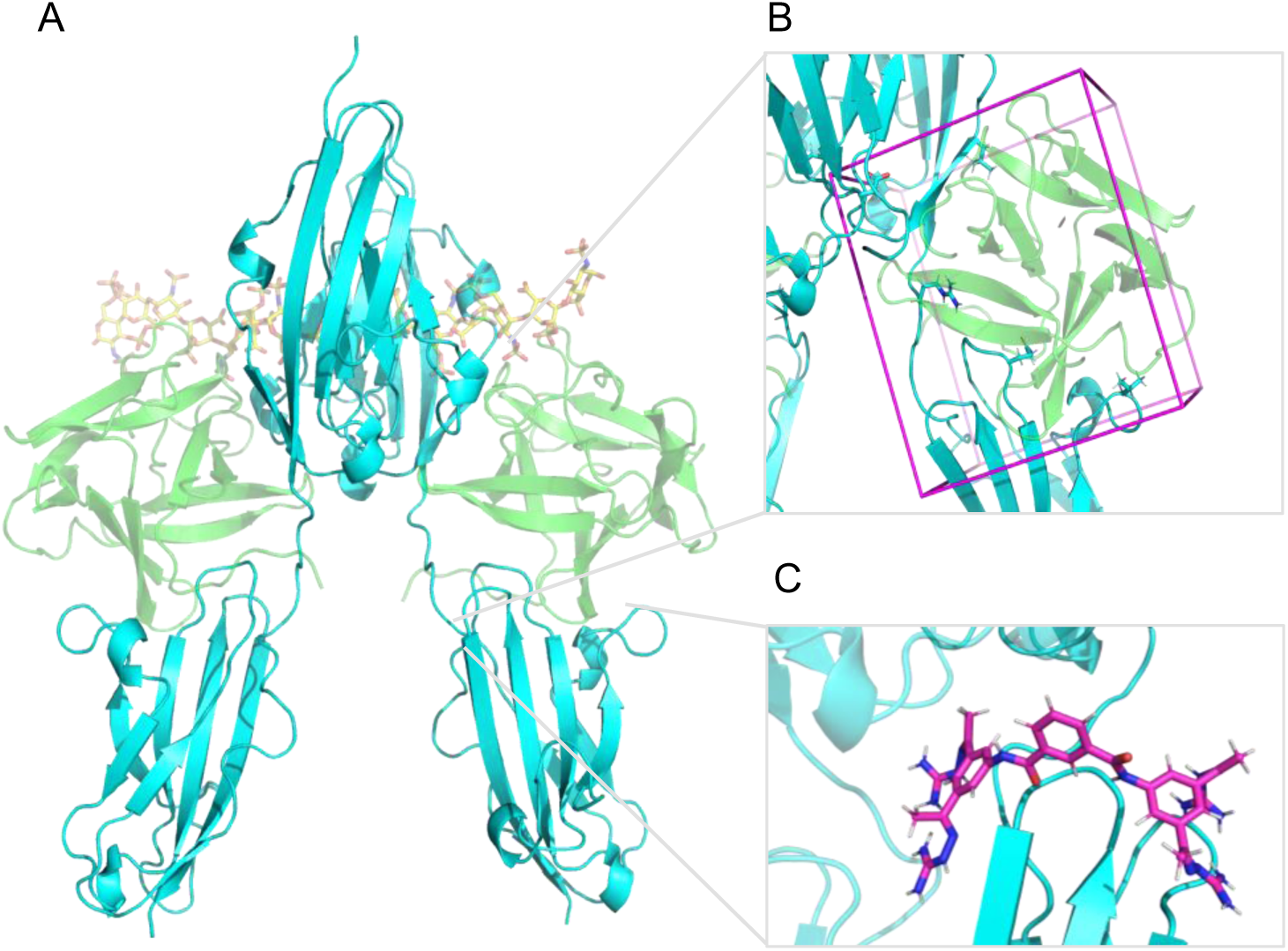
(A) Molecular representation of the FGF2/FGFR1 complex model used for simulation. FGFR1 in cyan, FGF2 in green. (B) Relevant hot spot residues (depicted as sticks) of FGFR1 in the FGF2/FGFR1 interaction site encapsulated by the docking box (magenta). (C) Most stable position of TCB-32 (magenta) after MD simulations on the FGF2/FGFR1 interaction site at FGFR1.

**Figure 2.**
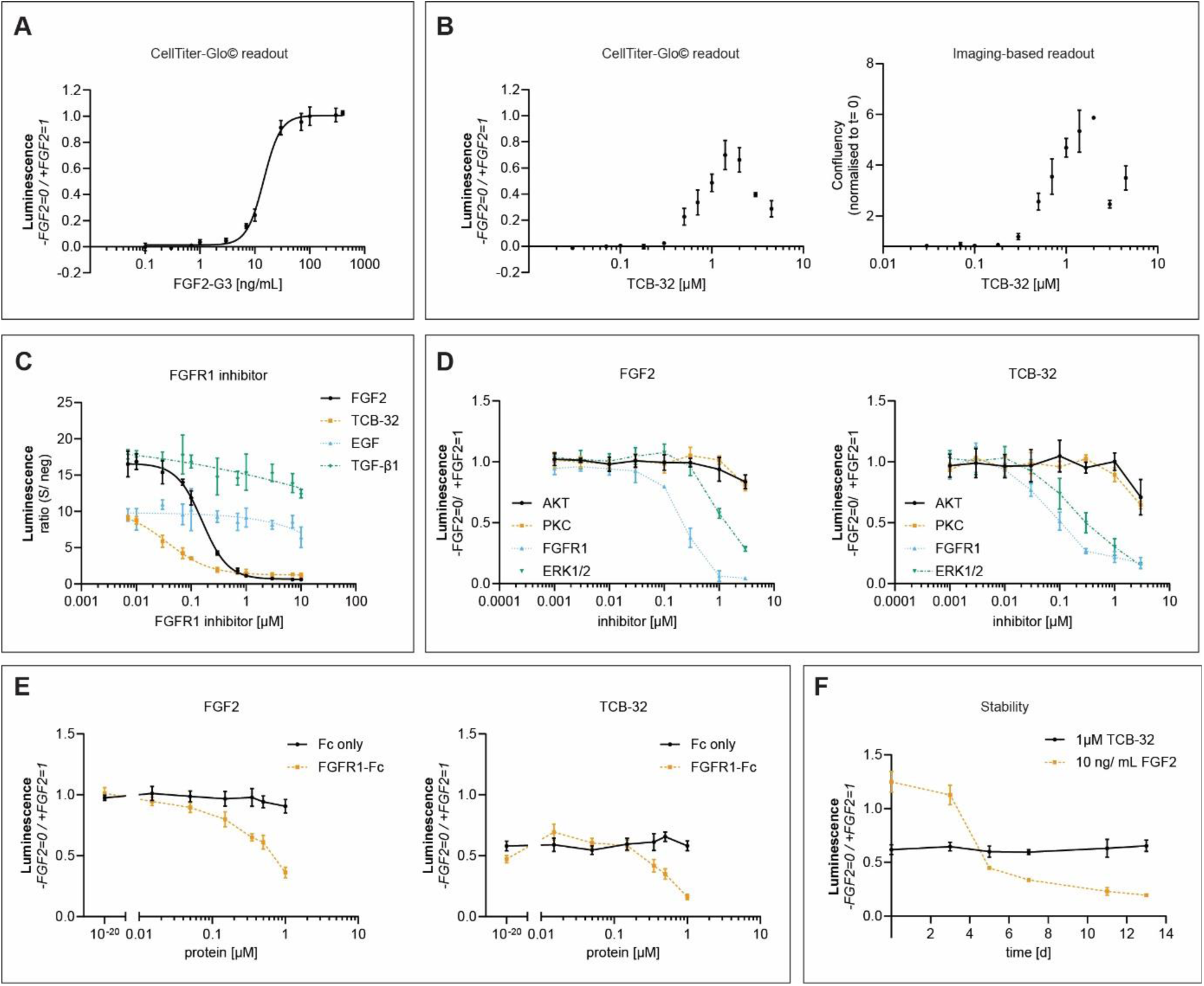
(A) Proliferation assay with FGF2 (n= 4, SD, z’=0.80). (B) Proliferation assay with TCB-32 using as readout (**left**) CellTiter-Glo^®^ 2.0 (n= 4, SD, z’= 0.73), (**right**) imaging after 48 hours (n= 4, SEM). (C) NIH 3T3 cells stimulated with 40 ng/ mL FGF2-G3, 1 μM TCB-32, 100 ng/ mL EGF and 2 ng/ mL TGF-β1 while being treated with increasing concentrations of FGFR1 inhibitor (n= 4, SD, z’= 0.78, outliers removed that showed ≥ 10x difference within replicates). (D) NIH 3T3 cells stimulated with (**left**) 100 ng/ mL FGF2-G3 (**right**) 1.4 μM TCB-32 while being treated with AKT (AZD5363), PKC (Bisindolylmaleimide 1), FGFR1 (PD166866) and ERK 1/2 (SCH772984) inhibitors. (n= 4, SD, z’= 0.67). (E) Stimulation of NIH 3T3 cells with (**left**) 30 ng/ mL FGF2-G3 and (**right**) 1.4 μM TCB-32 while being exposed to increasing concentrations of either FGFR1-Fc or only the Fc domain (n= 4, SD, z’= 0.75). (F) Samples of TCB-32 and wildtype FGF2 were incubated for several days in the cell-culture incubator and their activity tested in the proliferation assay in NIH 3t3 cells (n= 4, SD, z’= 0.94).

To accommodate such large size of the interaction site, molecules with a molecular weight over 500MW were selected from ZINC database (one of the most used ultra large chemical databases) ^27^ for docking, totalling 9 million molecules. After filtering and docking, the molecules (72 in total) showing the best GLIDESP docking score (below -6 kcal/m) were tested in MD simulation (**Fig. 1C**) and analysed for undesired physicochemical and biological properties and synthetic complexity. Based on this assessment only 7 potential FGFR1 binders remained, showing the complications of identifying agonistic small molecules that mimic binding of bFGF to such a large orthosteric binding site.

### Identification and cellular characterization of TCB-32

From the 7 small molecules tested in the proliferation assay, only one hit was identified. The remaining molecules were tested with biochemical (HTRF, nanoDSF) and cellular (One-Glo™ Luciferase Assay System) assays, but no further FGFR1-binding or -activating molecule was found. Our hit TCB-32 (**1**) is a tetravalent aromatic guanylhydrazone that was able to induce proliferation with 70% ± 10% of the level of FGF2-G3 in our assay conditions (**Fig. 2B left**). Evaluation of cell-imaging data showed that this is indeed due to an increase in cell number and not an artifact due to changes in cell size or altered ATP metabolism as reported previously for CellTiter-Glo^®^ 2.0 (**Fig. 2B right**).^28^ In contrast to FGF2-G3 (**Fig. 2A**), the dose-response curve was bell shaped at concentrations higher than 2 μM, which is likely due to precipitation of TCB-32 that is visible on images. A toxic effect was ruled out with an LDH-Glo^™^ Cytotoxicity assay in HT-1080 cells using serum-containing cell medium suggesting that toxicity starts at 30 μM (**Fig. S2A**). We also confirmed that TCB-32 was able to maintain its proliferative effect for at least 3 passages in C2C12 cells grown in serum-free cell medium (**Fig. S2B**). To proof our hypothesis that TCB-32 is binding to FGFR1 as determined by the virtual screen, we first investigated if the observed effect on proliferation is mediated through the FGFR1 pathway.

To determine the impact of TCB-32 on FGFR1 activity, we used the inhibitor PD166866 that was originally identified in NIH 3T3 cells to inhibit bFGF-stimulated FGFR1-tyrosine kinase activation.^29^ We added the inhibitor to the cell medium, incubated for 30 minutes and then stimulated proliferation by either adding EGF, TGF-β1, FGF2-G3 or TCB-32 (**Fig. 2C**). We chose EGF and TGF-β1 as negative control because those growth factors stimulate growth besides bFGF in Essential 6 medium and should not be affected by the inhibitor. Indeed, both controls remained unaffected by the treatment, which shows that the inhibitor is not toxic at the tested concentrations. In contrast, TCB-32 like FGF2-G3 completely lost its proliferative effect at 1 μM inhibitor concentrations. This illustrates that its action in NIH 3T3 cells is indeed FGFR1-dependent and not caused by another effect.

Signal transduction after binding of bFGF to FGFR1 leads to the activation of different pathways that have different biological outcomes. Therefore, we investigated whether FGF2-G3 and TCB-32 activate the same downstream pathway to promote proliferation of NIH 3T3 cells. For this reason, we used inhibitors against PKC, AKT and ERK 1/2 to block the components of the three major pathways downstream of FGFR1. We first added those inhibitors to the cell culture medium in a dose-response manner, briefly incubated and then stimulated the cells with either FGF2-G3 or TCB-32 (**Fig. 2D**). We observed that ERK 1/2 was able to inhibit proliferation at non-toxic concentrations while PKC and AKT inhibitors had not effect. This confirms previous reports that ERK1/2 drive proliferation in NIH 3T3 cells.^30^ With FGF2-G3 and TCB-32 both losing their activity with FGFR1 and ERK 1/2 inhibitors, this suggests that their proliferative effect is promoted by the same downstream pathways.

Next, we wanted to investigate if TCB-32 is activating the FGFR1 signalling pathway by directly interacting with FGFR1. We setup a competition assay, where we expect a decrease in proliferation if TBC-32 interacts with recombinantly instead of endogenously expressed FGFR1. To produce the recombinant protein, we cloned the extracellular domain of FGFR1 comprising domains 1 to 3 and added a Fc-domain at the C-terminus for dimerization. Then we expressed the protein in mammalian cells to maintain correct folding and post-translational modifications. We conducted the proliferation assay, where we added FGF2-G3 or TCB-32 at a fixed concentration and the recombinant protein in a dose-response manner simultaneously to the cell culture medium (**Fig. 2E**). While the Fc domain by itself did not have any effect on activity, addition of recombinant FGFR1-Fc lead to decreased activity of FGF2-G3 and TCB-32. This is evidence that both directly interact with FGFR1 in a cellular context, and that it is this interaction that leads to activation of the FGFR1 signalling pathway. Unfortunately, we were unable to gather direct evidence for a binding of TCB-32 to the orthosteric site of FGFR1 as postulated by the virtual structure-based screening using biochemical assays. We speculate that, due to the high positive charge of the aminoguanidine moieties, TCB-32 interacts with dyes and destabilises proteins in vitro.

Thermal stability is a main advantage of synthetic small molecules over proteins in cell culture media, since it improves consistency and paves the way for large-scale cellular biomass production. A prominent example would be the increase of biomass for cultivated meat or the increase in cell number for cell therapeutic applications. Therefore, we compared the stability between TCB-32 and wildtype bovine FGF2, which would be used to produce cultivated meat from porcine cells and would face less regulatory hurdles than FGF2-G3. We incubated the stock solutions of both reagents in the cell culture incubator and afterwards tested their activity in our cell proliferation assay in a time-dependent manner (**Fig. 2F**). While TCB-32 did not show loss of activity over the entire 13-day test period, wildtype bovine FGF2 showed already a decrease of activity after 3 days and was below TCB-32 levels after 5 days. This observation suggests superior stability of TCB-32 in comparison with the wildtype bovine FGF2 and that this molecule constitutes a promising starting point to optimise its potency and/ or efficacy. While the former would decrease the concentration of this molecule in the final cell biomass, the latter would lead to a higher proliferation rate similar than bFGF.

### Structure-activity relationships exploration of TCB-32

Since we were unable to prove our original hypothesis that TCB-32 binds to the orthosteric site, we introduced stepwise modifications of the molecule via chemical synthesis to identify the essential parts responsible for its activity and potentially find a possibility to increase its activity. The molecule was divided into four regions: aminoguanidine arms (**I**), linker connection (**II**), linker region (**III**) and tripod region (**IV**) (**Fig.3A**). We then introduced modifications in those four regions and measured their effects on potency and efficacy in our proliferation assay (**Table 1**). We focused our analysis mainly on potency, since it was very robust in contrast to efficacy, which was sensitive to small variations in cell numbers.

**Table 1:**
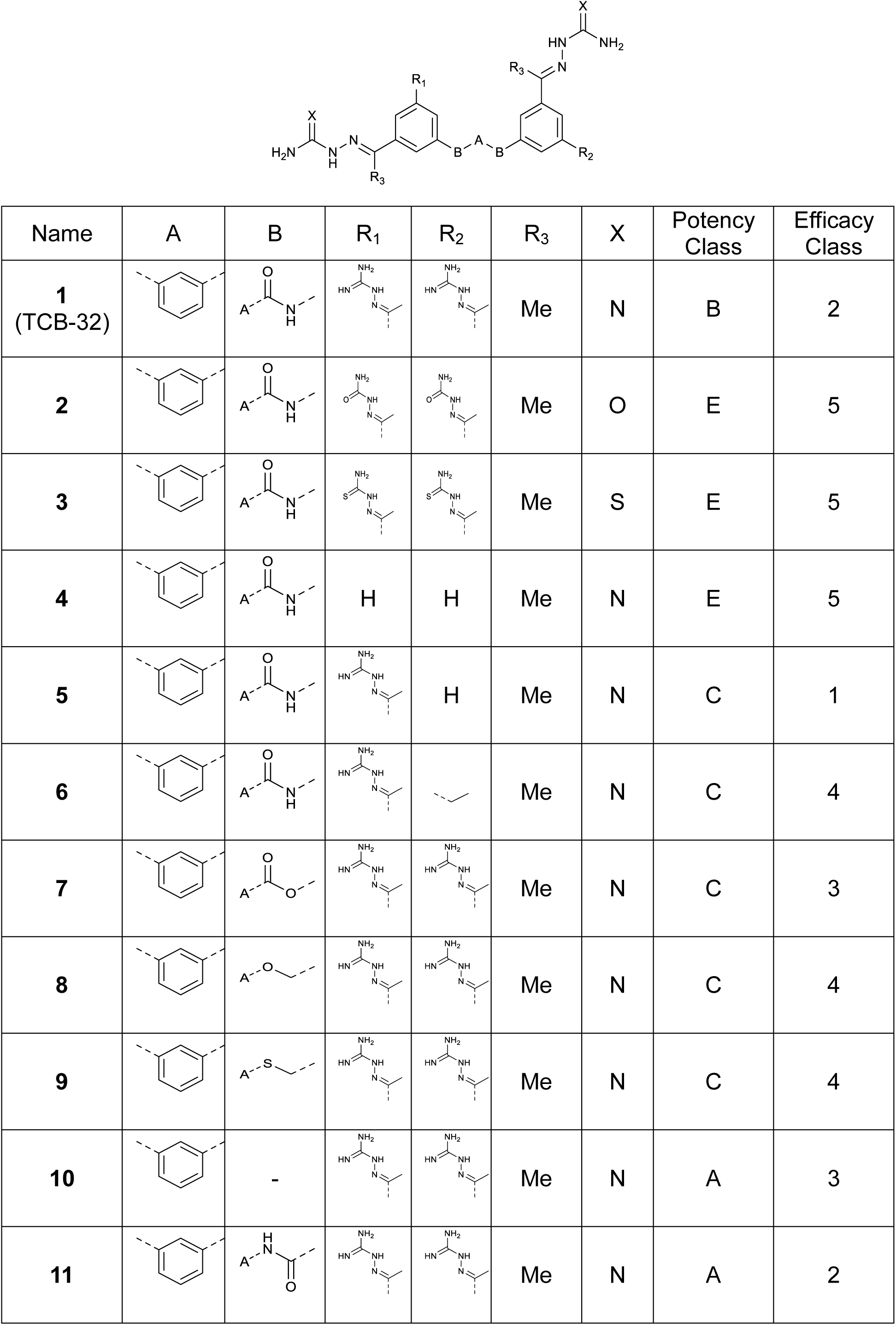
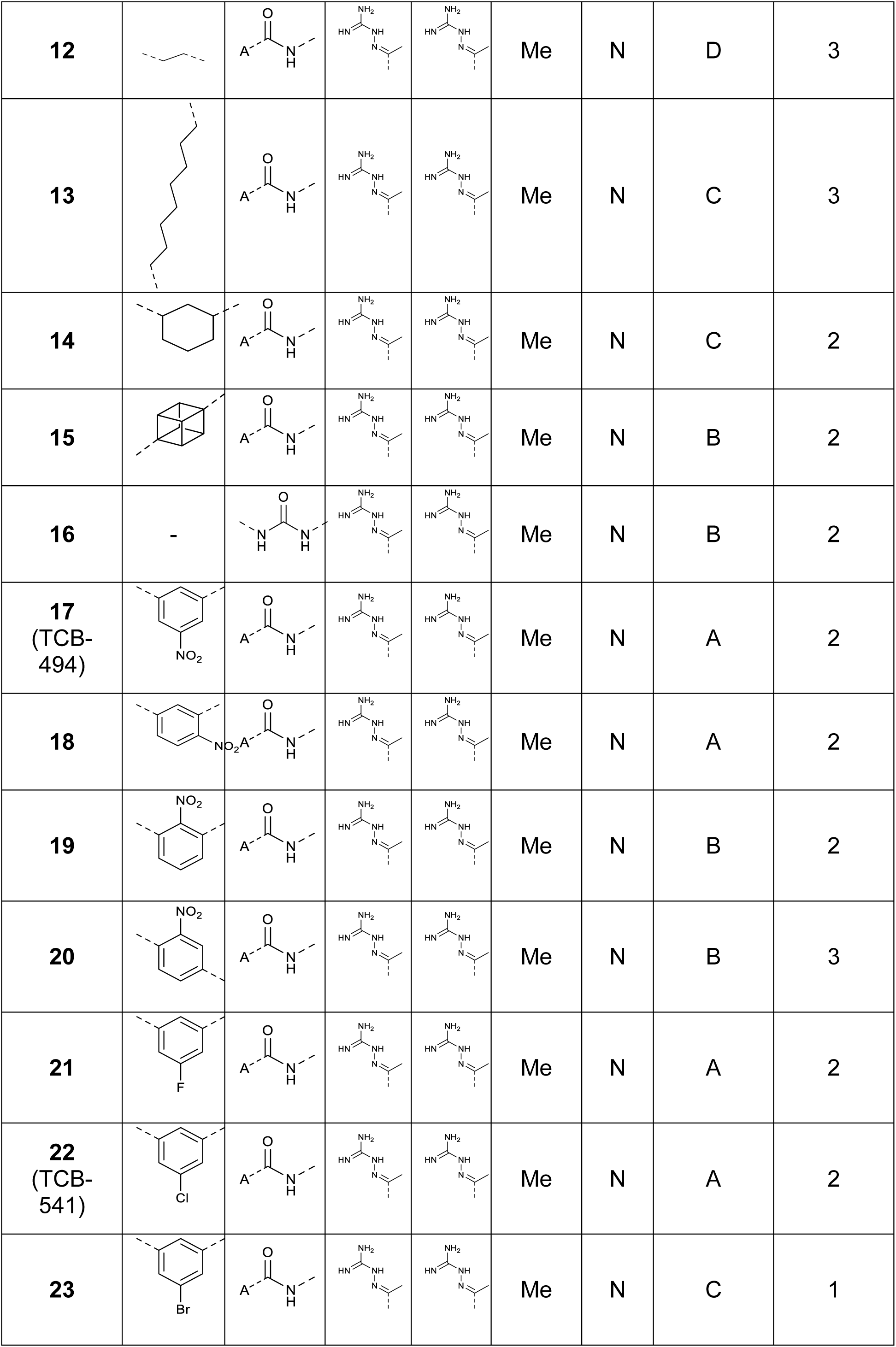
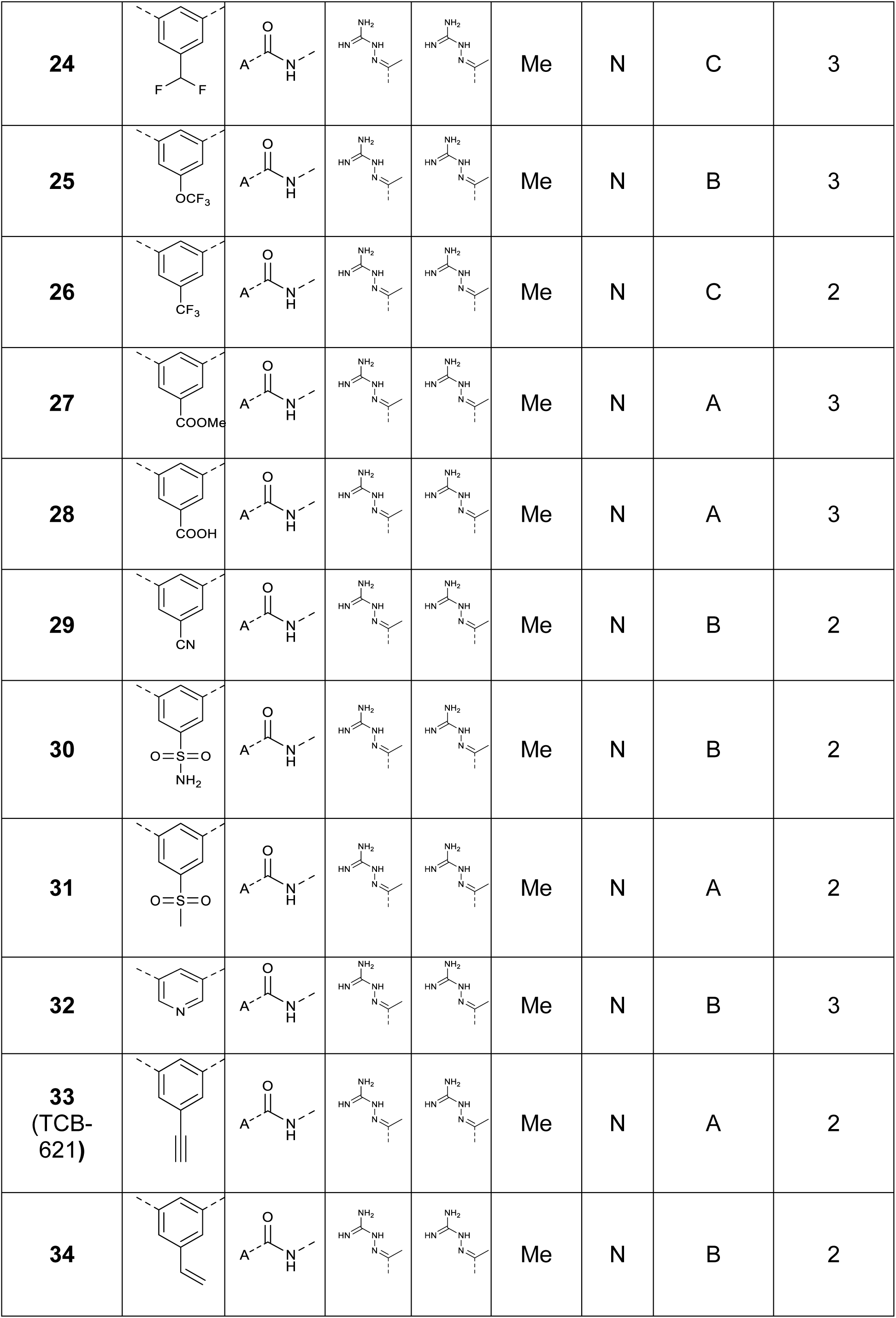
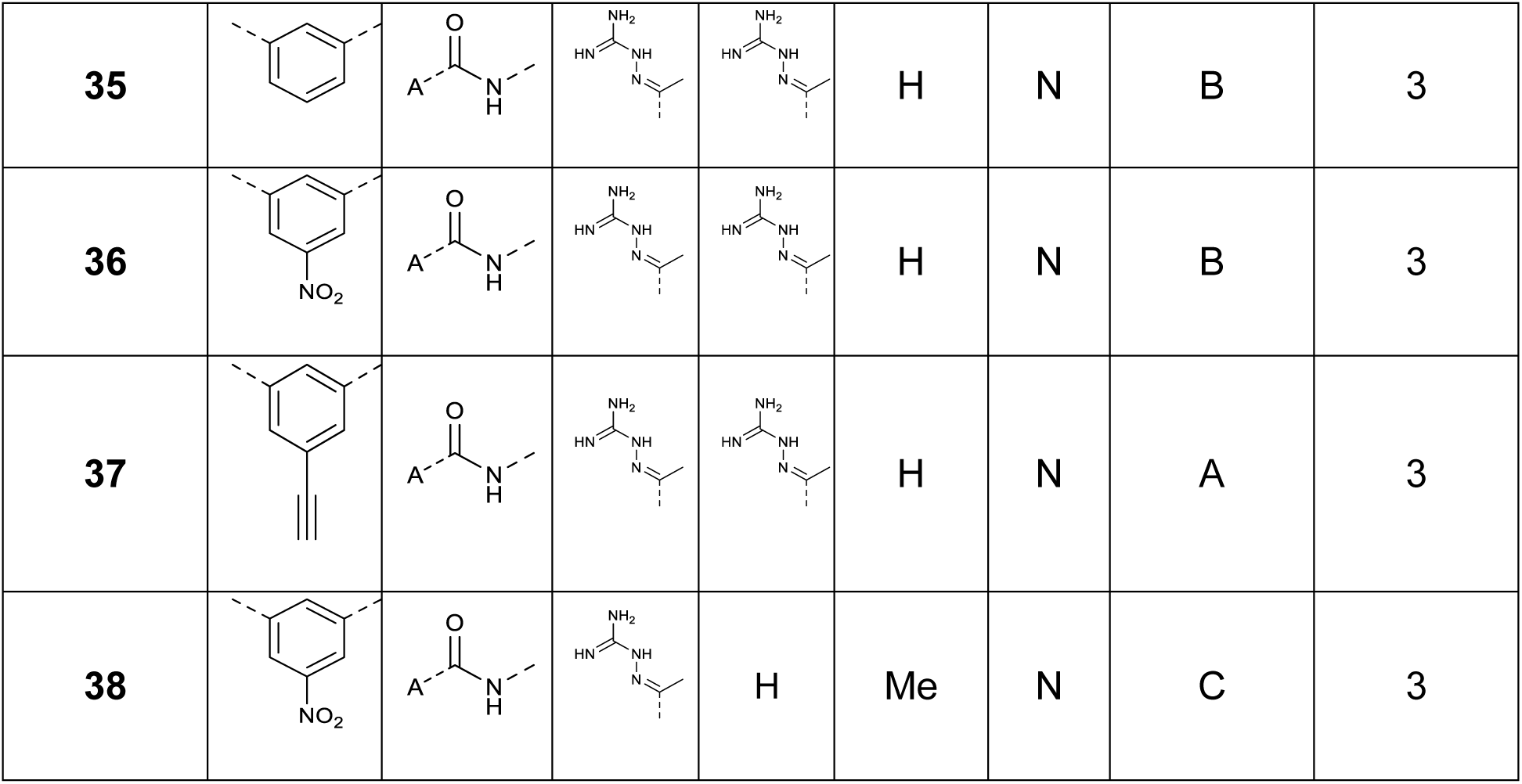
Overview of compounds described in this study.

| Name | A | B | R<sub>1</sub> | R<sub>2</sub> | R<sub>3</sub> | X | Potency Class | Efficacy Class |
| --- | --- | --- | --- | --- | --- | --- | --- | --- |
| **1**<br>(TCB-32) |  |  |  |  | Me | N | B | 2 |
| **2** |  |  |  |  | Me | O | E | 5 |
| **3** |  |  |  |  | Me | S | E | 5 |
| **4** |  |  | H | H | Me | N | E | 5 |
| **5** |  |  |  | H | Me | N | C | 1 |
| **6** |  |  |  |  | Me | N | C | 4 |
| **7** |  |  |  |  | Me | N | C | 3 |
| **8** |  |  |  |  | Me | N | C | 4 |
| **9** |  |  |  |  | Me | N | C | 4 |
| **10** |  | - |  |  | Me | N | A | 3 |
| **11** |  |  |  |  | Me | N | A | 2 |
| 12 |  |  |  |  | Me | N | D | 3 |
| 13 |  |  |  |  | Me | N | C | 3 |
| 14 |  |  |  |  | Me | N | C | 2 |
| 15 |  |  |  |  | Me | N | B | 2 |
| 16 | - |  |  |  | Me | N | B | 2 |
| 17<br>(TCB-494) |  |  |  |  | Me | N | A | 2 |
| 18 |  |  |  |  | Me | N | A | 2 |
| 19 |  |  |  |  | Me | N | B | 2 |
| 20 |  |  |  |  | Me | N | B | 3 |
| 21 |  |  |  |  | Me | N | A | 2 |
| 22<br>(TCB-541) |  |  |  |  | Me | N | A | 2 |
| 23 |  |  |  |  | Me | N | C | 1 |
| 24 |  |  |  |  | Me | N | C | 3 |
| 25 |  |  |  |  | Me | N | B | 3 |
| 26 |  |  |  |  | Me | N | C | 2 |
| 27 |  |  |  |  | Me | N | A | 3 |
| 28 |  |  |  |  | Me | N | A | 3 |
| 29 |  |  |  |  | Me | N | B | 2 |
| 30 |  |  |  |  | Me | N | B | 2 |
| 31 |  |  |  |  | Me | N | A | 2 |
| 32 |  |  |  |  | Me | N | B | 3 |
| 33<br>(TCB-621) |  |  |  |  | Me | N | A | 2 |
| 34 |  |  |  |  | Me | N | B | 2 |
| 35 |  |  |  |  | H | N | B | 3 |
| 36 |  |  |  |  | H | N | B | 3 |
| 37 |  |  |  |  | H | N | A | 3 |
| 38 |  |  |  | H | Me | N | C | 3 |

**Figure 3.**
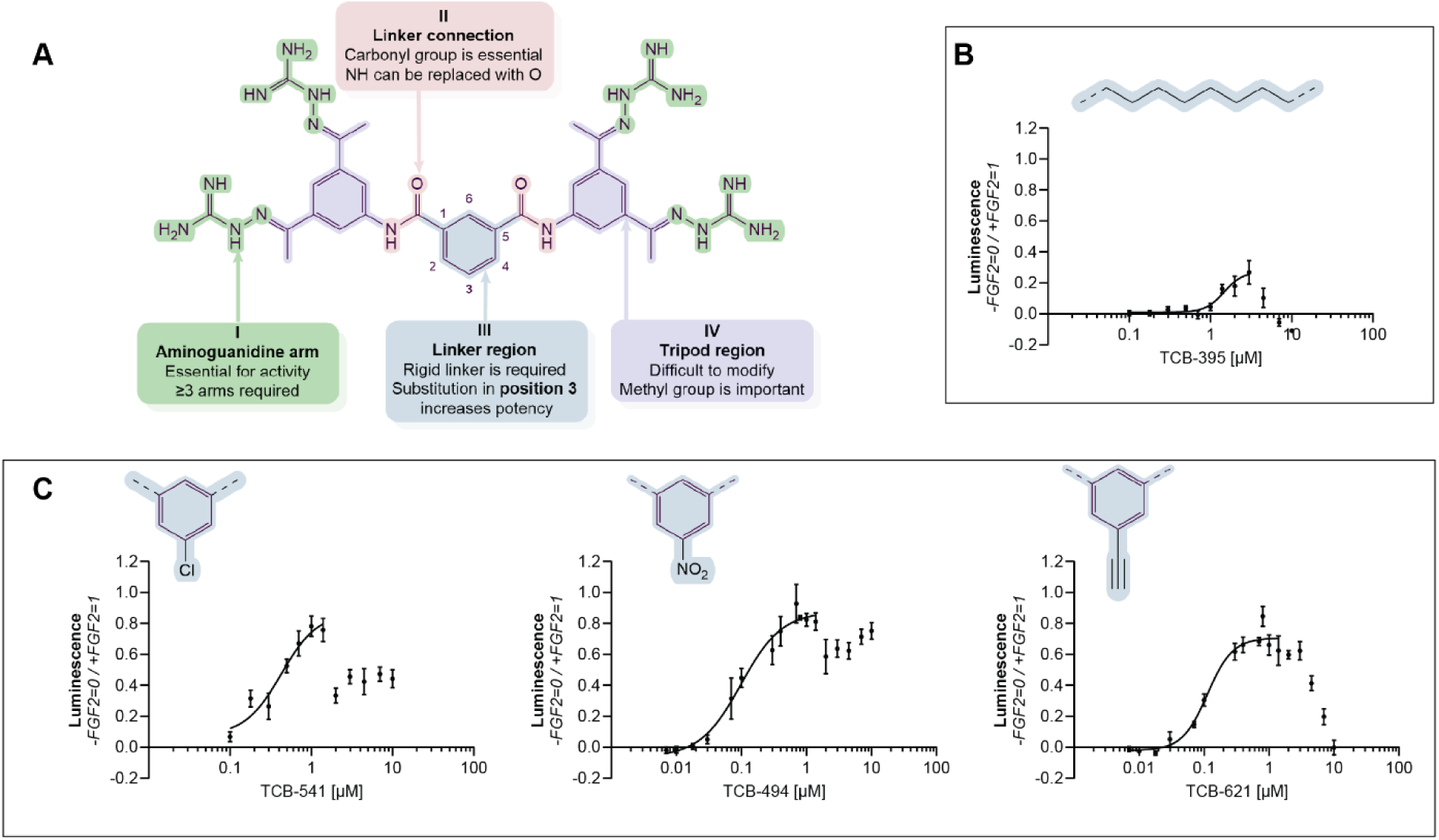
(A) Optimisation of TCB-32 based on the four different regions: **I** Aminoguanidine arm, **II** Linker connection, **III** Linker region and **IV** Tripod region. (B) Proliferative activity of the structurally similar compound **13** (CNI-1493/ Semapimod) (n= 4, SD, z’= 0.78). (C) Proliferative activity of the three most promising analogues (left) TCB-541 (n= 4, SD, z’= 0.78, (middle) TCB-494 and (right) TCB-621 (n= 4, SD, z’= 0.71).

Starting with region **I**, two analogues containing semicarbazone (**2**) and thiosemicarbazone (**3**) moieties were synthesized. Substitution of nitrogen to oxygen or sulphur led to a complete loss of activity. This highlighted the importance of positive charge and strong basicity of region **I.**^31^

Subsequently, the necessary number of aminoguanidine arms was evaluated. A series of molecules containing two (**4**) and three (**5**-**6**) arms was designed. While **4** did not show any activity at all, **5** showed only a slight decrease in potency as determined by EC50 compared to **1,** but at the same time had higher efficiency. This was a surprising discovery, especially since **6**, which lacks only the aminoguanidine part whereas maintaining aliphatic chain of the arm, had very insignificant activity. Since it was easier to screen changes in **II** and **III** having a structure containing 4 aminoguanidine arms (2 less synthetic steps), it was decided that when the best substituents in **II** and **III** will be identified, additional analogues containing 3 arms will be tested.

To identify the importance of **II**, several analogues containing ester (**7**), ether (**8**), thioester (**9**), or no **II** at all (**10**) were synthesized. The potency significantly decreased if the carbonyl group was not present, but it did not depend on the type of heteroatom. At the same time, complete lack of **II** lead to increase of potency, but significantly decreased the efficiency, probably, due to lower solubility. Interestingly, an exchange of positions of carbonyl group and amine did lead to a slight decrease of potency (**11**). Following these results, amide linker connection was chosen as the best, and was not modified anymore.

Modifications in **III** were found the easiest to vary due to the broad availability of aromatic and aliphatic dicarboxylic acids. First, more flexible saturated chains were evaluated (**12**-**14**). Interestingly, **13** is an investigational drug for Crohn’s disease called CNI-1493 (Semapimod) and was previously evaluated as the best analogue in the series of tetraguanilhydrazones.^32^ However, it was less active than TCB-32 in our experimental setup. To confirm the importance of structural rigidity, a bioisosteric cubane moiety (**15**) was installed, and it led to the similar activity compared to TCB-32. Lack of **III** was also (**16**) evaluated, and the activity was also like TCB-32. Since it is much easier to tune the structure in the aromatic ring, we focused on the installation of substituents in the phenyl ring.

Due to the in-house availability of 5-nitroisophthalic acid required for the synthesis of the 3,5-diacetylaniline, naturally we tested it first. This surprisingly led to the discovery of TCB-494 (**17**) that showed increased potency. This made it possible to fit the curve and revealed an EC50 of approx. 0.1 µM that is lower than the approx. 0.7 µM observed for TCB-32 (**Fig. 3C**). Worth to note that nitro groups in other positions of the aromatic ring did not lead to such improvement of potency (**18**-**20**).

Although the findings were quite encouraging, nitro groups can show mutagenic and genotoxic properties that potentially pose a limit for future applications.^33^ Although an increase in cytotoxicity was not observed in cell culture compared to TCB-32 (**Fig. S2A**), it constitutes an increased risk in a regulatory setting and therefore additional experiments to find alternative substituents were conducted. According to literature, ^34,35^ the optimal way to substitute nitro group is to use another electron-acceptor small substituents. Therefore, the series of molecules containing halogens (**21**-**23**), difluoromethyl or trifluoromethyl (**24**-**26**), carboxymethyl and carboxy (**27**-**28**), cyano (**29**), sulfonamide and sulfone (**30**-**31**) and pyridine ring (**32**). Substitution of nitro group by these groups decreased the potency measured by the EC50. Interestingly, TCB-541 (**22**) and **23** showed the highest efficacy in the series, and TCB-541 had also a decreased EC50. With the introduction of ethynyl group (TCB-621, **33**) the EC50 after curve fitting remained with approx. 0.1 μM the same compared to TCB-494 (**Fig. 3C**). Acetylene group is broadly exploited as a privileged structural feature in medicinal chemistry and can considered more metabolically stable compared to nitro group, when it is not activated for Michael addition and is used to increase stability of some steroids.^36^ Curiously, a double bond did not lead to the same result (**34**). This makes the acetylene group the most interesting structural feature for linker region modifications.

Additional attempts to decrease the EC50 further were not successful. Modifications in **IV** were proven very challenging due to difficulties of synthesis of molecules containing both keto-groups and amino-group, low activity of heterocyclic analogues and cumbersome synthesis of tetrasubstituted benzene rings. Therefore, the main modification tested was methyl group deletion (**35**-**37**) that did not lead to a better potency or efficacy compared to TCB-494 or TCB-621. In addition, 3-arms analogue of TCB-494 also showed increase of EC50 (**38**).

## DISCUSSION

Here, we have identified a tetravalent aromatic guanylhydrazone small molecule, TCB-32, that exhibits remarkable potency and efficacy in stimulating proliferation in NIH 3T3 cells and shows sensitivity to inhibitors targeting FGFR1 and ERK 1/2 signalling pathways. Through a comprehensive exploration of structural variations of all four structural regions of the molecule TCB-32, we were able to optimise the potency through substitutions at position 3 of the aromatic ring in the linker region. This led to the discovery of TCB-494 (**17**), featuring a nitro group, and TCB-621 (**33**), characterised by an acetylene group. In addition, substitution at position 3 with a halogen also increased efficacy without compromising potency as observed for TCB-541 (**22**).

Notably, CNI-1493 (Semapimod) shares its structural motif with our compound series and showed proliferation albeit reduced potency and efficacy in our assay conditions. Initially discovered as an inhibitor of macrophage arginine transport,^41^ it was found to exhibit therapeutic benefits in various clinical settings. Amongst others, it was proposed as treatment for cachexia as inhibitor of TNF ^42^ and suppressor for HIV-1 replication ^43^ and underwent several clinical trials for Crohn’s disease. As a result of a screening pipeline, CNI-1493 was further proposed to be a non-toxic antibiotic that kills metabolically dormant cells.^44^ It would be very interesting to investigate, if our series exhibits the same feature, since they only differ in the linker region compared to CNI-1493. This could either constitute a very useful side effect for cell culture applications or lead to a new series of antibiotics.

In terms of molecular mechanism, it was proposed that CNI-1493 inhibits TLR4 signalling by targeting its chaperone gp96, which in turn leads to the inhibition of downstream targets p38 MAPK and NF-kB.^35^ Several connections between TLR4 and FGFR1 have been reported, e.g., FGFR1 and TLR4 expression was shown to correlate in cancerous tissue leading to regulation of proliferation;^46^ inhibition of FGFR1 in hepatic stellate cells decreased NF-kB activation; and LPS triggered FGFR1 phosphorylation via TLR4.^47^ While the correlation is intriguing, the absence of biochemical data in the literature for CNI-1493 is noticeable and underlines the obstacles to identify a direct interaction due to the inherent chemical characteristics of this chemical scaffold. Therefore, it cannot be ruled out that tetravalent aromatic guanylhydrazones unspecifically aggregate cell surface receptors, thereby leading to their activation (FGFR1) or inhibition (TLR4).

It is therefore of interest to optimise the tetravalent guanylhydrazone scaffold to improve its inherent chemical characteristics. AI/ML models can be used to gain further insights into the formation of aggregates,^38,39^ and were used to evaluate the TCB-32 analogue series.^40^ It was shown that the central aromatic scaffold, the linker and the tripod region of TCB-32, is a strong predictor of the formation of aggregates, which may only be balanced by its substituents. In the future, it would be interesting to test, if there is a possibility to replace all aromatic rings with alternatives less prone for aggregation.

In summary, our data shows that it is possible to replace a growth factor in serum-free medium with a synthetic small molecule. This approach will improve the scalability of cells and will ultimately remove current bottlenecks in cell culture applications like cell therapy or cultivated meat.

### Limitations of the study

This study contains several limitations that are important to consider. First, the action of TCB-32 was only tested in NIH 3T3 and C2C12 using serum-free, and HT-1080 using serum-containing medium. Testing other cell lines and serum-free media is required to determine the universality of our findings. Second, it was beyond the scope of this work to identify the molecular mechanism of the interaction between TCB-32 and FGFR1. Finally, only modifications of the tetravalent guanylhydrazone that are within the patent cited in *Declaration of interests* are presented and discussed to avoid invention disclosure.

### Significance

New applications in cell therapy or cultivated meat require the cultivation of a large volume of cells in bioreactors. Serum-containing medium is superior in producing high cell numbers but shows high batch-to-batch variation, has potential for exogenous contamination and is highly unethical since it is sourced from bovine fetuses as a byproduct of the meatpacking industry. This is not sustainable in the long-term view for food and clinical applications, and serum-free medium is required to meet the growing demands of the industry. Three major obstacles for the wider application of serum-free medium are consistency, costs and scalability, which is mainly the result of protein components that are produced in biological systems. It would therefore be desirable to replace the proteins with chemical compounds that can be produced by an easy scalable manufacturing process. In this study, we present a guanylhydrazone-based class of chemicals to replace protein component bFGF in serum-free medium. The class was found by virtual structure-based screening and exhibits a proliferative effect of up to 70% of bFGF, activates the FGFR1-signalling pathway and binds FGFR1 in a cellular context. In addition, it shows improved thermostability compared to bFGF and thus increases the consistency of serum-free medium. This study provides proof that it is possible to replace protein components in serum-free media with chemical compounds, thereby offering a promising strategy to enable the large-scale production of cells for applications like cell therapy or cultivated meat.

## RESOURCE AVAILABILITY

### Lead contact

Further information should be directed to the lead contact, Hamid Noori

### Materials availability

All the materials generated in this study are available upon request from The Cultivated B GmbH.

### Data and code availability

- All data points to create the graphs are provided in Data S1
- This paper does not report original code
- Any additional information required to reanalyse the data reported in this paper is available from the lead contact upon request.

## STAR*METHODS (STAR★Methods: Cell Press)

### KEY RESOURCES TABLE

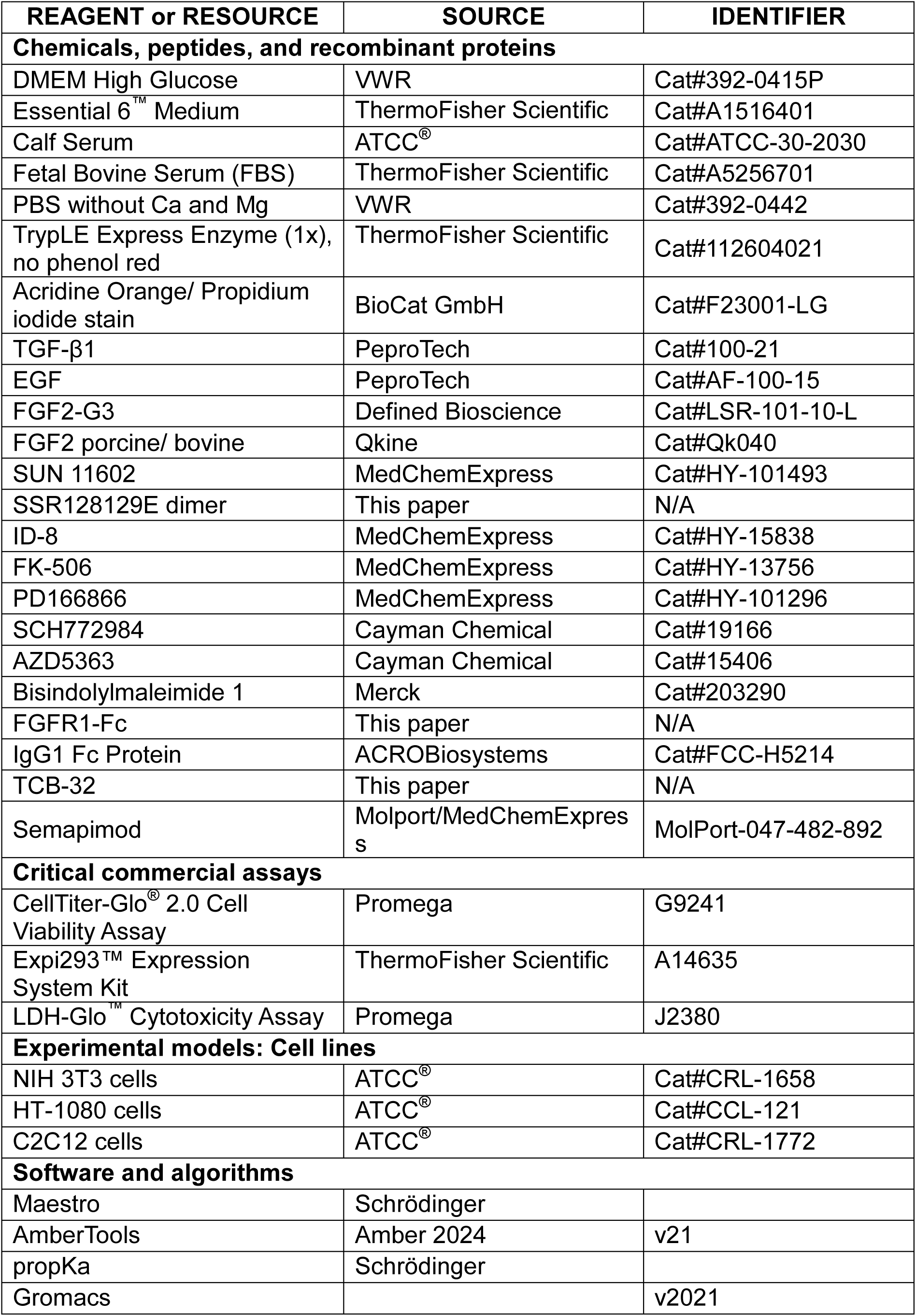
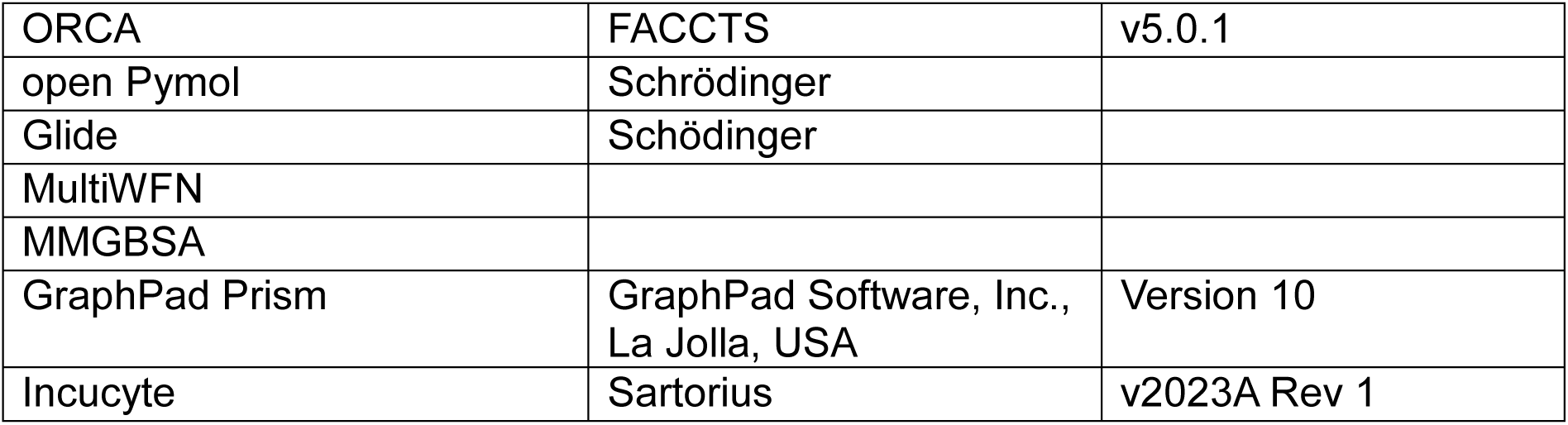

| REAGENT or RESOURCE | SOURCE | IDENTIFIER |
| --- | --- | --- |
| **Chemicals, peptides, and recombinant proteins** |  |  |
| DMEM High Glucose | VWR | Cat#392-0415P |
| Essential 6™ Medium | ThermoFisher Scientific | Cat#A1516401 |
| Calf Serum | ATCC® | Cat#ATCC-30-2030 |
| Fetal Bovine Serum (FBS) | ThermoFisher Scientific | Cat#A5256701 |
| PBS without Ca and Mg | VWR | Cat#392-0442 |
| TrypLE Express Enzyme (1x), no phenol red | ThermoFisher Scientific | Cat#112604021 |
| Acridine Orange/ Propidium iodide stain | BioCat GmbH | Cat#F23001-LG |
| TGF-β1 | PeproTech | Cat#100-21 |
| EGF | PeproTech | Cat#AF-100-15 |
| FGF2-G3 | Defined Bioscience | Cat#LSR-101-10-L |
| FGF2 porcine/ bovine | Qkine | Cat#Qk040 |
| SUN 11602 | MedChemExpress | Cat#HY-101493 |
| SSR128129E dimer | This paper | N/A |
| ID-8 | MedChemExpress | Cat#HY-15838 |
| FK-506 | MedChemExpress | Cat#HY-13756 |
| PD166866 | MedChemExpress | Cat#HY-101296 |
| SCH772984 | Cayman Chemical | Cat#19166 |
| AZD5363 | Cayman Chemical | Cat#15406 |
| Bisindolylmaleimide 1 | Merck | Cat#203290 |
| FGFR1-Fc | This paper | N/A |
| IgG1 Fc Protein | ACROBiosystems | Cat#FCC-H5214 |
| TCB-32 | This paper | N/A |
| Semapimod | Molport/MedChemExpress | MolPort-047-482-892 |
| **Critical commercial assays** |  |  |
| CellTiter-Glo® 2.0 Cell Viability Assay | Promega | G9241 |
| Expi293™ Expression System Kit | ThermoFisher Scientific | A14635 |
| LDH-Glo™ Cytotoxicity Assay | Promega | J2380 |
| **Experimental models: Cell lines** |  |  |
| NIH 3T3 cells | ATCC® | Cat#CRL-1658 |
| HT-1080 cells | ATCC® | Cat#CCL-121 |
| C2C12 cells | ATCC® | Cat#CRL-1772 |
| **Software and algorithms** |  |  |
| Maestro | Schrödinger |  |
| AmberTools | Amber 2024 | v21 |
| propKa | Schrödinger |  |
| Gromacs |  | v2021 |
| ORCA | FACCTS | v5.0.1 |
| open Pymol | Schrödinger |  |
| Glide | Schrödinger |  |
| MultiWFN |  |  |
| MMGBSA |  |  |
| GraphPad Prism | GraphPad Software, Inc.,<br>La Jolla, USA | Version 10 |
| Incucyte | Sartorius | v2023A Rev 1 |

### METHOD DETAILS

#### Structure Analysis

Pig FGFR1 sequence K7GQJ1 (uniprot number) was used for homology modelling, by fitting onto the two human structures PDB ID 1fq9 ^25^ and 1cvs ^26^ using Maestro from the Schrödinger software suite. The topology was created using the tleap module from the AmberTools21 ^51^ using the AMBER forcefield (FF14SB+TIP3P) ^52,53^for protein and water and the compatible GLYCAM (GLYCAM_06j-1) forcefield ^54^ for heparin. Using the acpype program ^55,56^, the topology was translated to Gromacs format. The protonation state of amino acids was predicted using propKa from the Schrödinger software suite.^57,58^ The Na^+^ and Cl^-^ ions were used to achieve a neutral charge and a salt concentration of 0.15M using updated ion parameters.^59^ The system was embedded in a 10 Å layer of TIP3P water molecules leading to a box size of 125 × 125 × 125 Å^3^ and 177 Na^+^ and 179 Cl^-^ ions leading to 169949 atoms.

The system was simulated using Gromacs2021.^60,61^ After a first minimization step using steepest descent method of 1000 steps, up to a convergence criteria of 1000 kJ/mol nm2 of maximum force, the system was gradually heated up to 300 K with an increase of 50 K every 650 ps for a total of 2 ns while keeping the entire system highly restrained (1000 kJ/mol nm2 ) except for the solvent and solute hydrogens. Equilibration started with relaxing the sidechains using an NPT ensemble, scaling the pressure to 1 bar and Berendsen barostat for 1 ns followed by Parrinello–Rahman barostat for an additional 2 ns. In three subsequent 100 ps simulations, restraints on the protein backbone and heparin were gradually decreased (500, 250, 50). Production simulations consist of three replicas for 2 μs each, starting from the same structure with different initial velocities.

Principal component analysis was done using the Amber cpptraj ^62^ module by aligning the three-replica trajectory, considering 10 000 frames for each replica. Clustering analyses were done using DBSCAN and k-means from the Amber cpptraj module ^62^ and the time-lagged t-SNE package ^63^ to select the starting structures for following QM/MM calculations. Interaction energies between amino acid residues of FGF and FGFR were calculated using the ORCA program package ^64,65^, using the local energy decomposition scheme ^66–68^ with HFLD and a def2-TZVP(-f) basis set. Figures and graphical analysis were done using pymol.^69^

#### Virtual Screening

Molecules with a weight over 500MW were downloaded from ZincDB^27^, totalling 9,040,170 substances. 3D structures were prepared using Schrödingers Phase database tool ^70^, leading to 14,157,489 enantiomers and considering the most probable protonation states using PropKa. Molecules were docked on FGFR using Autodock4-GPU ^71^ with default parameters on a box on the orthosteric site with coordinates [105.940, 14.281,136.741] with dimensions [30,24,34]. The top 5% of molecules were redocked using Schödinger’s GlideSP ^72^ given the interactions between bFGF and FGFR1 as input. Required interactions included Arg250, Gln284@NH, Gln284@OE1, and Asp282. Top molecules were chosen for MD simulations of 25 ns each with the same protocol as discussed before. Charges for the molecule were calculated using standard Hartree-Fock, 6-31G* calculation using the ORCA quantum chemistry package ^64,65^ in combination with MultiWFN.^73^ To avoid distortion of the D3 domains, a 500 kJ/mol distance constraint was applied. Binding free energies were calculated using the MM-GBSA ^74,75^ method on 100 frames of the equilibrated parts of the trajectories, using GB-Neck2 ^76^ solvation model.

The molecules with very low solubility in water (clogS lower than -10), known DNA- intercalators, warheads/precursors for cell dyes and molecules with errors in the structures in database entry were filtered. While 3 commercially available molecules were purchased, the synthetic accessibility of the remaining molecules was determined according to the following four criteria: easy difficulty: 1-3 steps; medium difficulty: 3-6 steps; hard difficulty: 6+ steps; no suggested synthesis due to unknown synthesis of building blocks. A total of 9 molecules with easy and medium difficulty entered the synthesis, but only four of them were successfully synthesized. This means a total of 7 compounds was tested in the proliferation assay.

#### Optimization

To gain further insights into the formation of aggregates, AI/ML models were used to to evaluate the TCB-32 analogue series.^50^

#### Compounds & proteins

Growth factors TGF-β1 (PeproTech, #100-21), EGF (PeproTech, #AF-100-15), FGF2- G3 (Defined Bioscience, #LSR-101-10-L), and porcine/ bovine FGF2 wildtype (Qkine, #Qk040) were resuspended in PBS and 5% Trehalose to a concentration of 100 μg/ mL.

Compounds SUN 11602 (MCE, #HY-101493, dimeric SSR12129E (synthesised in-house), ID-8 (MCE, #HY-15838), FK-506 (MCE, HY-13756), PD166866 (MCE, #HY- 101296), SCH772984 (Cayman, #19166), AZD5363 (Cayman, #Cay15406) and Bisindolylmaleimide 1 (Merck, #203290) were prepared at 10 mM in DMSO.

The extracellular domain of porcine FGFR1 (aa 1-374) - tagged C-terminally with the Fc domain of human IgG1 and followed by a non-biotinylated Avi-Tag - was expressed using Gibco’s Expi293™ expression system (ThermoFisher) according to the Expi293™ user guide. The supernatant was further purified by Protein A chromatography using a HiTrap® MabSelect Sure column (Cytiva) on an Äkta Pure™ 25 (Cytiva). The protein was eluted with elution buffer (20 mM MES, 3.6 M MgCl2, pH= 6.6) and buffer exchanged into TBS buffer (pH= 8.0) with either a Vivaspin20 10 kDa concentrator (Sartorius) or an Amicon® Ultra-15 10 kDa concentrator (Merck). A second buffer exchange into the final buffer (PBS, 10% Glycerol, pH= 7.4) was performed with an Amicon® Ultra-15 10 kDa concentrator (Merck). The concentration of the protein was determined by UV/ Vis spectrometry and its quality checked by reducing and non-reducing SDS-PAGE. The protein was aliquoted, flash frozen in liquid nitrogen and stored at -80°C. As negative control, the Fc domain of human IgG1 (ACROBiosystems) was used.

#### Proliferation Assay

NIH 3T3 cells (ATCC) were grown in DMEM High Glucose (VWR) supplemented with 10% Calf Serum (CS, VWR) in a T75-flask. At 80-90% confluency, medium was aspirated, cells were washed with 10 mL PBS (VWR) and detached with 2 mL TrypLE Express (ThermoFisher). Cells were resuspended in 8 mL of Essential 6 medium (ThermoFisher) without bFGF and TGF-β1 and counted with Acridine Orange/Propidium iodide stain(Biocat) using the LUNA-FL Dual Fluorescence Cell Counter (Logos). Cells were diluted to 9.000 cells/ mL with Essential 6 medium (ThermoFisher). 50 μL of the cell dilution was dispensed to wells B2 to O23 of a collagen-coated white 384-well plate (Greiner) using a 24-channel pipette. The outer wells were filled with 50 μL PBS. The plate was centrifuged for 30 s at 300 x g and incubated at 37°C and 5% CO2.

After 2-3 hours, the compound of interest was added to the well in varying concentrations, a maximum total volume of 0.2 μL (0.4% of total volume) and in a randomised manner using a phyton-based script developed in-house with the I-DOT non-contact liquid handler (Dispendix). The plate was centrifuged for 30 s at 300 x g and incubated at 37°C and 5% CO2.

For live-cell imaging, the plate was imaged every 4 hours in the Incucyte S3 (Sartorius). The confluency was analyzed using ―AI confluence‖ as segmentation method. Data was normalized to the first scan at time 0 and displayed as ratio. Data is based on 4 replicated shown as SEM. The endpoint was measured after 64-70 hours by adding 30 μL of CellTiter-Glo^®^ 2.0 reagent (Promega) to each cell-containing well. The plate was shaken for 2 min at 500 rpm and then incubated for 10 min at 22°C. Luminescence was recorded using the SpectraMax i3x Multimode microplate reader (Molecular Devices). Data was normalized by using the negative control (= DMSO) as 0, and the positive control (= 100 ng/ mL FGF2-G3) as 1. For plate-to-plate comparison, the z’ factor was calculated as z’ = 1 – ((3* (SDPos – SDNeg))/ (Pos-Neg)), and the signal to noise ratio was determined as the ratio of the positive and negative control. Data is based on 4 replicates shown as SD, if outliers were removed this is mentioned in the figure legend. Data was fitted to the equation: Y= Bottom + (X^Hillslope)*(Top-Bottom)/ (X^Hillslope + EC50^Hillslope). To rank the various TCB-32 analogues, each plate contained the complete dose-response of TCB-32 for normalisation.

To rank molecules according to their potency, we determined the concentration when the sample reached its maximum activity and used the following classification: A ≤ 1 µM; B > 1 and ≤ 1.4 µM; C > 1.4 and ≤ 4.5 µM; D > 4.5 and ≤ 10 µM; E = not activity. For efficacy, we used the top value normalised to TCB-32 and used the following classification: 1 ≥ 0.6; 2 < 0.6 and ≥ 0.4; 3 < 0.4 and ≥ 0.2; 4 < 0.2 and ≥ 0.05; 5 < 0.05. All graphs were generated using Prism (GraphPad).

#### LDH-Glo^™^ Cytotoxicity assay

HT-1080 cells (ATCC) were grown in DMEM High Glucose supplemented with 10% FBS (ThermoFisher) in a T75-flask. At 80-90% confluency, medium was aspirated, cells were washed with 10 mL PBS and detached with 2 mL TrypLE Express. Cells were resuspended in 8 mL of standard growth medium and counted with Acridine Orange/ Propidium iodide stain using the LUNA-FL Dual Fluorescence Cell Counter. Cells were diluted to 24.000 cells/ mL in standard growth medium with 5% FBS. 50 μL of the cell dilution was dispensed to wells B2 to O23 of a white 384-well plate using a 24-channel pipette. The outer wells were filled with 50 μL PBS. The plate was centrifuged for 30 s at 300 x g and incubated at 37°C and 5% CO2.

After 2-3 hours, TCB-32 was added to the well in varying concentrations and a maximum total volume of 0.2 μL (0.4% of total volume) DMSO using the I-DOT non-contact liquid handler. The plate was centrifuged for 30 s at 300 x g and incubated at 37°C and 5% CO2.

Reagents of the LDH-Glo™ Cytotoxicity assay (Promega) were prepared according to the manufacturer’s advice. The LDH reaction reagent was prepared by mixing LDH Detection Enzyme Mix and Reductase Substrate at a ratio of 200 to 1. The maximum LDH release sample was prepared by adding 1 μL of Triton X-100 to 50 μL of Vehicle-Only Cells for 15 minutes before collecting the samples.

After 24 hours, 2 μL of the cell culture medium was diluted in 8 μL of LDH Storage Buffer (200 mM Tris-HCl (pH 7.3), 10% Glycerol, 15% BSA) in a white 384-well plate. 10 μL of the LDH reaction reagent was then added to the samples and incubated for 60 minutes at room temperature. Luminescence was recorded using the SpectraMax i3x Multimode microplate reader. Viability of the cells was determined using CellTiter- Glo^®^ 2.0 reagent as described for the proliferation assay. Graphs were generated using Prism. Cytotoxicity was calculated as: Cytotoxicity = (Experimental LDH Release – Medium Background)/ (Maximum LDH Release Control – Medium Background). Viability was calculated as: Viability = Treated sample/ DMSO Control. Data is based on 4 replicates shown as SD.

#### Synthesis of TCB-32 and analogues

All chemicals and solvents were purchased in reagent grade from commercial suppliers (Acros®, SigmaAldrich®, Merck®, ChemPur®, Chemspace®, ABCR®, VWR®, Roth®, Thermo Fischer Scientific®) and used as received unless otherwise specified.

Reaction progress was monitored using thin-layer chromatography (TLC), or ultra-high performance liquid chromatography-mass spectrometry (UHPLC-MS). TLC was performed on pre-coated silica gel plates with a fluorescence indicator (254 nm excitation wavelength) and visualized under UV light. UHPLC-MS data was acquired on an Agilent Infinity II 1260 instrument with a 1260 Infinity II liquid autosampler, Poroshell 120 EC-C18 columns, acetonitrile/water gradients, and formic acid modifier. The column eluate was analysed using a 1260 Infinity II Diode Array Detector WR scanning from 200 to 400 nm and LC/MSD iQ mass spectrometer scanning in both positive and negative ion modes from 140 to 1000 Da.

Typical elution system, if not specified: A/B gradient (0-2.5 min 100% A ,2.5-9 min linear gradient from 100% A to 100% of B ,9-10 min linear gradient from 100% of B to 100% of A, A= 98/2 water/acetonitrile +0.1% formic acid, B= 2/98 water/acetonitrile + 0.1% formic acid).

Normal phase flash column chromatography was performed on automated flash chromatography (FC) using Pure C-810 Buchi instrument and Silica 60 A cartridges.

Reverse-phase chromatography was performed on automated flash chromatography (FC) using Pure C-810 Buchi instrument and pre-packed BGB Scorpius C18-HP 100 A cartriges or HPLC Agilent Infinity II Preparative HPLC using InfinityLab ZORBAX Eclipse Plus C18 column. UV detection was used to trigger fraction collection.

Nuclear magnetic resonance spectroscopy (NMR) chemical shifts are given in parts per million downfield from tetramethylsilane and were recorded Bruker spectrometers operaring on 300 or 400 MHz for ^1^H NMR and 75 MHz or 100 MHz for ^13^C NMR. Chemical shifts are expressed in parts per million (ppm, δ) referenced to the solvent residual peaks (^1^H: CHCl_3_, 7.26 ppm; DMSO-d5, 2.50 ppm; ^13^C: CDCl_3,_ 77.00 ppm; DMSO-d6, 39.52 ppm). The peak shapes are described as follows: s, singlet; d, doublet; t, triplet; q, quartet; quin, quintet; m, multiplet; br s, broad singlet. Coupling constants were assigned as observed. The obtained spectra were evaluated with the program MestReNova Mass spectrometry data is reported from UHPLC-MS analyses. Mass spectrometry (MS) was performed using electrospray ionization (ESI).

## QUANTIFICATION AND STATISTICAL ANALYSIS

Data are presented as mean ± standard deviation (SD), except for data based on images that is presented as mean ± standard error of mean (SEM). The n values indicated in the figures refer to biological replicates. Statistical analysis was conducted using GraphPad Prism 10 and data was fitted to the equation: Y= Bottom + (X^Hillslope)*(Top-Bottom)/ (X^Hillslope + EC50^Hillslope). For plate-to-plate comparison, the z’ factor was calculated as z’= 1 – ((3* (SDPos – SDNeg))/ (Pos- Neg)).

## SUPPLEMENTAL INFORMATION

Document S1. Figures S1 and S2

Document S2. Synthesis of all described small molecules

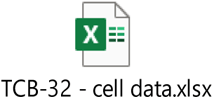

Document S3. Table with raw data from all cellular experiments

## Supporting information

Synthesis of all described small molecules

## ACKNOWLEDGMENTS

The authors wish to thank Nicolai Bluthardt for his support in the purification of FGFR1-Fc. Study was funded by The Cultivated B GmbH.

## AUTHOR CONTRIBUTIONS

Conceptualisation H.N., S.L., A.S., M.F., M.Z., G.D., C.S., M.B., K.K., C.H. and J.P.; supervision H.N. and S.L.; methodology M.F., G.D., A.S., T.D., K.K., C.S. and F.H.; software A.S., C.S. and T.D.; investigation M.F., G.D., A.S., K.K., C.S., M.B., F.H. and M.S.; Resources C.H. and M.B.; writing – original draft G.D., M.F., and C.S.; writing – review and editing, M.F., G.D. and H.N.

## DECLRATION OF INTERESTS

H. Noori is the manager of The Cultivated B. GmbH.

The authors wish to disclose pending patent application PCT/EP2024/075415 related to this work. H.N., S.L., A.S., M.B., M.F., M.Z., G.D., K.K., C.H. and F.H. are inventors on this patent.

