## Supplementary material for "Discovery and optimization of a guanylhydrazone-based small molecule to replace bFGF for cell culture applications": Figures S1 and S2

### Supplementary data

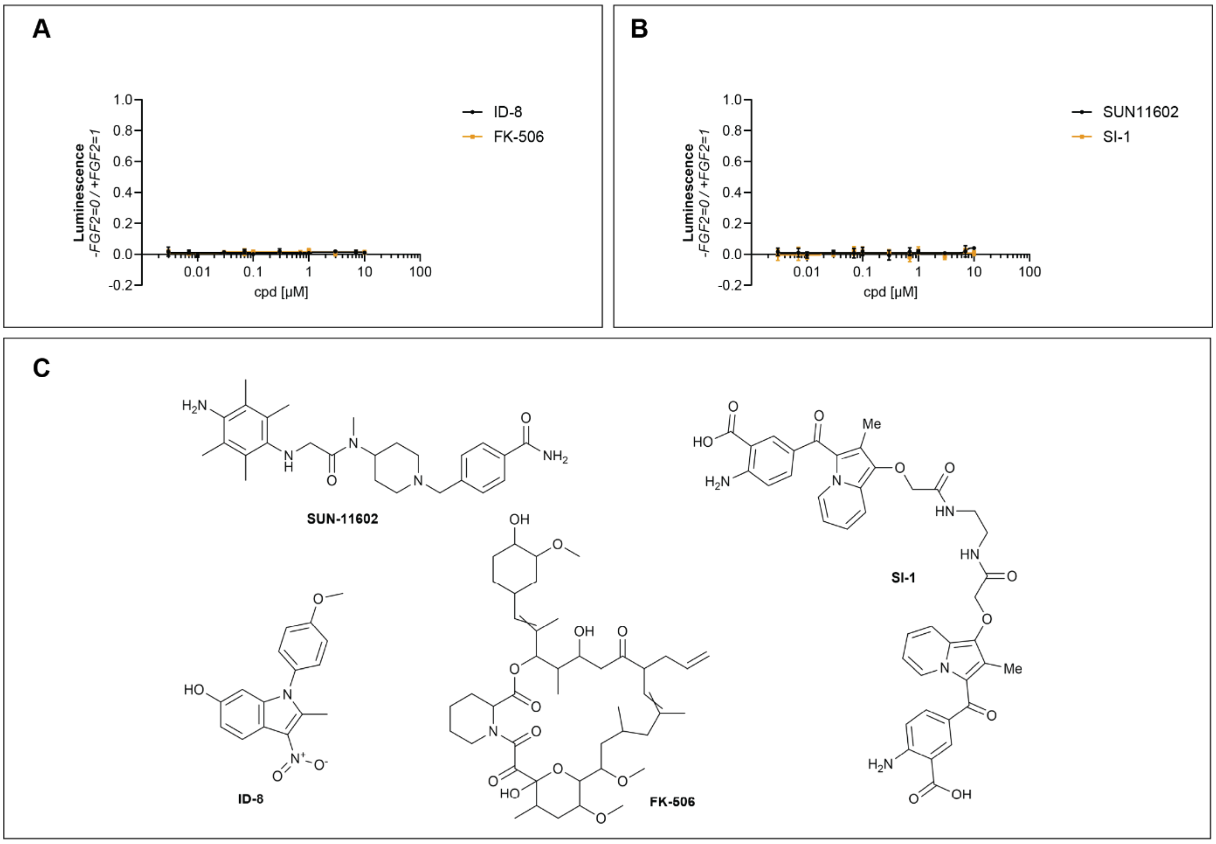


### Figure S1

Proliferation assay of NIH 3T3 cells stimulated with (**A**) ID-8 and FK-506 (n= 4, SD, z'= 0.86), (**B**) SUN 11602 and SI-1 (n= 4, SD, z'= 0.65). (**C**) The chemical structures of the potential FGFR1 pathway activators from the literature.

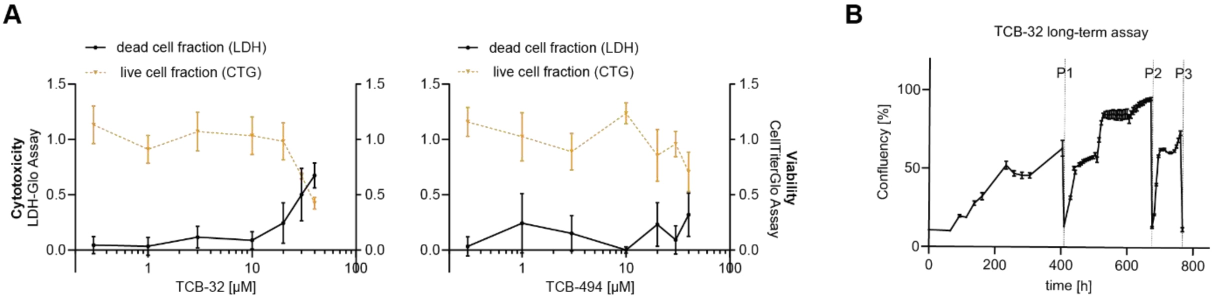


### Figure S2

(**A**) LDH-Glo™ cytotoxicity (black curve, left y-axis) and CellTiter-Glo® 2.0 cell viability assay (orange curve, right y-axis) in HT-1080 cells using various concentrations of TCB-32 (left) and TCB-494 (right) (n= 4, SD, row I was removed as outlier since LDH reaction reagent was not added).

(**B**) Cultivation of C2C12 cells over 3 passages in Essential 6 with 1 µM TCB-32 using live-cell imaging (n= 3, SEM).
