## Supplementary material for "Discovery and optimization of a guanylhydrazone-based small molecule to replace bFGF for cell culture applications": Synthesis of all described small molecules

### Synthesis of building blocks.

#### Synthesis of 3,5-diacetylaniline.

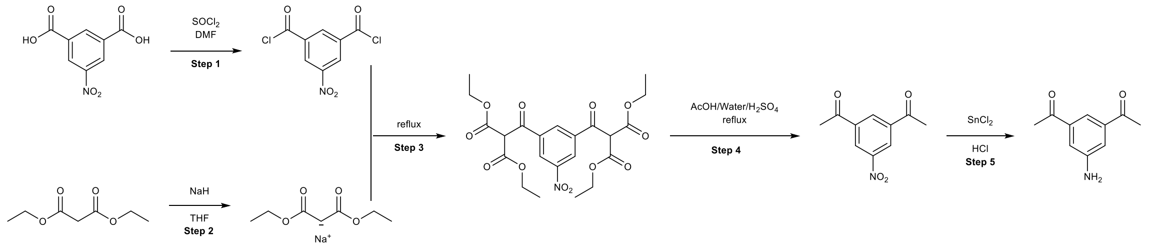


The synthesis was conducted according to [1].

**Steps 1-4. Synthesis of 3,5-diacetylnitrobenzene.**

To a suspension of 5-nitroisophthalic acid (10 g, 60 mmol) in 20 mL of thionyl chloride, 5 drops of dry DMF were added and the reaction mixture was refluxed for 2 h giving a pale-yellow solution. The solvent was removed in vacuo yielding a yellow solid, which was used without further purification for the next step.

Diethyl malonate (17.5 ml, 0.11 mmol) was dissolved in 225 mL of THF, 60% sodium hydride in mineral oil (5 g, 0.12 mol) of was added portion wise and the mixture was heated for 3 h under reflux. To this solution 5-Nitroisophthalic acid dichloride acquired in the previous step was dissolved in 40 ml of THF and was added dropwise. After complete addition, the solution was refluxed overnight, and then the solvent was evaporated under reduced pressure. The brown slurry was dissolved in 100 mL of DCM and 100 ml of water, separated and aqueous layers were extracted with DCM (2*100 ml). DCM was removed in vacuo yielding a yellow oil. The oil was dissolved in a mixture of glacial acetic acid/conc. sulfuric acid/water (100 mL: 5 mL: 20 mL) and refluxed overnight. After that, 250 mL water was added, and the mixture was extracted with DCM (3*200 mL). The organic layers combined, the solvent was removed in vacuo and the resulting residue was purified by flash chromatography on silica gel eluting with 0-60% Cyclohexane-EtOAc obtaining the desired product as a slightly yellow solid in 25% yield (2.5 g).

MS $(M+H)^+$: 208.4

$^1$H NMR (300 MHz, $CDCl_3$) δ 8.87 (d, J = 1.6 Hz, 2H), 8.74 (t, J = 1.6 Hz, 1H), 2.68 (s, 6H).

$^{13}$C NMR (75 MHz, $CDCl_3$) δ 195.0, 148.8, 138.8, 132.7, 126.6, 26.8.

**Step 5. Synthesis of 3,5-diacetylaminobenzene.**

In a round-bottom flask tin (II)chloride (10.2 g, 45 mmol) was dissolved in 33 of mL conc. HCl and warmed up to 50 °C followed by the slow addition of 2.5 g of product of the step 1. Afterwards, the solution was stirred for further 15 minutes and then poured, under gas formation, in a mixture of 55 g potassium carbonate and 200 mL ice/water. The product was extracted with EtOAc and the combined organic layers were dried over sodium sulfate. Finally, the solvent was removed in vacuo obtaining the product as a bright yellow solid in 54% yield (1.2 g).

MS $(M+H)^+$: 177.9.

$^1$H NMR (300 MHz, $CDCl_3$) 7.63 (s, 1H), 7.36 (s, 2H), 5.63 (s, 2H), 2.55 (s, 6H).

$^{13}$C NMR (75 MHz, $CDCl_3$) 197.9, 149.4, 137.9, 117.0, 115.5, 26.8.

**Synthesis of 1-(3-amino-5-ethylphenyl)ethan-1-one**.

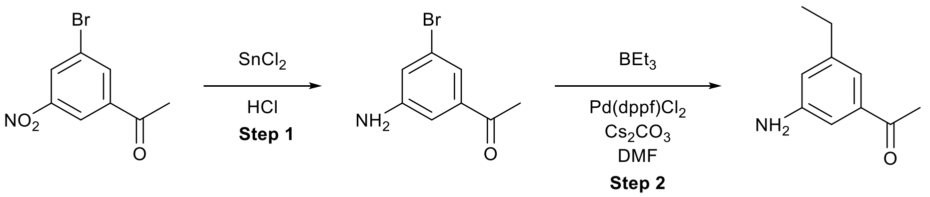


**Step 1**. Synthesis of 1-(3-amino-5-bromophenyl)ethan-1-one.

In a round-bottom flask tin(II)chloride (2.1 g, 10 mmol) was dissolved in 6 of mL conc. HCl and warmed up to 50 °C followed by the slow addition of 1-(3-bromo-5-nitrophenyl)ethenone (500 mg, 2.04 mmol). Afterwards, the solution was stirred for further 15 min and then poured, under gas formation, in a mixture of 11 g potassium carbonate and 40 mL ice/water. The product was extracted with EtOAc and the combined organic layers were dried over sodium sulfate. Finally, the solvent was removed in vacuo obtaining a bright yellow solid in 85% yield (340 mg).
$^1$H NMR (300 MHz, $CDCl_3$) δ 7.34 (s, 1H), 7.08 (s, 1H), 6.93 (s, 1H), 3.62 (s, 2H), 2.46 (s, 4H).
$^{13}$C NMR (75 MHz, $CDCl_3$) δ 197.1, 153.2, 148.0, 139.5, 123.3, 121.8, 121.5, 112.9, 26.7.

**Step 2**. Synthesis of 1-(3-amino-5-ethylphenyl)ethan-1-one.
The synthesis was adopted from [2] for intermediate 9.
In a three-necked flask, product from step 1 (330 mg, 1.6 mmol) and triethylborane (1 M in THF, 2.3 ml), $Cs_2CO_3$ (2.7 g. 6.3 mmol) and 13 ml of DMF were placed. The mixture was evacuated and flushed with nitrogen three times and $PddppfCl_2$ (30 mg, 0.04 mmol) was added. The reaction was stirred overnight at 70 °C. After the end of the reaction, it was cooled to room temperature and diluted with 15 ml of EtOAc. The organic layer was washed with water, separated and concentrated under reduced pressure. The residue was purified by silica gel column chromatography eluting with cyclohexane to cyclohexane:ethylacetate 40:60 obtaining 1-(3-amino-5-ethylphenyl)ethan-1-one as a light orange solid in 46% yield (120 mg).
MS $(M+H)^+$: 163.9.
$^1$H NMR (300 MHz, $CDCl_3$) δ 7.18 (s, 1H), 7.11 (s, 1H), 6.76 (s, 1H), 3.68 (s, 2H), 2.60 (q, $J$ = 7.6 Hz, 2H), 2.54 (s, 3H), 1.22 (t, $J$ = 7.5 Hz, 3H).
$^{13}$C NMR (75 MHz, $CDCl_3$) δ 198.6, 146.1, 145.9, 138.3, 124.7, 119.7, 118.8, 112.3, 31.5, 26.8, 15.4.

**Synthesis of 3,5-diacetylphenol**.

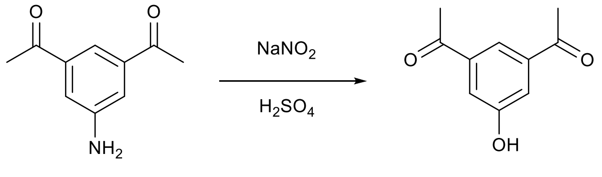


The synthesis was conducted according to [3].
To a solution of 3,5-diacetylaniline (0.3 g, 1.5 mmol) in 3% (v/v) $H_2SO_4$ (0.15 ml of acid in 4.85 mL water) at 0 °C was added $NaNO_2$ (0.13 g, 1.83 mmol). After 30 min at 0 °C urea (50 mg) was added. The mixture was warmed to room temperature and then was heated to 80 °C over 30 min. The mixture was cooled and extracted with EtOAc. The organic solvents were evaporated under reduced pressure obtaining 3,5-diacetylphenol as red solid in 80% yield (210 mg).

MS $(M+H)^+$: 178.7.
$^1H$ NMR (300 MHz, $CDCl_3$) δ 8.01 (s, 1H), 7.63 (s, 1H), 2.58 (s, 5H).
$^{13}C$ NMR (75 MHz, $CDCl_3$) δ 197.7, 156.8, 138.8, 120.7, 119.6, 26.8.

**Synthesis of 3,5-diacetylbromobenzene**.

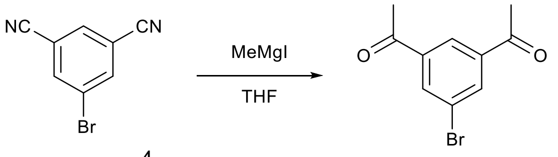


The synthesis was adopted from [4].

To the solution of 5-bromoisophthalonitrile (10 mmol, 2.07 g) in dry THF (100 mL) was added dropwise a solution of methylmagnesium iodide (3 M solution in $Et_2O$, 40 mmol, 13 ml) at 0 °C. The reaction mixture was stirred under nitrogen atmosphere at rt overnight. 150 mL of water and 150 mL of 3 M HCl were added, and the reaction mixture was extracted with ethyl acetate (3 x 200 mL). The organic extracts were combined and the solvents were evaporated under reduced pressure. The residue was purified by silica gel column chromatography eluting with cyclohexane to cyclohexane:ethylacetate 40:60 obtaining 3,5-diacetylbromobenzene as a light orange solid in 50% yield (1.2 g).
MS $(M+H)^+$: 242.5.
$^1H$ NMR (300 MHz, $CDCl_3$) δ 8.34 (s, 1H), 8.19 (s, 2H), 2.58 (s, 6H).
$^{13}C$ NMR (75 MHz, $CDCl_3$) δ 195.9, 139.0, 135.4, 126.5, 123.5, 26.7.

**Synthesis of 3,5-diacetylbenzoic acid**.

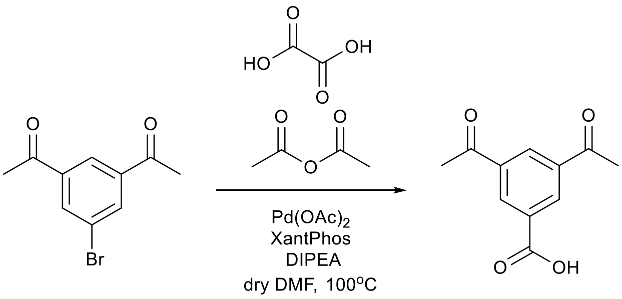


The synthesis was adopted from [5].

A 50 mL Schlenk tube equipped with a stir bar was charged with oxalic acid dihydrate (570 mg, 6.3 mmol), palladium (II) acetate (9.4 mg, 0.042 mg), Xantphos (24.3 mg, 0.042 mmol), intermediate 50 (500 mg, 2.1 mmol), acetic anhydride (0.6 ml, 6.3 mmol), N,N-diisopropylethylamine (1.08 ml, 6.3 mmol) and N,N-dimethylformamide (10 ml) under air. The tube was evacuated and backfilled with nitrogen (5 times). The mixture was stirred in a preheated oil bath (100 °C) for 21h. The reaction mixture was diluted with ethyl acetate (10 mL), acidified with 2 M hydrochloric acid, and washed with brine. The organic phase was dried over anhydrous $MgSO_4$ and concentrated in vacuo. The residue was purified by silica gel column chromatography eluting with cyclohexane to cyclohexane:ethylacetate 50:50 to obtain the desired product as a yellow solid in 60% yield (260 mg).
MS $(M+H)^+$: 206.8.
$^1H$ NMR (300 MHz, DMSO-$d_6$) δ 8.63 (s, 3H), 2.70 (s, 6H)
$^{13}C$ NMR (75 MHz, DMSO-$d_6$) δ 197.0, 166.1, 137.6, 132.5, 132.0, 131.3, 27.0.

**Synthesis of 5-(difluoromethyl)isophthalic acid.**

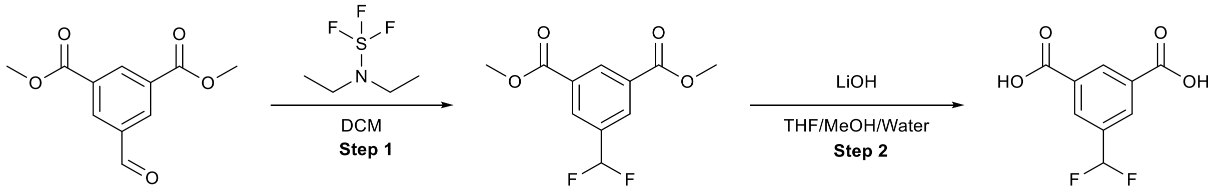


The synthesis was adopted from [5].

**Step 1**: Synthesis of dimethyl 5-(difluoromethyl)isophthalate.

To a solution of 1,3-dimethyl 5-formylbenzene-1,3-dicarboxylate (444 mg, 2 mmol) in dichloromethane (13 ml) was added (diethylamino)sulphur trifluoride (0.5 ml, 38.6 mmol) and the reaction mixture was stirred at room temperature for 18 h. To the mixture was added additional dichloromethane and saturated aqueous sodium hydrogen carbonate solution and the two layers were separated. The solvent was evaporated, and the residue was purified by silica gel column chromatography eluting with cyclohexane to cyclohexane:ethylacetate 60:40 to obtain the desired product.

MS $(M+H)^+$: 244.9.

$^1$H NMR (300 MHz, $CDCl_3$) δ 8.71 (s, 1H), 8.29 (s, 2H), 6.66 (t, $J$ = 56.0 Hz, 1H), 3.90 (s, 6H).

$^{19}$F NMR (283 MHz, $CDCl_3$) δ -111.77 (s, 2F).

$^{13}$C NMR (75 MHz, $CDCl_3$) δ 165.3, 135.4 (t, $J$ = 23.3 Hz), 132.7 (t, $J$ = 1.6 Hz), 131.4, 130.9 (t, $J$ = 5.9 Hz), 113.5 (t, $J$ = 240.4 Hz), 52.6.

**Step 2:** Synthesis of 5-(difluoromethyl)isophthalic acid.

The product from the previous step was dissolved in a mixture of THF/MeOH/Water (1 ml/1 ml/1 ml) and LiOH (400 mg) was added. The resulting mixture was stirred overnight. After the completion of the reaction, 1M HCl was added (5 ml) and the mixture was extracted with EtOAc. The organic layers were combined, dried over the sodium sulfate and evaporated under the reduced pressure yielding the product as white solid.

Yield, over two steps: 370 mg (89%).

MS $(M+H)^+$: 216.9.

$^1$H NMR (300 MHz, DMSO-$d_6$) δ 13.52 (s, 1H), 8.59 (s, 2H), 8.33 (s, 2H), 7.23 (t, $J$ = 55.4 Hz, 1H).

$^{19}$F NMR (283 MHz, DMSO-$d_6$) δ -106.19 (s, 2F).

$^{13}$C NMR (75 MHz, DMSO-$d_6$) δ 165.7, 135.1 (t, $J$ = 30.3 Hz), 132.1, 132.0, 130.3 (t, $J$ = 6.0 Hz), 113.9 (t, $J$ = 240.4 Hz).

**Synthesis of 5-(trifluoromethoxy)isophthalic acid.**

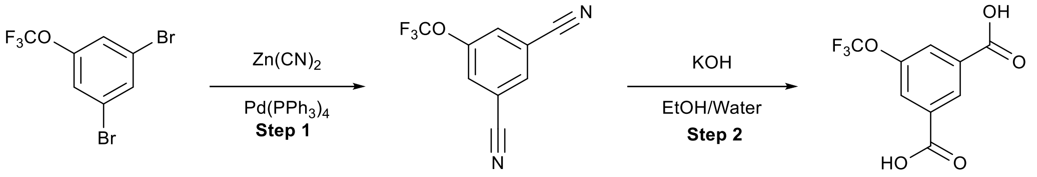


The synthesis was conducted according to [6].

**Step 1**: Synthesis of 5-(trifluoromethoxy)isophthalonitrile.

A 25 mL round bottom flask equipped with a magnetic stir bar and a condenser was charged with the 1,3-dibromo-5-(trifluoromethoxy)benzene (0.6g, 1.9 mmol) and DMF (4 ml). The reaction mixture was degassed using nitrogen gas. $Pd(PPh_3)_4$ (0.65 g, 0.56 mmol) and $Zn(CN)_2$ (0.24 g, 2.06 mmol) were added afterwards. The mixture

was stirred at 85 °C overnight. The solvent was evaporated under the reduced pressure, and the residue was purified by silica gel column chromatography eluting with cyclohexane to cyclohexane:ethylacetate 50:50 to obtain the desired product.
Mass was not observed in ESI.
$^1$H NMR (300 MHz, $CDCl_3$) δ 7.91 (t, *J* = 1.4 Hz, 1H), 7.76 (s, 1H).
$^{19}$F NMR (283 MHz, $CDCl_3$) δ -58.06 (s, 3F).
$^{13}$C NMR (75 MHz, $CDCl_3$) δ 149.6, 147.5, 133.5, 128.3, 120.0 (q, *J* = 268.0 Hz), 116.0, 115.2.

**Step 2**: Synthesis of 5-(trifluoromethoxy)isophthalic acid.
The product from the previous step in EtOH (6 ml) was treated with 1N KOH (6 ml, 336 mg of KOH) and refluxed at 80 °C overnight. After cooling to room temperature the volatiles were removed via rotary evaporation. The mixture was adjusted to pH=l-2 with concentrated HCl and the mixture was extracted with EtOAc. The combined organics were dried over $Na_2SO_4$. The organic layers were combined, dried over the sodium sulfate and evaporated under the reduced pressure yielding the product as white solid.
Yield, over two steps: 205 mg (41%).
MS $(M-H)^-$: 248.8.
$^1$H NMR (300 MHz, DMSO-$d_6$) δ 13.41 (s, 2H), 8.45 (t, *J* = 1.5 Hz, 1H), 8.03 (s, 2H).
$^{19}$F NMR (283 MHz, DMSO-$d_6$) δ -57.05 (s, 1F).
$^{13}$C NMR (75 MHz, DMSO-$d_6$) δ 165.2, 148.4, 133.7, 128.6, 125.4, 119.9 (d, *J* = 257.8 Hz).

**Synthesis of 5-cyanoisophthalic acid.**

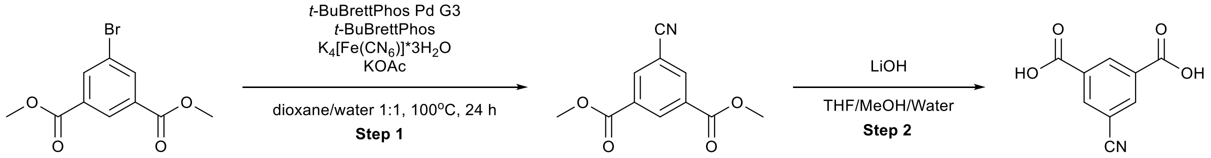


The synthesis was adopted from [7].
**Step 1**: Synthesis of dimethyl 5-cyanoisophthalate.
To a screw-top test tube equipped with a magnetic stir bar was added *t*-BuBrettPhos Pd G3 (43 mg, 0.05 mmol), *t*-BuBrettPhos (24 mg, 0.05 mmol), $K_4[Fe(CN)_6]*3H_2O$ (184 mg, 0.5 mmol) and dimethyl 5-bromoisophthalate (273 mg, 1 mmol). After sealing with a Teflon-lined screw-cap septum, the vessel was evacuated and refilled with nitrogen (three cycles). Degassed dioxane (2.5 mL), KOAc (12 mg, 0.125 mmol) and water (2.5 mL) were then added to the reaction tube via syringe. The test tube was placed in an oil bath preheated to 100 °C. After 4 h of stirring at 100 °C, the reaction mixture was then cooled to room temperature. The contents of the test tube were transferred to a separation funnel using EtOAc (15 mL) and brine (15 mL), and the organic layer was separated from the aqueous layer. The solvents were evaporated and the residue was purified by chromatography on silica gel eluting with cyclohexane to cyclohexane:ethylacetate 50:50 to obtain the product in 72% yield (158 mg).
MS $(M+H)^+$: 210.5.
$^1$H NMR (300 MHz, $CDCl_3$) δ 8.87 (s, 1H), 8.48 (d, *J* = 1.6 Hz, 2H), 3.99 (s, 6H).
$^{13}$C NMR (75 MHz, $CDCl_3$) δ 164.4, 136.9, 134.5, 132.2, 117.1, 113.7, 53.1.

**Step 2**: Synthesis of 5-cyanoisophthalic acid.

The product from the previous step was dissolved in mixture of THF/MeOH/Water (1 ml/1 ml/1 ml) and LiOH (200 mg) was added. The resulting mixture was stirred overnight. After the completion of the reaction, 1 M HCl was added (5 ml) and the mixture was extracted with EtOAc. The organic layers were combined, dried over sodium sulfate and evaporated under the reduced pressure obtaining the product as white solid in 91% yield (125 mg).
MS $(M-H)^-$: 189.9.
$^1$H NMR (300 MHz, DMSO-$d_6$) δ 13.65 (s, 1H), 8.65 (s, 1H), 8.51 (d, *J* = 1.6 Hz, 2H).
$^{13}$C NMR (75 MHz, DMSO-$d_6$) δ 165.0, 136.5, 133.7, 132.6, 117.3, 112.7.

**Synthesis of 5-sulfamoylisophthalic acid.**

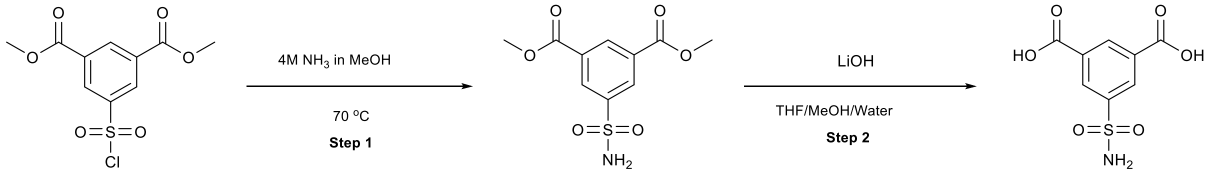


**Step 1**: Synthesis of dimethyl 5-sulfamoylisophthalate.
1,3-dimethyl 5-(chlorosulfonyl)benzene-1,3-dicarboxylate (200 mg, 0.69 mmol) was dissolved in 3 ml of MeOH and 2 ml of 4M ammonia in methanol were added. The mixture was stirred in the microwawe vial with a closed cap at 70 °C overnight. After the completion of the reaction the solvents were evaporated under the reduced pressure to afford the desired product as white solid.
MS $(M-H)^-$: 271.9.
$^1$H NMR (300 MHz, $CDCl_3$) δ 8.73 – 8.65 (m, 3H), 3.90 (s, 6H).
$^{13}$C NMR (75 MHz, $CDCl_3$) δ 164.8, 145.0, 133.1, 131.3, 131.1, 130.5, 52.6.

**Step 2**: Synthesis of 5-sulfamoylisophthalic acid.
The product from the previous step was dissolved in mixture of THF/MeOH/Water (1 ml/1 ml/1 ml) and LiOH (400 mg) was added. The resulting mixture was stirred overnight. After the completion of the reaction, 1M HCl was added (5 ml) and the mixture was extracted with EtOAc. The organic layers were combined, dried over the sodium sulfate and evaporated under the reduced pressure yielding the product as white solid.
Yield, over two steps: 105 mg (63%).
MS $(M+H)^+$: 245.8.
$^1$H NMR (300 MHz, DMSO-$d_6$) δ 13.71 (s, 2H), 8.60 (t, *J* = 1.6 Hz, 1H), 8.56 (d, *J* = 1.6 Hz, 2H), 7.64 (s, 2H).
$^{13}$C NMR (75 MHz, DMSO-$d_6$) δ 165.4, 145.2, 132.5, 132.3, 130.1.

**Synthesis of 5-(methylsulfonyl)isophthalic acid.**

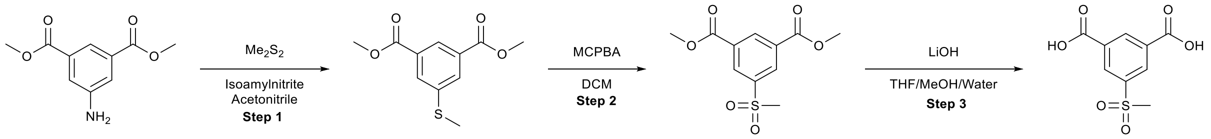


The synthesis was conducted according to [8].
**Step 1**: Synthesis of dimethyl 5-(methylthio)isophthalate.
To a solution of starting aniline (5 g, 23.8 mmol) in anhydrous acetonitrile (50 mL) at RT was added dimethylsulfide (3.2 ml, 36 mmol) followed by slow addition of isoamyl nitrite (1.6 ml, 12 mmol). The reaction was stirred and heated to reflux for 4 hours then cooled to 60°C and stirred overnight. The solvents were evaporated under the

reduced pressure, and the residue was purified by silica gel column chromatography eluting with cyclohexane to cyclohexane:ethylacetate 0:100 obtaining the product as a light orange solid in 8% yield (500 mg).
MS $(M+H)^+$: 240.9.
$^1$H NMR (300 MHz, $CDCl_3$) δ 8.33 (t, *J* = 1.6 Hz, 1H), 7.99 (d, *J* = 1.5 Hz, 2H), 3.87 (s, 6H), 2.48 (s, 3H).
$^{13}$C NMR (75 MHz, $CDCl_3$) δ 165.9, 140.4, 131.1, 130.9, 127.0, 52.5, 15.5.

**Step 2**: Synthesis of dimethyl 5-(methylsulfonyl)isophthalate.
To a solution of dimethyl 5-(methylthio)isophthalate (223 mg, 1 mmol) in DCM (10 mL) was added 70% mCPBA (614 mg, 2.5 mmol) and the reaction was stirred at RT overnight. The final mixture was diluted with saturated aqueous sodium bicarbonate and extracted with DCM. The combined organic layers were dried over Na2S04 and the solvents were evaporated under the reduced pressure. The residue was purified by silica gel column chromatography eluting with cyclohexane to cyclohexane:ethylacetate 0:100 obtaining the intermediate as a white solid.
MS $(M+H)^+$: 272.9.
$^1$H NMR (300 MHz, $CDCl_3$) δ 8.96 (t, *J* = 1.6 Hz, 1H), 8.80 (d, *J* = 1.6 Hz, 2H), 4.03 (s, 6H), 3.14 (s, 3H).

$^{13}$C NMR (75 MHz, $CDCl_3$) δ 161.8, 144.2, 132.3, 125.2, 53.0, 44.3.

**Step 3**: Synthesis of 5-(methylsulfonyl)isophthalic acid.
The product from the previous step was dissolved in a mixture of THF/MeOH/Water (1 ml/1 ml/1 ml) and LiOH (400 mg) was added. The resulting mixture was stirred overnight. After the completion of the reaction, 1M HCl was added (5 ml) and the mixture was extracted with EtOAc. The organic layers were combined, dried over the sodium sulfate and evaporated under the reduced pressure yielding the product as white solid.
Yield, over steps 2-3: 95 mg (39%).
MS $(M+H)^+$: 244.8.
$^1$H NMR (300 MHz, DMSO-$d_6$) δ 13.81 (s, 2H), 8.70 (t, *J* = 1.6 Hz, 1H), 8.60 (d, *J* = 1.6 Hz, 2H), 3.36 (s, 3H).
$^{13}$C NMR (75 MHz, DMSO-$d_6$) δ 165.2, 142.0, 134.1, 132.7, 131.4, 43.2.
**Synthesis of 5-ethynylisophthalic acid.**

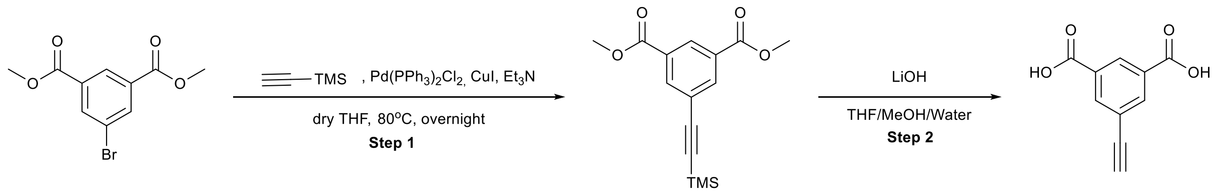


The synthesis was conducted according to [9].
**Step 1:** Synthesis of dimethyl 5-((trimethylsilyl)ethynyl)isophthalate.
A 25 mL round bottom flask equipped with a magnetic stir bar and a condenser was charged with the 5-bromoisophthalic acid dimethyl ester (819 mg, 3 mmol), trimethylsilylacetylene (589 mg, 6 mmol), THF (10 ml) and $Et_3N$ (2 ml). The reaction mixture was degassed using nitrogen gas. $Pd(PPh_3)_2Cl_2$ (84 mg, 0.12 mmol) and CuI (46 mg, 0.24 mmol) were added afterwards. The mixture was stirred at 70 °C overnight. The solvent was evaporated, and the residue was purified by silica gel column chromatography eluting with cyclohexane to cyclohexane:ethylacetate 60:40 to obtain the desired product as light orange solid in 72% yield (617 mg).

MS $(M+H)^+$: 291.6.
$^1$H NMR (300 MHz, $CDCl_3$) δ 8.60 (s, 1H), 8.29 (s, 2H), 3.95 (s, 6H), 0.26 (s, 9H)
$^{13}$C NMR (75 MHz, $CDCl_3$) δ 165.7, 137.0, 131.0, 130.4, 124.4, 102.9, 96.9, 52.7, -0.1

**Step 2**: Synthesis of 5-ethynylisophthalic acid.
The product from the previous step was dissolved in mixture of THF/MeOH/Water (5 ml/5 ml/5 ml) and LiOH (1 g) was added. The resulting mixture was stirred overnight. After the completion of the reaction, 1M HCl was added (25 ml) and the mixture was extracted with EtOAc. The organic layers were combined, dried over the sodium sulfate and evaporated under the reduced pressure yielding the product as white solid in 91% yield (530 mg).
MS $(M+H)^+$: 190.7.
$^1$H NMR (300 MHz, DMSO-$d_6$) δ 13.55 (s, 2H), 8.48 (t, *J* = 1.7 Hz, 1H), 8.19 (d, *J* = 1.6 Hz, 2H), 4.45 (s, 1H).
$^{13}$C NMR (75 MHz, DMSO-$d_6$) δ 165.7, 135.8, 132.0, 130.0, 122.7, 82.6, 81.5.
**Synthesis of 5-vinylisophthalic acid.**

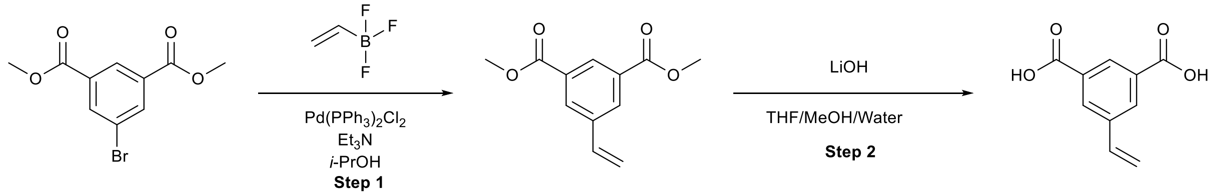


The synthesis was conducted according to [10].
**Step 1**: Synthesis of dimethyl 5-vinylisophthalate.
To a stirred in nitrogen atmosphere solution of dimethyl 5-bromoisophthalate (1.00 g, 3.7 mmol), potassium trifluoro(vinyl)borate (0.98 g, 7.3 mmol) and triethylamine (1.12 g, 11.1 mmol) in degassed isopropanol (15 mL) was added bis(triphenylphosphine)palladium(II) chloride (0.26 g, 0.37 mmol) and refluxed for 18 h. Afterwards, the solvent was removed under the reduced pressure and the residue was purified by silica gel column chromatography eluting with cyclohexane to cyclohexane:ethylacetate 70:30 to obtain the desired product as a yellow oil in 82% yield (670 mg).
MS $(M+H)^+$: 220.8.
$^1$H NMR (301 MHz, $CDCl_3$) δ 8.48 (t, *J* = 1.7 Hz, 1H), 8.17 (d, *J* = 1.7 Hz, 2H), 6.70 (dd, *J* = 17.6, 10.9 Hz, 1H), 5.84 (d, *J* = 17.3 Hz, 1H), 5.33 (d, *J* = 11.0 Hz, 1H), 3.88 (s, 6H).
$^{13}$C NMR (76 MHz, $CDCl_3$) δ 166.2, 138.3, 135.0, 131.3, 130.9, 129.7, 116.5, 52.4.

**Step 2**: Synthesis of dimethyl 5-vinylisophthalic acid.
The product from the previous step (220 mg, 1 mmol) was dissolved in mixture of THF/MeOH/Water (1 ml/1 ml/1 ml) and LiOH (80 mg, 2 mmol) was added. The resulting mixture was stirred overnight. After the completion of the reaction, 1M HCl was added (5 ml) and the mixture was extracted with EtOAc. The organic layers were combined, dried over the sodium sulfate and evaporated under the reduced pressure yielding the product as white solid in 88% yield (170 mg).
MS $(M+H)^+$: 192.9.
$^1$H NMR (300 MHz, DMSO-$d_6$) δ 13.25 (s, 2H), 8.36 (t, *J* = 1.6 Hz, 1H), 8.22 (d, *J* = 1.6 Hz, 2H), 6.91 (dd, *J* = 17.7, 11.1 Hz, 1H), 5.99 (d, *J* = 17.7 Hz, 1H), 5.41 (d, *J* = 11.1 Hz, 1H).
$^{13}$C NMR (75 MHz, DMSO-$d_6$) δ 166.4, 138.0, 135.1, 131.7, 130.5, 129.1, 116.6.

**Synthesis of 3,5-di(1,3-dioxolan-2-yl)aniline.**

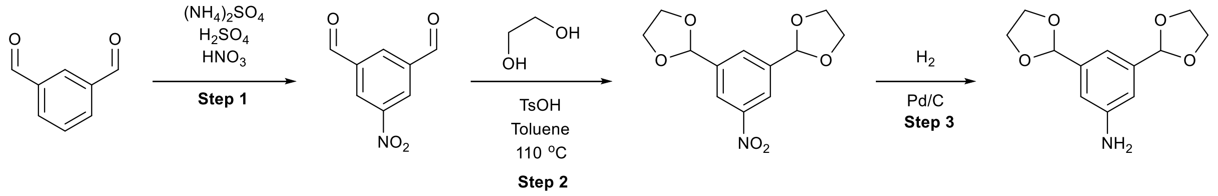


The synthesis was adopted from [11,12].

**Step 1:** Synthesis of 5-nitroisophthalaldehyde.

To a mixture of 10 g of ammonium sulfate in 16.5 ml of sulfuric and and 3.5 ml of nitric acid the solution of isophthalaldehyde (2.5 g, 18.6 mmol) in 15 ml of sulfuric acid was added. The mixture was stirred for two days and then poured in ice. The precipitate was filtered and washed with water to afford the desired product as a white solid in 40% yield (1.32 g).

MS $(M+H)^+$: 179.9.

$^1$H NMR (300 MHz, $CDCl_3$) δ 10.23 (s, 2H), 8.95 (s, 2H), 8.74 (m, 1H).

$^{13}$C NMR (75 MHz, $CDCl_3$) δ 188.6, 138.3, 134.6, 128.6.

**Step 2**: Synthesis of 2,2'-(5-nitro-1,3-phenylene)bis(1,3-dioxolane).

The mixture of the product from step 1 (820 mg, 4.8 mmol), ethylene glycol (500 mg), and p-toluenesulfonic acid (4 mg) in toluene (25 ml) was refluxed overnight in flask connected to Dean-Stark apparatus. Afterwards, the solvent was removed under the reduced pressure and the residue was purified by silica gel column chromatography eluting with cyclohexane to cyclohexane:ethylacetate 0:100 to obtain the desired product as a yellow oil in 59% yield (760 mg).

MS $(M+H)^+$: 267.7.

$^1$H NMR (300 MHz, $CDCl_3$) δ 8.27 (s, 2H), 7.84 (s, 1H), 5.82 (s, 2H), 4.16 – 3.89 (m, 8H).

$^{13}$C NMR (75 MHz, $CDCl_3$) δ 148.3, 140.7, 130.7, 122.1, 102.1, 65.5.

**Step 3:** Synthesis of 3,5-di(1,3-dioxolan-2-yl)aniline.

Product from step 2 was dissolved in EtOH (20mL). 10% Pd/C (40 mg) was added afterwards, and the mixture was stirred in hydrogen atmosphere (1 atm) in stainless steel miniclave. After the end of the reaction, the catalyst was filtered, solvent removed under the reduced pressure to obtain the desired product as a yellow oil in 81% yield (550 mg).

MS $(M+H)^+$: 237.9.

$^1$H NMR (300 MHz, $CDCl_3$) δ 6.99 (s, 1H), 6.81 (d, *J* = 1.5 Hz, 2H), 5.76 (s, 2H), 4.22 – 3.90 (m, 8H).

$^{13}$C NMR (75 MHz, $CDCl_3$) δ 146.6, 139.5, 114.8, 113.6, 103.5, 65.2.

#### Amide bond formation method.

During the synthesis of **15** it was found that the developed literature[1] conditions are not suitable for the challenging amide bond formation with 1,4-cubanedicarboxilic acid (entry 1, table S1). 3,5-diacetylaniline is weak nucleophile and steric hinderance in 1,4-cubanedicarboxilic acid further decreases the probability of successful transformation. Typical conditions for amide bond formation (entries 2,3) were also not successful. Thorough literature analysis was conducted to identify modern methods for such challenging transformations, and method described in was very efficient[13], and the corresponding diamide product was isolated in 44% after a simple filtration. Very encouraged by these results, we adopted this method as standard and used it for synthesis of all other TCB-32 analogs due to its simplicity, precipitation of the product and direct activation of carboxylic acid, eliminating the requirement for the synthesis of acyl chloride.

**Table S1.** Screening of conditions for challenging amide bond reaction.

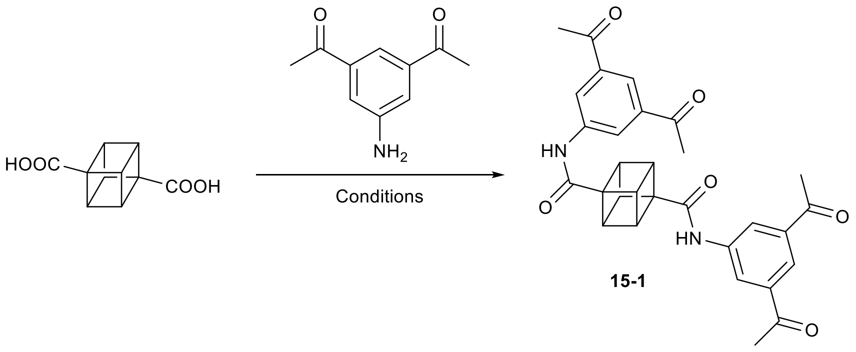


| Number | Conditions | Yield, % |
|---|---|---|
| 1. | 1. $SOCl_2$. 2. Py, DCM | 0% |
| 2. | EDC*HCl, HOBt, DIPEA, DCM | <5% |
| 3. | HATU, DIPEA, THF | <5% |
| 4. [14] | BTFFH, DIPEA, DCM | <5% |
| 5. [15] | EDC*HCl, HOBt, DMAP, DCM | 20% |
| 6. [13] | TCFH, NMI, ACN | 44% |

### Synthesis of TCB-32 analogs.

#### General procedure 1 for synthesis of guanilhydrazones.

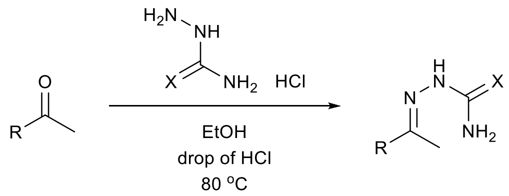

Corresponding ketone was dissolved in 5 ml 96% ethanol, 100 mg of hydrazine derivative and two drops of concentrated HCl were added. The reaction was heated to 80 °C and stirred at this temperature overnight.
Aminoguanidines:
After the completion of the reaction the solvent was evaporated, and the residue was purified using reverse-phase column chromatography eluting with water-acetonitrile with formic acid modifier. The resulting solution was lyophilized yielding the product as a solid di, tri, or tetraformate salt.
Semicarbazide and thiosemicarbazide:
The product was filtered and dried.

#### Region I.

**2.**

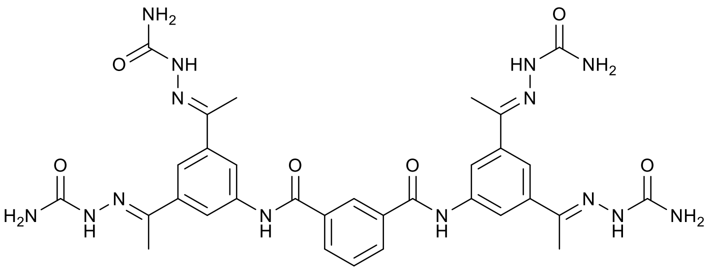

The compound were synthesized according to general procedure 1 using **1-1** (50 mg) and semicarbazide.
Yield: 52 mg (52%).
MS was not observed.
$^{1}$H NMR (400 MHz, DMSO-$d_6$) δ 10.50 (s, 2H), 10.37 (s, 4H), 8.65 (s, 8H), 8.60 (s, 1H), 8.30 (s, 4H), 8.21 (d, *J* = 9.5 Hz, 2H), 8.03 (s, 2H), 7.75 (t, *J* = 7.8 Hz, 1H), 2.38 (s, 12H).

**3.**

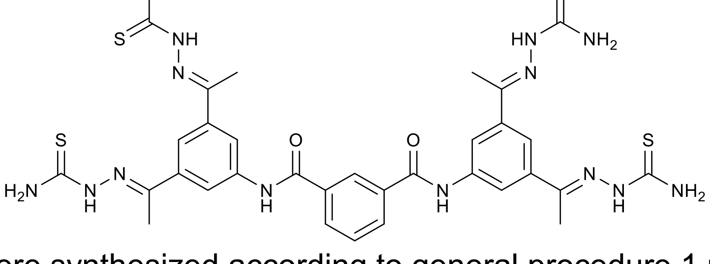

The compound were synthesized according to general procedure 1 using **1-1** (50 mg) and thiosemicarbazide.
Yield: 36 mg (33%).
MS was not observed.
$^{1}$H NMR (400 MHz, DMSO-$d_6$) δ 10.73 (s, 2H), 9.44 (s, 4H), 8.84 (s, 9H), 8.37 (s, 4H), 8.22 (d, *J* = 6.0 Hz, 1H), 7.86 (s, 2H), 7.72 (t, *J* = 7.8 Hz, 1H), 2.26 (s, 12H).

**4.**

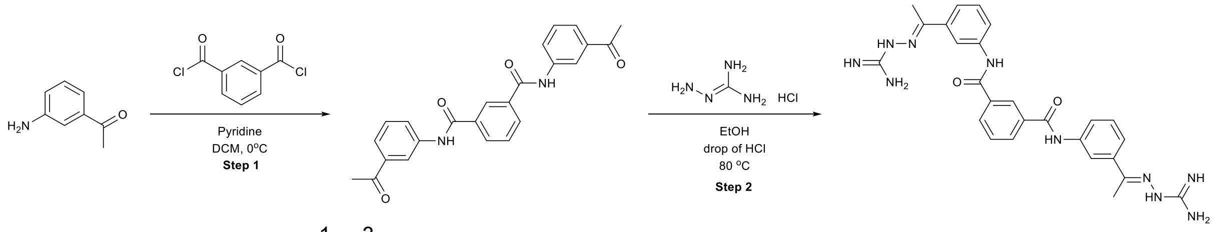


**Step 1**: synthesis of N[1],N[3]-bis(3-acetylphenyl)isophthalamide.
3-aminoacethophenone (38 mg, 0.28 mmol) was dissolved in 2 ml of dry $CH_2Cl_2$, 0.06 ml of pyridine were added, and the reaction mixture cooled down to 0 °C using an ice cooling bath. After that isophthaloyl chloride (28 mg, 0.14 mmol, dissolved in 1 ml of dry DCM) was added slowly. Then, the reaction mixture was warmed up to rt and stirred overnight. The solvent was evaporated, and the residue was used in the next reaction without additional purification.
**Step 2:** synthesis of **4**.
The compound was synthesized according to general procedure 1 using the product from the previous step.
Yield, over two steps: 40 mg (48%).
MS $(M+H)^+$: 512.9.
$^1$H NMR (300 MHz, DMSO-$d_6$) δ 10.52 (s, 2H), 8.61 (s, 1H), 8.41 (s, 2H), 8.23 – 8.14 (m, 4H), 7.92 (d, *J* = 7.5 Hz, 2H), 7.74 (m, 10H), 7.40 (t, *J* = 8.0 Hz, 2H), 2.33 (s, 6H).

**Synthesis of 5-6.**

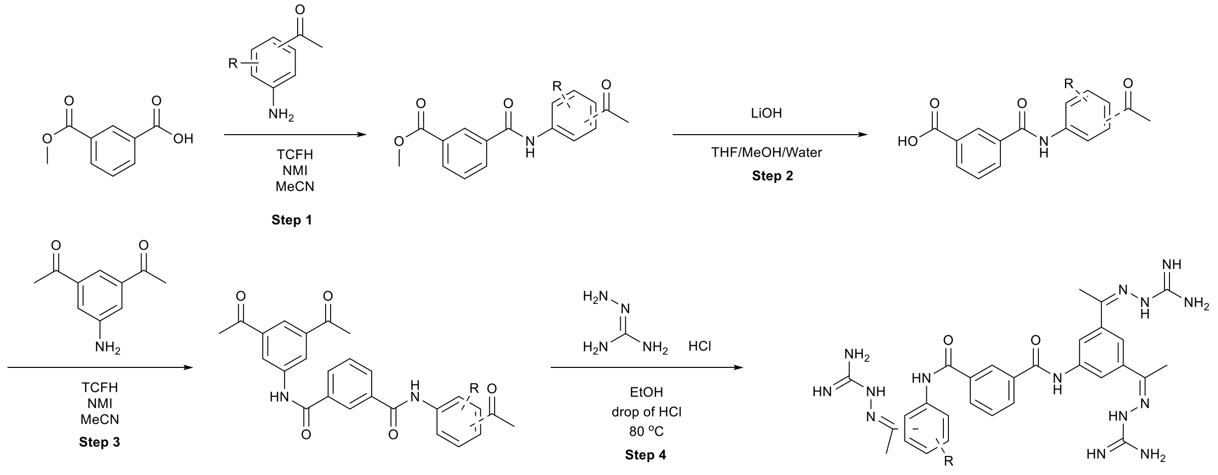


**Step 1.**
The 3-(methoxycarbonyl)benzoic acid (180 mg, 1 mmol) and aminoacetophenone (1 mmol) were dissolved in 3 ml of ACN. 0.2 mL N-methylimidazole and TCFH (210 mg, 0.75 mmol) were added sequentially. The reaction was stirred overnight. After the completion of the reaction solvents were evaporated, and the residue was purified using flash chromatography eluting with Cyclohexane: Ethylacetate 100:0% to Cyclohexane: Ethylacetate 0:100% yielding product as white solid**.**
**Step 2.**
The product from previous step was dissolved in mixture of THF/MeOH/Water (1 ml/1 ml/1 ml) and LiOH (200 mg) was added. The resulting mixture was stirred overnight. After the completion of the reaction, 1 M HCl was added (5 ml) and the mixture was extracted with EtOAc. The organic layers were combined, dried over sodium sulfate and evaporated under reduced pressure yielding the desired product as white solid.
**Step 3**.

The product from previous step (0.14 mmol) and intermediate 1 (50 mg, 0.28 mmol) were dissolved in 1 ml of MeCN. 0.06 mL N-methylimidazole and TCFH (73 mg, 0.28 mmol) were added sequentially. The reaction was stirred overnight. After the completion of the reaction the product was filtered off yielding the desired product as white solid.

**Step 4**.

The compound was synthesized according to general procedure 1 using the product from the previous step.

**5.**

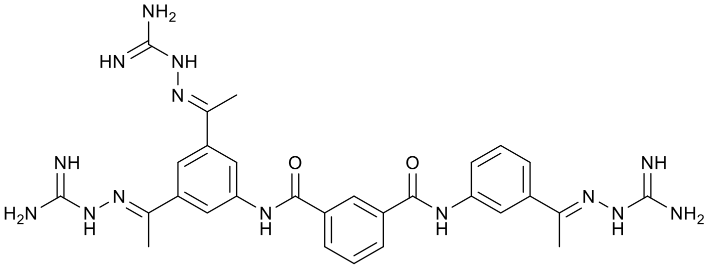


**Step 1:** synthesis of methyl 3-((3-acetylphenyl)carbamoyl)benzoate.
Starting materials: 3-aminoacetophenone (135 mg).
MS $(M+H)^+$: 298.0.
$^1$H NMR (300 MHz, DMSO-$d_6$) δ 10.59 (s, 1H), 8.51 (s, 1H), 8.30 (s, 1H), 8.20 (d, *J* = 9.5 Hz, 1H), 8.11 (d, *J* = 9.3 Hz, 1H), 8.04 (d, *J* = 8.1 Hz, 1H), 7.72 – 7.61 (m, 2H), 7.47 (t, *J* = 7.9 Hz, 1H), 3.86 (s, 3H), 2.53 (s, 3H).
$^{13}$C NMR (75 MHz, DMSO) δ 197.61, 165.72, 164.70, 146.65, 139.32, 137.24, 135.06, 132.40, 132.17, 129.88, 129.10, 128.30, 124.89, 123.87, 119.75, 52.40, 26.74.

**Step 2**: synthesis of 3-((3-acetylphenyl)carbamoyl)benzoic acid.
Yield, over steps 1-2: 118 mg (40%).
MS $(M+H)^+$: 284.0.
$^1$H NMR (300 MHz, DMSO-$d_6$) δ 10.56 (s, 1H), 8.49 (s, 1H), 8.29 (s, 1H), 8.15 (d, *J* = 9.3 Hz, 1H), 8.08 (d, *J* = 7.8 Hz, 1H), 8.01 (d, *J* = 10.4 Hz, 0H), 7.68 – 7.56 (m, 2H), 7.45 (t, *J* = 8.0 Hz, 1H), 2.42 (s, 3H).
$^{13}$C NMR (75 MHz, DMSO) δ 197.62, 166.77, 164.87, 139.37, 137.23, 134.92, 132.33, 132.03, 131.04, 129.04, 128.89, 128.44, 124.88, 123.80, 119.75, 26.74.

**Step 3:** synthesis of $N^1$-(3-acetylphenyl)-$N^3$-(3,5-diacetylphenyl)isophthalamide.
Starting material: 3-((3-acetylphenyl)carbamoyl)benzoic acid (40 mg).

**Step 4**: Synthesis of **5**.
Yield, over steps 3-4: 25 mg (24%).
MS $(M+H)^+$: 611.1.
$^1$H NMR (300 MHz, DMSO-$d_6$) δ 10.57 (s, 1H), 10.54 (s, 1H), 8.64 (s, 1H), 8.41 (s, 3H), 8.28 (s, 2H), 8.23 – 8.18 (m, 3H), 8.04 (s, 1H), 7.93 (d, *J* = 7.4 Hz, 1H), 7.86 – 7.62 (m, 12H), 7.41 (t, *J* = 8.0 Hz, 1H), 2.38 (s, 6H),
2.33 (s, 3H).

**6.**

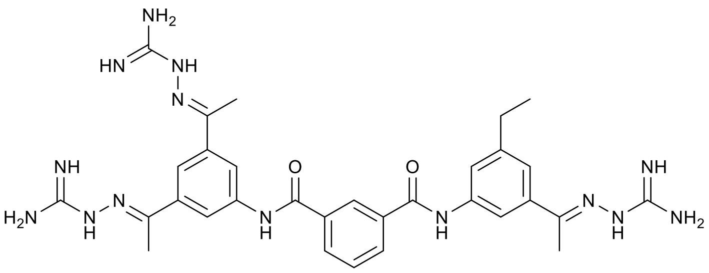


**Step 1:** synthesis of methyl 3-((3-acetyl-5-ethylphenyl)carbamoyl)benzoate.
Starting material: 1-(3-amino-5-ethylphenyl)ethan-1-one (164 mg).
MS $(M+H)^+$: 326.1.
$^1$H NMR (300 MHz, DMSO-$d_6$) δ 10.53 (s, 1H), 8.51 (s, 1H), 8.20 (d, *J* = 9.6 Hz, 1H), 8.15 (d, *J* = 1.8 Hz, 1H), 8.11 (s, 1H), 7.89 (s, 1H), 7.66 (t, *J* = 7.8 Hz, 1H), 7.53 (s, 1H), 3.86 (s, 3H), 2.65 (q, *J* = 7.6 Hz, 2H), 2.53 (s, 3H), 1.18 (t, *J* = 7.6 Hz, 3H).
$^{13}$C NMR (75 MHz, DMSO) δ 197.72, 165.72, 164.62, 144.76, 139.30, 137.32, 135.10, 132.39, 132.13, 129.87, 129.10, 128.26, 124.28, 123.28, 117.54, 52.40, 28.15, 26.78, 15.41.

**Step 2:** synthesis of 3-((3-acetyl-5-ethylphenyl)carbamoyl)benzoic acid.
Yield, over step 1-2: 105 mg (33%).
MS $(M+H)^+$: 312.1.
$^1$H NMR (300 MHz, DMSO-$d_6$) δ 13.27 (s, 1H), 10.58 (s, 1H), 8.58 (s, 1H), 8.28 – 8.22 (m, 2H), 8.16 (d, *J* = 7.8 Hz, 1H), 7.97 (s, 1H), 7.68 (t, *J* = 7.8 Hz, 1H), 7.58 (s, 1H), 2.70 (q, *J* = 7.5 Hz, 2H), 2.59 (s, 3H), 1.24 (t, *J* = 7.6 Hz, 3H).
$^{13}$C NMR (75 MHz, DMSO) δ 197.71, 166.78, 164.79, 144.73, 139.37, 137.31, 134.95, 132.29, 132.00, 131.04, 128.88, 128.41, 124.26, 123.19, 117.54, 28.15, 26.77, 15.39.

**Step 3:** synthesis of $N^1$-(3-acetyl-5-ethylphenyl)-$N^3$-(3,5-diacetylphenyl) isophthalamide.
Starting material: 3-((3-acetyl-5-ethylphenyl)carbamoyl)benzoic acid (40 mg). .
**Step 4:** synthesis of **6**.
Yield, over steps 3-4: 15 mg (14%).
MS $(M+H)^+$: 639.6.
$^1$H NMR (300 MHz, DMSO-$d_6$) δ 10.34 (s, 1H), 10.26 (s, 1H), 8.44 (s, 1H), 8.18 (s, 3H), 8.08 (s, 2H), 8.00 (m, 2H), 7.82 (s, 2H), 7.60 (s, 1H), 7.53 (t, *J* = 7.8 Hz, 1H), 7.60-7.25 (m, 13 H), 2.48 (q, *J* = 7.8 Hz, 1H), 2.18 (s, 6H), 2.12 (s, 3H), 1.05 (t, *J* = 7.6 Hz, 3H).

### Region II.

#### Synthesis of 7.

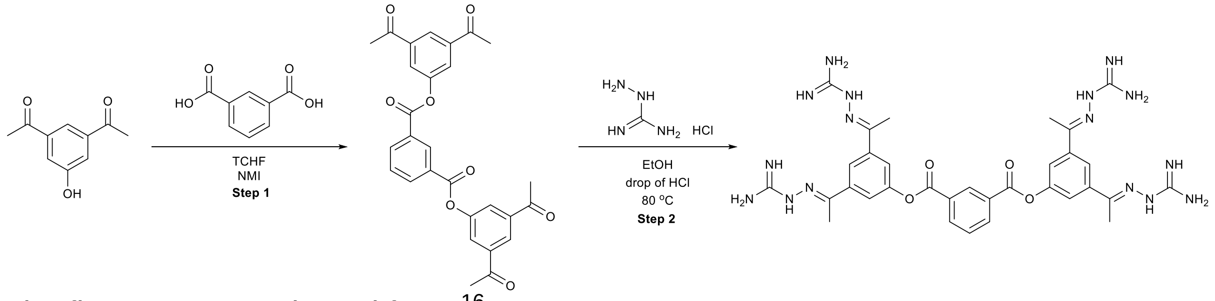


The first step was adopted from [16].

**Step 1**: Synthesis of **7-1**.

The isophtalic acid (23 mg, 0.14 mmol equiv) and 3,5-diacetylphenol (50 mg, 0.29 mmol) were dissolved in 1 ml of DCM. Pyridine (90 mg) and TCFH (78 mg, 0.28 mmol) were added sequentially. The reaction was stirred overnight. After the completion of the reaction, volatiles were evaporated under reduced pressure and the residue was purified using flash chromatography eluting with Cyclohexane: Ethylacetate 100:0% to Cyclohexane: Ethylacetate 0:100% yielding the desired product as white solid.

MS $(M+H)^+$: 487.6.

$^1$H NMR (300 MHz, DMSO-$d_6$) δ 8.81 (s, 1H), 8.47 (dd, *J* = 7.8, 1.9 Hz, 2H), 8.33 (s, 2H), 8.15 (d, *J* = 1.5 Hz, 4H), 7.85 (t, *J* = 7.8 Hz, 1H), 2.62 (s, 12H).

$^{13}$C NMR (75 MHz, DMSO-$d_6$) δ 196.75, 163.67, 150.99, 138.57, 135.16, 130.95, 129.98, 129.53, 126.05, 125.21, 27.00.

**Step 2**: Synthesis of **7**.

The compound was synthesized according to general procedure 1 using the product from the previous step.

Yield, over two steps: 21 mg (17%).

MS $(M+H)^+$: 711.8

$^1$H NMR (300 MHz, DMSO-$d_6$) δ 8.89 (s, 1H), 8.55 (s, 2H), 8.36 (s, 4H), 8.07 (s, 2H), 7.96 (s, 4H), 7.92 (t, *J* = 7.9 Hz, 1H), 7.57 (s, 16H), 2.38 (s, 12H).

#### Synthesis of 8.

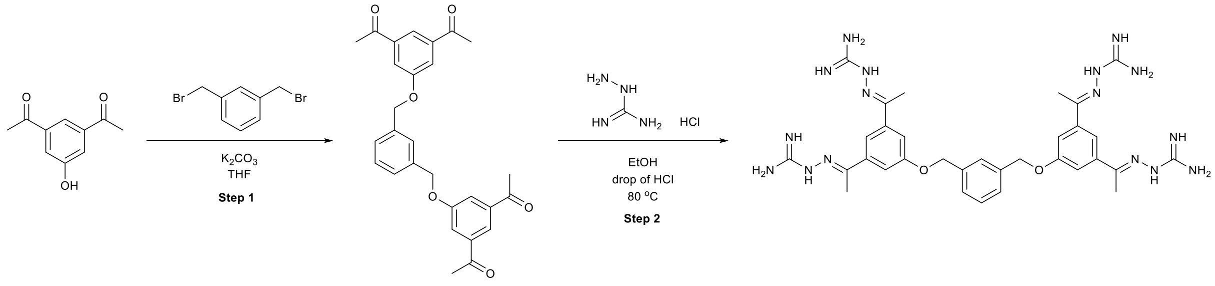


**Step 1**: Synthesis of **8-1**.

To a solution of 3,5-diacetylphenol (80 mg, 0.5 mmol) and $K_2CO_3$ (200 mg, 1.45 mmol) in DMF (1 mL), α,α′-dibromo-m-xylene (65 mg, 0.25 mmol) was added. The resulting mixture was stirred at 60 °C overnight. After the completion of the reaction, volatiles were evaporated under reduced pressure and the residue was purified using flash chromatography eluting with Cyclohexane: Ethylacetate 100:0% to Cyclohexane: Ethylacetate 0:100% yielding the desired product as white solid.

MS $(M+H)^+$: 459.1

$^1$H NMR (300 MHz, DMSO-$d_6$) δ 7.98 (s, 2H), 7.72 (s, 4H), 7.57 (s, 1H), 7.41 (s, 3H), 5.23 (s, 4H), 2.57 (s, 12H).

$^{13}C$ NMR (75 MHz, DMSO-d$_6$) δ 197.27, 138.47, 136.80, 128.71, 127.43, 126.96, 120.22, 118.47, 69.63, 26.96.
**Step 2**: Synthesis of **8**.
The compound was synthesized according to general procedure 1 using the product from the previous step.
Yield, over two steps: 28 mg (23%).
MS $(M+H)^+$: 683.7.
$^1H$ NMR (300 MHz, DMSO-d$_6$) δ 8.39 (s, 4H), 7.77 (s, 18H), 7.63 (s, 1H), 7.58 (s, 4H), 7.48 (m, 3H), 5.26 (s, 4H), 2.35 (s, 12H).
**Synthesis of 9.**

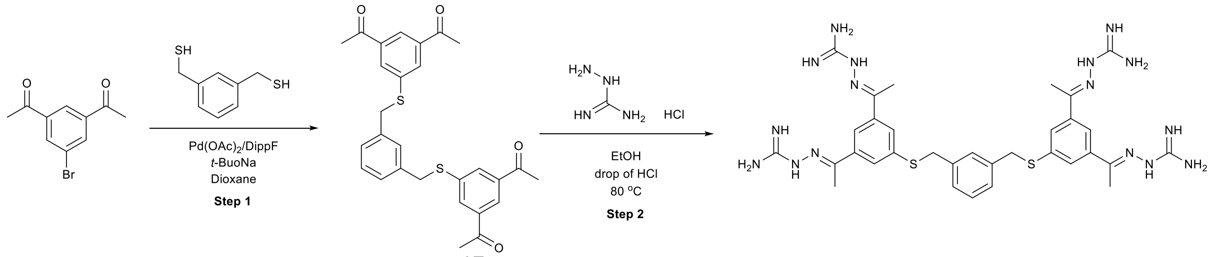


The first step was adopted from [17].
**Step 1**: Synthesis of **9-1**.
A 8 ml microwave vial was charged with 3,5-diacetylbromobenzene (120 mg, 0.5 mmol), 1,3-phenylendimethanthiol, (42.5 mg, 0.25 mmol), Pd(OAc)$_2$ (2.24 mg, 0.01 mmol), dippf (5 mg, 0.011 mmol) , and *t*-BuONa (57 mg, 0.6 mmol). The reaction tube was sealed. The septum was pierced with a needle attached to a Schlenk line, and the tube was evacuated and backfilled with nitrogen (this process was repeated a total of three times). 1,4-dioxane (1.0 mL) were added via a syringe. The reaction was stirred at 85 °C overnight. After the completion of the reaction, the solvents were evaporated, and the residue was purified using flash chromatography eluting with Cyclohexane: Ethylacetate 100:0% to Cyclohexane: Ethylacetate 0:100% yielding product as white solid**.**
MS $(M+H)^+$: 491.2.
$^1H$ NMR (300 MHz, $CDCl_3$) δ 8.19 (s, 2H), 7.90 (s, 4H), 7.27 – 7.02 (m, 4H), 4.09 (s, 4H), 2.51 (s, 12H).
**Step 2**: Synthesis of **9**.
The compound was synthesized according to general procedure 1 using the product from the previous step.
Yield, over two steps: 2 mg (0.5%).
MS $(M+H)^+$: 715.7.
$^1H$ NMR (300 MHz, DMSO-d$_6$) δ 8.32 (s, 4H), 7.90 (s, 2H), 7.74 (s, 4H), 7.49 (s, 1H), 7.25 (m, 3H), 7.02 (s, 16H), 4.32 (s, 4H), 2.28 (s, 12H).
**Synthesis of 10.**

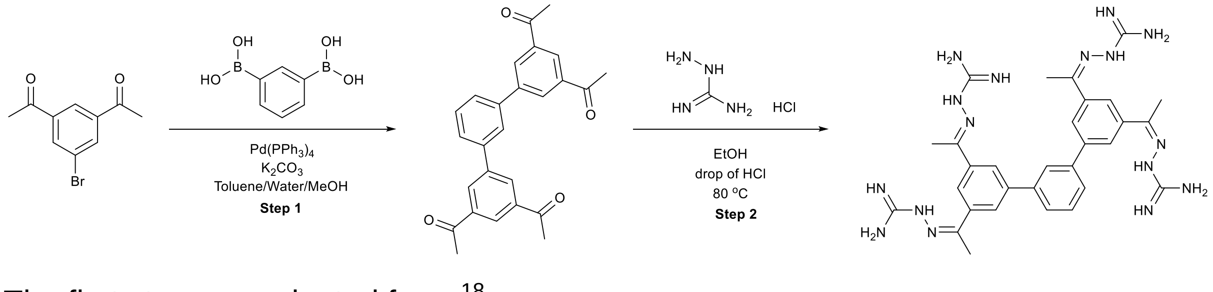


The first step was adopted from [18].
**Step 1**: Synthesis of **10-1**.
A 100 mL round bottom flask equipped with a magnetic stir bar and a condenser was charged with 3,5-diacetylbromobenzene (120 mg, 0.5 mmol), 1,3-benzenediboronic

acid (41 mg, 0.25 mmol) and $K_2CO_3$ (550 mg, 4 mmol). The solids were suspended in 12 ml/5 ml/1.5 ml toluene/MeOH/$H_2O$ (18.5 mL), degassed, and the atmosphere was exchanged with nitrogen. Afterwards, $Pd(PPh_3)_4$ (84 mg, 72 µmol) was added. The mixture was stirred at 80 °C overnight. The solvent was evaporated and the residue was purified by silica gel column chromatography eluting with cyclohexane to cyclohexane:ethylacetate 20:80 to obtain the desired product as a white solid.

MS $(M+H)^+$: 399.3.
$^1$H NMR (300 MHz, DMSO-$d_6$) δ 8.45 (d, $J$ = 1.6 Hz, 4H), 8.38 (t, $J$ = 1.6 Hz, 2H), 8.12 (s, 1H), 7.84 – 7.81 (m, 2H), 7.66 – 7.61 (m, 1H), 2.68 (s, 12H).

$^{13}$C NMR (75 MHz, DMSO-$d_6$) δ 197.65, 147.03, 141.09, 139.64, 137.85, 131.02, 129.88, 127.23, 126.27, 126.11, 27.13.

**Step 2**: Synthesis of **10**.
The compound was synthesized according to general procedure 1 using the product from the previous step.
Yield, over two steps: 6 mg (1%).
MS $(M+H)^+$: 623.8.
$^1$H NMR (300 MHz, DMSO-$d_6$) δ 8.39 (s, 4H), 8.21 (s, 6H), 8.07 (s, 1H), 7.84 (d, $J$ = 9.5 Hz, 2H), 7.76 – 7.55 (m, 17H), 2.44 (s, 12H).

**Synthesis of 11.**

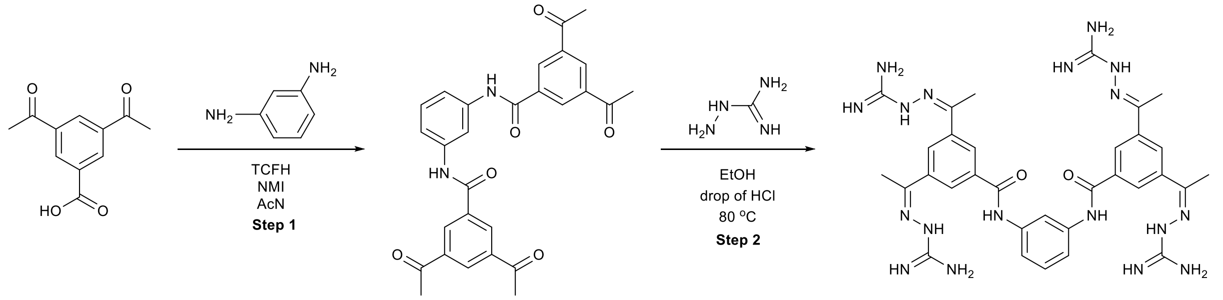


**Step 1**: Synthesis of **11-1**.
3,5-diacetylbenzoic acid (58 mg, 0.14 mmol) and m-phenylenediamine (15 mg, 0.14 mmol) were dissolved in 1 ml of MeCN. N-methylimidazole (0.06 ml) and TCFH (73 mg, 0.28 mmol) were added sequentially. After the completion of the reaction, the solids were filtrated, washed with ethanol and used in the next step without additional purification.
**Step 2**: Synthesis of **11**.
The compound was synthesized according to general procedure 1 using the product from the previous step.
Yield, over two steps: 14 mg (11%).
MS $(M+H)^+$: 826.8.
$^1$H NMR (300 MHz, DMSO-$d_6$) δ 10.48 (s, 2H), 8.38 (s, 4H), 8.38 – 8.29 (m, 5H), 7.64 (s, 16H), 7.54 (d, $J$ = 10.2 Hz, 2H), 7.39 – 7.34 (m, 1H), 2.43 (s, 12H).

#### Region III.

##### Synthesis of 1, 12, 14-15, 17-26, 29-34.

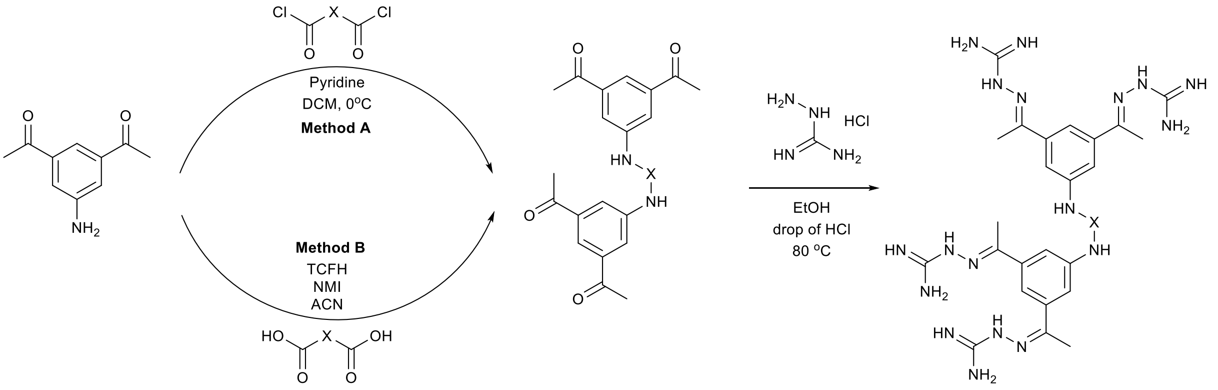


##### Synthesis of examples 1, 12, 14, 32.

Step 1:

The corresponding diacid (0.14 mmol) was suspended in thionyl chloride (0.5 mL) and five drops of dry DMF were added. The mixture was heated to 80 °C and stirred for 2 h. The solvent was removed in vacuo yielding the diacid dichloride. Respective products were used without further purification.

Step 2:

3,5-diacetylpyridine (50 mg, 0.28 mmol) was dissolved in 2 ml of dry $CH_2Cl_2$, 0.06 ml pyridine was added and the reaction mixture cooled down to 0 °C using an ice bath. The corresponding diacid chloride from the previous step (dissolved in 1 ml dry DCM) was added slowly. Then, the reaction mixture was warmed up to rt and stirred overnight. The solvent was evaporated, and the residue was used in the next reaction without additional purification.

Step 3:

The corresponding diamide from the previous step was dissolved in 5 ml 96% ethanol, 100 mg of aminoguanidine hydrochloride and two drops of concentrated HCl were added. The reaction was heated to 80 °C and stirred at this temperature overnight. After the completion of the reaction the solvent was evaporated, and the residue was purified using reverse-phase column chromatography eluting with water-acetonitrile with formic acid modifier. The resulting solution was lyophilized yielding the product as a solid tetra formate salt.

##### Synthesis of examples 15, 17-26, 29-31, 33-34.

Step 1:

The acid (0.14 mmol) and 3,5-diacetylaniline (50 mg, 0.28 mmol) were dissolved in 1 ml of MeCN. 0.06 mL N-methylimidazole and TCFH (73 mg, 0.28 mmol) were added sequentially. The reaction was stirred overnight. After the completion of the reaction, the product was filtered and used in the next step.

Step 2:

The corresponding diamide from the previous step was dissolved in 5 ml of 96% ethanol, 100 mg of aminoguanidine hydrochloride and two drops of concentrated HCl were added. The reaction was heated to 80 °C and stirred at this temperature overnight. After the completion of the reaction the solvent was evaporated, and the residue was purified using reverse-phase chromatography eluting with water-

acetonitrile with formic acid modifier. The resulting solution was lyophilized yielding the product as a solid tetraformate salt.
**1** (TCB-32).

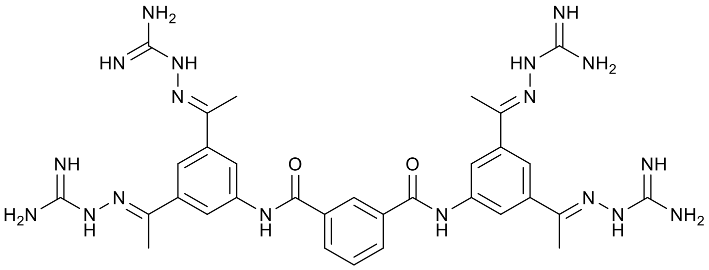

Starting material: isophthalic acid (23 mg).
**1-1.**
MS $(M+H)^+$: 483.1.
$^1$H NMR (300 MHz, DMSO $d_6$) δ 10.82 (s, 1H), 8.68 – 8.66 (m, 5H), 8.29 – 8.20 (m, 4H), 7.76 (t, *J* = 7.7 Hz, 1H), 2.67 (s, 12H).
$^{13}$C NMR (75 MHz, DMSO) δ 197.3, 165.3, 139.9, 137.6, 134.6, 131.1, 128.9, 127.2, 123.6, 123.2, 26.9.
Yield, over two steps: 24%.
MS $(M+H)^+$: 709.3.
$^1$H NMR (300 MHz, DMSO-$d_6$) δ 10.56 (s, 2H), 8.65 (s, 2H), 8.38 (s, 4H), 8.26 (s, 4H), 8.21 (d, J = 7.7 Hz, 2H), 8.03 (s, 2H), 7.74 (t, J = 7.8 Hz, 1H), 7.64 (brs, 16H), 2.37 (s, 12H).
$^{13}$C NMR (101 MHz, DMSO $d_6$) δ 166.5, 165.0, 157.5, 150.4, 138.9, 138.7, 135.0, 130.7, 128.7, 127.0, 119.8, 119.3, 14.7.

**12.**

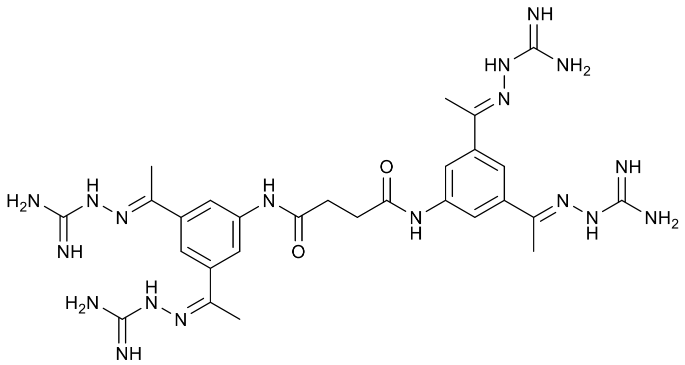

Starting material: succinic acid (16 mg).
Yield, over two steps: 12 mg (8%).
MS $(M+H)^+$: 661.5.
$^1$H NMR (300 MHz, DMSO-$d_6$) δ 10.15 (s, 2H), 8.39 (s, 4H), 8.04 (s, 4H), 7.94 (s, 2H), 7.71 (s, 16H), 2.70 (s, 4H), 2.32 (s, 12H).

**14.**

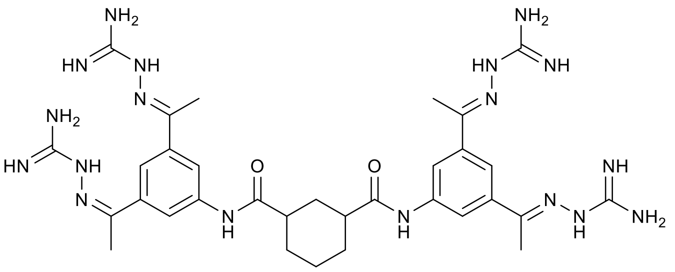

Starting material: cyclohexane-1,3-dicarboxylic acid (24 mg).
Yield, over two steps: 20 mg (16%).
Mixture of cis and trans.
MS $(M+H)^+$: 715.4.
$^1$H NMR (300 MHz, DMSO-$d_6$) δ 10.04 (s, 2H), 9.98 (s, 2H), 8.38 (s, 8H), 8.10 (s, 4H), 8.07 (s, 4H), 7.95 (s, 4H), 7.63 (s, 32H), 2.33 (s, 24H), 1.92 (m, 8H), 1.71 (m, 8H), 1.41 (m, 4H).

**15.**

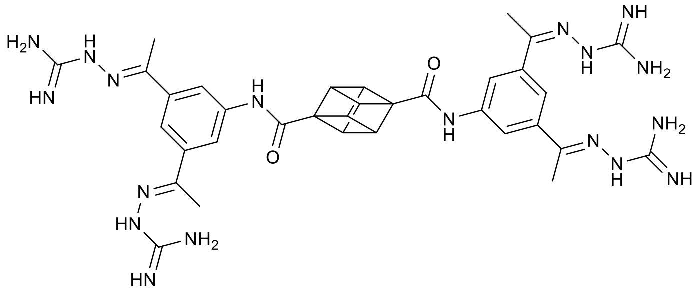

Starting material: cubane-1,4-dicarboxylic acid (31 mg).
**15-1.**
MS (M+H)+: 510.37.
$^1$H NMR (300 MHz, DMSO-$d_6$) δ 10.12 (s, 2H), 8.54 (s, 4H), 8.18 (s, 2H), 4.32 (s, 6H), 2.65 (s, 12H).
**15.**
Yield, over two steps: 13 mg (10%).
MS (M+H)+: 735.3.
$^1$H NMR (300 MHz, DMSO-$d_6$) δ 9.81 (s, 2H), 8.38 (s, 4H), 8.13 (s, 4H), 7.99 (s, 2H), 7.65 (s, 16H), 4.31 (s, 6H), 2.35 (s, 12H).

**16.**

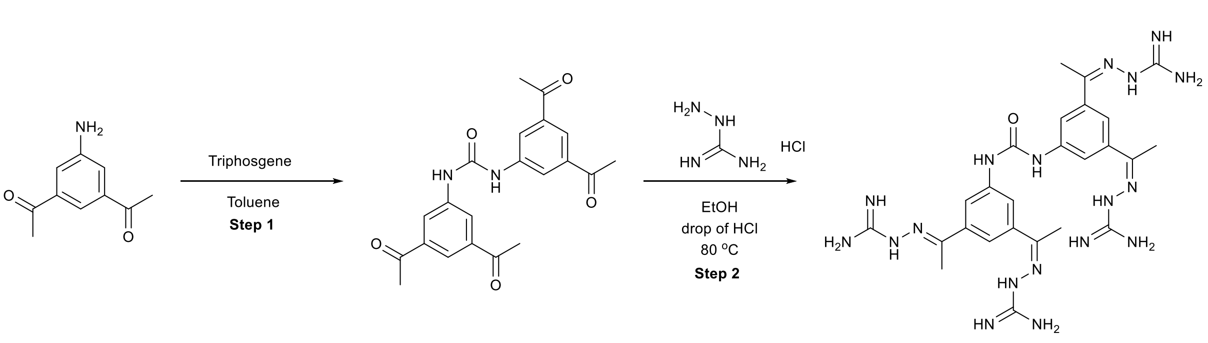


**Step 1:** Synthesis of **16-1**.

3,5-diacetylaniline (120 mg, 0.68 mmol) was suspended in toluene (2 mL) and stirred in an ice bath. A solution of triphosgene (66 mg, 0.2 mmol) in toluene (0.5 mL) was added. The suspension was allowed to warm to rt, and was stirred overnight at rt. The product precipitated, was filtered and used in the next step without additional purification.

**Step 2:** Synthesis of **16**.

The compound was synthesized according to general procedure 1 using the product from the previous step.
Yield, over two steps: 15 mg (3%).
MS $(M+H)^+$: 605.6.
$^1$H NMR (300 MHz, DMSO-$d_6$) δ 9.74 (s, 2H), 8.39 (s, 4H), 7.96 (s, 4H), 7.85 (s, 2H), 7.45 (s, 16H), 2.34 (s, 12H).

**17 (TCB-494).**

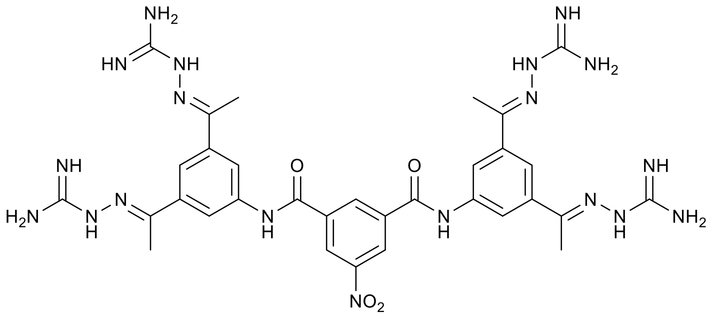

Starting material: 5-nitroisophthalic acid (29 mg).
Yield, over two steps: 8 mg (6%).
MS $(M+H)^+$: 754.7.
$^1$H NMR (300 MHz, DMSO-$d_6$) δ 10.93 (s, 2H), 9.11 (s, 1H), 9.04 (s, 2H), 8.37 (s, 4H), 8.26 (s, 4H), 8.04 (s, 2H), 7.43 (s, 16H), 2.38 (s, 12H).

**18.**

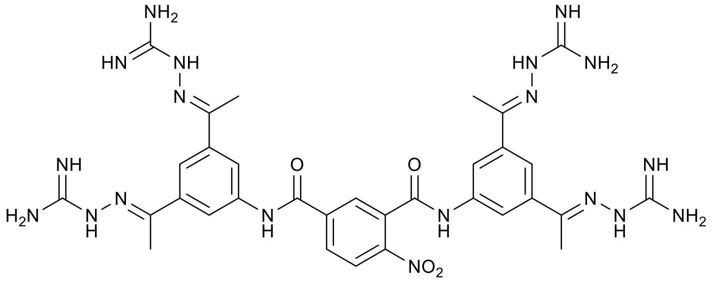

Starting material: 4-nitroisophthalic acid (29 mg).
Yield, over two steps: 20 mg (15%).
MS $(M+H)^+$: 754.7.
$^1$H NMR (300 MHz, DMSO-$d_6$) δ 11.03 (s, 1H), 10.84 (s, 1H), 8.46 (s, 1H), 8.36 (s, 4H), 8.37 – 8.29 (m, 2H), 8.26 (s, 2H), 8.16 (s, 2H), 8.11 – 8.02 (m, 2H), 7.50 (brs, 16H), 2.37 (s, 6H), 2.36 (s, 6H).

**19.**

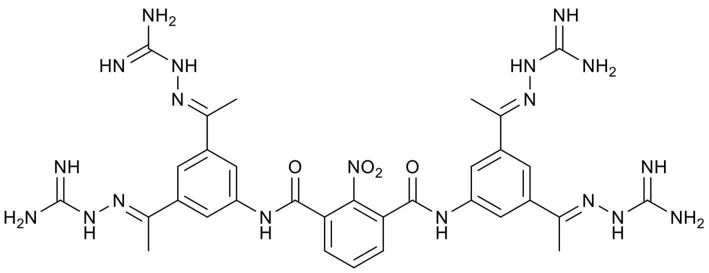

Starting material: 2-nitroisophthalic acid (29 mg).
Yield, over two steps: 7 mg (5%).
MS $(M+H)^+$: 754.7.
$^1$H NMR (300 MHz, DMSO-$d_6$) δ 10.89 (s, 2H), 8.25 (s, 4H), 8.08 (s, 4H), 8.03 (s, 1H), 8.01 (s, 2H), 7.97 (s, 2H), 6.49 (s, 16H), 2.31 (s, 12H).

**20.**

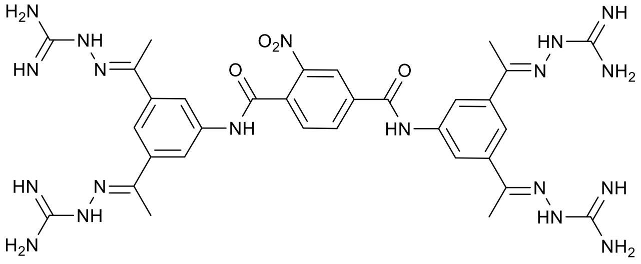

Starting material: 2-nitroterephthalic acid (29 mg).
Yield, over two steps: 11 mg (8%).
MS $(M+H)^+$: 754.6.
$^1$H NMR (300 MHz, DMSO-$d_6$) δ 10.98 (s, 1H), 10.82 (s, 1H), 8.77 (s, 1H), 8.49 (d, *J* = 9.6 Hz, 1H), 8.38 (s, 4H), 8.24 (s, 2H), 8.13 (s, 2H), 8.06 (s, 2H), 8.01 (d, *J* = 8.0 Hz, 1H), 7.64 (s, 8H), 7.58 (s, 8H), 2.38 (s, 6H), 2.36 (s, 6H).

**21.**

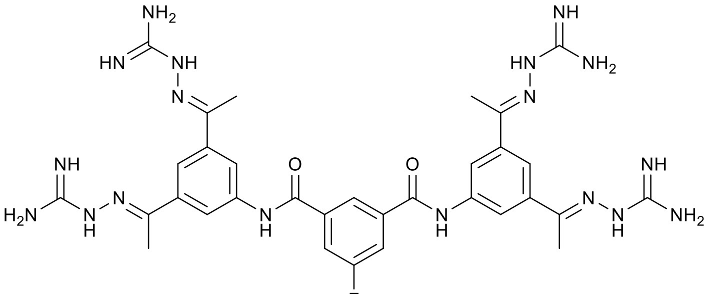

Starting material: 5-fluoroisophthalic acid (26 mg).
Yield, over two steps: 21 mg (16%).
MS $(M+H)^+$: 727.7.
H NMR (300 MHz, DMSO-$d_6$) δ 10.65 (s, 2H), 8.55 (s, 1H), 8.39 (s, 4H), 8.25 (s, 4H), 8.11 – 8.01 (m, 4H), 7.58 (s, 16H), 2.37 (s, 12H).

**22 (TCB-541).**

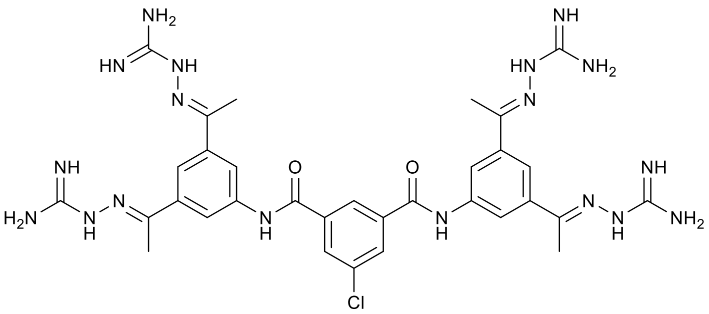

Starting material: 5-chloroisophthalic acid (28 mg).
Yield, over two steps: 8 mg (6 %).
MS $(M+H)^+$: 743.8
$^1$H NMR (300 MHz, DMSO-$d_6$) δ 10.73 (s, 2H), 8.72 (s, 1H), 8.32-8.29 (m, 8H), 8.06 (s, 2H), 7.40 (s, 16H), 2.38 (s, 12H).

**23.**

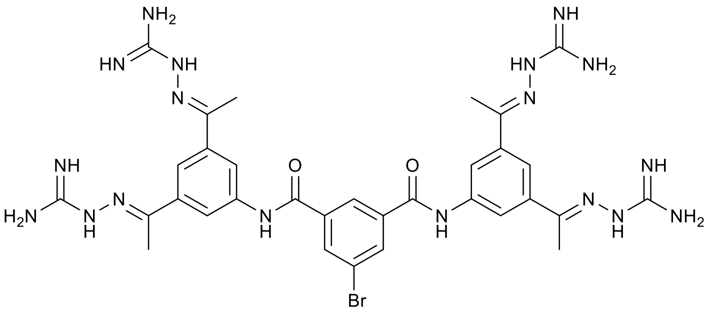

Starting material: 5-bromoisophthalic acid (34 mg).
Yield, over two steps: 12 mg (9%).
MS $(M+H)^+$: 787.6.
$^1$H NMR (300 MHz, DMSO-$d_6$) δ 10.66 (s, 2H), 8.67 (s, 1H), 8.42 (s, 2H), 8.37 (s, 4H), 8.25 (s, 4H), 8.03 (s, 2H), 7.49 (s, 16H), 2.37 (s, 12H).

**24.**

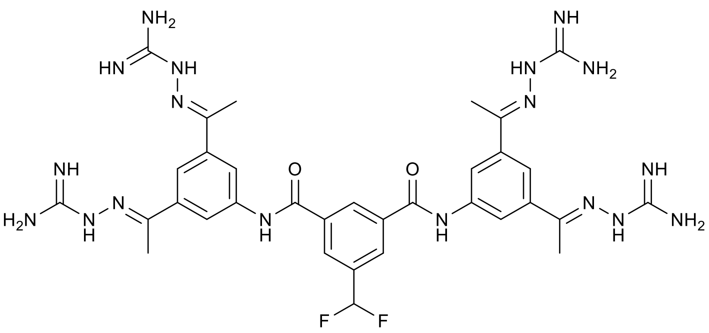

Starting material: 5-(difluoromethyl)isophthalic acid (30 mg).
Yield, over two steps: 32 mg (24 %).
MS $(M+H)^+$: 759.8.
$^1$H NMR (300 MHz, DMSO-$d_6$) δ 10.82 (s, 2H), 8.92 (s, 1H), 8.43 (s, 2H), 8.39 (s, 1H), 8.31 (s, 32H), 8.07 (s, 2H), 7.75 (s, 16H), 7.28 (t, *J* = 55.4 Hz, 1H), 2.39 (s, 12H).

**25.**

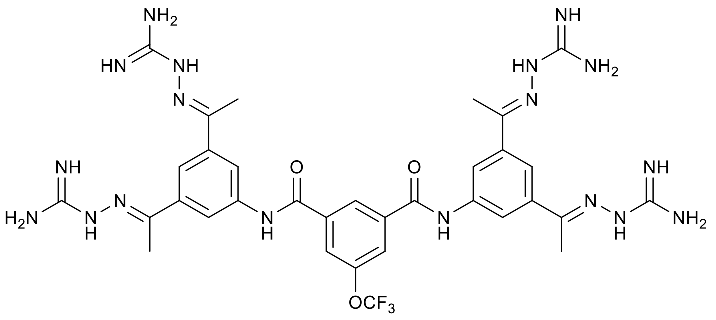

Starting material: 5-(trifluoromethoxy)isophthalic acid (35 mg).
Yield, over two steps: 18 mg (13 %).
MS $(M+H)^+$: 793.8.
$^1H$ NMR (300 MHz, DMSO-$d_6$) δ 10.76 (s, 2H), 8.77 (s, 1H), 8.25 (s, 4H), 8.19 (s, 2H), 8.03 (s, 2H), 7.49 (s, 16H), 2.37 (s, 12H).

**26.**

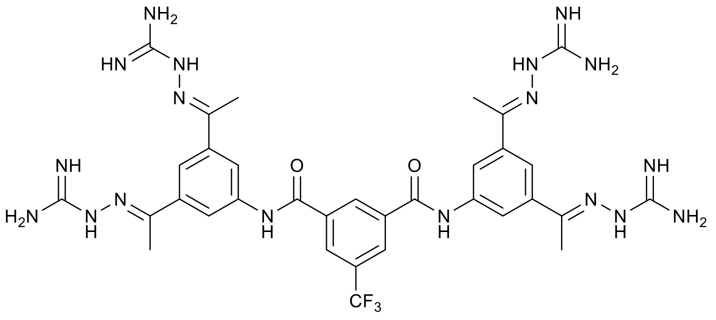

Starting material: 5-trifluoromethylisophthalic acid (33 mg).
Yield, over two steps: 28 mg (21 %).
MS $(M+H)^+$: 777.4.
$^1H$ NMR (300 MHz, DMSO-$d_6$) δ 10.81 (s, 2H), 8.96 (s, 1H), 8.57 (s, 2H), 8.38 (s, 4H), 8.24 (s, 4H), 8.03 (s, 2H), 7.46 (s, 16H), 2.37 (s, 12H).

**Synthesis of 27-1.**

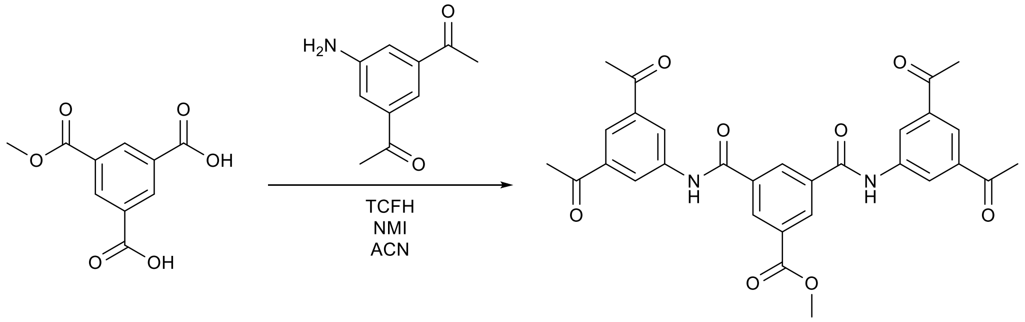


The 5-(methoxycarbonyl)benzene-1,3-dicarboxylic acid (250 mg, 1.1 mmol) and 3,5-diacetylaniline (400 mg, 2.3 mmol) were dissolved in 8 ml of MeCN. 0.47 mL N-methylimidazole and TCFH (640 mg, 2.2 mmol) were added sequentially. The reaction was stirred overnight. After the completion of the reaction the product was filtered and washed with ethanol to obtain intermediate 63 as white solid in 27% yield (150 mg).
MS $(M+H)^+$: 543.2.

$^{1}H$ NMR (300 MHz, DMSO) δ 11.01 (s, 2H), 8.94 (s, 1H), 8.81 (d, *J* = 1.8 Hz, 2H), 8.68 (d, *J* = 1.6 Hz, 4H), 8.27 (s, 2H), 3.99 (s, 3H), 2.69 (s, 12H).

**Synthesis of 28-1.**

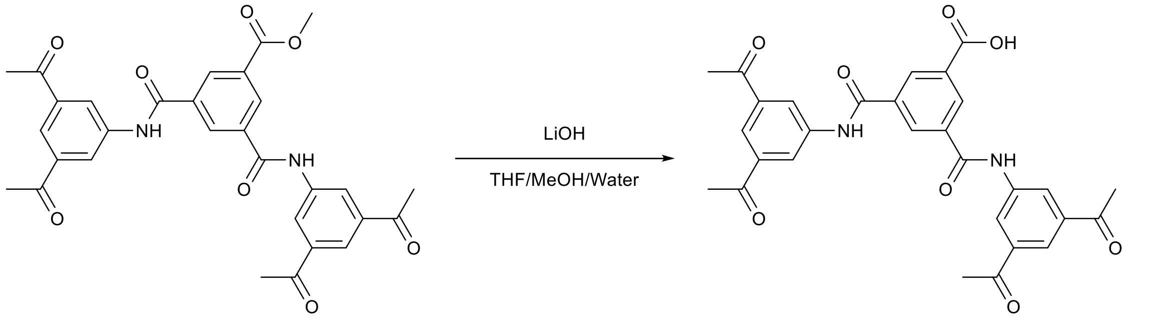


**27-1** (120 mg, 0.22 mmol) was dissolved in a mixture of THF/MeOH/Water (2 ml/2 ml/2 ml) and LiOH (300 mg) was added. The resulting mixture was stirred overnight. After completion of the reaction, 1 M HCl was added (5 ml) and solid was filtered and dried yielding **28-1** in 81% yield (90 mg).
MS $(M+H)^{+}$: 528.8.
$^{1}H$ NMR (300 MHz, DMSO) δ 11.02 (s, 1H), 8.91 (s, 1H), 8.81 (d, *J* = 1.7 Hz, 2H), 8.69 (d, *J* = 1.5 Hz, 4H), 8.26 (s, 2H), 2.68 (d, *J* = 2.1 Hz, 12H).

**Synthesis of 27-28.**

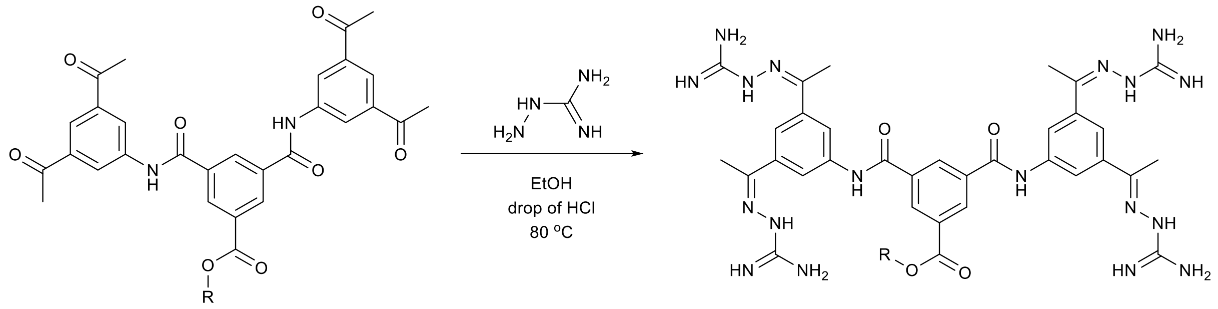


The compounds were synthesyzed according to general procedure 1 using 30 mg of respective tetraketone.

**27.**

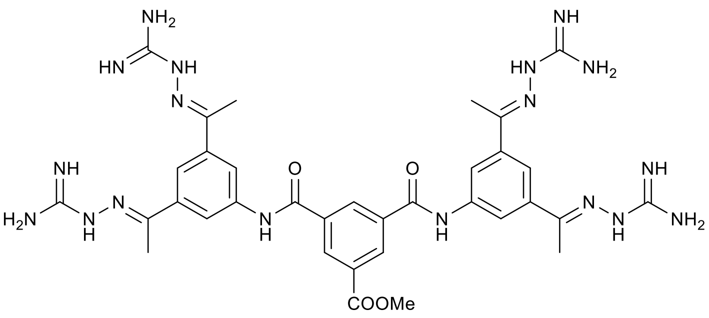

Starting material: **27-1**.
Yield: 31 mg (59%).
MS $(M+H)^{+}$: 767,9.
$^{1}H$ NMR (300 MHz, DMSO-$d_6$) δ 10.81 (s, 2H), 8.94 (s, 1H), 8.77 (d, J = 1.6 Hz, 2H), 8.40 (s, 4H), 8.26 (s, 4H), 8.05 (s, 2H), 7.71 (s, 16H), 3.98 (s, 3H), 2.38 (s, 12H).

**28.**

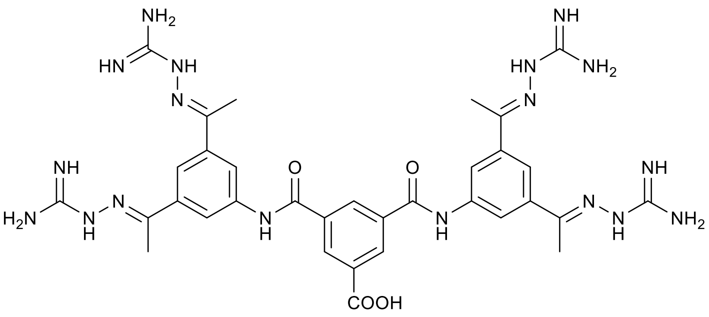

Starting material: **28-1**.
Yield: 9 mg (17%).
MS $(M+H)^+$: 753,7.
$^1$H NMR (300 MHz, DMSO-$d_6$) δ 10.64 (s, 2H), 8.83 (s, 2H), 8.75 (s, 1H), 8.38 (s, 4H), 8.33 (s, 4H), 8.09 (s, 2H), 7.71 (s, 16H), 2.44 (s, 12H).

**29.**

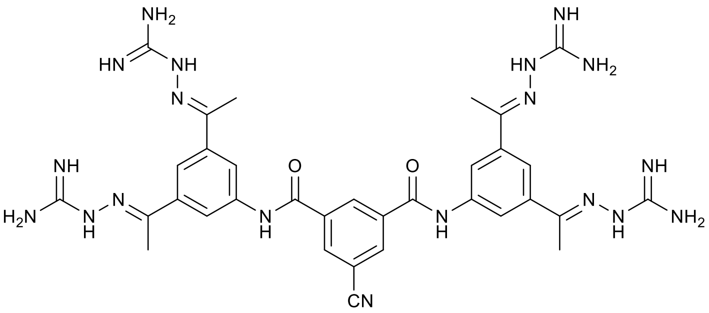

Starting material: 5-cyanoisophthalic acid (29 mg).
Yield, over two steps: 3 mg (2%).
MS $(M+H)^+$: 734.6.
$^1$H NMR (300 MHz, DMSO-$d_6$) δ 10.68 (s, 2H), 8.88 (s, 1H), 8.65 (s, 2H), 8.32 (s, 4H), 8.21 (s, 4H), 7.99 (s, 2H), 7.01 (s, 16H), 2.35 (s, 12H).

**30.**

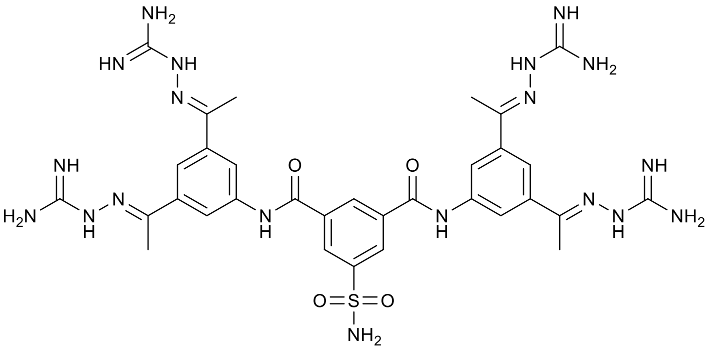

Starting material: 5-sulfamoylisophthalic acid (34 mg).
Yield, over two steps: 10 mg (7 %).
MS $(M+H)^+$: 788.9.
$^1$H NMR (300 MHz, DMSO-$d_6$) δ 10.88 (s, 2H), 8.95 (s, 1H), 8.65 (d, *J* = 1.5 Hz, 2H), 8.37 (s, 4H), 8.30 (s, 4H), 8.04 (s, 2H), 7.67 (s, 2H), 7.49 (s, 16H), 2.38 (s, 12H).

**31.**

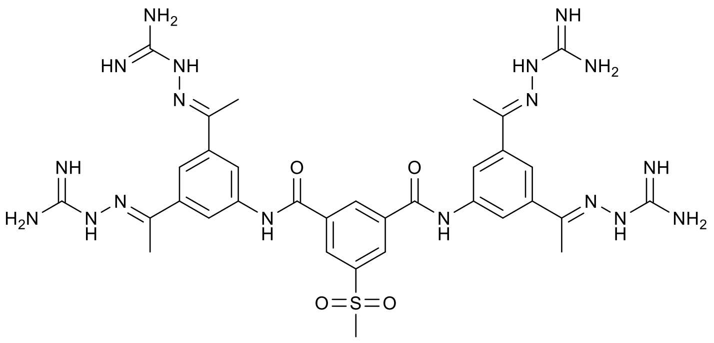

Starting material: 5-(methylsulfonyl)isophthalic acid (34 mg).
Yield, over two steps: 15 mg (11 %).
MS $(M+H)^+$: 787.8.
$^1$H NMR (300 MHz, DMSO-$d_6$) δ 10.85 (s, 2H), 8.98 (s, 1H), 8.73 (s, 2H), 8.33 (s, 4H), 8.24 (s, 4H), 8.01 (s, 2H), 7.18 (s, 16H), 3.40 (s, 3H), 2.36 (s, 12H).

**32.**

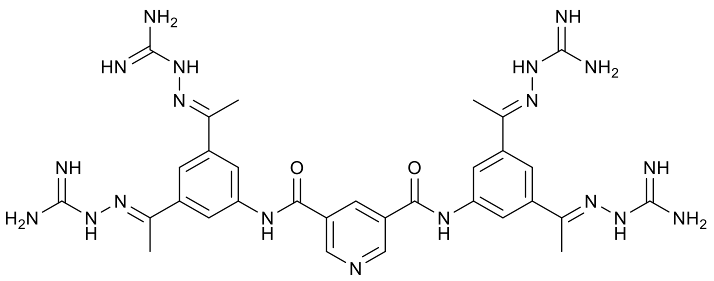

Starting material: pyridine-3,5-dicarboxylic acid (23 mg).
Yield, over two steps: 10 mg 8%.
MS $(M+H)^+$: 710.5.
$^1$H NMR (300 MHz, DMSO-$d_6$) δ 10.75 (s, 2H), 9.33 (s, 2H), 8.94 (s, 1H), 8.36 (s, 4H), 8.23 (s, 4H), 8.02 (s, 2H), 7.29 (s, 16H), 2.36 (s, 12H).

**33 (TCB-621).**

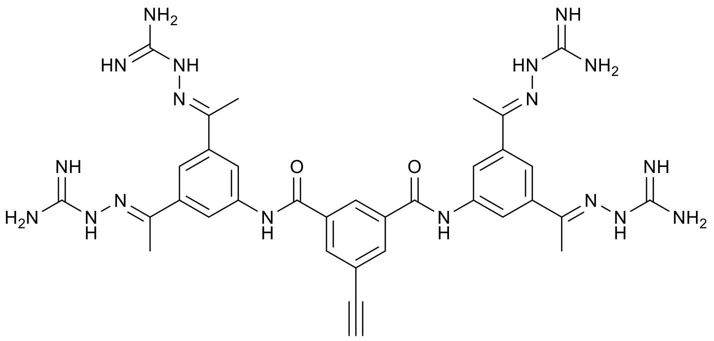

Starting material: 5-ethynylisophthalic acid (27 mg).
Yield, over two steps: 16 mg (12%).
MS $(M+H)^+$: 733.7.
$^1$H NMR (300 MHz, DMSO-$d_6$) δ 10.61 (s, 2H), 8.65 (s, 1H), 8.37 (s, 4H), 8.32 (s, 2H), 8.24 (s, 2H), 8.03 (s, 2H), 7.51 (s, 16H), 4.49 (s, 1H), 2.37 (s, 12H).
$^{13}$C NMR (101 MHz, DMSO-$d_6$) δ 167.3, 164.5, 157.9, 150.9, 139.2, 139.1, 136.0, 133.9, 128.2, 122.7, 119.9, 83.1, 79.7, 15.2.

**34.**

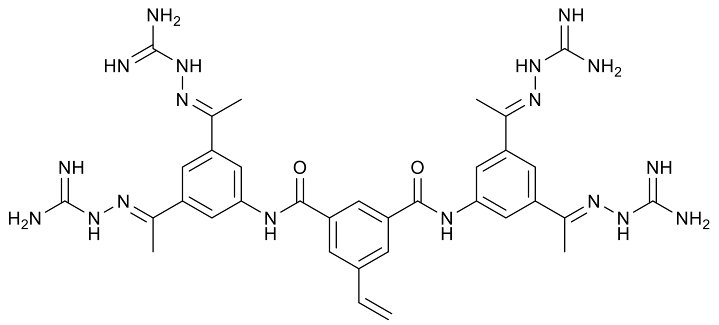

Starting material: 5-vinylisophthalic acid (27 mg).
Yield, over two steps: 28 mg (21%).
MS $(M+H)^+$: 735,8.
$^1$H NMR (300 MHz, DMSO-$d_6$) δ 10.61 (s, 2H), 8.54 (s, 1H), 8.39 (s, 4H), 8.31 (s, 2H), 8.27 (d, *J* = 1.6 Hz, 4H), 8.04 (s, 2H), 7.74 (s, 16H), 6.96 (dd, *J* = 17.6, 11.2 Hz, 1H), 6.13 (d, *J* = 17.6 Hz, 1H), 5.51 (d, *J* = 11.4 Hz, 1H), 2.38 (s, 12H).

#### Region IV

##### Synthesis of 35-37.

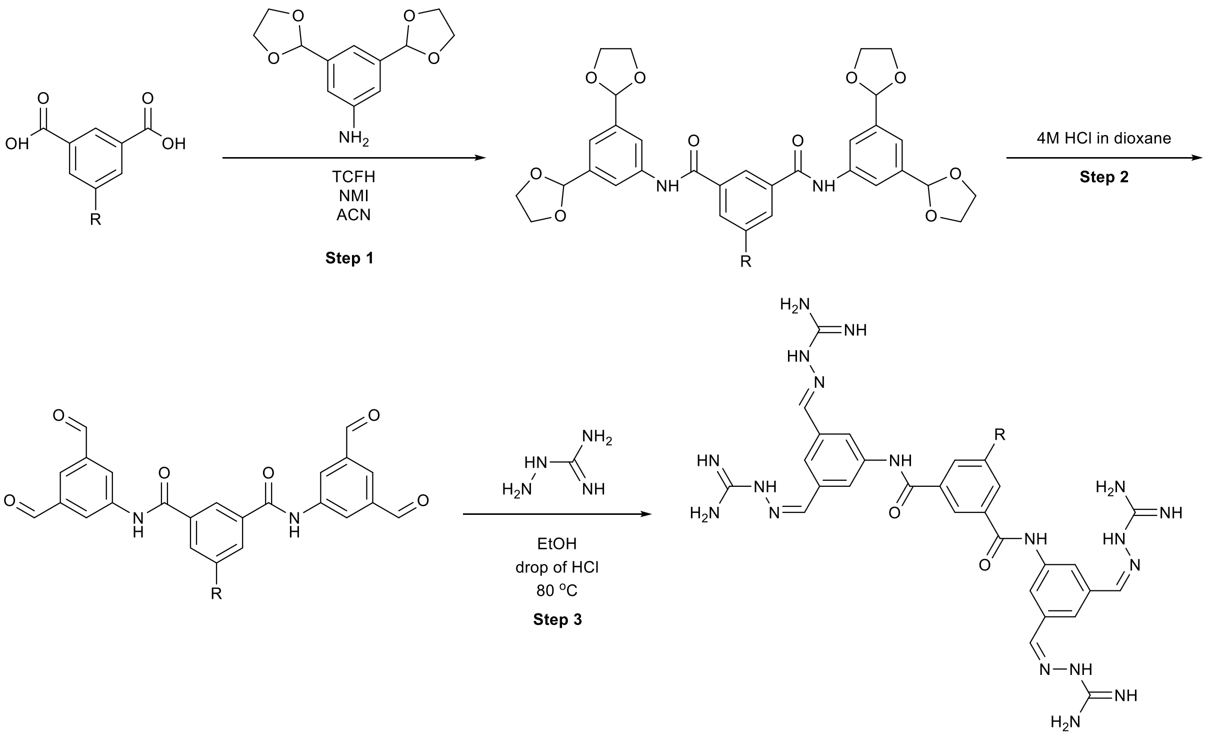


##### Step 1:

The acid (0.5 mmol) and 3,5-di(1,3-dioxolan-2-yl)aniline (237 mg, 1 mmol) were dissolved in 4 ml of MeCN. 0.24 mL *N*-methylimidazole and TCFH (300 mg, 1.07 mmol) were added sequentially. The reaction was stirred overnight. After the completion of the reaction, the solvent was evaporated and the residue was purified by silica gel column chromatography eluting with cyclohexane to cyclohexane:ethylacetate 0:100 to obtain the desired product as a light yellow oil.

##### Step 2:

The product from the previous step was dissolved in 4 M HCl in dioxane (3 ml) and stirred at room temperature for 3 h. After the completion of the reaction the solvents were evaporated, and the residue was used in the next steps without additional purification.

##### Step 3:

The compounds were synthesized according to general procedure using the product from the previous step.

**35.**

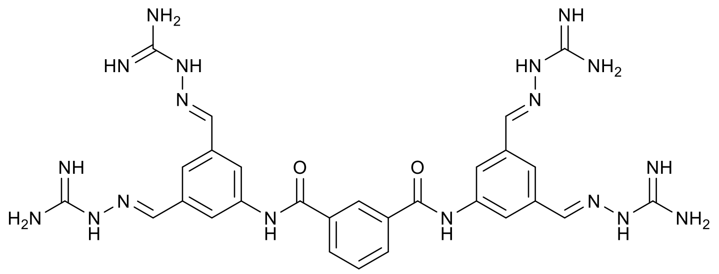

**Step 1:** Synthesis of **35-1**.
Starting material: isophthalic acid (88 mg).
MS $(M+H)^+$: 605,0.
$^1$H NMR (300 MHz, DMSO-$d_6$) δ 10.54 (s, 2H), 8.59 (t, *J* = 1.8 Hz, 1H), 8.17 (dd, *J* = 7.8, 1.8 Hz, 2H), 7.95 (d, *J* = 1.6 Hz, 4H), 7.70 (t, *J* = 7.8 Hz, 1H), 7.26 (s, 2H), 5.77 (s, 4H), 4.13 – 3.86 (m, 16H).

**Step 2:** Synthesis of **35-1**.
MS $(M+H)^+$: 429,4.
$^1$H NMR (300 MHz, DMSO-$d_6$) δ 11.06 (s, 2H), 10.14 (s, 4H), 8.75 (s, 1H), 8.71 (s, 4H), 8.26 (d, *J* = 7.5 Hz, 1H), 8.24 (s, 2H), 7.78 (t, *J* = 7.8 Hz, 1H).

**Step 3:** Synthesis of **35**.
Yield, over three steps: 20 mg (5%).
MS $(M+H)^+$: 653,8.
$^1$H NMR (300 MHz, DMSO-$d_6$) δ 10.59 (s, 2H), 8.64 (s, 1H), 8.37 (s, 4H), 8.24 – 8.17 (m, 2H), 8.11 (s, 4H), 8.07 (s, 6H), 7.74 (t, *J* = 7.8 Hz, 1H), 7.41 (s, 16H).

**36.**

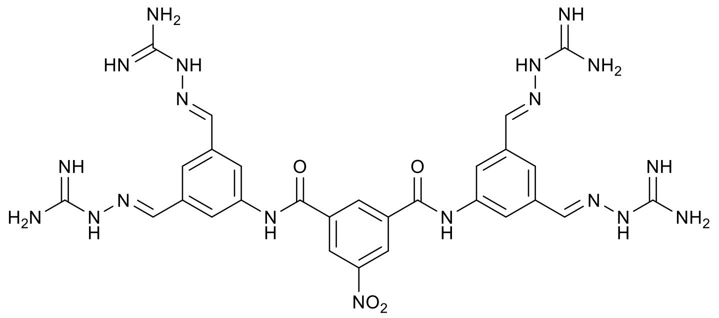

**Step 1:** Synthesis of **36-1**.
Starting material: 5-nitroisophtalic acid (102 mg).
MS $(M+H)^+$: 650.2.
$^1$H NMR (300 MHz, DMSO-$d_6$) δ 10.78 (s, 2H), 8.97 (s, 1H), 8.95 (s, 2H), 7.90 (d, *J* = 1.6 Hz, 4H), 7.23 (s, 2H), 5.72 (s, 4H), 4.06 – 3.82 (m, 16H).

**Step 2:** Synthesis of **36-2**.
MS $(M+H)^+$: 474.4.

**Step 3:** Synthesis of **36**.
Yield, over 3 steps: 15 mg (2%).
MS $(M+H)^+$: 698,2.

$^1$H NMR (300 MHz, DMSO-d$_6$) δ 10.94 (s, 2H), 9.09 (s, 1H), 9.05 (s, 2H), 8.36 (s, 4H), 8.12 (s, 3H), 8.09 (s, 2H), 8.08 (s, 4H), 7.37 (s, 16H).

**37.**

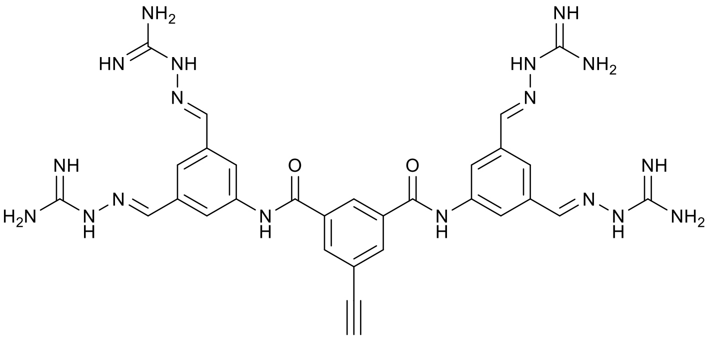


The molecule was done on scale of 80 mg of 3,5-di(1,3-dioxolan-2-yl)aniline (0.25 mmol).

**Step 1:** Synthesis of **37-1**.
Starting material: 5-ethynylisophthalic acid (23 mg, 0.125 mmol).
MS $(M+H)^+$: 629.6.
$^1$H NMR (300 MHz, DMSO-d$_6$) δ 10.53 (s, 2H), 8.52 (s, 1H), 8.22 (d, *J* = 1.8 Hz, 2H), 7.88 (d, *J* = 1.5 Hz, 4H), 7.20 (s, 2H), 5.70 (s, 4H), 4.40 (s, 1H), 4.14 – 3.81 (m, 16H).

**Step 2:** Synthesis of **37-2**.
MS $(M+H)^+$: 452,9.

**Step 3:** Synthesis of **37.**
Yield, over 3 steps: 6 mg (6%).
MS $(M+H)^+$: 677.7.
$^1$H NMR (300 MHz, DMSO-d$_6$) δ 10.67 (s, 2H), 8.64 (t, *J* = 1.7 Hz, 1H), 8.38 (s, 4H), 8.32 (d, *J* = 1.8 Hz, 2H), 8.11 (s, 2H), 8.10 (s, 4H), 8.07 (s, 2H), 7.57 (s, 4H), 4.49 (s, 1H).

**Synthesis of 38.**

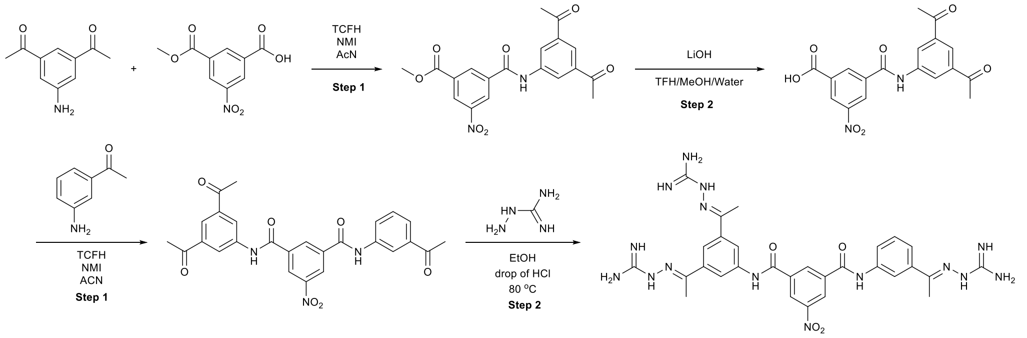


**Step 1**: Synthesis of methyl 3-((3,5-diacetylphenyl)carbamoyl)-5-nitrobenzoate.
The 5-Nitroisophthalic acid monomethyl ester (1.22 g, 5.6 mmol), intermediate 1 (1 g, 5.6 mmol) were dissolved in 20 ml of acetonitrile. *N*-methylimidazole (1.2 ml) and TCFH (1.46 g) were added sequentially in a single portion. The reaction was stirred overnight. After the completion of the reaction. The product was filtered and used in the next step without additional purification.
MS $(M+H)^+$: 384.9.

$^{1}H$ NMR (300 MHz, $CDCl_3$) δ 9.92 (s, 1H), 9.16 (t, *J* = 2.0 Hz, 1H), 8.99 (t, *J* = 1.6 Hz, 1H), 8.97 – 8.90 (m, 1H), 8.71 (d, *J* = 1.6 Hz, 2H), 8.30 (t, *J* = 1.5 Hz, 1H), 4.02 (s, 3H), 2.69 (s, 3H).
$^{13}C$ NMR (75 MHz, $CDCl_3$) δ 197.48, 164.28, 163.23, 139.16, 138.01, 136.43, 134.26, 132.38, 127.18, 126.81, 124.49, 124.07, 121.79, 53.10, 26.89.

**Step 2**: Synthesis of 3-((3,5-diacetylphenyl)carbamoyl)-5-nitrobenzoic acid.
The product from the previous step was dissolved in a mixture of THF/MeOH/Water (3 ml/3 ml/3 ml) and LiOH (800 mg) was added. The resulting mixture was stirred overnight. After completion of the reaction, 1 M HCl was added (5 ml) and the mixture was extracted with EtOAc. The organic layers were combined, dried over sodium sulfate and evaporated under reduced pressure yielding intermediate 56 as green solid.
Yield, over two steps: 1.13 g (55%).
MS $(M+H)^+$: 371.0.
$^{1}H$ NMR (300 MHz, DMSO-$d_6$) δ 11.06 (s, 1H), 9.06 (t, *J* = 1.9 Hz, 1H), 8.96 (d, *J* = 1.6 Hz, 1H), 8.78 (d, *J* = 1.5 Hz, 1H), 8.63 (d, *J* = 1.6 Hz, 2H), 8.25 (s, 1H), 2.68 (s, 6H).
$^{13}C$ NMR (75 MHz, DMSO-$d_6$) δ 197.13, 164.99, 162.75, 148.06, 139.42, 137.59, 136.04, 134.02, 132.91, 126.51, 126.28, 123.79, 123.65, 26.88.

**Step 3**: Synthesis of **38-3.**
3-((3,5-diacetylphenyl)carbamoyl)-5-nitrobenzoic acid (50 mg, 0.14 mmol) and 3-aminoacetophenone (19 mg, 0.14 mmol) were dissolved in 1 ml of ACN. 0.06 mL N-methylimidazole and TCFH (73 mg, 0.28 mmol) were added sequentially. The reaction was stirred overnight. After the completion of the reaction the product was filtered (if precipitated) to obtain the desired product.

**Step 4**: Synthesis of **38.**
The compound was synthesized according to general procedure 1 using the product from the previous step.
Yield, over two steps: 22 mg (20%).
MS $(M+H)^+$: 656.7.
$^{1}H$ NMR (300 MHz, DMSO-$d_6$) δ 10.93 (s, 1H), 10.88 (s, 1H), 9.08 (s, 1H), 9.01 (s, 2H), 8.38 (s, 3H), 8.25 (s, 2H), 8.17 (s, 1H), 8.05 (s, 1H), 7.91 (d, *J* = 8.0 Hz, 1H), 7.76 (d, *J* = 8.0 Hz, 1H), 7.61 (s, 12H), 7.42 (t, *J* = 8.0 Hz, 1H), 2.38 (s, 6H), 2.33 (s, 3H).

#### TCB-32 upscaling.

Step 1:
The isophthalic acid (450 mg, 2.8 mmol) and 3,5-acetylaniline (1000 mg, 5.6 mmol) were dissolved in 20 ml of MeCN. 1.2 mL N-methylimidazole and TCFH (1.46 g, 5.6 mmol) were added sequentially. The reaction was stirred overnight. After the completion of the reaction, the product was filtered to obtain the desired product as white solid.
Step 2:
The corresponding diamide from the previous step was dissolved in 40 ml of 96% ethanol, 100 mg of aminoguanidine hydrochloride and two drops of concentrated HCl were added. The reaction was heated to 80 °C and stirred at this temperature overnight. After the completion of the reaction the solvent was evaporated, and the residue was purified using reverse-phase chromatography (40 g cartridge, 10 runs)

eluting with water-acetonitrile with formic acid modifier. The resulting solution was lyophilized yielding the product as a solid tetraformate salt.
Yield, over two steps: 1050 mg (42%).

#### TCB-494 upscaling.

Step 1:
The 5-nitroisophthalic acid (590 mg, 2.8 mmol) and 3,5-acetylaniline (1000 mg, 5.6 mmol) were dissolved in 20 ml of MeCN. 1.2 mL N-methylimidazole and TCFH (1.46 g, 5.6 mmol) were added sequentially. The reaction was stirred overnight. After the completion of the reaction. The product was either filtered to obtain the desired product as white solid.
Step 2:
The corresponding diamide from the previous step was dissolved in 40 ml of 96% ethanol, 100 mg of aminoguanidine hydrochloride and two drops of concentrated HCl were added. The reaction was heated to 80 °C and stirred at this temperature overnight. After the completion of the reaction the solvent was evaporated, and the residue was purified using reverse-phase chromatography (120 g cartridge, two runs) eluting with water-acetonitrile with formic acid modifier. The resulting solution was lyophilized yielding the product as a solid tetraformate salt.
Yield, over two steps: 500 mg (19%).

#### Synthesis of SI-1.

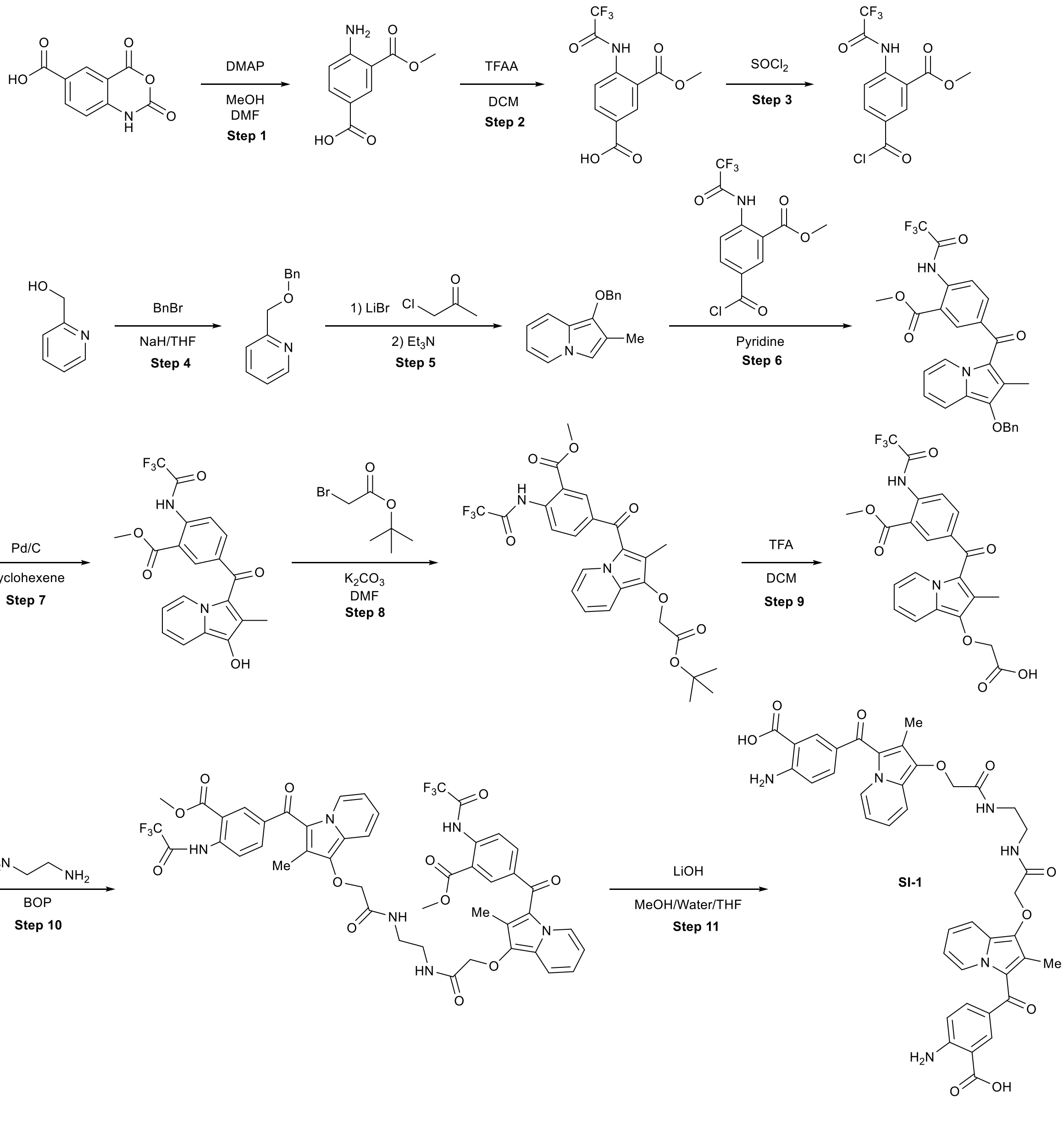


The synthesis was conducted according to [19] with additional modifications.

**Step 1**: Synthesis of 4-amino-3-(methoxycarbonyl)benzoic acid.

150 mg (1.2 mmol) of 4-dimethylaminopyridine are added to 2,4-dioxo-1,4-dihydro-2H-3, 1-benzoxazine-6-carboxylic acid (2.5 g, 12 mmol) in solution in 10 ml of dimethylformamide and 10 ml of methanol, and the mixture was heated at 60° C. for 3 hours. After the completion of the reaction, the solvents were evaporated under reduced pressure. The residue was taken up in water and extracted with ethyl acetate. The organic phase was washed with a saturated sodium chloride solution, dried over sodium sulphate and concentrated under reduced pressure yielding the product as white powder in 64 % yield (1500 mg).

MS $(M+H)^+$: 196.0.

$^1$H NMR (300 MHz, $CDCl_3$+DMSO) δ 11.93 (s, 1H), 8.41 (d, J = 2.1 Hz, 1H), 7.80 (dd, J = 8.8, 2.2 Hz, 1H), 6.90 – 6.70 (m, 3H), 3.82 (s, 3H).

**Step 2**: Synthesis of 3-(methoxycarbonyl)-4-(2,2,2-trifluoroacetamido)benzoic acid.

2 ml of trifluoroacetic anhydride are rapidly added to 4-amino-3-(methoxycarbonyl) benzoic acid (1500 mg, 7.6 mmol) in suspension in 25 ml of dichloromethane. The solution is stirred for 30 minutes at room temperature. The mixture was diluted with ethylacetate and extracted with water. The organic solvents were evaporated yielding the product as white solid in 94% yield (2.1 g).

The NMR looked like a mixture of compounds, but in HPLC only one peak of the product was observed. It was assumed that such observation is connected to the diastereomers of the amide bond, and no additional investigations were not conducted.

MS $(M+H)^+$: 291.8.
$^1$H NMR (300 MHz, $CDCl_3$) δ 13.21 (s, 1H), 12.43 (s, 1H), 8.81 – 8.65 (m, 2H), 8.32 – 8.17 (m, 1H), 3.97 (d, J = 6.9 Hz, 3H).

$^{13}$C NMR (75 MHz, $CDCl_3$) δ 168.2, 168.0, 167.0, 164.5, 155.7, 155.2, 142.6, 142.2, 136.1, 135.9, 133.0, 132.9, 127.0, 126.0, 120.4, 120.3, 117.4, 116.0, 115.9, 113.7, 53.1, 53.0.

**Step 3**: Synthesis of methyl 5-(chlorocarbonyl)-2-(2,2,2-trifluoroacetamido)benzoate.

1 ml of thionyl chloride and 1 drop of DMF were added to MF-6 suspension (1.5 g, 5.1 mmol) in 2 ml of DCM and the mixture was stirred for 90 minutes. If the clear solution did not form, more thionyl chloride was added and stirred for another 90 min. After the completion of the reaction the solvents were evaporated, and the residue was used further without additional purification. The yield was assumed as quantitative.

**Step 4**: Synthesis of 2-((benzyloxy)methyl)pyridine.

2-hydroxymethylpyridine (2.72 g, 25 mmol) was added portion-wise to the suspension of NaH (60% dispersion in oil, 4.1 g, 25 mmol)) in 50 ml of THF at 0°C. under nitrogen atmosphere, followed, after 10 minutes, benzylbromide (4.27 g, 25 mmol). The mixture was stirred at ambient temperature overnight. The reaction medium was carefully diluted with ethyl acetate and washed with water. The remaining NaH can lead to the extensive formation of gases. The organic phase is washed with a saturated aqueous solution of sodium chloride, dried over sodium sulphate, and concentrated under reduced pressure. The residue was purified by column chromatography 0-10% Cy-EtOAc yielding 2-((benzyloxy)methyl)pyridine as yellow oil in 68% yield (3.4 g).

MS $(M+H)^+$: 200.0.
$^1$H NMR (300 MHz, $CDCl_3$) δ 8.51 – 8.42 (m, 1H), 7.63 (td, J = 7.6, 1.8 Hz, 1H), 7.43 (d, J = 7.8 Hz, 1H), 7.35 – 7.17 (m, 5H), 7.12 (dd, J = 8.2, 5.5 Hz, 1H), 4.63 (s, 2H), 4.58 (s, 2H).

$^{13}$C NMR (75 MHz, $CDCl_3$) δ 158.4, 148.9, 137.9, 136.7, 128.4, 127.8, 127.7, 122.4, 121.4, 73.0, 72.9.

**Step 5:** Synthesis of 1-(benzyloxy)-2-methylindolizine.

The mixture of 2-((benzyloxy)methyl)pyridine (3.4 g, 17 mmol), 3 ml of chloroacetone, and 3 g of lithium bromide in 30 ml of acetonitrile is refluxed for 16 hours. Then 10 ml of water and 2.34 g of $K_2CO_3$ were added. The mixture was refluxed overnight._The pyridinium salt was formed. Then 4 ml of triethylamine was added. After refluxing for 3 hours, the mixture was poured into water and the mixture was extracted with ethyl acetate. The colour is very dark, the separation of phases is difficult, and special care should be taken. The organic solvents were evaporated, and the residue was purified by column chromatography 0-20% (Cy-EtOAc) yielding 1-(benzyloxy)-2-methylindolizine as a brown oil in 33% yield (1.33 g).

MS $(M+H)^+$: 256.0. (intermediate quaternaty salt)
MS $(M+H)^+$: 237.9. (indolizine)
$^1$H NMR (300 MHz, $CDCl_3$) δ 7.83 – 7.10 (m, 7H), 6.93 (s, 1H), 6.47 (t, J = 7.8 Hz, 1H), 6.29 (t, J = 6.7 Hz, 1H), 5.02 (s, 2H), 2.22 (s, 3H).

$^{13}$C NMR (75 MHz, $CDCl_3$) δ 138.0, 134.4, 128.3, 128.3, 127.9, 124.2, 122.8, 116.5, 115.8, 114.7, 109.1, 107.6, 29.7, 9.1.

**Step 6:** Synthesis of methyl 5-(1-(benzyloxy)-2-methylindolizine-3-carbonyl)-2-(2,2,2-trifluoroacetamido)benzoate.

1 ml of pyridine and 1-(benzyloxy)-2-methylindolizine (4.9 mmol) were added to the solution of product from step 3 (1.6 g. 5.17 mmol**)** in 40 ml of DCM. The reaction medium that has turned greenish was stirred overnight. It was concentrated to dryness and the residue was purified by column chromatography using 0-100% Cy-EtOAc as the eluent to give the product as an orange solid in 41% yield (1.05 g).

MS $(M+H)^+$: 511.1.
$^1$H NMR (300 MHz, DMSO-$d_6$) δ 11.90 (s, 1H), 9.54 (d, *J* = 7.1 Hz, 1H), 8.13 (d, *J* = 8.5 Hz, 1H), 8.04 (d, *J* = 2.2 Hz, 1H), 7.82 (dd, *J* = 8.4, 2.1 Hz, 1H), 7.50 (d, *J* = 8.8 Hz, 1H), 7.21 – 7.10 (m, 5H), 6.91 (t, *J* = 7.7 Hz, 1H), 4.93 (s, 2H), 3.82 (s, 3H), 1.62 (s, 3H).

$^{13}$C NMR (75 MHz, DMSO-$d_6$) δ 182.1, 166.7, 146.7, 138.5, 137.9, 137.0, 136.2, 133.7, 130.5, 130.3, 128.7, 128.3, 128.2, 127.2, 124.2, 124.0, 122.7, 120.4, 116.8, 115.3, 114.0, 76.2, 52.9, 11.0.

**Step 7**. Synthesis of methyl 5-(1-hydroxy-2-methylindolizine-3-carbonyl)-2-(2,2,2-trifluoroacetamido)benzoate.

1.9 ml of cyclohexene were added to 1g of the product from step 6 (1 g, 1.95 mmol) in 30 ml of ethanol, in the presence of 150 mg of 10% Pd/C, and the medium is heated under reflux for one hour. The reaction medium is cooled to room temperature, the solvent evaporated under the reduced pressure and the residue was purified using column chromatography column chromatography using 0-100% Cy-EtOAc as the eluent to give the product as orange solid in 62% yield (510 mg).

MS $(M+H)^+$: 420.9.

$^{1}H$ NMR (300 MHz, DMSO-$d_6$) δ 11.90 (s, 1H), 9.56 (d, J = 7.1 Hz, 1H), 8.75 (s, 1H), 8.13 (d, J = 8.5 Hz, 1H), 8.02 (s, 1H), 7.82 (dd, J = 8.4, 2.2 Hz, 1H), 7.60 (d, J = 8.8 Hz, 1H), 7.16 – 7.05 (m, 1H), 6.84 (t, J = 6.9 Hz, 1H), 3.81 (s, 3H), 1.67 (s, 3H).

$^{13}C$ NMR (75 MHz, DMSO-$d_6$) δ 181.3, 166.7, 154.9, 138.9, 137.7, 134.9, 133.6, 130.5, 128.3, 127.0, 122.9, 122.7, 120.3, 118.4, 117.5, 116.7, 115.6, 113.6, 52.9, 11.0.

**Step 8:** Synthesis of methyl 5-(1-(2-(tert-butoxy)-2-oxoethoxy)-2-methylindolizine-3-carbonyl)-2-(2,2,2-trifluoroacetamido)benzoate

In a 30 mL microwave vial, products from step 7 (500 mg, 1.21 mmol) and $K_2CO_3$ (340 mg, 2.4 mmol) were added to 10 ml of DMF. The cap was sealed. *Tert*-butyl 2-bromoacetate (540 mg, 0.4 ml, 0.28 mmol) was added and the reaction was stirred at RT for two hours. The solvent was evaporated under the reduced pressure, and the residue was purified using column chromatography using 0-100% Cy-EtOAc as the eluent to give yielding the product as a yellow-green solid in 50% yield (320 mg).

MS $(M+H)^+$: 535.0.

$^{1}H$ NMR (300 MHz, DMSO-$d_6$) δ 11.97 (s, 1H), 9.60 (d, *J* = 7.1 Hz, 1H), 8.20 (d, *J* = 8.5 Hz, 1H), 8.13 (s, 1H), 7.92 (d, *J* = 10.6 Hz, 1H), 7.78 (d, *J* = 10.2 Hz, 1H), 7.39 – 7.21 (m, 1H), 7.00 (t, *J* = 7.7 Hz, 1H), 4.57 (s, 2H), 3.88 (s, 3H), 1.83 (s, 3H), 1.42 (s, 9H).

**Step 9**. Synthesis of 2-((3-(3-(methoxycarbonyl)-4-(2,2,2-trifluoroacetamido)benzoyl)-2-methylindolizin-1-yl)oxy)acetic acid.

Product from step 8 (300 mg, 0.56 mmol) was dissolved in 20 ml of 50-50 mixture of DCM/TFA and stirred at room temperature for 3 h. After the completion of the reaction, the solvents were evaporated yielding the product as a brown solid in 59% yield (160 mg).

MS $(M+H)^+$: 479.0.

$^{1}H$ NMR (300 MHz, DMSO-$d_6$) δ 11.98 (s, 1H), 9.61 (d, J = 7.1 Hz, 1H), 8.20 (d, J = 8.5 Hz, 1H), 8.14 (d, J = 2.2 Hz, 1H), 7.93 (dd, J = 8.4, 2.1 Hz, 1H), 7.79 (d, J = 8.9 Hz, 1H), 7.35 – 7.20 (m, 1H), 7.00 (t, J = 7.0 Hz, 1H), 4.60 (s, 2H), 3.89 (s, 3H), 1.83 (s, 3H).

**Steps 10-11:** Synthesis of SI-1.

0.12 ml of triethylamine and BOP (270 mg, 0.45 mmol) are added to the solution of the product from step 9 (150 mg, 0.31 mmol) in 10 ml of DMF. After 15 minutes of ethylenediamine (8 mg, 0.15 mmol) in 1 ml of DMF was added. The reaction medium was stirred at room temperature overnight. The solvents were evaporated, and residue was dissolved in a mixture of THF/MeOH/Water (2/2/2 ml). 200 mg of LiOH was added and the mixture was stirred overnight. After the completion of the reaction the solvents were evaporated, and the residue was purified by RP-HPLC eluting with 0-100% Water/Acetonitrile with formic acid as a modifier. The resulting solution was lyophilized yielding SI-1 as yellow solid in 6% yield (7 mg).

MS $(M+H)^+$: 760.9.

$^1H$ NMR (300 MHz, DMSO-$d_6$) δ 9.18 (d, J = 7.3 Hz, 2H), 8.43 – 8.31 (m, 2H), 8.04 (d, J = 2.2 Hz, 2H), 7.63 (d, J = 8.9 Hz, 2H), 7.56 (dd, J = 8.7, 2.3 Hz, 2H), 7.02 (dd, J = 9.3, 6.2 Hz, 2H), 6.92 – 6.73 (m, 4H), 4.41 (s, 6H), 3.34 (s, 4H), 1.99 (s, 6H).

**Synthesis of molecules from screening.**

**VS-1.**

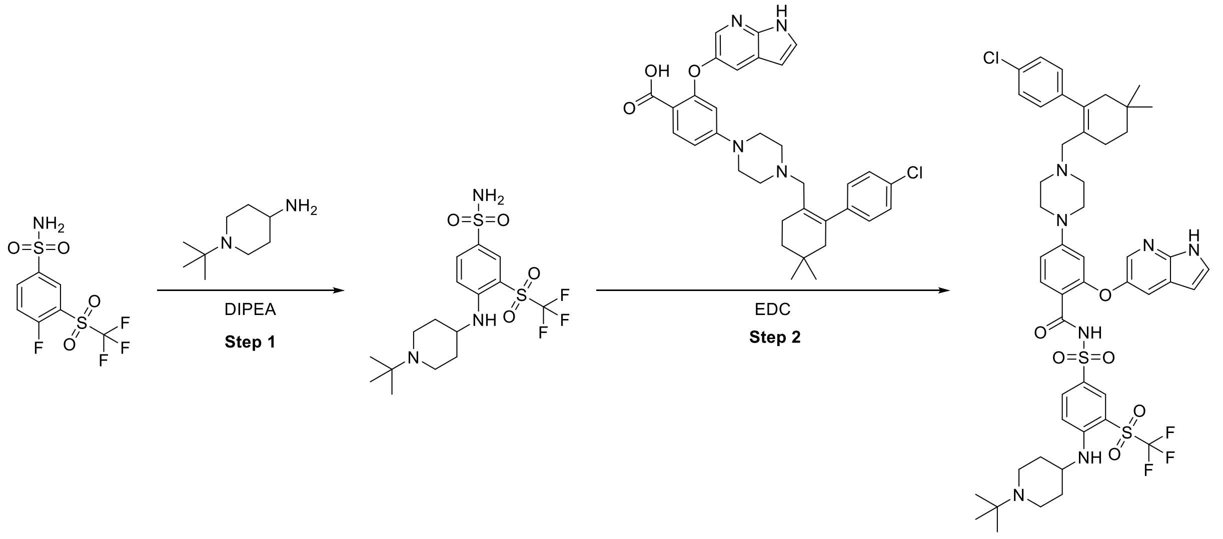


The synthesis was conducted according to [20]

**Step 1. Synthesis of VS-1-1.**

To a mixture of 4-fluoro-3-(trifluoromethylsulfonyl)benzenesulfonamide (307 mg) and 1-tert-Butylpiperidin-4-amine (156 mg) in tetrahydrofuran (4 ml) was added Hunig's Base (1 ml). The mixture was stirred overnight. The mixture was diluted with ethyl acetate (100 ml) and water (300 ml) . The layers were separated and the organic phase was washed with water, brine and dried over $Na_2SO_4$. After filtration, the solvent was evaporated to provide the title compound. Yield 440 mg (99%).

MS $(M+H)^+$: 444.0.

$^1$H NMR (300 MHz, $CDCl_3$) δ 8.05 (d, J = 2.3 Hz, 1H), 7.79 (dd, J = 9.2, 2.3 Hz, 1H), 6.72 – 6.56 (m, 2H), 6.44 (s, 2H), 3.32 (d, J = 7.4 Hz, 1H), 2.83 – 2.63 (m, 2H), 2.28 – 2.09 (m, 2H), 1.86 (m, 2H), 1.50 – 1.30 (m, 2H), 0.91 (s, 9H).

$^{13}$C NMR (75 MHz, $CDCl_3$) δ 150.3, 147.0, 135.5, 132.3, 130.4, 121.9, 117.5, 113.0, 107.5, 59.8, 53.9, 53.15, 49.0, 43.5, 31.4, 25.6, 20.6, 13.7.

**Step 1. Synthesis of VS-1.**

2-(1H-Pyrrolo[2,3-b]py-5-yloxy)-4-(4-(4'-Cl-5,5-diMe-3,4,5,6-tetrahydro-1,1'-biPh-2-yl)methylpiperazin-1-yl)benzoic acid (120 mg), VS-1-1 (92 mg), l-ethyl-3-[3-(dimethylamino)propyl]carbodiimide hydrochloride (EDC hydrochloride) (82 mg), and 4-dimethylaminopyridine (26 mg) were stirred in $CH_2Cl_2$ (3 ml) for 24 hours. After the end of the reaction, it was chromatographed on silica gel with 0-10% DCM-MeOH and afterwards by RP-HPLC eluting with 0-100% Water/Acetonitrile with formic acid as a modifier. The resulting solution was lyophilized yielding VS-1 as solid in 19% yield (40 mg).

MS $(M+H)^+$: 996.2.

$^1H$ NMR (300 MHz, CDCl3) δ 10.92 (s, 1H), 8.24 (d, J = 2.3 Hz, 1H), 8.01 (q, J = 2.2 Hz, 2H), 7.75 (d, J = 9.1 Hz, 1H), 7.50 (d, J = 2.6 Hz, 1H), 7.31 – 7.26 (m, 1H), 7.06 (d, J = 8.4 Hz, 2H), 6.79 (m, 3H), 6.66 (d, J = 9.5 Hz, 1H), 6.42 – 6.35 (m, 1H), 6.35 – 6.29 (m, 1H), 5.84 (d, J = 2.5 Hz, 1H), 3.37 (d, J = 7.4 Hz, 1H), 3.01 – 2.69 (m, 6H), 2.58 (s, 2H), 2.16 – 1.74 (m, 11H), 1.56 – 1.36 (m, 2H), 1.33 – 1.17 (m, 3H), 0.95 (s, 9H), 0.78 (s, 7H).

13C NMR (75 MHz, CDCl3) δ 162.0, 159.2, 155.2, 151.1, 146.3, 144.4, 141.8, 137.6, 136.0, 135.3, 134.7, 133.2, 131.2, 129.4, 128.8, 127.8, 127.4, 125.9, 121.9, 120.2, 119.8, 117.6, 112.5, 108.6, 108.4, 108.0, 100.2, 60.0, 53.2, 51.8, 49.1, 46.5, 43.5, 34.9, 31.4, 29.3, 28.8, 27.8, 26.5, 25.6, 25.2.

**VS-2.**

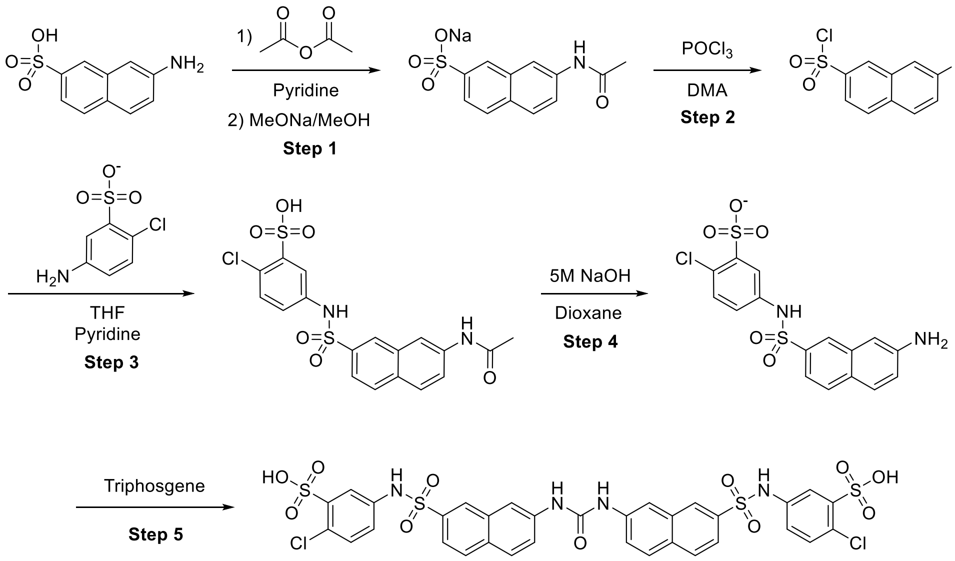


The synthesis was conducted according to[21] with additional modifications.

**Step 1**: Synthesis of sodium 7-acetamidonaphthalene-2-sulfonate.

The sodium salt of starting material (7-Amino-2-naphthalenesulfonate) (10.21g, 45 mmol) was dissolved in hot 1M NaOH (500ml), filtered, precipitated by acidifying with HCl (conc) and filtered again.

Purified starting material (5.06g. 22.5 mmol) were slowly added (ca. 15 min) to 30ml of a 1:1 mixture of pyridine and acetic anhydride while cooling with an ice bath and afterwards stirred overnight. Afterwards, the mixture was poured into a $Et_2O$/THF mixture (200ml/100ml), decanted, and the sticky oil was dried over the weekend. The resulted solid was dissolved in 30 of dry MeOH, and NaOMe (1.85 g) in 20 ml of MeOH was added to it. The precipitated solid was filtered and dried, yielding the product as white solid in 72% yield (4.67 g).

$^1H$ NMR (300 MHz, DMSO-$d_6$) δ 10.16 (s, 1H), 8.22 – 8.13 (m, 1H), 7.97 (d, *J* = 1.6 Hz, 1H), 7.83 (d, *J* = 8.9 Hz, 1H), 7.77 (d, *J* = 8.5 Hz, 1H), 7.67 (dd, *J* = 8.8, 2.1 Hz, 1H), 7.59 (dd, *J* = 8.5, 1.6 Hz, 1H), 2.50 (p, *J* = 1.8 Hz, 4H), 2.10 (s, 3H).

$^{13}C$ NMR (75 MHz, DMSO-$d_6$) δ 168.6, 146.7, 146.0, 137.4, 132.5, 129.3, 128.0, 127.0, 123.4, 122.5, 120.4, 115.5, 24.1.

**Step 2**: Synthesis of 7-acetamidonaphthalene-2-sulfonyl chloride.

sodium 7-acetamidonaphthalene-2-sulfonate (1g, 3.48 mmol) was suspended in 10ml of $POCl_3$ at 0 °C and 0.5 ml of dimethylacetamide was added dropwise. Afterwards, the mixture was stirred overnight and poured into ice and was left in fridge overnight. The precipitate was filtered, affording the product as off-white solid in 70% yield (0.69 g), which was used in the next step without additional purification.

**Step 3.** Synthesis of 5-((7-acetamidonaphthalene)-2-sulfonamido)-2-chlorobenzenesulfonic acid.

4- chloroaniline-3 sulfonic acid (178 mg, 0.86 mmol) was suspended in THF (2ml), Pyridine (0.45 ml) was added to the suspension, and the mixture was cooled to 0°C. Product from the previous step (200 mg, 0.71 mmol) was added portion wise, and the mixture was stirred overnight. Afterwards, the solvents were evaporated, and 1M HCl (5ml) was added. This solution was extracted with EtOAc several times, dried over MgSO4 and concentrated, yielding the product in 10% yield (32 mg).

MS (M-H): 453.0.

**Step 4-5**. Synthesis of VS-2.

NaOH solution (5M, 1.05ml) and Dioxane (0.05 ml) were added, and stirred at 55°C overnight. Afterwards, the mixture was filtered through PTFE syringe filter, conc. $H_2SO_4$ was slowly added until solid precipitated. The liquids were pipetted, and the product was dried. The resulted solid was dissolved/suspended in acetate buffer (1M, pH ca. 4.7, 0.41 ml), THF (0.25 ml) and NaOH solution (5M, 0.016 ml), was added in several portions over the course of 24 h until no starting material was observed in HPLC. The solvents were evaporated, and the residue was purified by RP-HPLC eluting with 0-100% Water/Acetonitrile with formic acid as a modifier. The resulting solution was lyophilized yielding VS-2 as solid in 33% yield (9.8 mg).

MS (M-H): 848.8.

$^1$H NMR (300 MHz, DMSO-d6) δ 10.47 (s, 2H), 9.21 (s, 2H), 8.30 (d, *J* = 1.8 Hz, 2H), 8.18 (d, *J* = 2.1 Hz, 2H), 7.99 (d, *J* = 8.7 Hz, 2H), 7.94 (d, *J* = 9.0 Hz, 2H), 7.81 (dd, *J* = 8.9, 2.2 Hz, 2H), 7.71 (d, *J* = 2.7 Hz, 2H), 7.59 (dd, *J* = 8.6, 1.9 Hz, 2H), 7.21 (d, *J* = 8.6 Hz, 2H), 7.11 (dd, *J* = 8.6, 2.7 Hz, 2H).

**VS-3.**

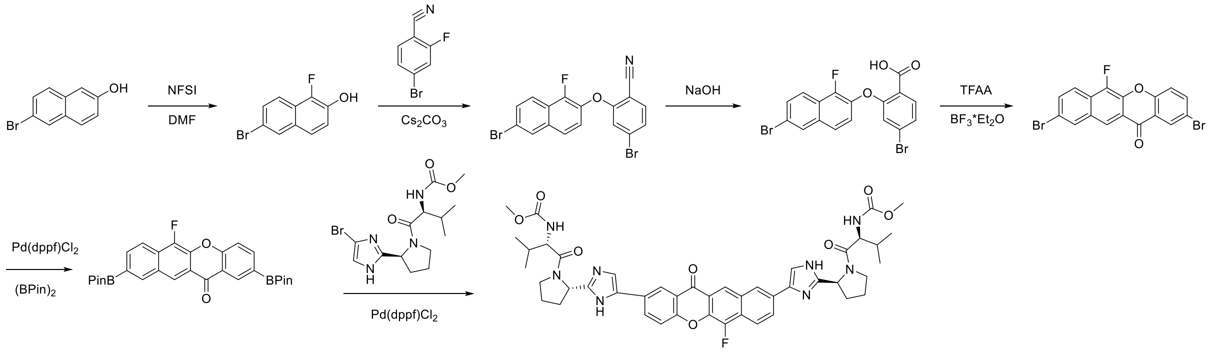

The synthesis was conducted according to[21] with additional modifications.

**Step 1:** Synthesis of 6-bromo-1-fluoronaphthalen-2-ol.

6-bromonaphthalen-2-ol (5 g, 22.41 mmol) was dissolved in DMF (50 mL) in a 250 mL flask. N-fluorobenzenesulfonimide (21.20 g, 67.2 mmol) was added, and the solution was stirred at 25°C over weekend. The solution was concentrated. The residue was purified by column chromatography on silica gel, eluting with cyclohexane and EtOAc (0% to 100%) to give 6-bromo-l-fluoronaphthalen-2-ol as brown solid in 59% yield (3.20 g).

MS was not observed in ESI.

$^1$H NMR (301 MHz, $CDCl_3$) δ 7.67 – 7.54 (m, 2H), 7.46 (d, J = 1.5 Hz, 1H), 7.30 (d, J = 10.2 Hz, 1H), 6.20 (dt, J = 10.0, 2.7 Hz, 1H).

$^{19}$F NMR (283 MHz, $CDCl_3$) δ -101.14 (s, 1F).

$^{13}$C NMR (75 MHz, $CDCl_3$) δ 187.1, 186.7, 186.4, 144.08, 144.06, 144.03, 133.8, 133.7, 133.7, 132.6, 129.2, 129.2, 129.1, 126.6, 126.5, 124.6, 124.6, 108.5, 105.3, 102.0.

**Step 2.** Synthesis of 4-bromo-2-((6-bromo-1-fluoronaphthalen-2-yl)oxy)benzonitrile.

5-bromo-2-fluorobenzonitrile (2.3 g, 11.5 mmol), 6-bromonaphthalen-2-ol (3.15 g, 11.4 mmol), and $Cs_2CO_3$ (7.47 g, 22.9 mmol) were added into a 500 ML flask. DMF (115 ml) was added. The solution was stirred at 80°C overnight. EtOAc (150 mL) and water (100 mL) were added. The organic layer was separated and washed with water (100 mL x 2), brine, dried over anhydrous $Na_2SO_4$, filtered, and concentrated. The product was purified by silica chromatography eluting with cyclohexane/EtOAc 0% to 50% yielding the product as off-white solid in 41 % (3.3 g of mixture).

After purification, there was a 30% impurity of the starting 5-bromo-2-fluorobenzonitrile, but the subsequent reactions were carried out without additional purification.

MS was not observed in ESI.

$^1$H NMR (300 MHz, $CDCl_3$) δ 7.98 (d, *J* = 1.9 Hz, 1H), 7.89 (d, *J* = 8.9 Hz, 1H), 7.78 – 7.43 (m, 3H), 7.25 (dd, *J* = 9.0, 7.5 Hz, 1H), 7.06 (t, *J* = 8.6 Hz, 1H), 6.59 (d, *J* = 9.1 Hz, 1H).

$^{19}$F NMR (283 MHz, $CDCl_3$) δ -108.21 (s, 1F).

**Step 3.** Synthesis of 4-bromo-2-((6-bromo-1-fluoronaphthalen-2-yl)oxy)benzoic acid.

Product from the previous step was dissolved (3.3 g) was dissolved in ethanol (40 ml). NaOH (3.1 g in 31 ml of water) was added. The solution was stirred at 85°C for overnight. The solution was neutralized by 100 ml of 6M HCl and extracted with ethylacetate. The solvents were evaporated, affording the product (1.5 g, 45%) with

30% mixture of 5-bromo-2-fluorobenzoic acid, which was used in the next step without additional purification.

MS $(M-H)^-$: 438.9.

$^1$H NMR was difficult to access due to impurity.

$^{19}$F NMR (283 MHz, DMSO-$d_6$) δ -112.43 (s, 1F).

**Step 4.** Synthesis of 2,9-dibromo-6-fluoro-12H-benzo[b]xanthen-12-one.

Product from the previous step (1.5 g) and $CH_2Cl_2$ (18 ml) were added into a flask and cooled to 0°C. TFAA (0.6 ml, 4.2 mmol) was added via syringe and the solution was stirred at 0°C for 30 min. Then $BF_3$*$OEt_2$ (0.09 ml) was added via Hamilton syringe and the solution was stirred at 0°C for 30 min, then at 25°C overnight. A suspension was formed. The solid was collected by filtration to give the product as yellow solid in 50% yield (500 mg, calculated to the amount of the SM in the mixture).

MS was not observed in ESI.

$^1$H NMR (301 MHz, DMSO-$d_6$) δ 8.79 (s, 1H), 8.72 (s, 1H), 8.32 (d, J = 2.5 Hz, 1H), 8.17 (d, J = 9.2 Hz, 1H), 8.10 (dd, J = 8.9, 2.6 Hz, 1H), 7.97 (s, 2H), 7.78 (d, J = 8.9 Hz, 1H).

$^{19}$F NMR (283 MHz, DMSO-$d_6$) δ -143.29 (s, 1F).

**Step 5**: Synthesis of 6-fluoro-2,9-bis(4,4,5,5-tetramethyl-1,3,2-dioxaborolan-2-yl)-12H-benzo[b]xanthen-12-one

The mixture of the product from previous step (150 mg, 0.355 mmol), bis(pinacolato)diboron (225 mg, 0.9 mmol), KOAc (200 mg, 2 mmol) and Pd(dppf)$Cl_2$ (60 mg, 0.081 mmol) in dioxane (30 mL) was stirred at 100 $^oC$ overnight under nitrogen. The residue was purified by column chromatography on silica gel, eluting with cyclohexane and EtOAc (0% to 100%) 32% yield (60 mg).

MS was not observed in ESI.

$^1$H NMR (300 MHz, $CDCl_3$) δ 8.77 (d, J = 1.6 Hz, 1H), 8.71 (d, J = 1.8 Hz, 1H), 8.50 (s, 1H), 8.10 (d, J = 8.4 Hz, 2H), 7.96 (d, J = 8.5 Hz, 1H), 7.51 (d, J = 8.4 Hz, 1H), 1.34 (s, 12H), 1.31 (s, 12H).

$^{19}$F NMR (283 MHz, $CDCl_3$) δ -144.37 (s, 1F).

**Step 6**: synthesis of VS-3.

The mixture of MF-62 (50 mg, 0.1 mmol), methyl ((S)-1-((S)-2-(4-bromo-1H-imidazol-2-yl)pyrrolidin-1-yl)-3-methyl-1-oxobutan-2-yl)carbamate (82 mg, 0.22 mmol), $Na_2CO_3$ (53 mg, 0.5 mmol) and Pd(dppf)$Cl_2$ (11 mg, 0.015 mmol) in THF/$H_2O$/DMF (v/v=5/2/1, 3 mL) was stirred at 80°C overnight under $N_2$ protection. After that, the mixture was washed with water and extracted with ethyl acetate, washed with brine and dried over anhydrous sodium sulfate. After filtrated, the filtrate was concentrated in vacuum, the residue was dissolved in DMSO and purified by RP-HPLC yielding 1 mg of the product (1% yield).

MS $(M+H)^+$: 849.5.

$^1H$ NMR (300 MHz, DMSO-$d_6$) δ 11.93 (d, J = 10.7 Hz, 2H), 8.64 (s, 1H), 8.56 (d, J = 2.5 Hz, 2H), 8.28 – 8.11 (m, 3H), 7.77 – 7.65 (m, 3H), 7.30 (m, 2H), 5.12 (brs, 2H), 4.10 (t, J = 8.2 Hz, 2H), 3.84 (brs, 4H), 3.55 (s, 6H), 2.31 – 1.79 (m, 10H), 1.01 – 0.75 (m, 12H).

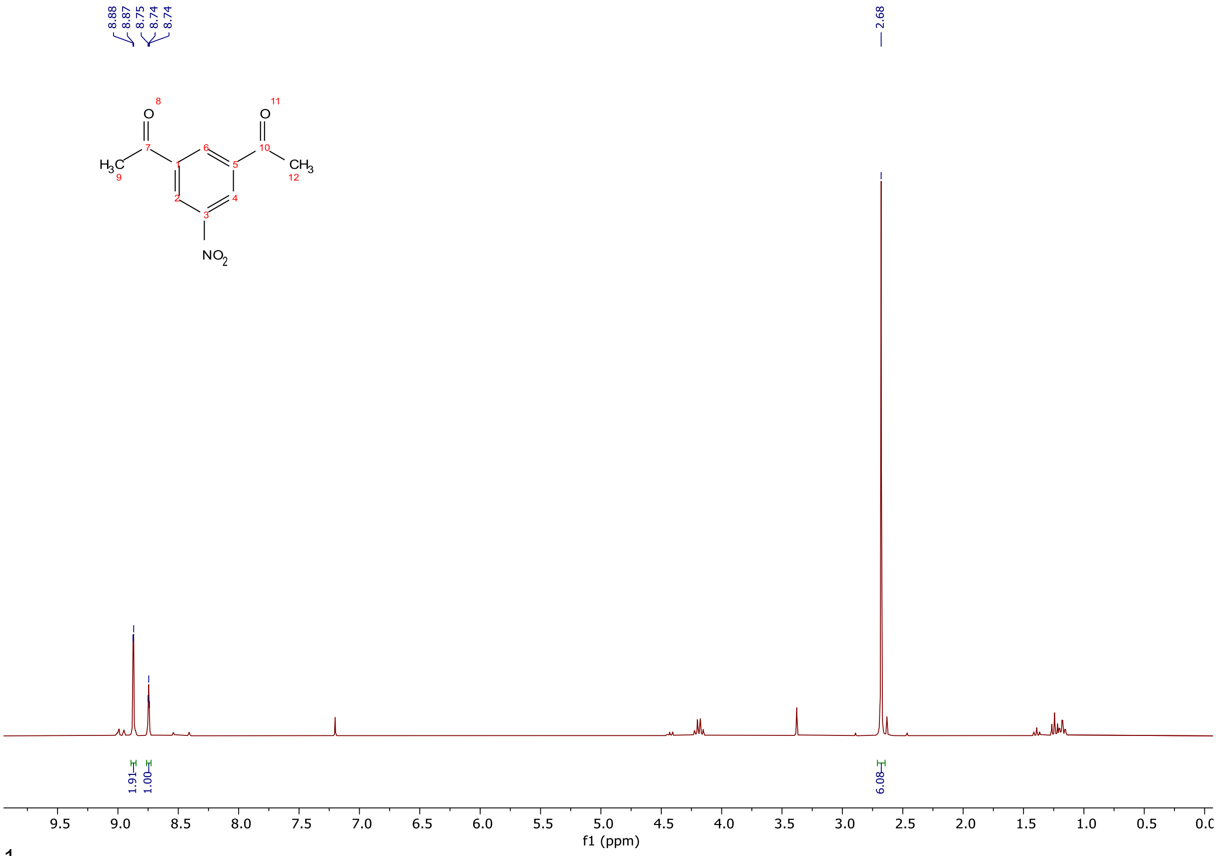


$^1$H NMR spectra (300 MHz, $CDCl_3$) of 3,5-diacetylnitrobenzene.

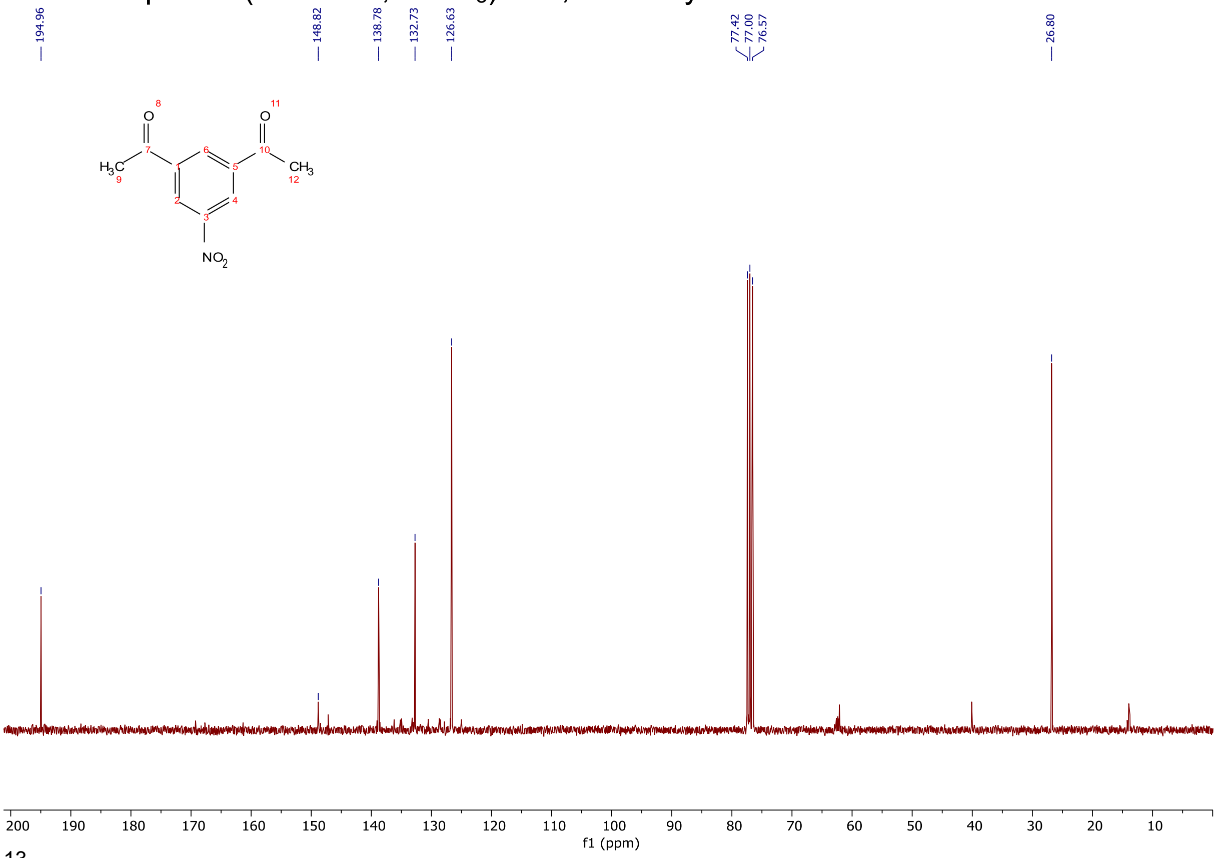


$^{13}$C NMR spectra (75 MHz, $CDCl_3$) of 3,5-diacetylnitrobenzene.

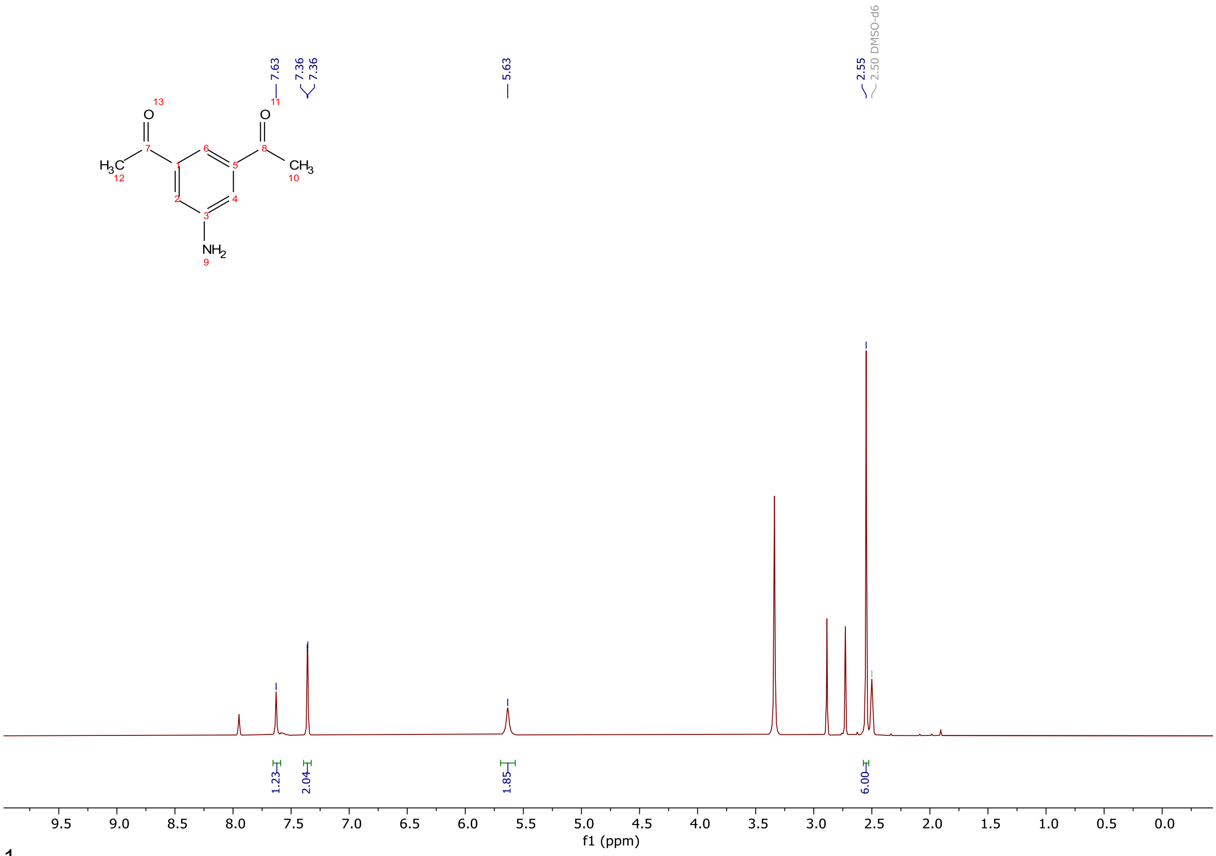


$^1$H NMR spectra (300 MHz, DMSO-$d_6$) of 3,5-diacetylaminobenzene.

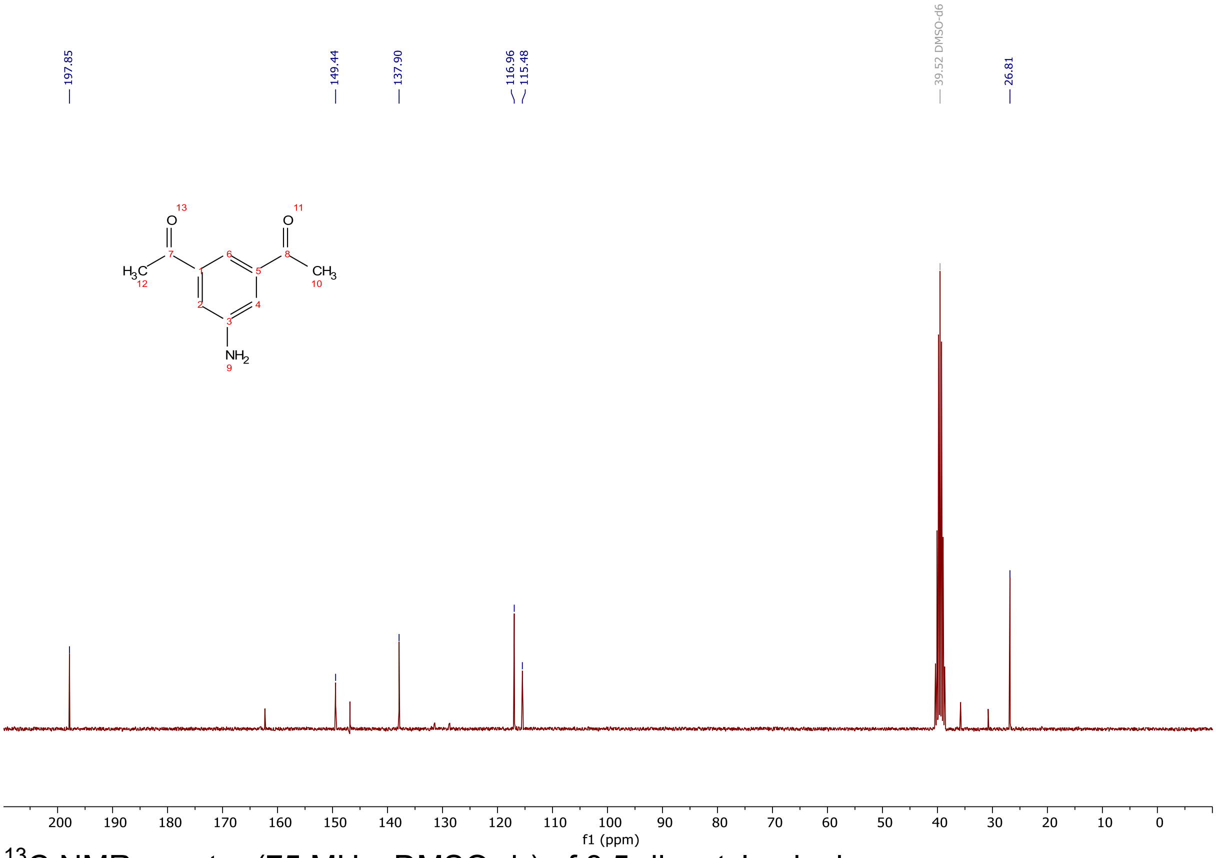


$^{13}$C NMR spectra (75 MHz, DMSO-$d_6$) of 3,5-diacetylaminobenzene.

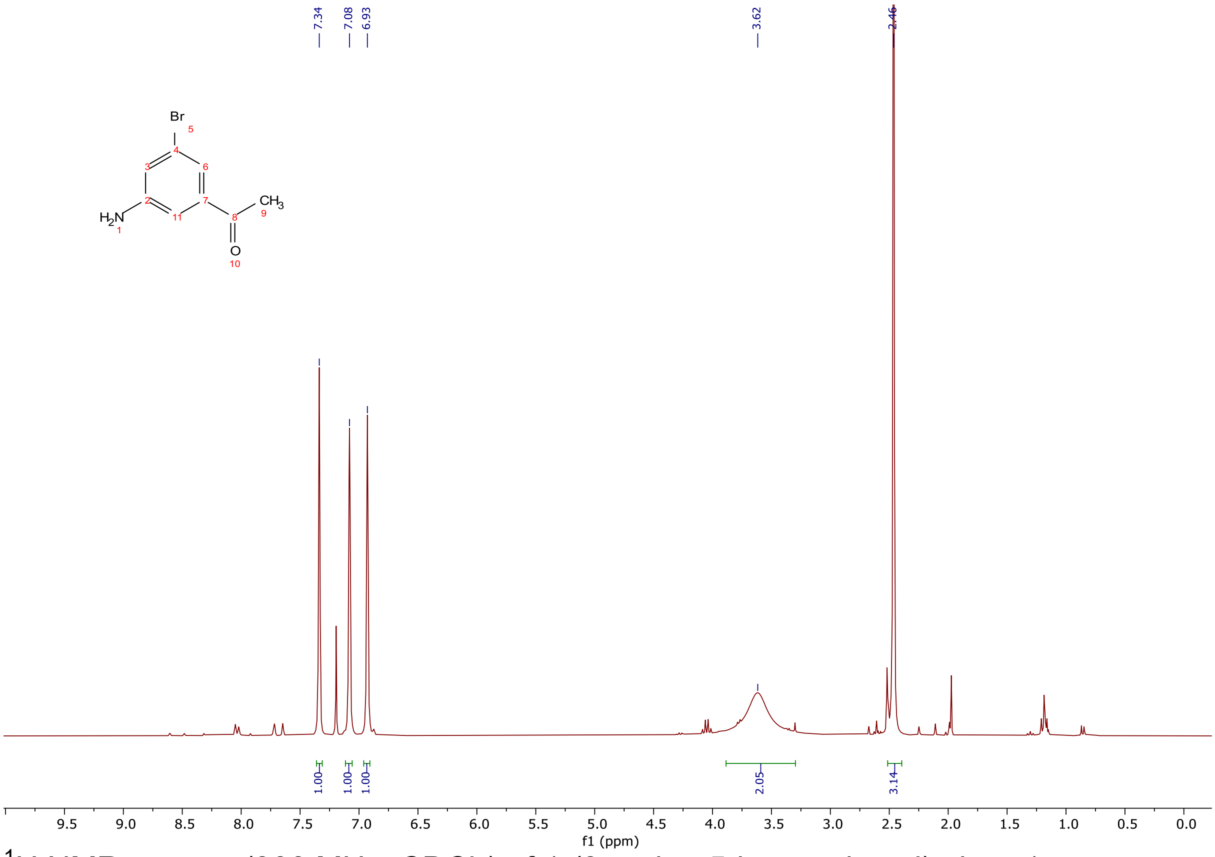


$^{1}H$ NMR spectra (300 MHz, $CDCl_3$) of 1-(3-amino-5-bromophenyl)ethan-1-one.

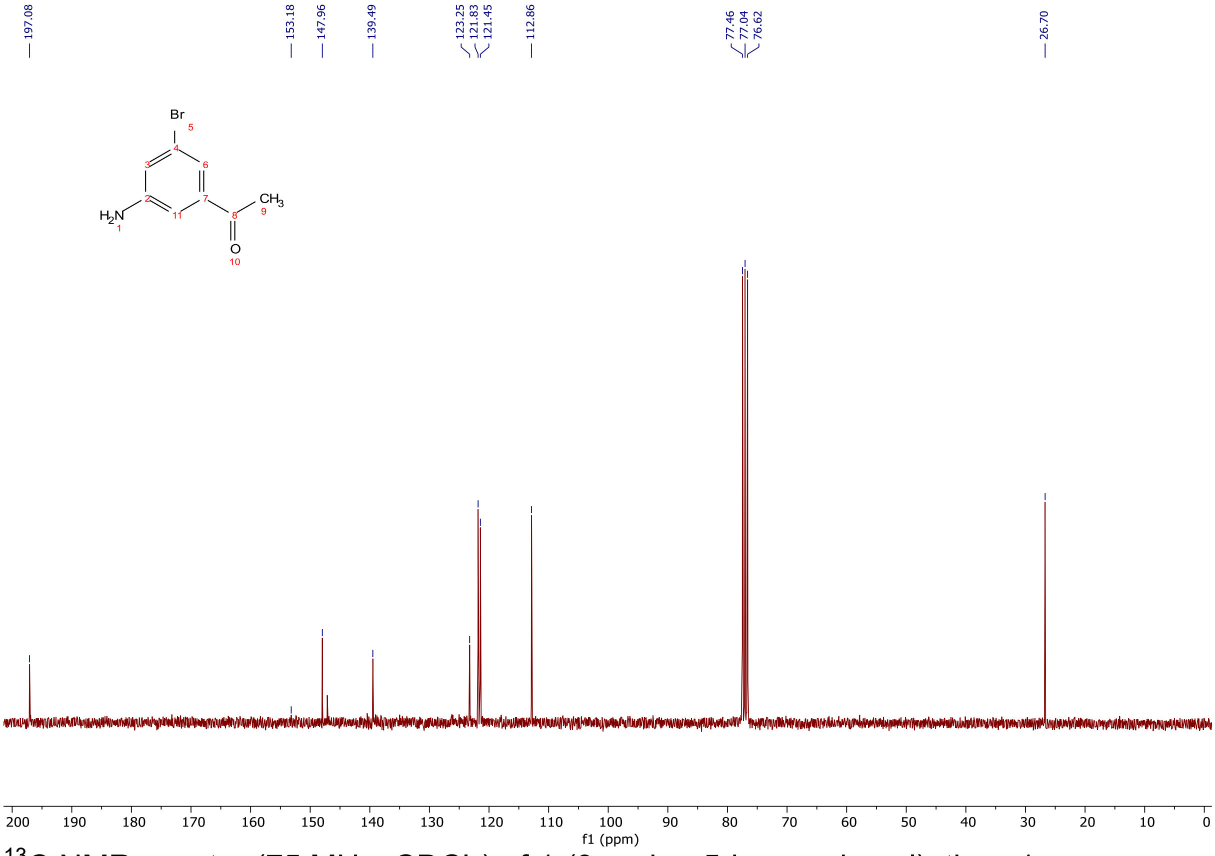


$^{13}C$ NMR spectra (75 MHz, $CDCl_3$) of 1-(3-amino-5-bromophenyl)ethan-1-one.

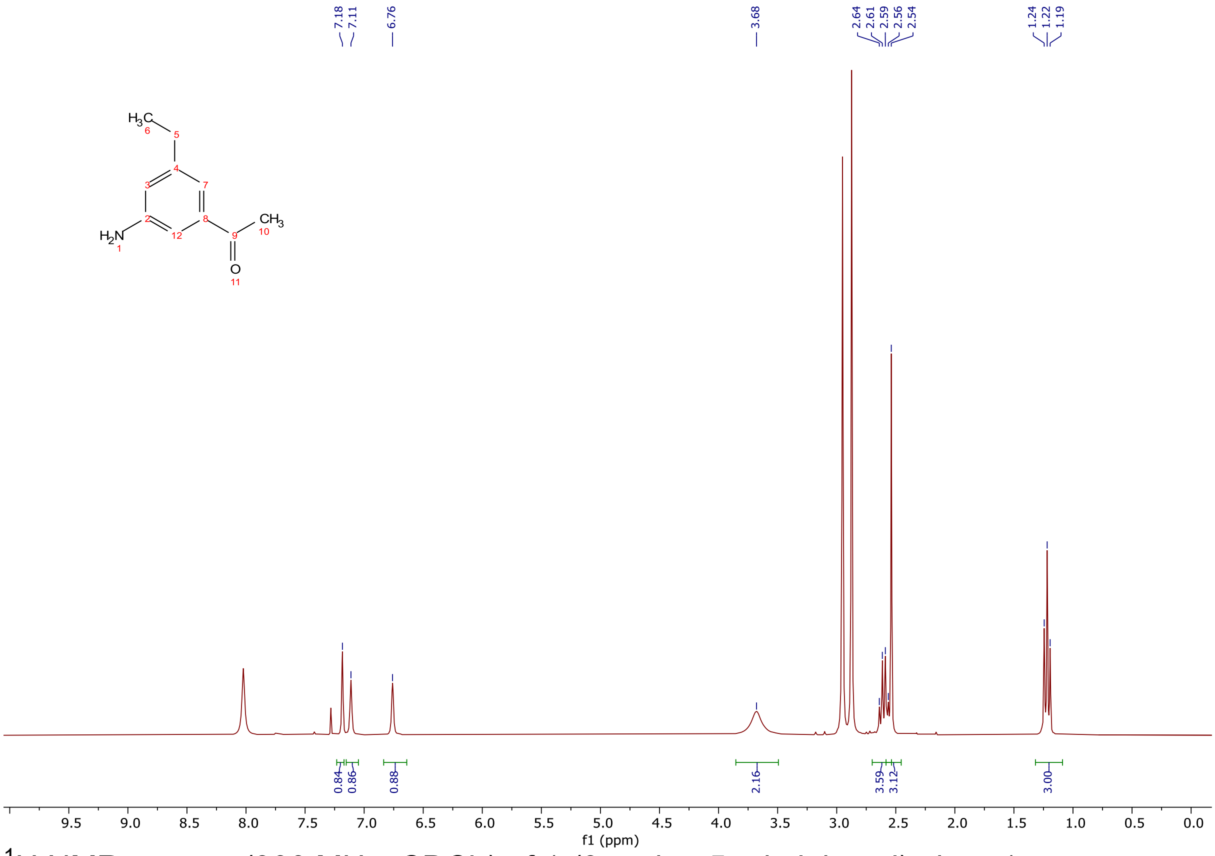


$^1$H NMR spectra (300 MHz, $CDCl_3$) of 1-(3-amino-5-ethylphenyl)ethan-1-one.

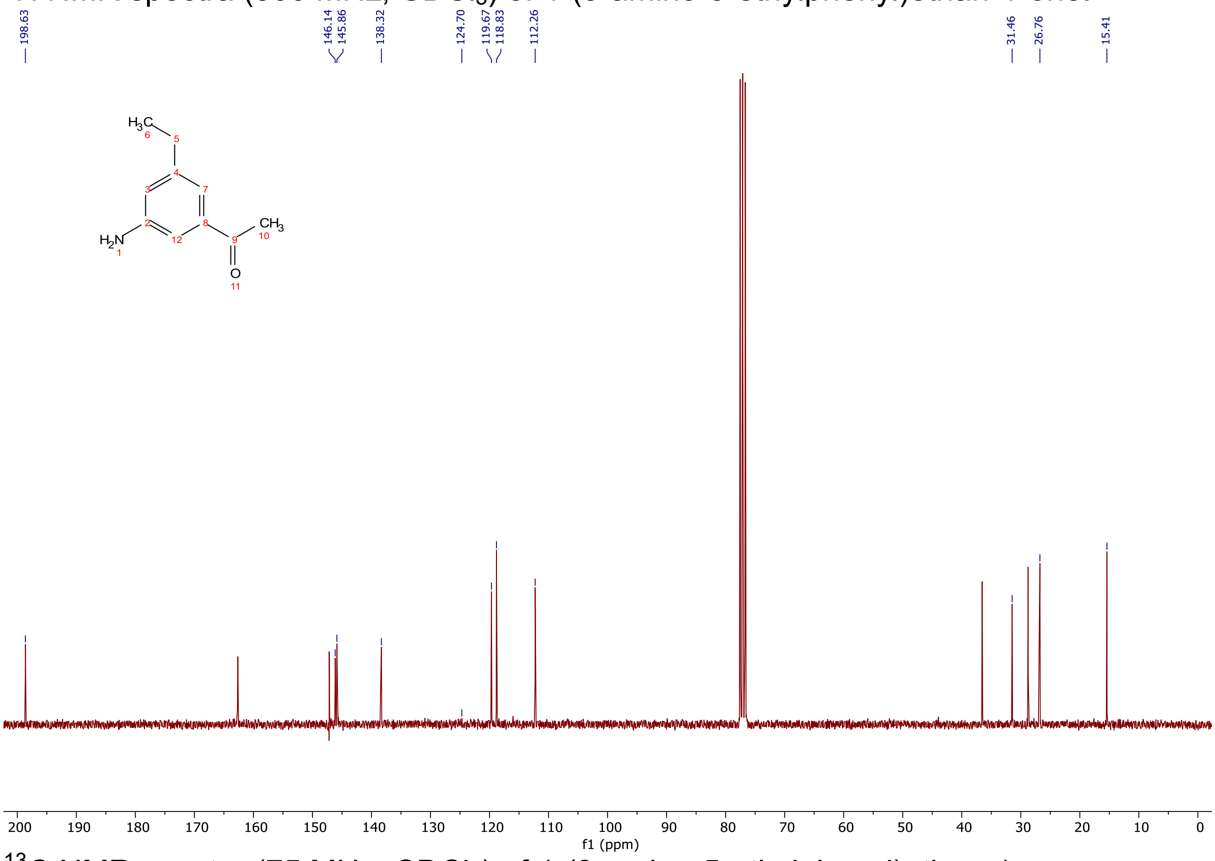


$^{13}$C NMR spectra (75 MHz, $CDCl_3$) of 1-(3-amino-5-ethylphenyl)ethan-1-one.

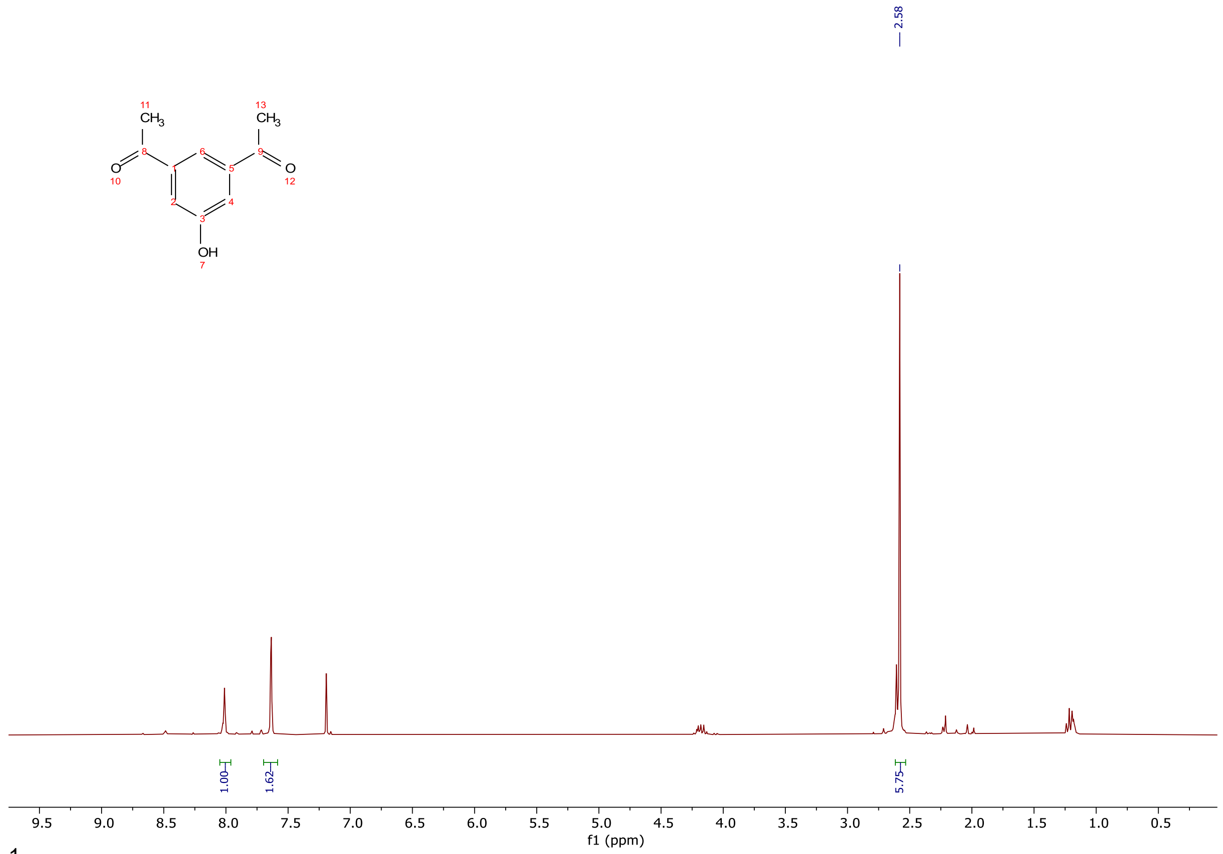


$^{1}$H NMR spectra (300 MHz, $CDCl_3$) of 3,5-diacetylphenol.

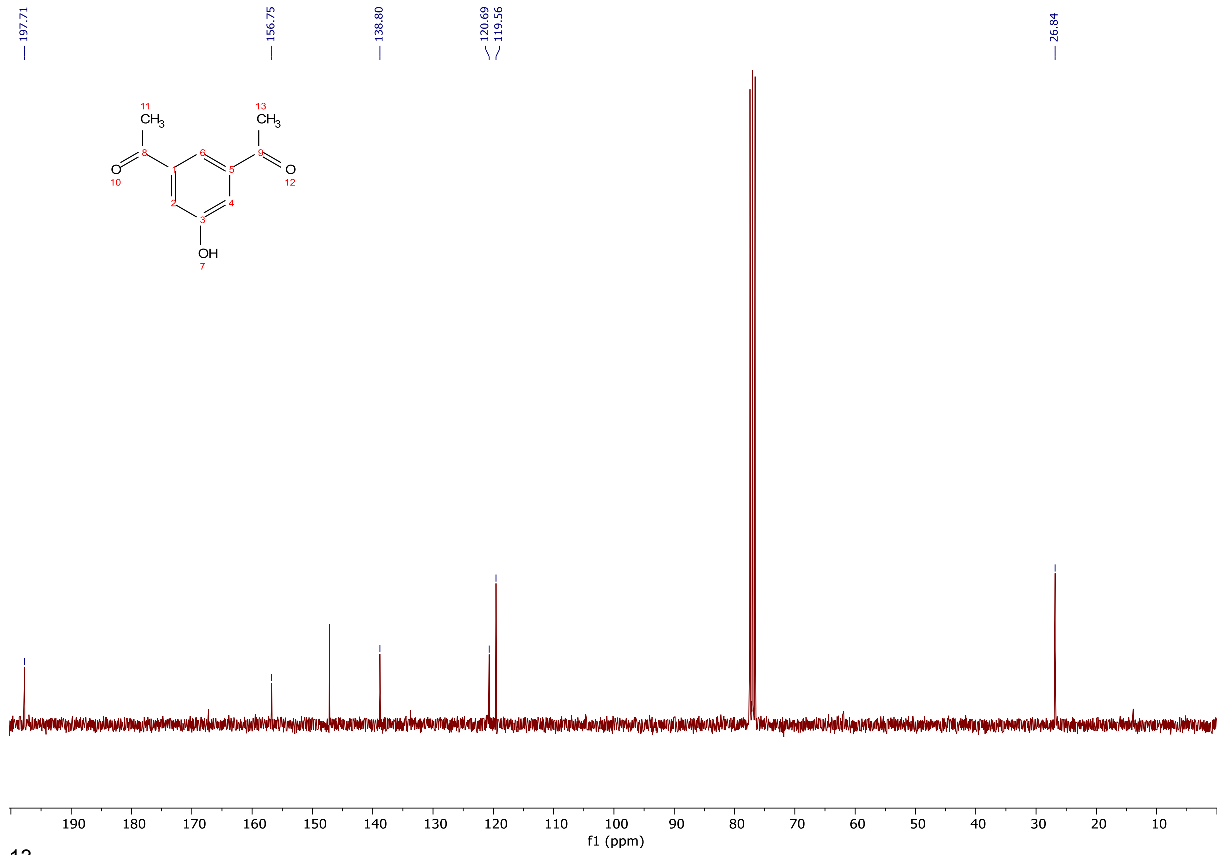


$^{13}$C NMR spectra (75 MHz, $CDCl_3$) of 3,5-diacetylphenol.

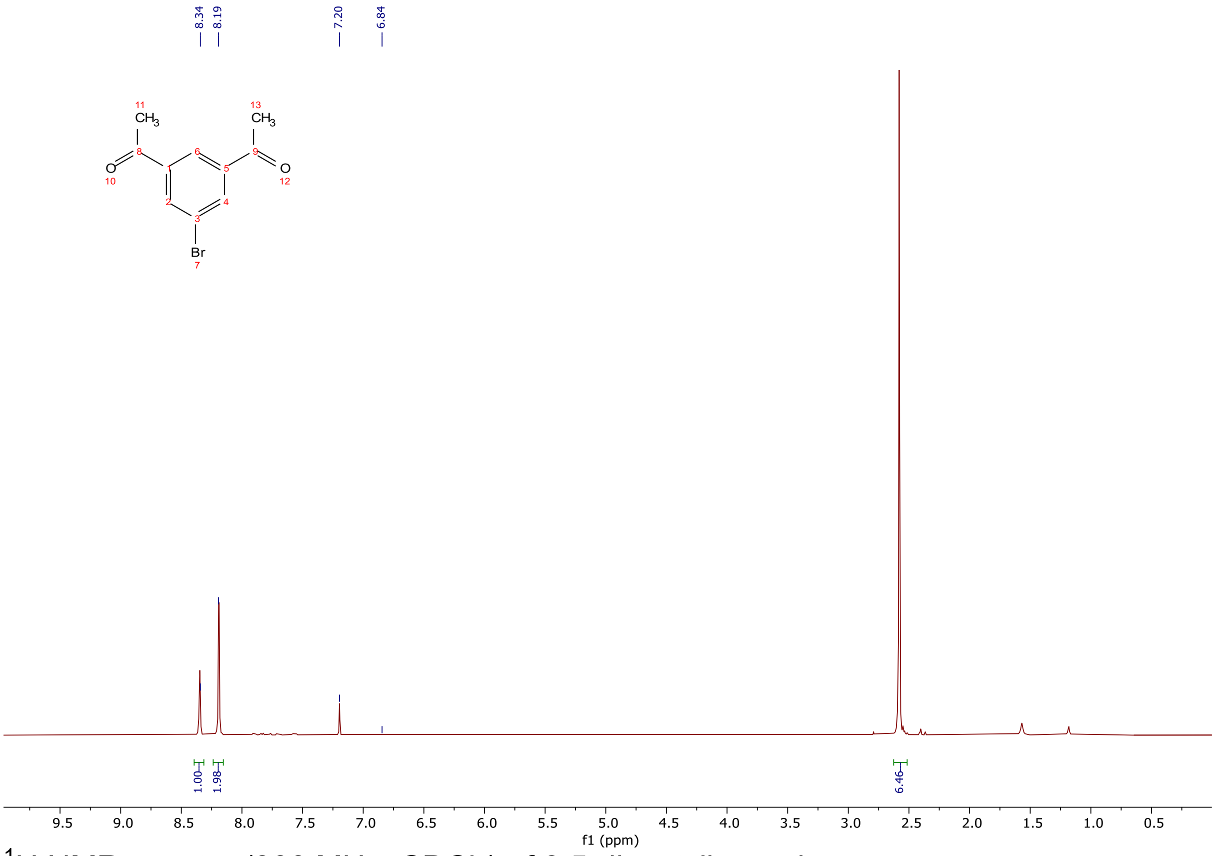


$^{1}$H NMR spectra (300 MHz, $CDCl_3$) of 3,5-diacetylbromobenzene.

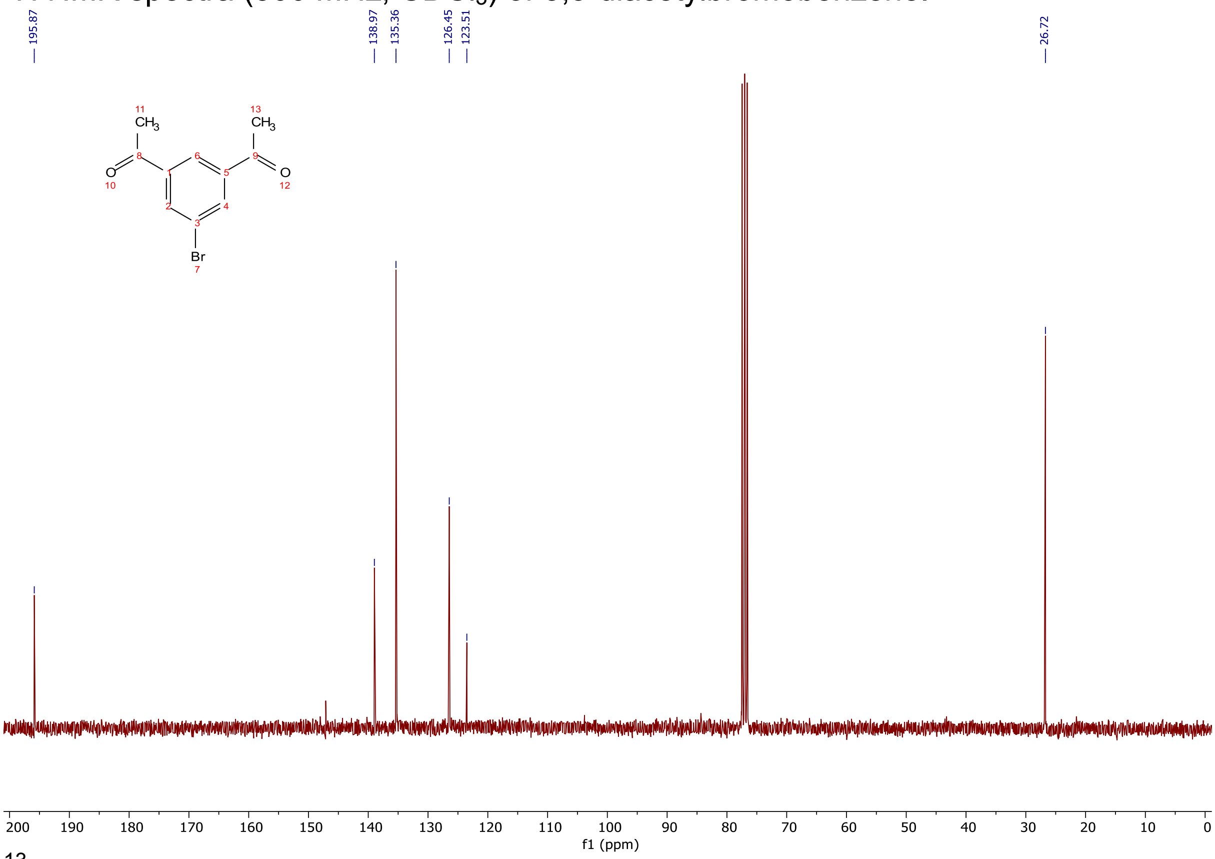


$^{13}$C NMR spectra (75 MHz, $CDCl_3$) of 3,5-diacetylbromobenzene.

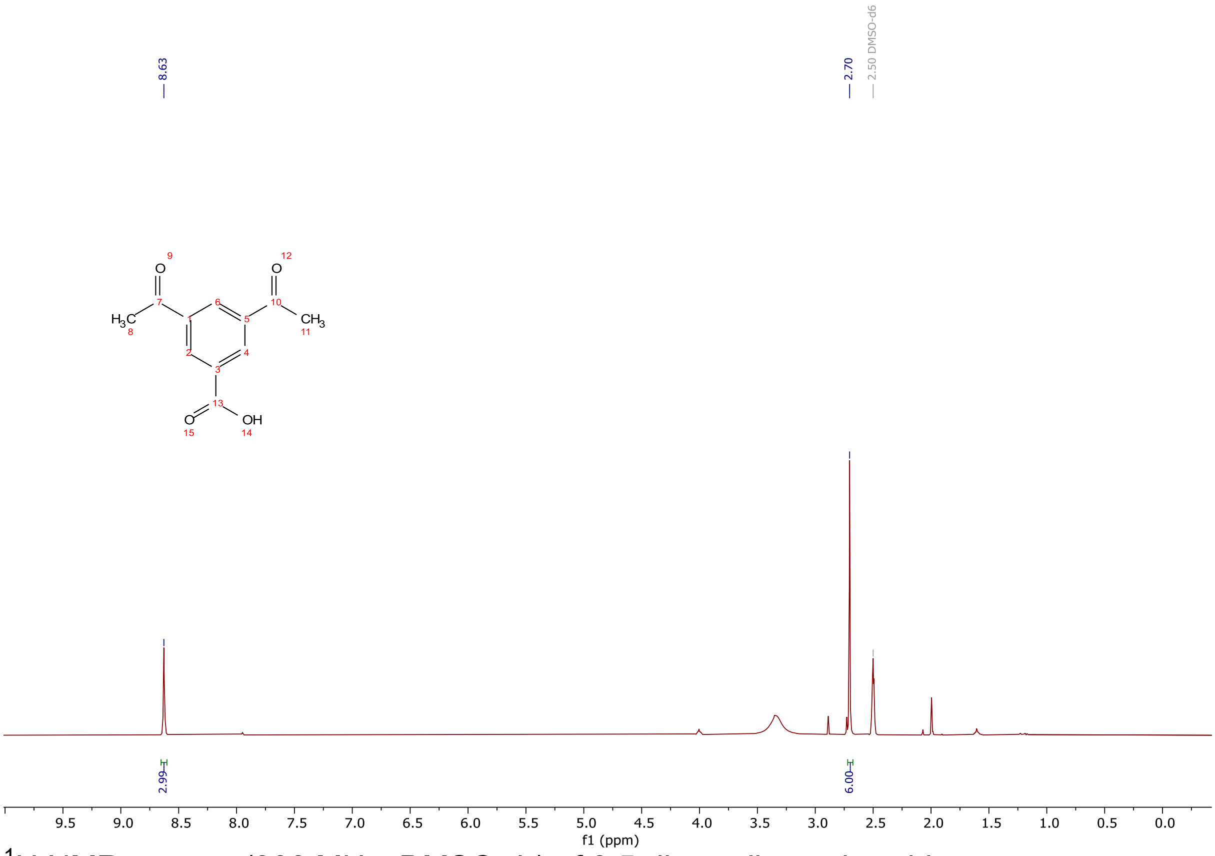

$^{1}H$ NMR spectra (300 MHz, DMSO-$d_6$) of 3,5-diacetylbenzoic acid.

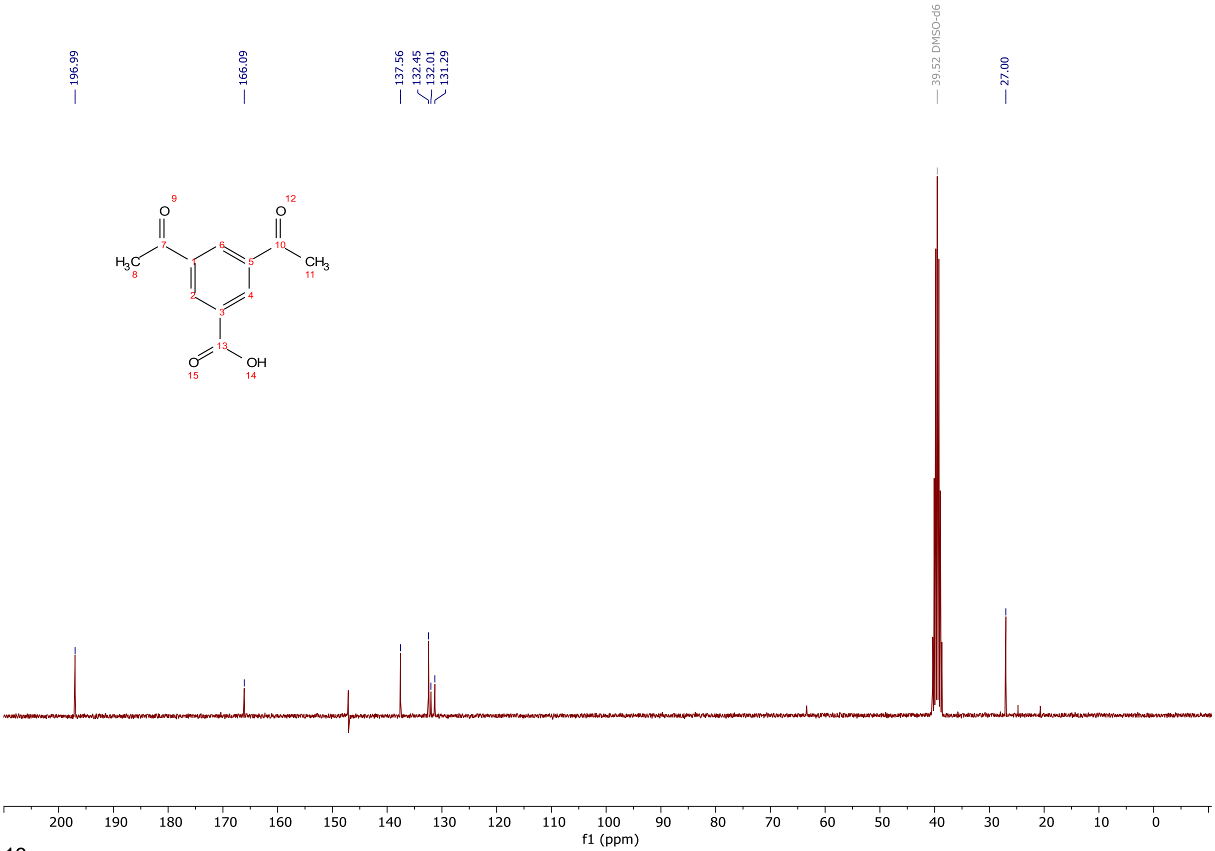

$^{13}C$ NMR spectra (300 MHz, DMSO-$d_6$) of 3,5-diacetylbenzoic acid.

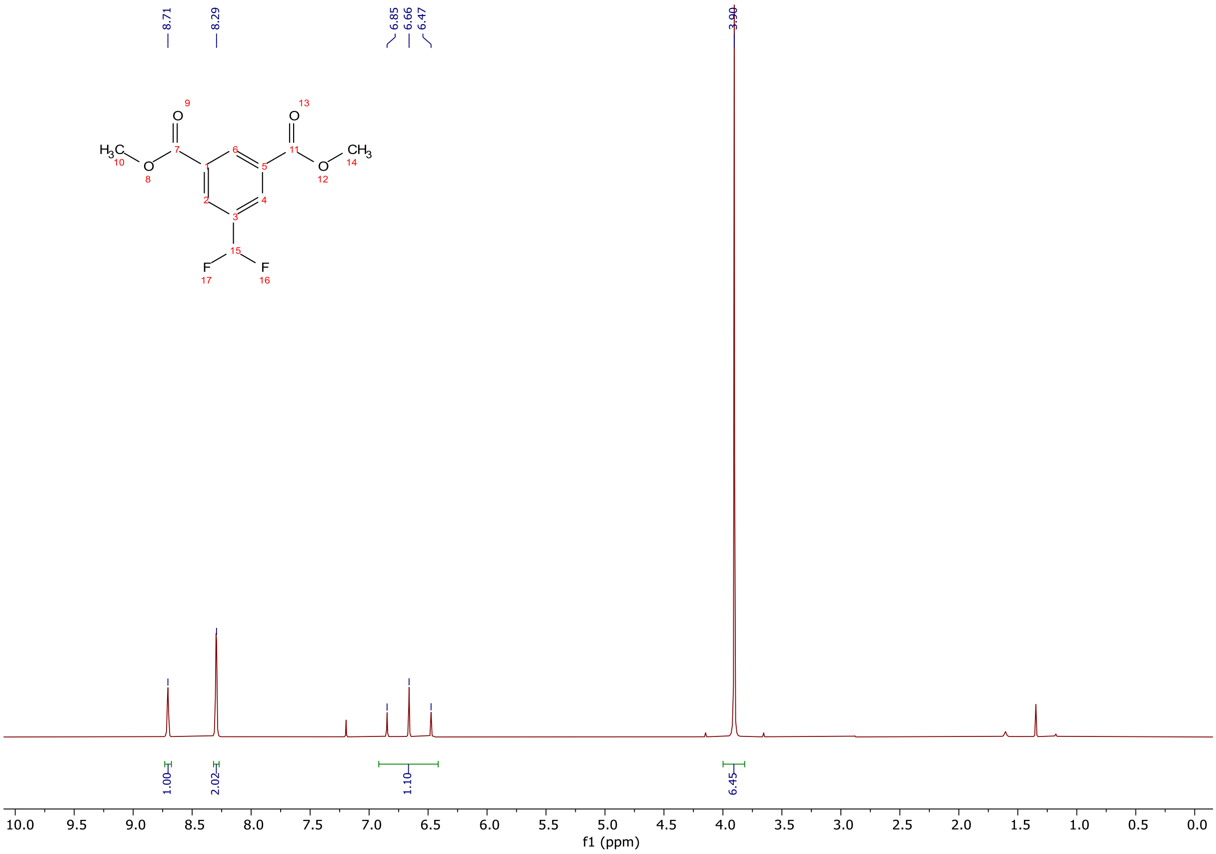


[1]H NMR spectra (300 MHz, $CDCl_3$) of 5-(difluoromethyl)isophthalate.

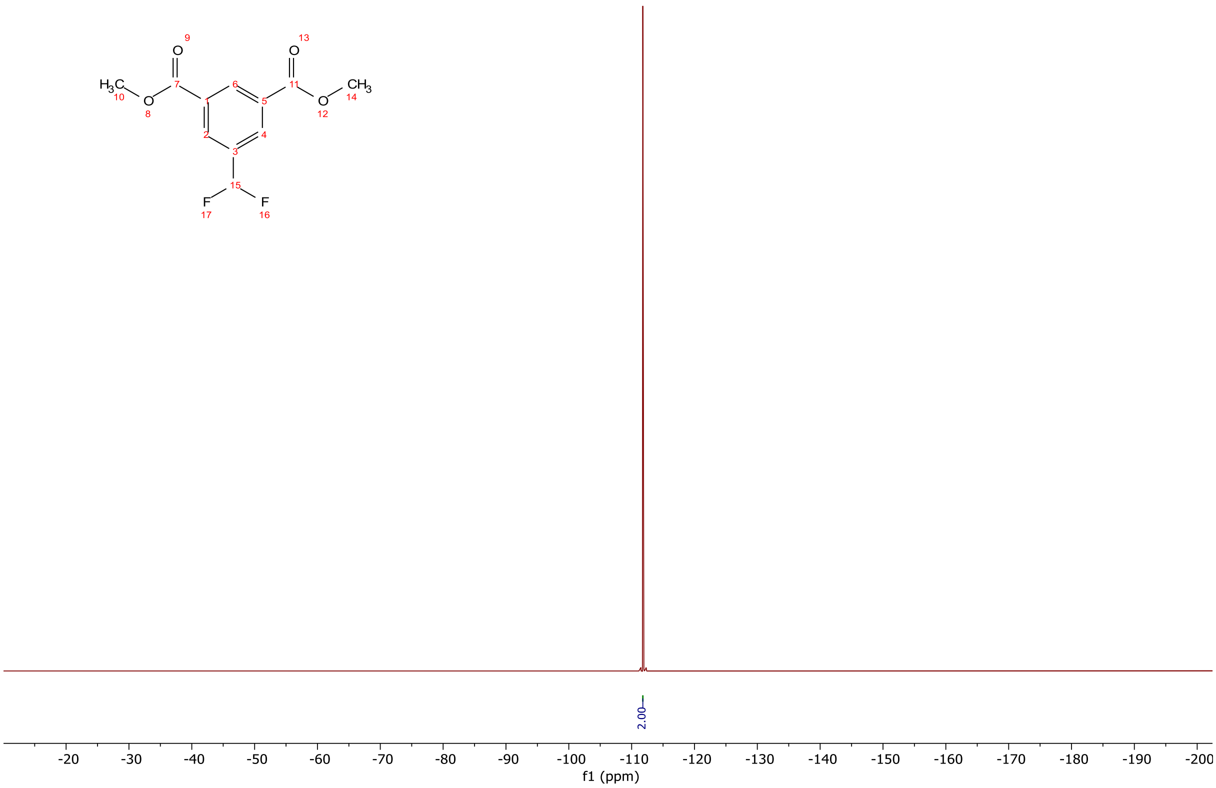


[19]F NMR spectra (283 MHz, $CDCl_3$) of 5-(difluoromethyl)isophthalate.

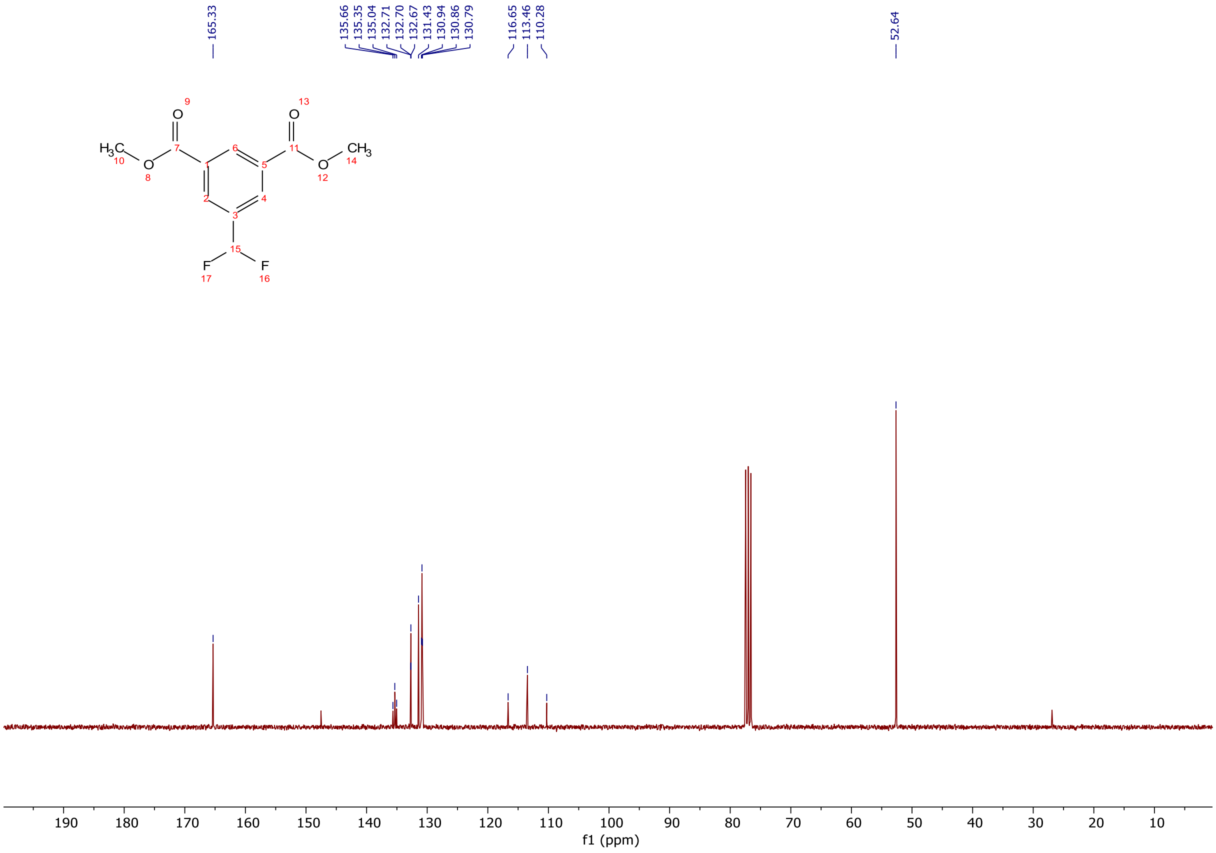


$^{13}C$ NMR spectra (75 MHz, $CDCl_3$) of 5-(difluoromethyl)isophthalate.

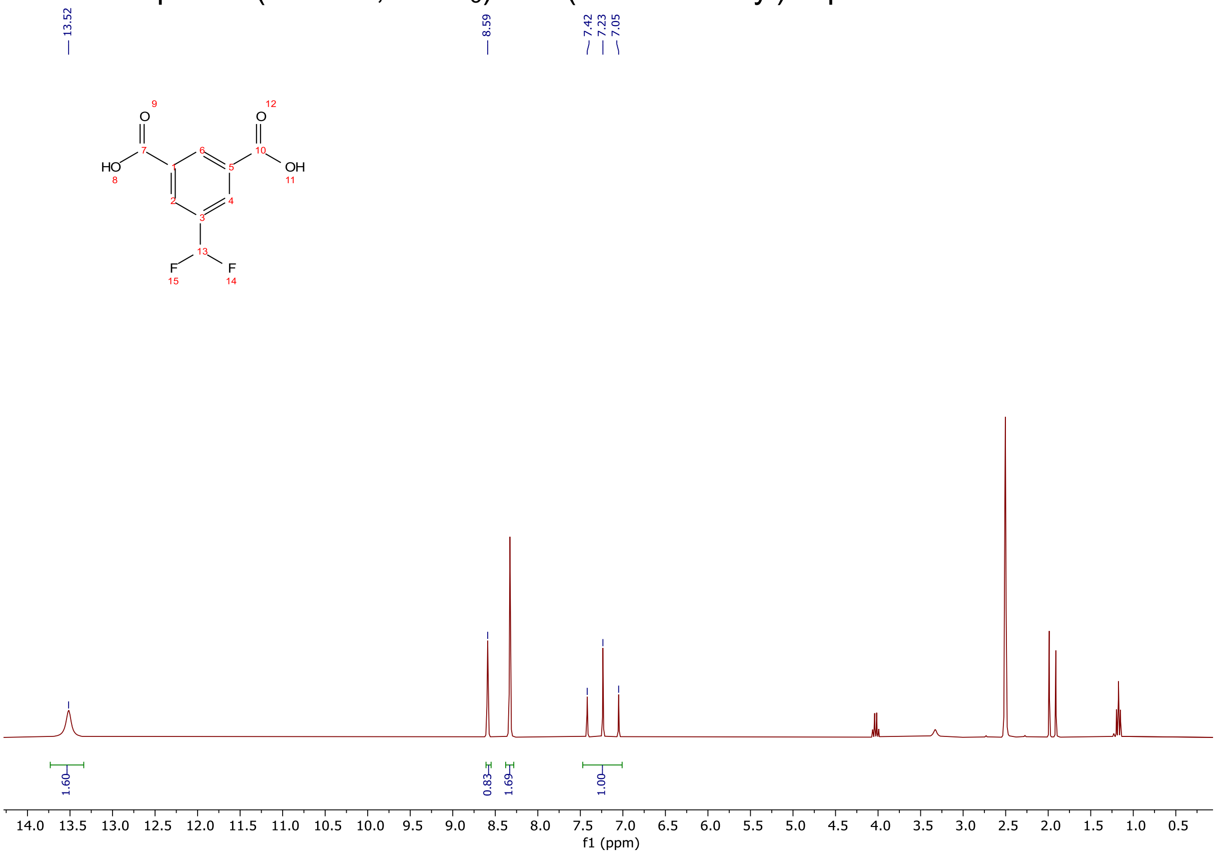


$^{1}H$ NMR spectra (300 MHz, DMSO-$d_6$) of 5-(difluoromethyl)isophthalic acid.

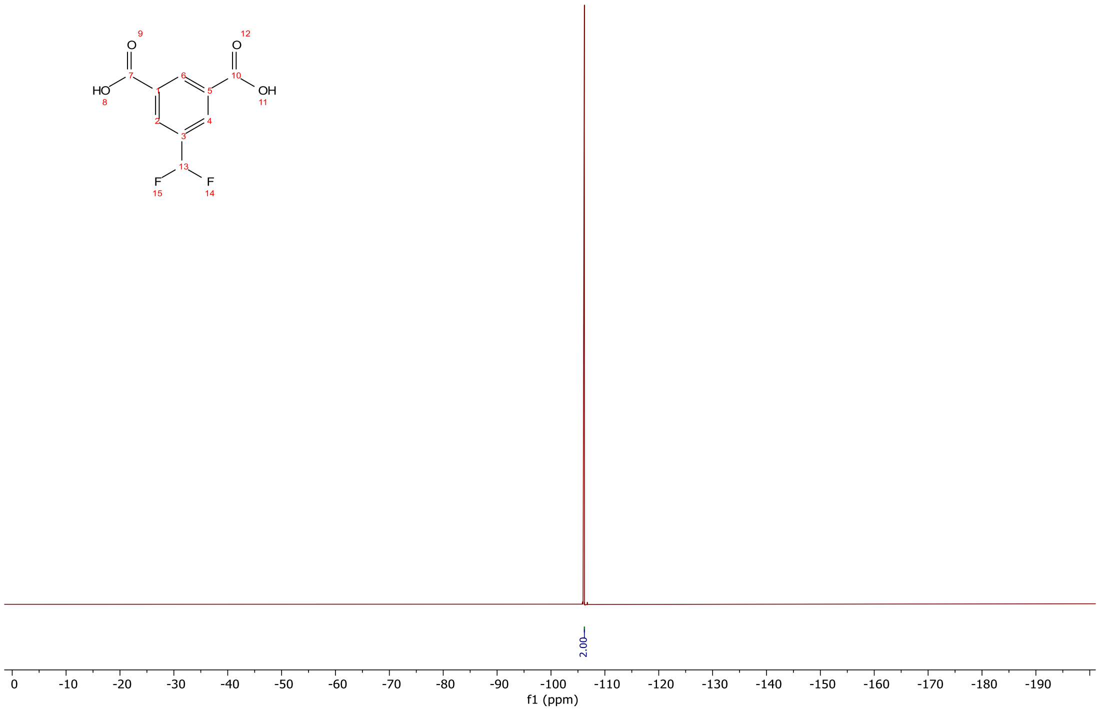


$^{19}$F NMR spectra (283 MHz, DMSO-d$_6$) of 5-(difluoromethyl)isophthalic acid.

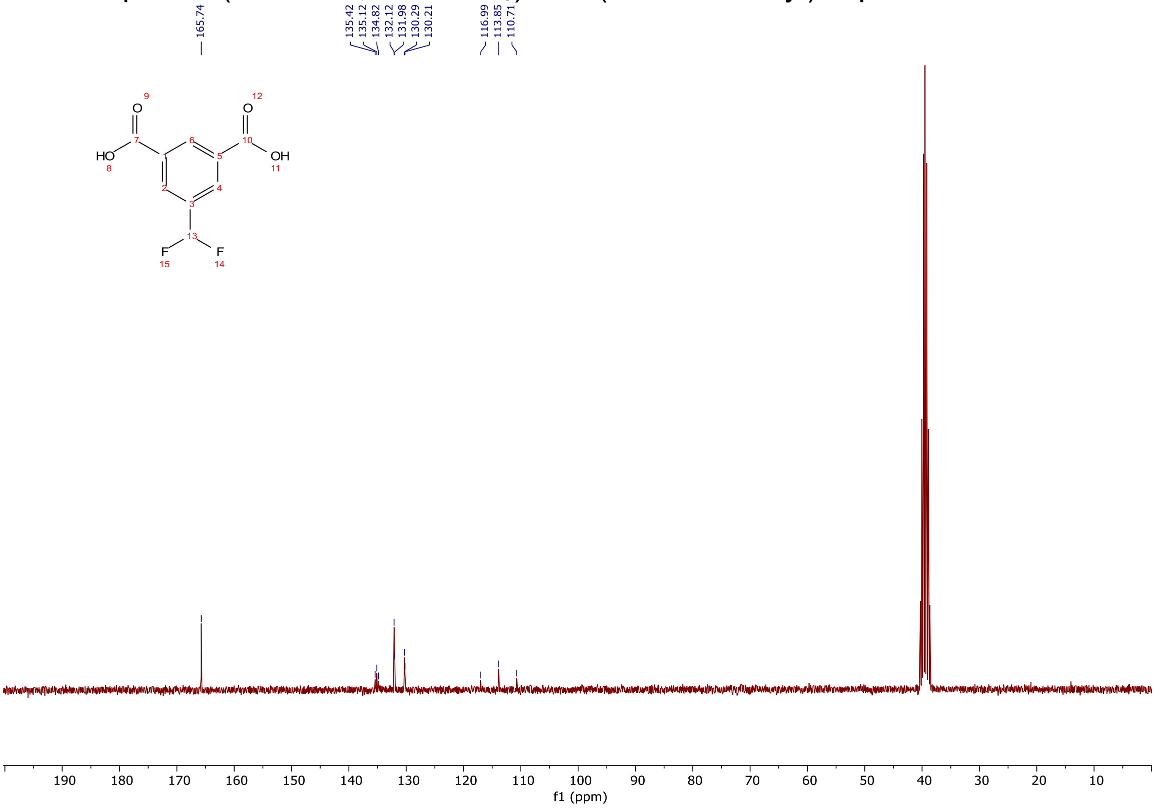


$^{13}$C NMR spectra (75 MHz, DMSO-d$_6$) of 5-(difluoromethyl)isophthalic acid.

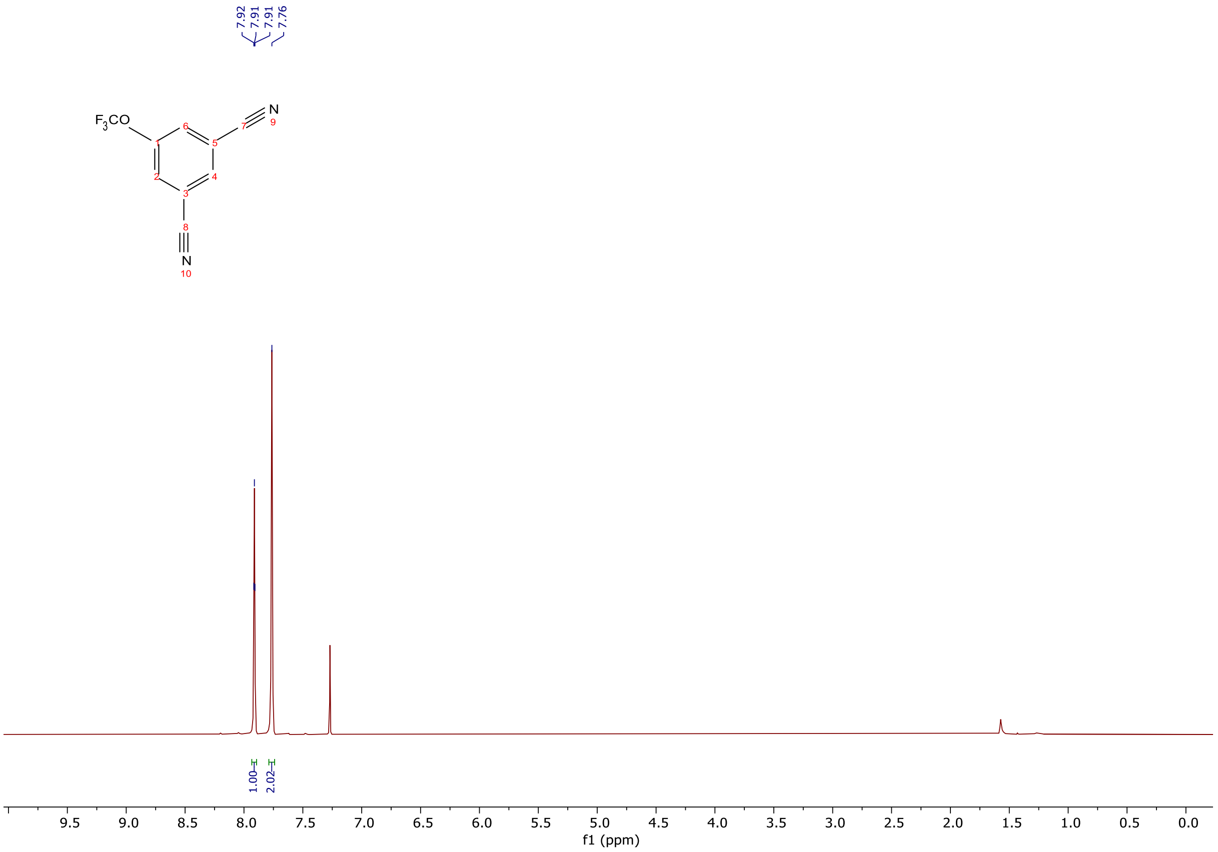


$^{1}H$ NMR spectra (300 MHz, $CDCl_3$) of 5-(trifluoromethoxy)isophthalonitrile.

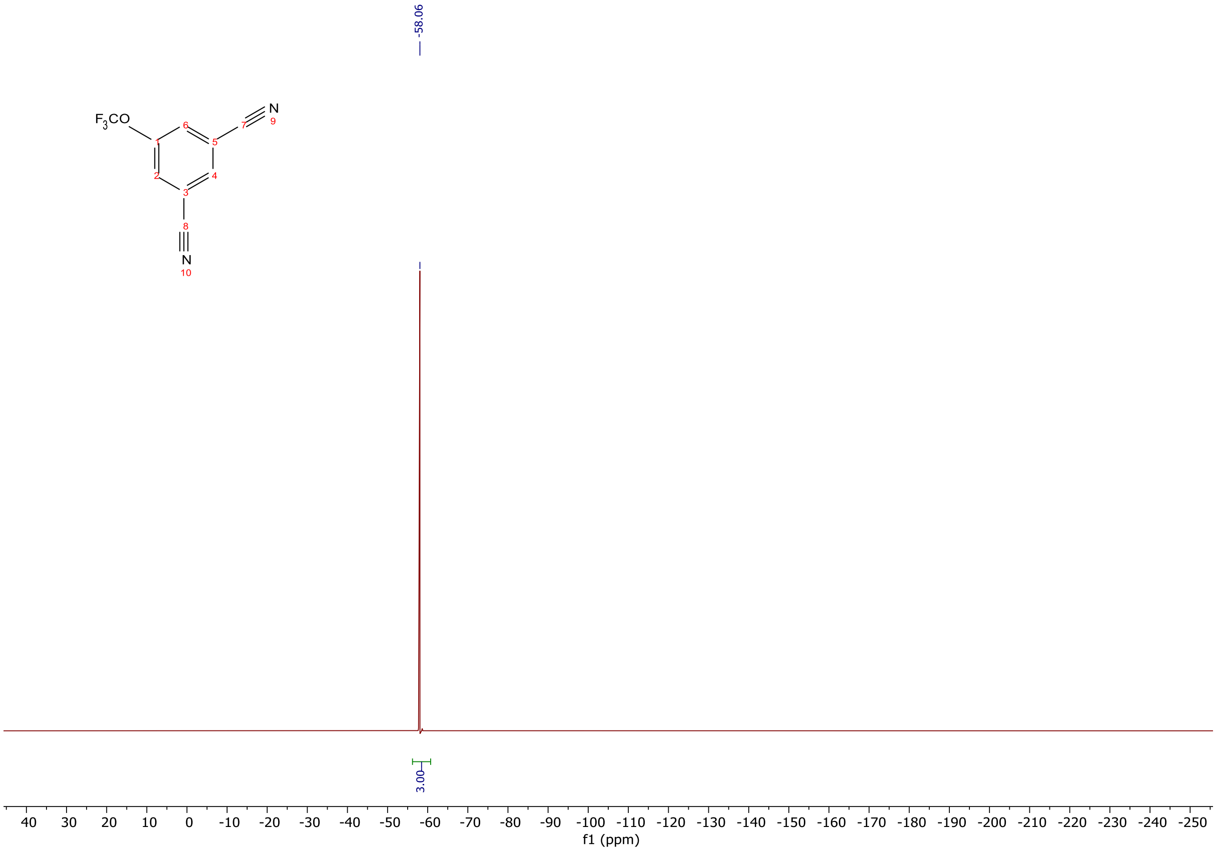


$^{19}F$ NMR spectra (283 MHz, $CDCl_3$) of 5-(trifluoromethoxy)isophthalonitrile.

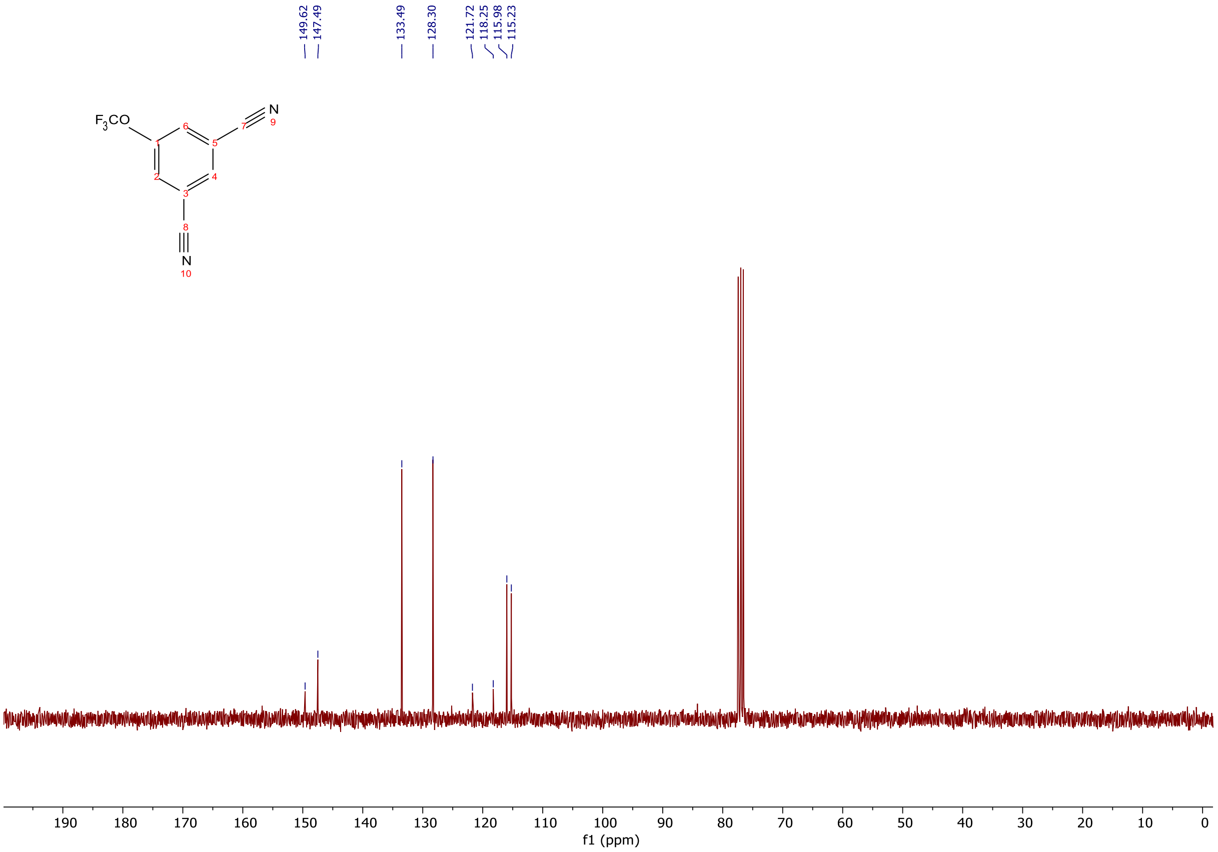


$^{13}C$ NMR spectra (75 MHz, $CDCl_3$) of 5-(trifluoromethoxy)isophthalonitrile.

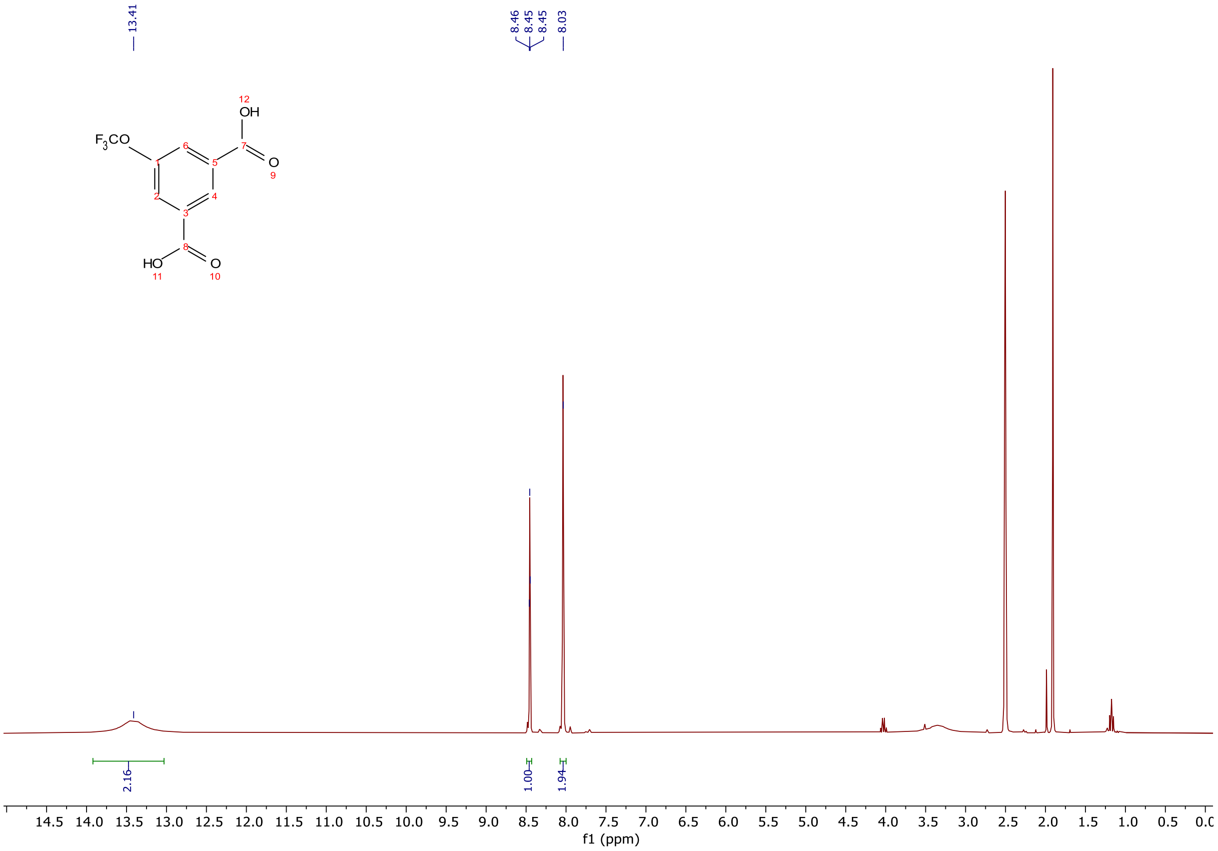


$^{1}H$ NMR spectra (300 MHz, DMSO-$d_6$) of 5-(trifluoromethoxy)isophthalic acid.

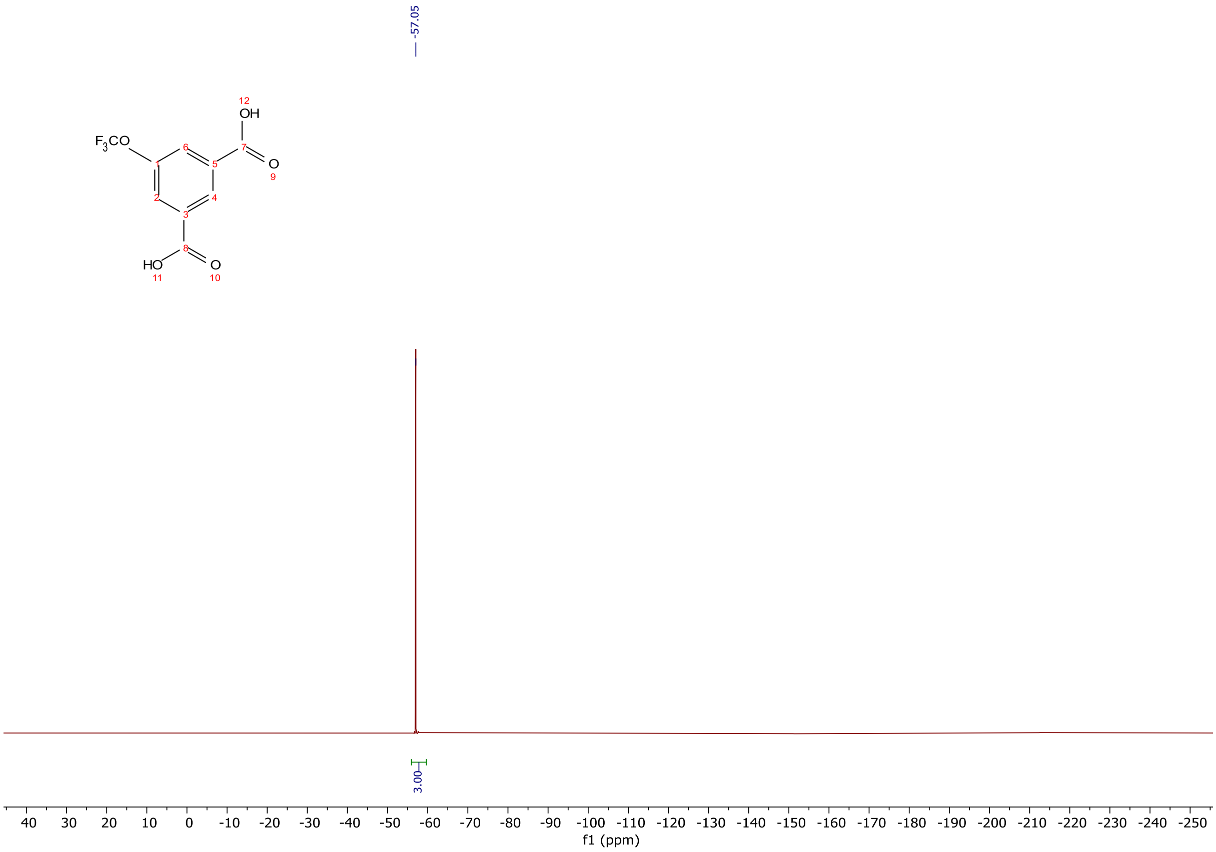


$^{19}F$ NMR spectra (283 MHz, DMSO-$d_6$) of 5-(trifluoromethoxy)isophthalic acid.

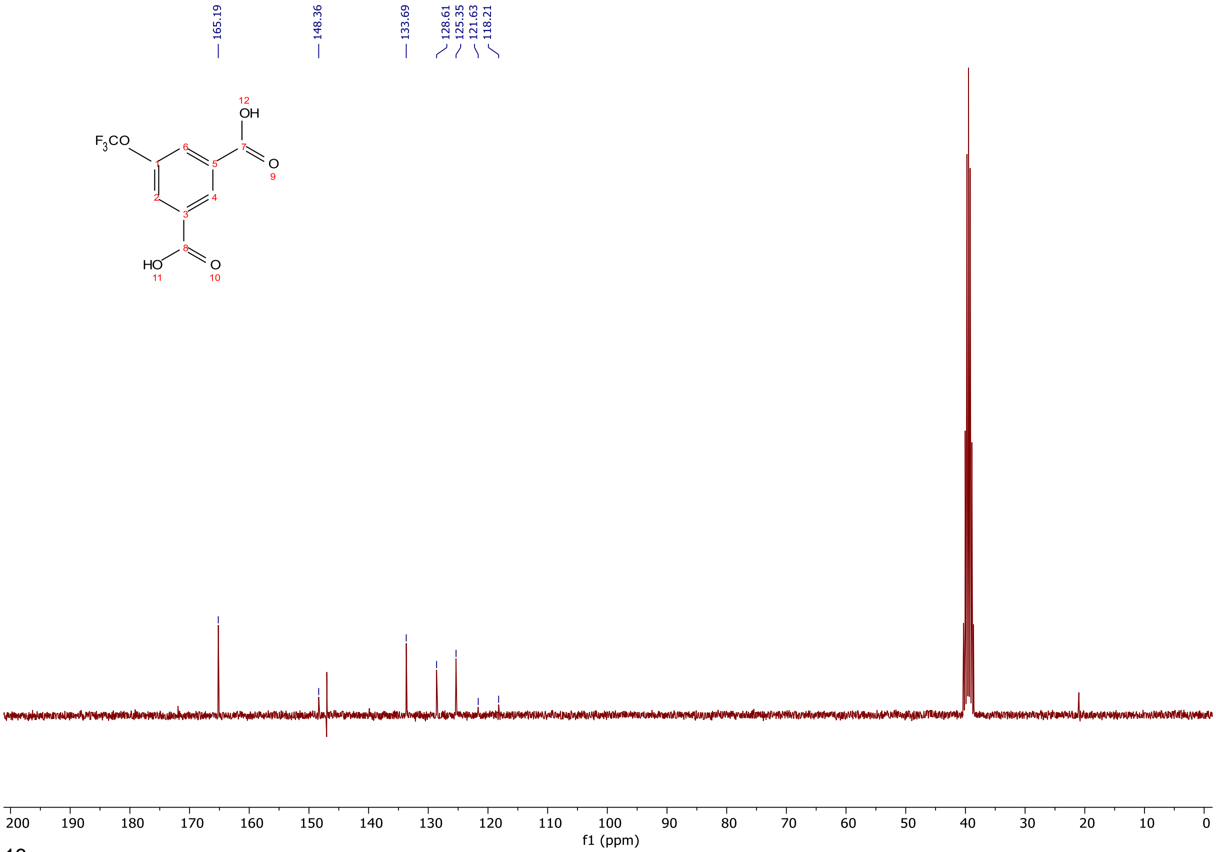


$^{13}C$ NMR spectra (75 MHz, DMSO-$d_6$) of 5-(trifluoromethoxy)isophthalic acid.

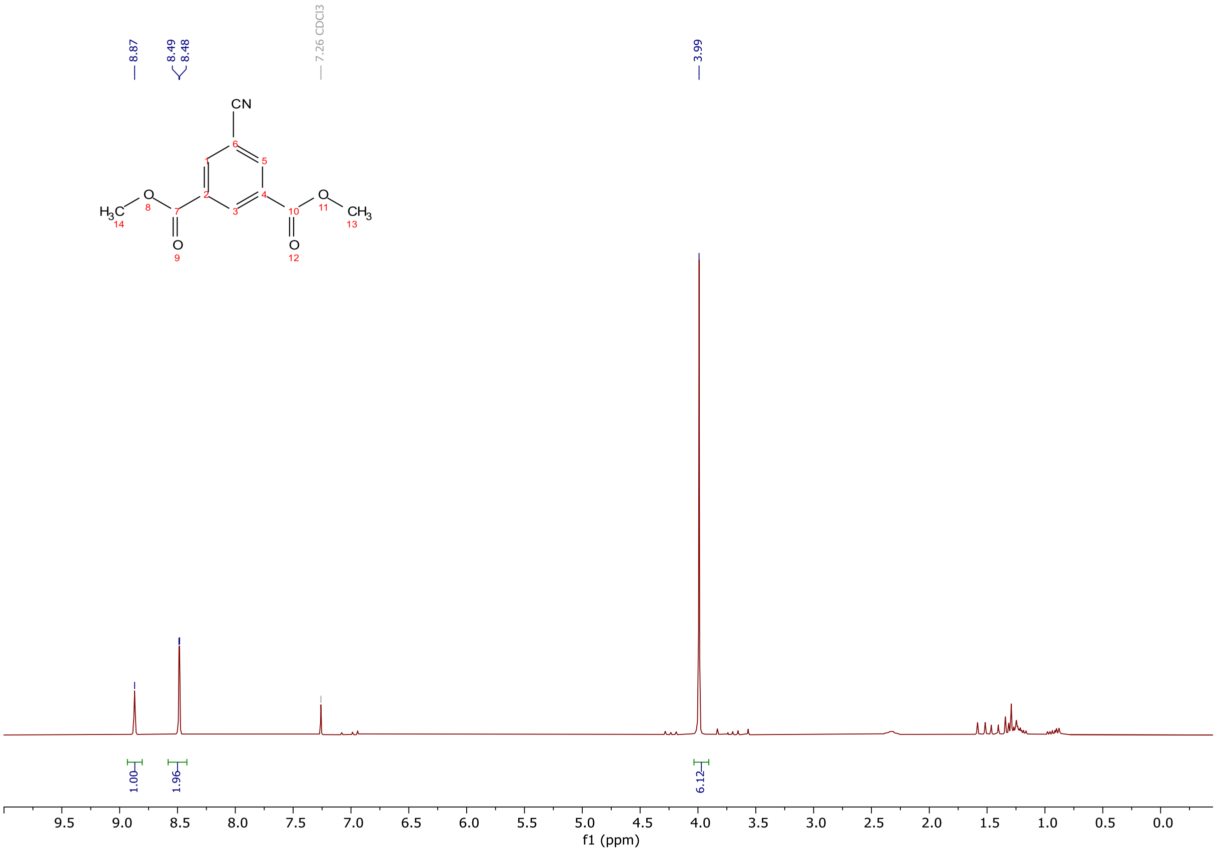


$^{1}H$ NMR spectra (300 MHz, $CDCl_3$) of dimethyl 5-cyanoisophthalate.

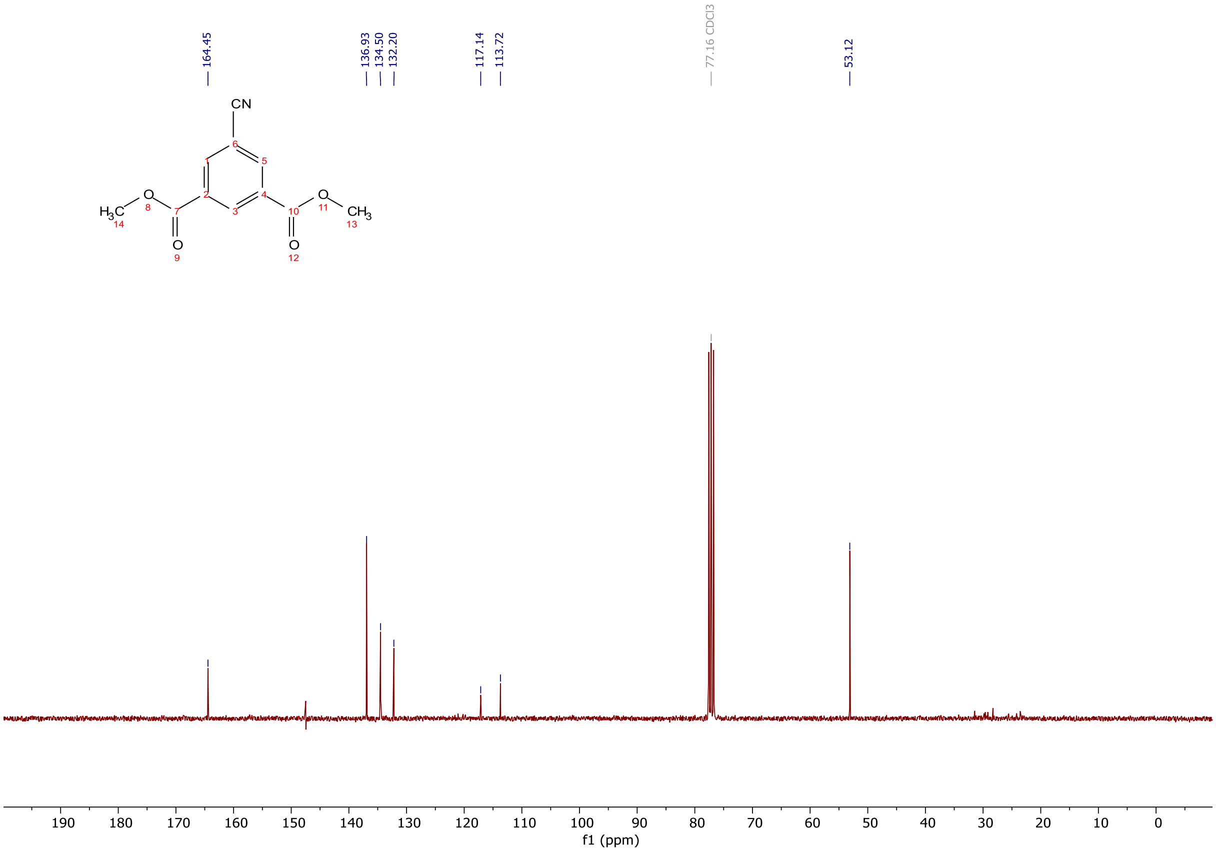


$^{13}C$ NMR spectra (75 MHz, $CDCl_3$) of dimethyl 5-cyanoisophthalate.

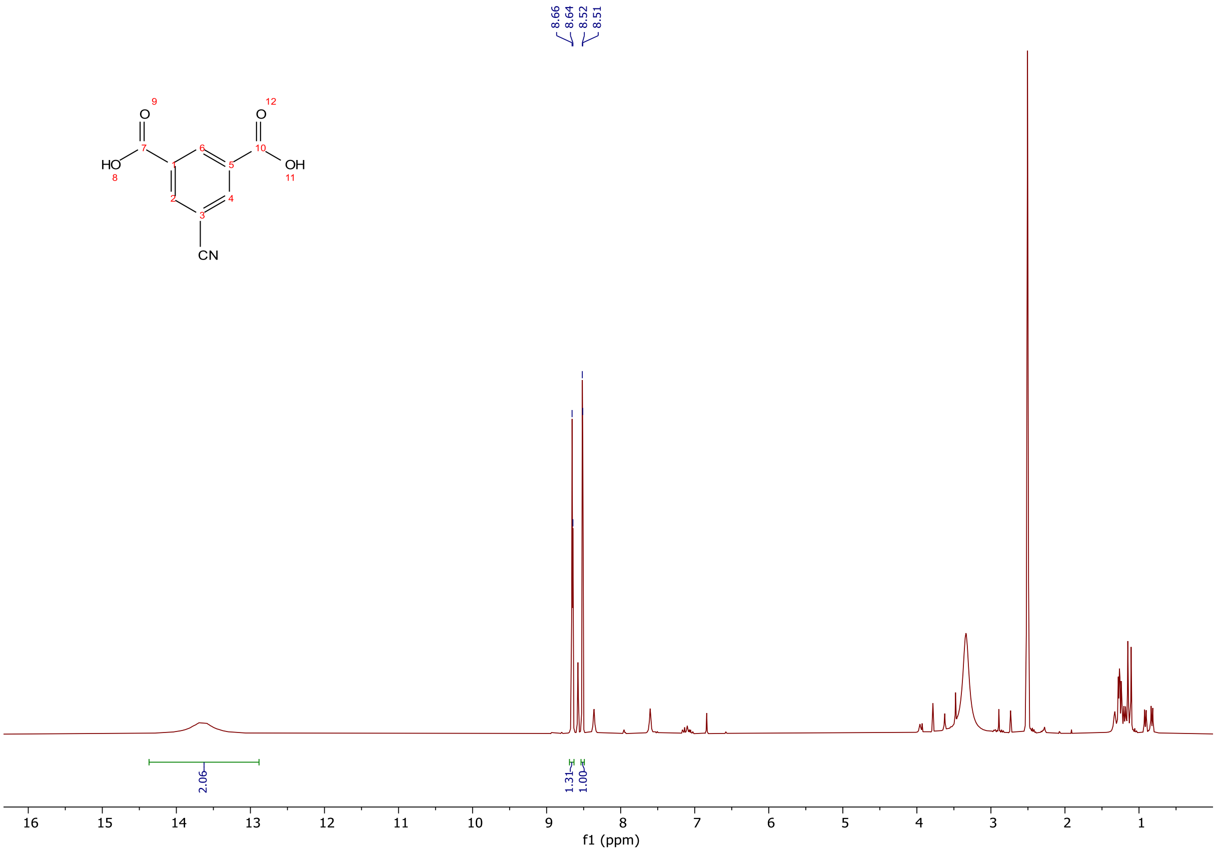


$^{1}$H NMR spectra (300 MHz, DMSO-$d_6$) of 5-cyanoisophthalic acid.

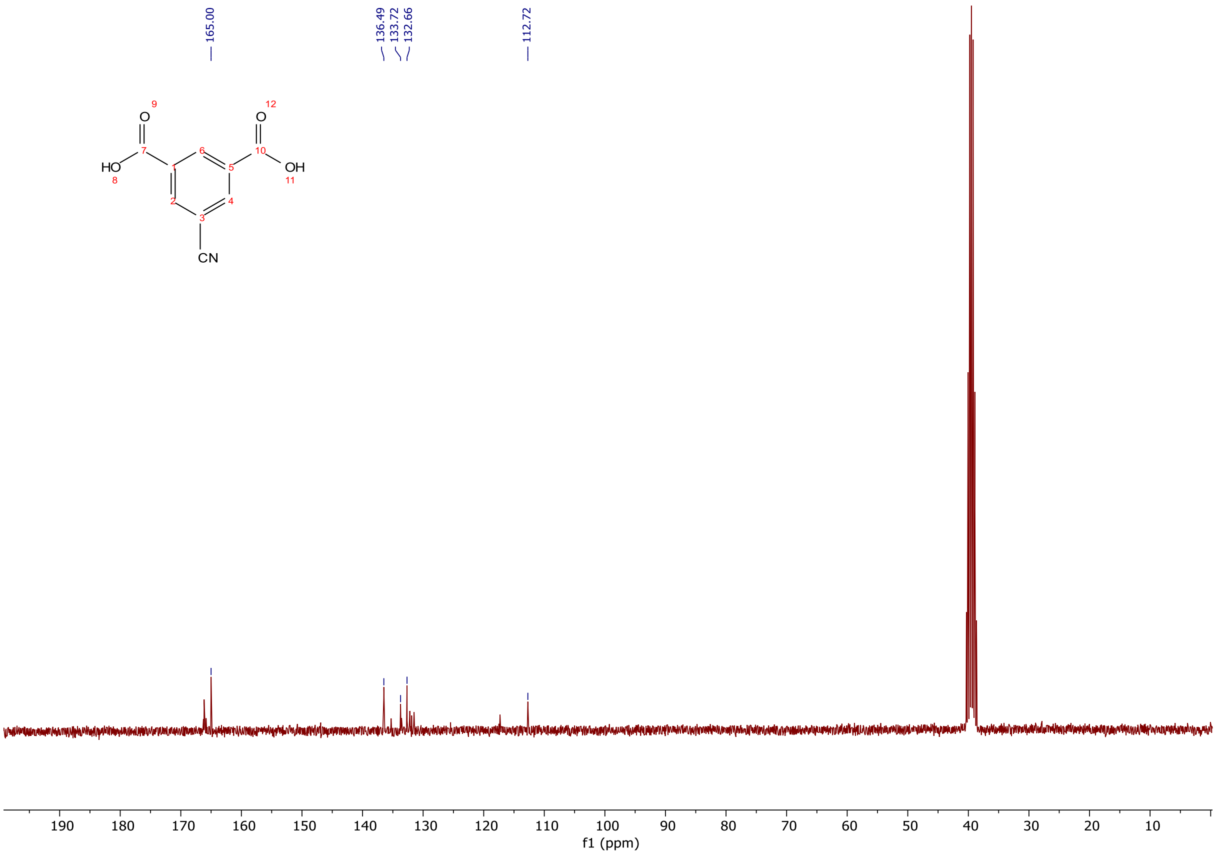


$^{13}$C NMR spectra (75 MHz, DMSO-$d_6$) of 5-cyanoisophthalic acid.

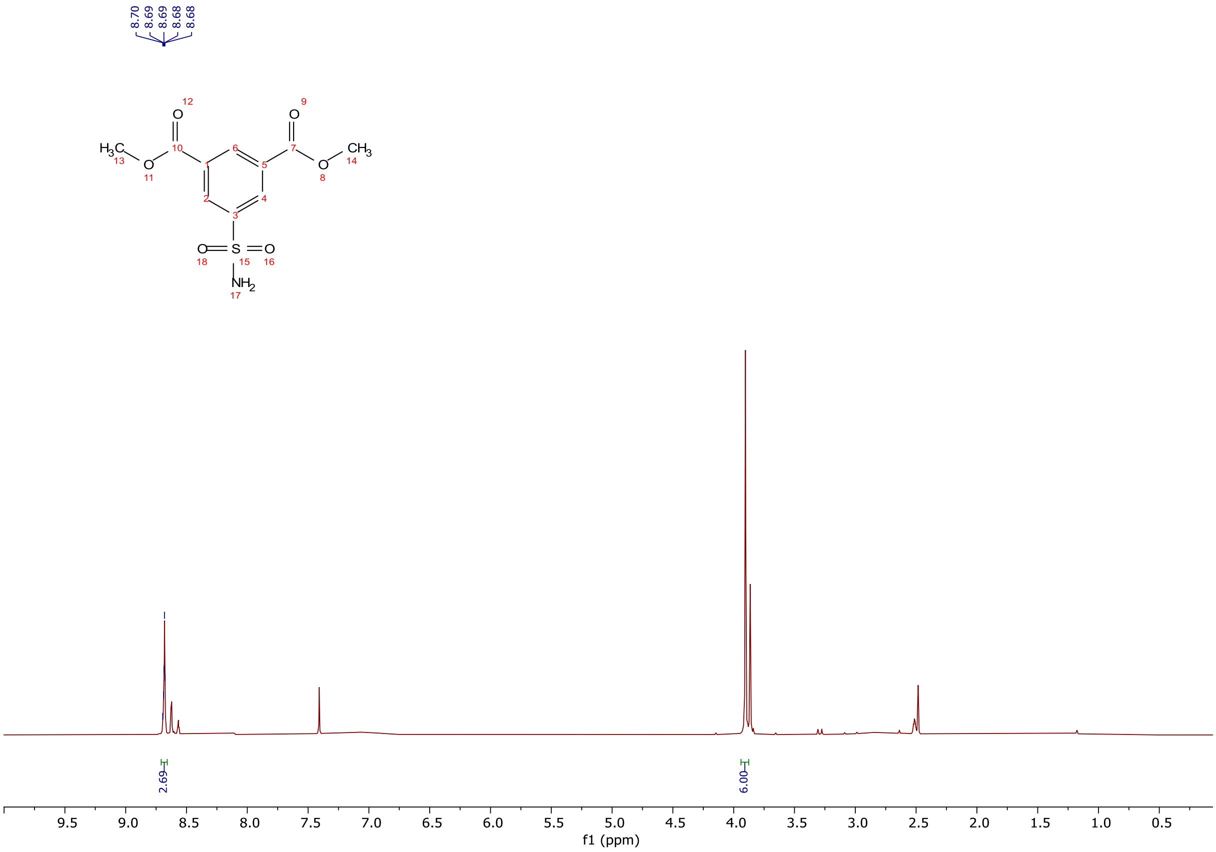

$^{1}H$ NMR spectra (300 MHz, $CDCl_3$) of dimethyl 5-sulfamoylisophthalate.

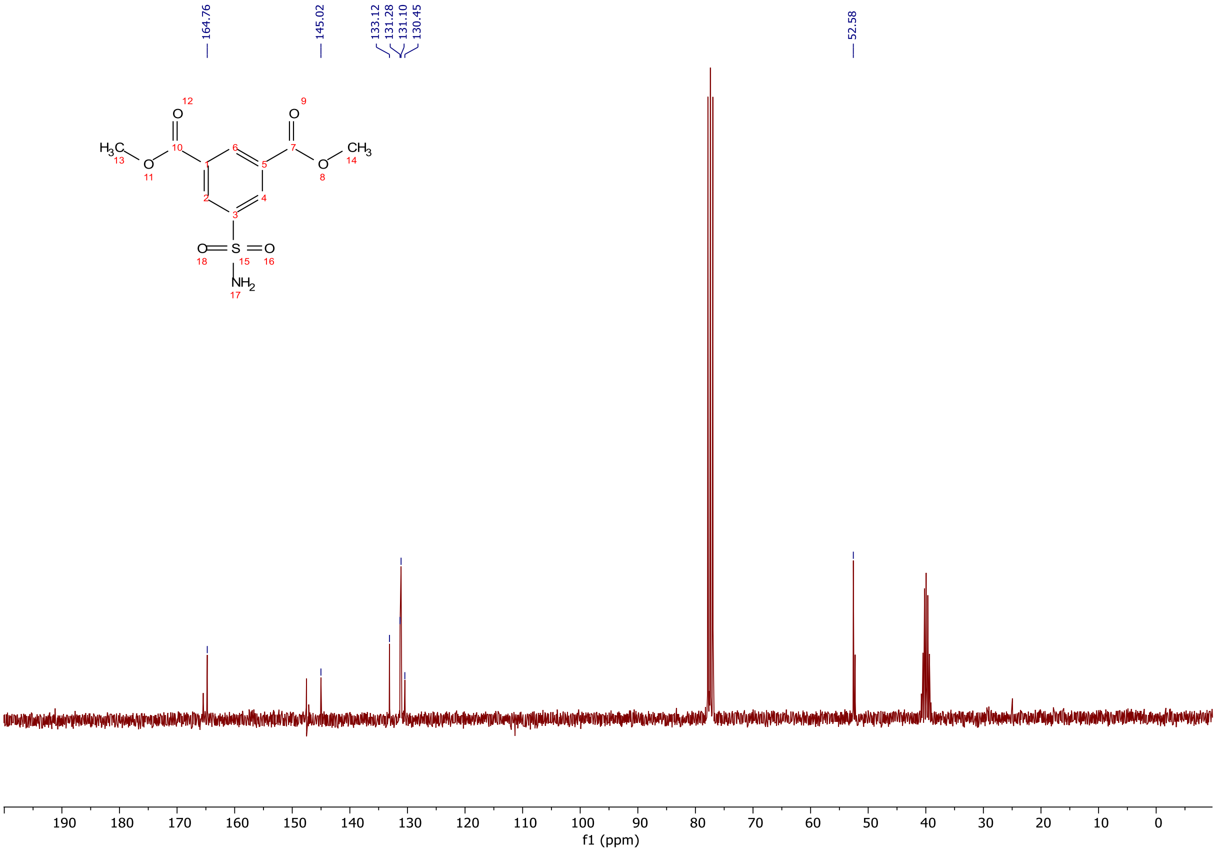

$^{13}C$ NMR spectra (75 MHz, $CDCl_3$) of dimethyl 5-sulfamoylisophthalate.

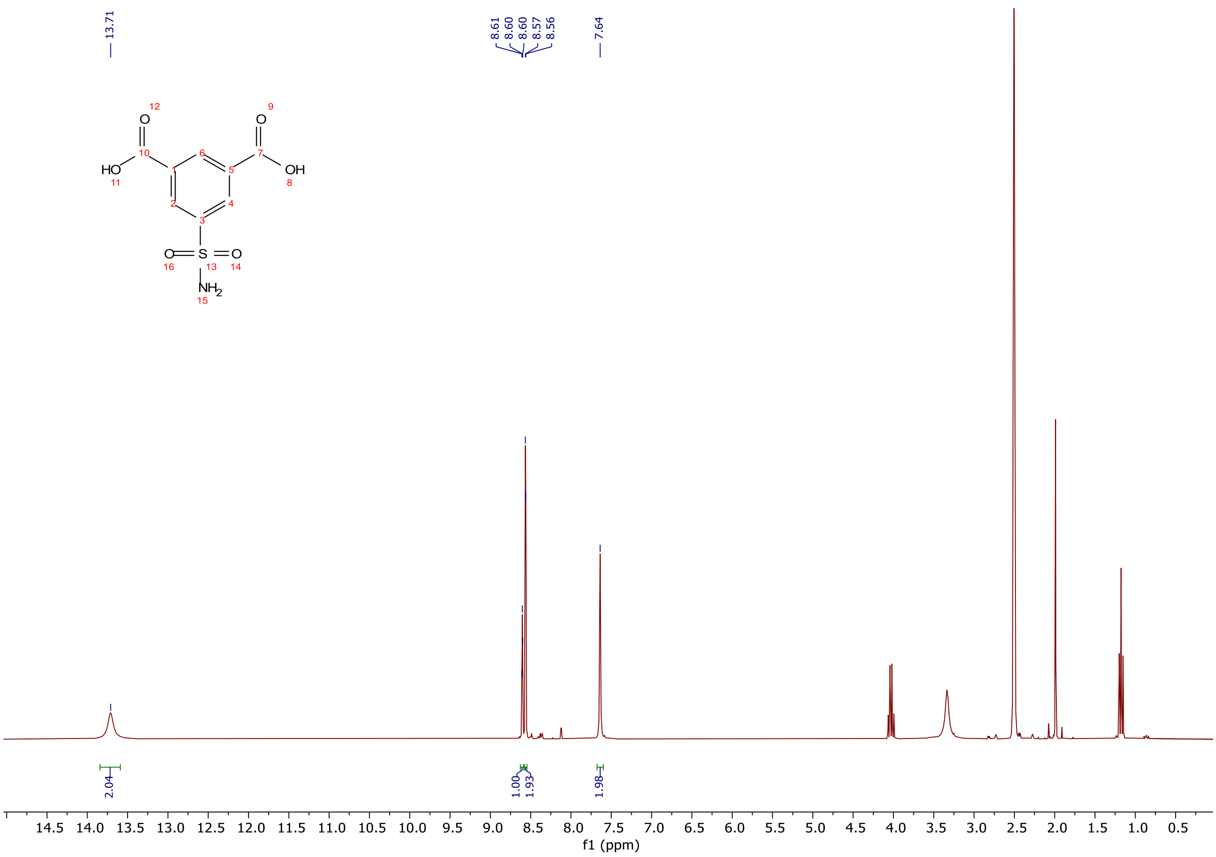


$^{1}H$ NMR spectra (300 MHz, DMSO-$d_6$) of 5-sulfamoylisophthalic acid.

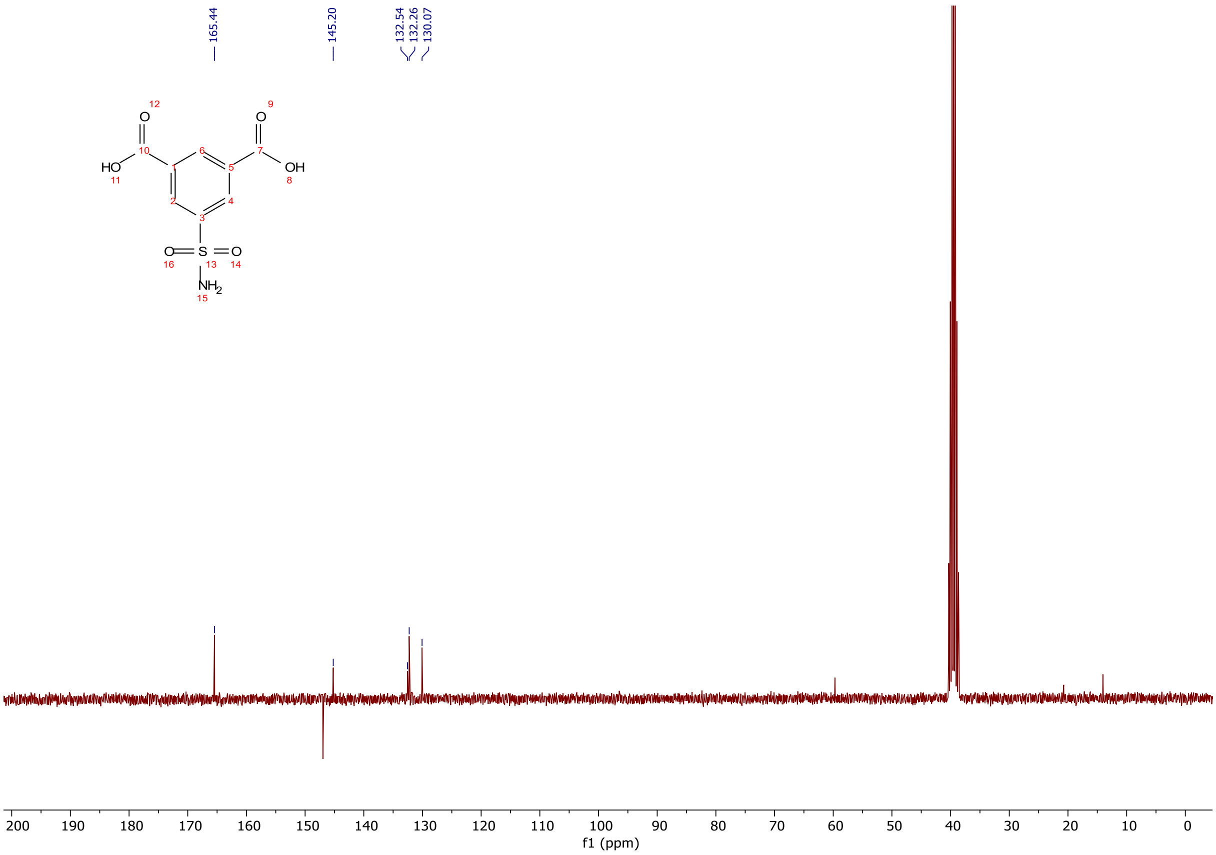


$^{13}C$ NMR spectra (75 MHz, DMSO-$d_6$) of 5-sulfamoylisophthalic acid.

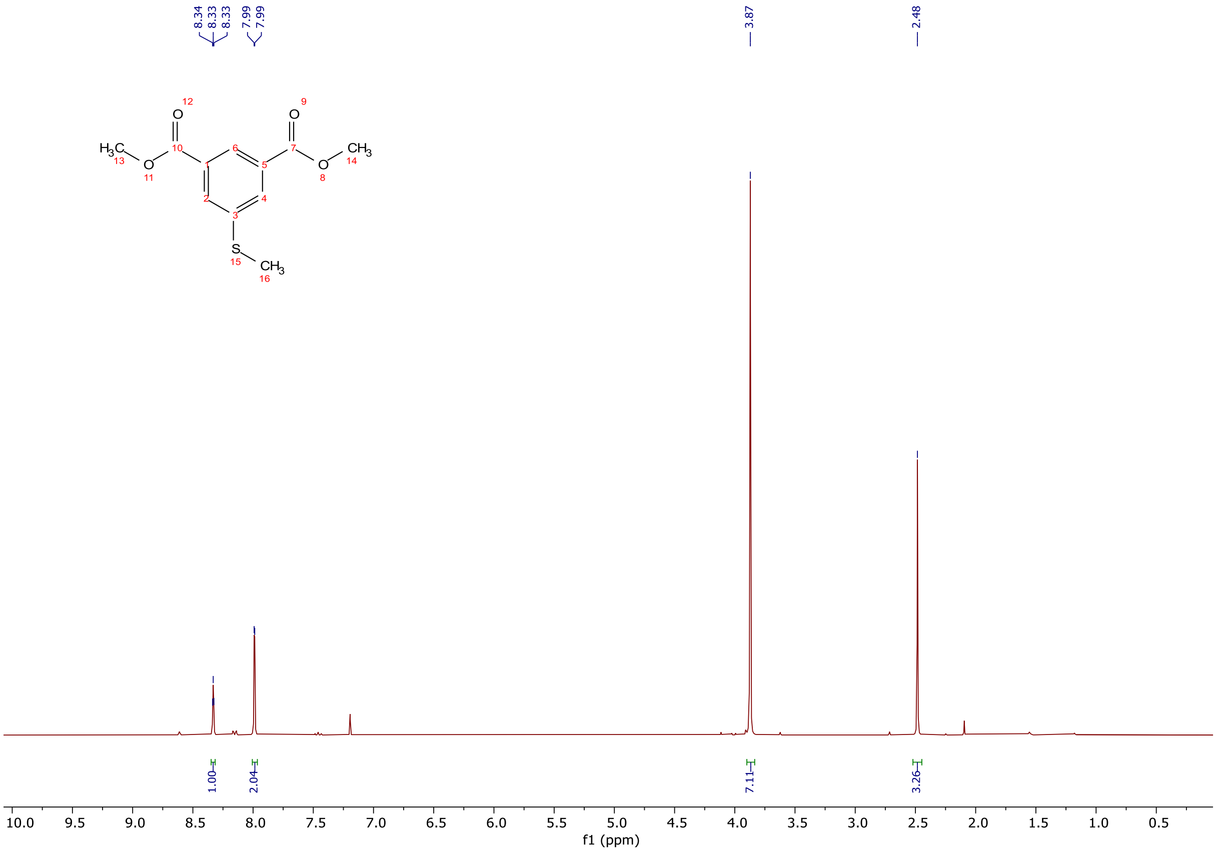


$^{1}$H NMR spectra (300 MHz, $CDCl_3$) of dimethyl 5-(methylthio)isophthalate.

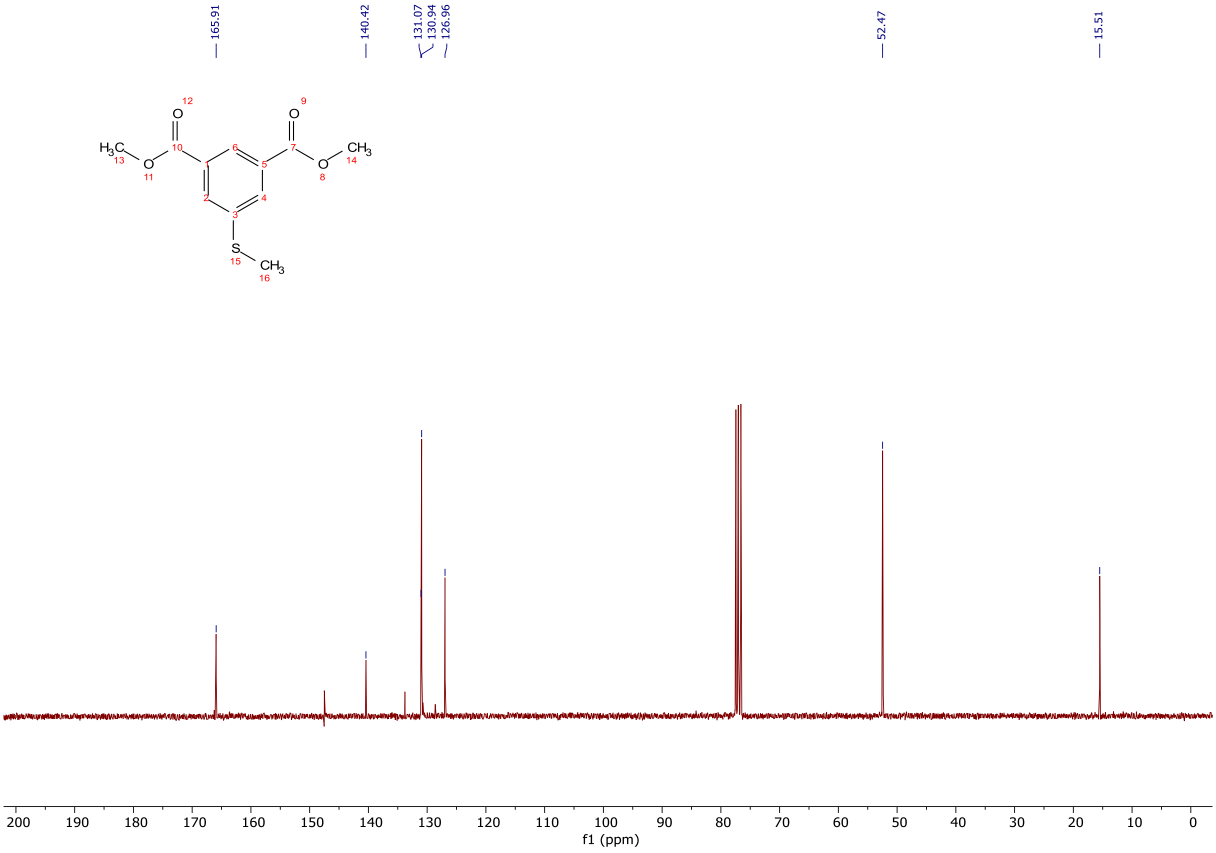


$^{13}$C NMR spectra (75 MHz, $CDCl_3$) of dimethyl 5-(methylthio)isophthalate.

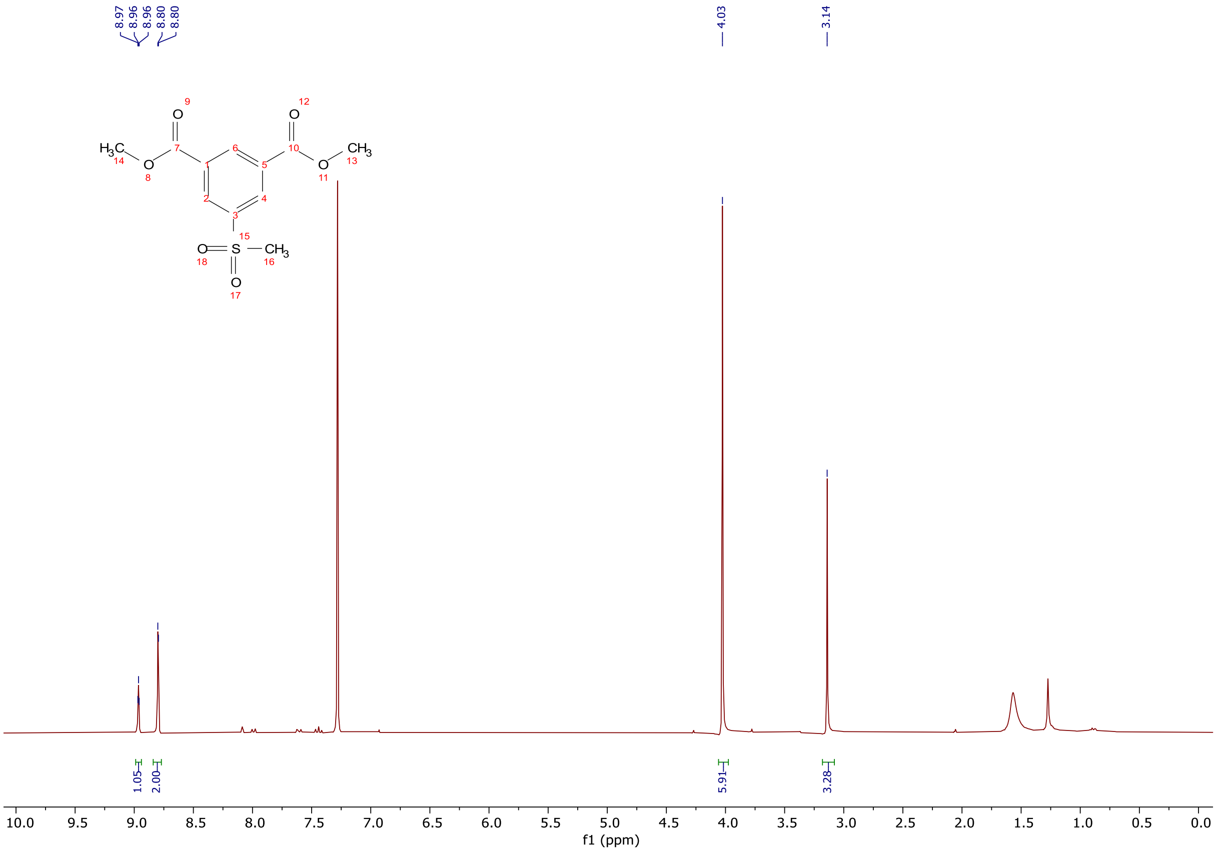


$^{1}H$ NMR spectra (300 MHz, $CDCl_3$) of dimethyl 5-(methylsulfonyl)isophthalate.

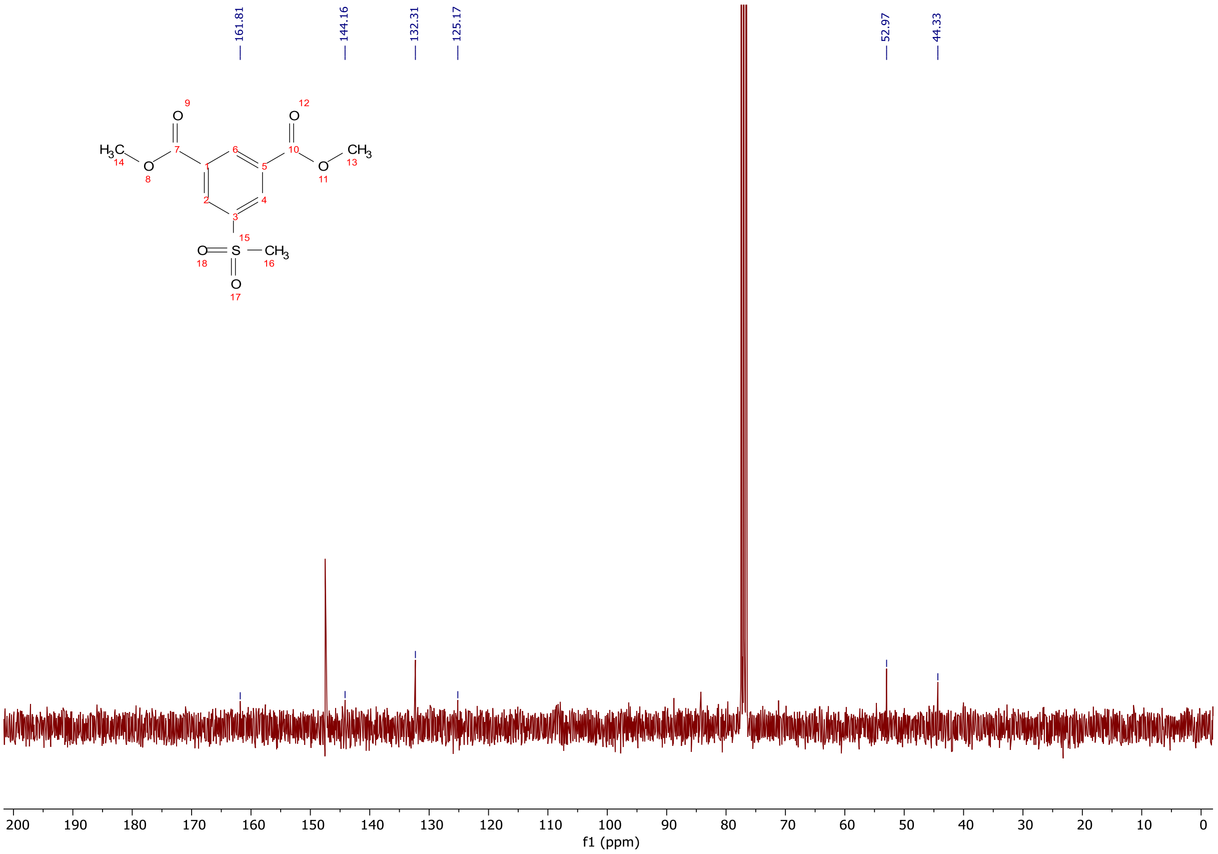


$^{13}C$ NMR spectra (75 MHz, $CDCl_3$) of dimethyl 5-(methylsulfonyl)isophthalate.

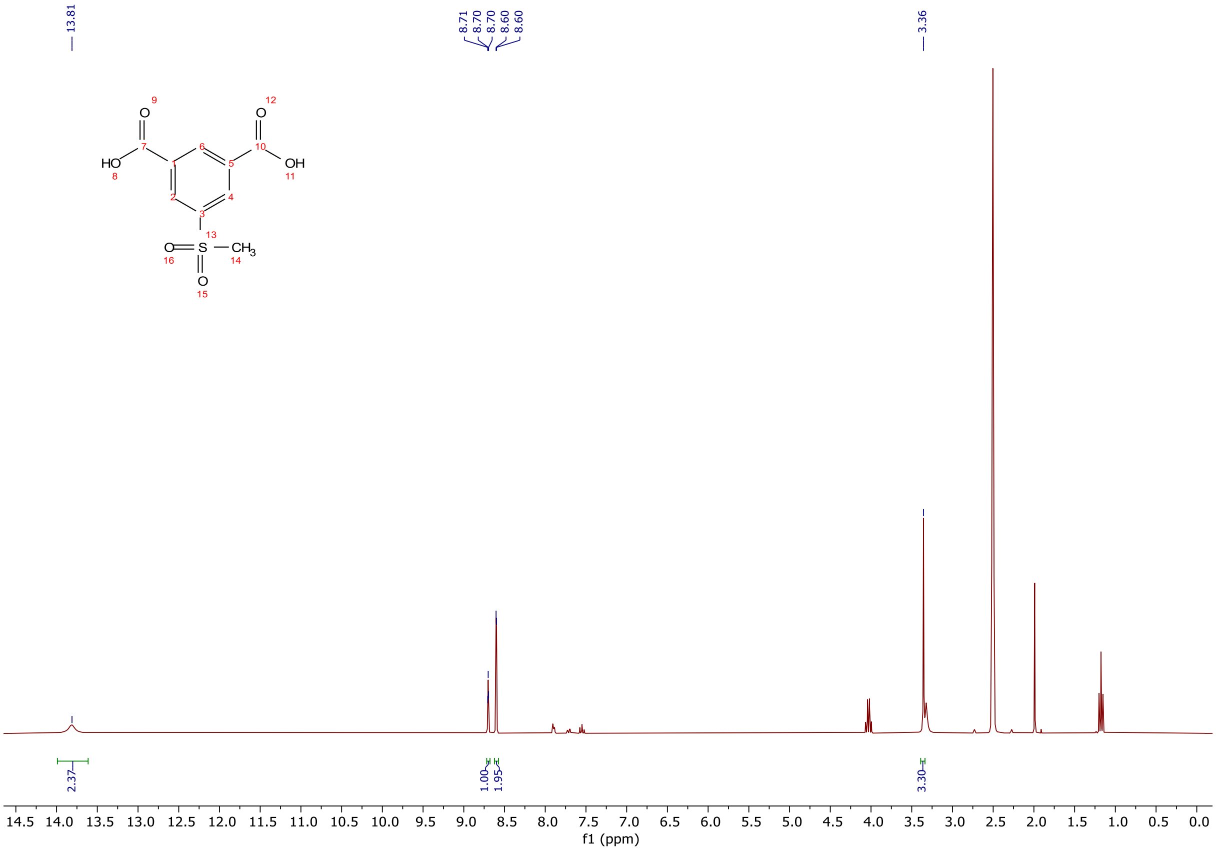


$^{1}H$ NMR spectra (300 MHz, DMSO-$d_6$) of 5-(methylsulfonyl)isophthalic acid.

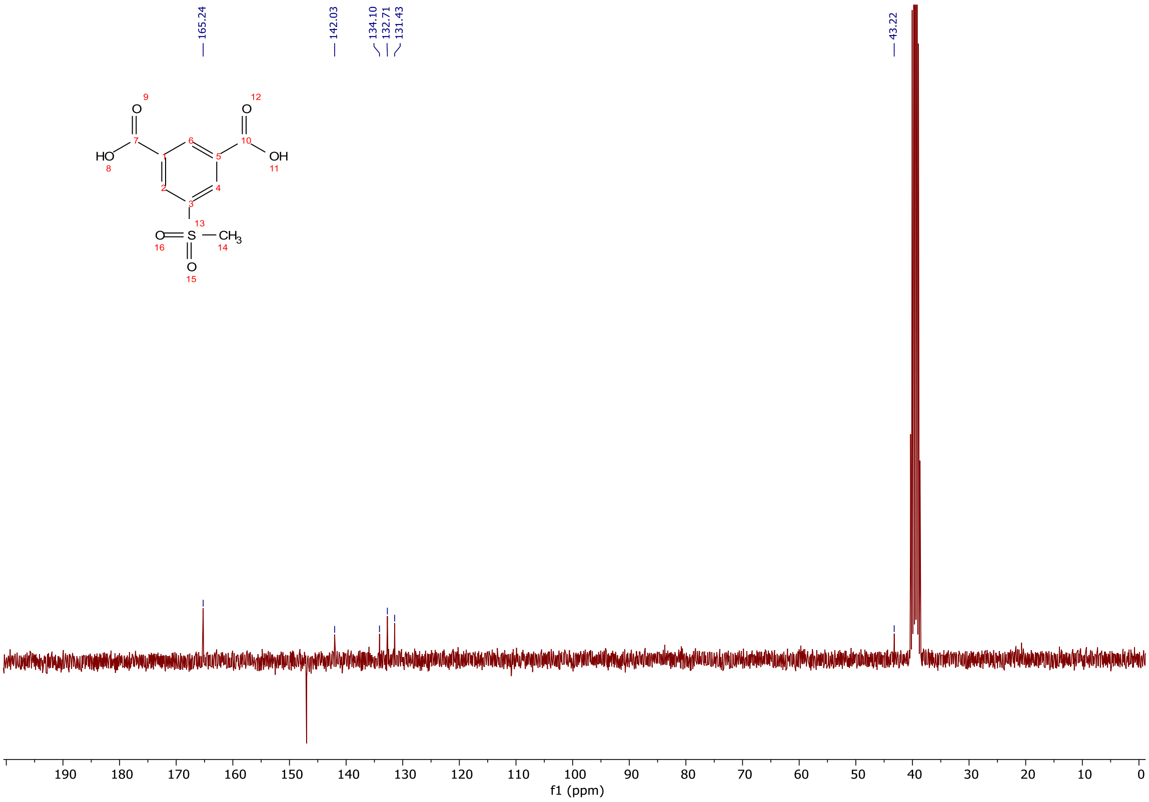


$^{13}C$ NMR spectra (75 MHz, DMSO-$d_6$) of 5-(methylsulfonyl)isophthalic acid.

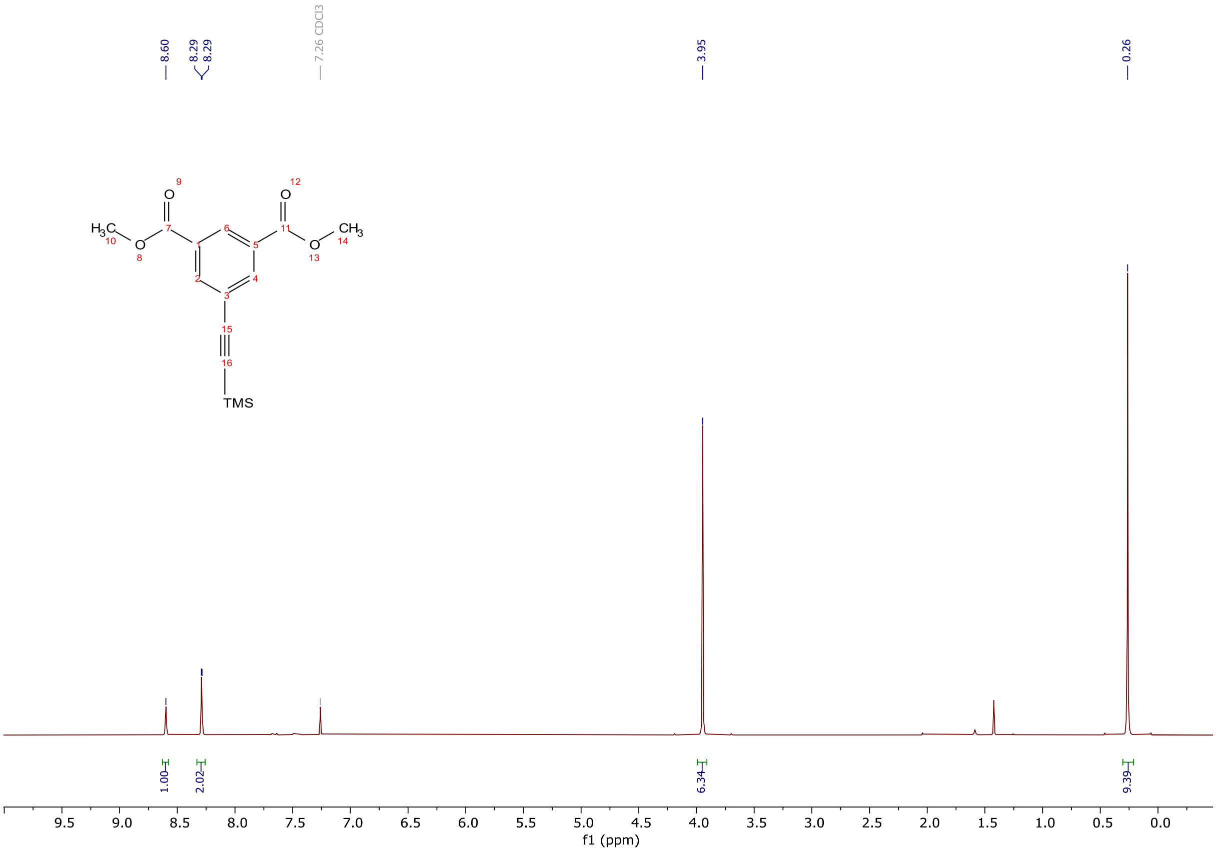


$^{1}H$ NMR spectra (300 MHz, $CDCl_3$) of dimethyl 5-((trimethylsilyl)ethynyl)isophthalate.

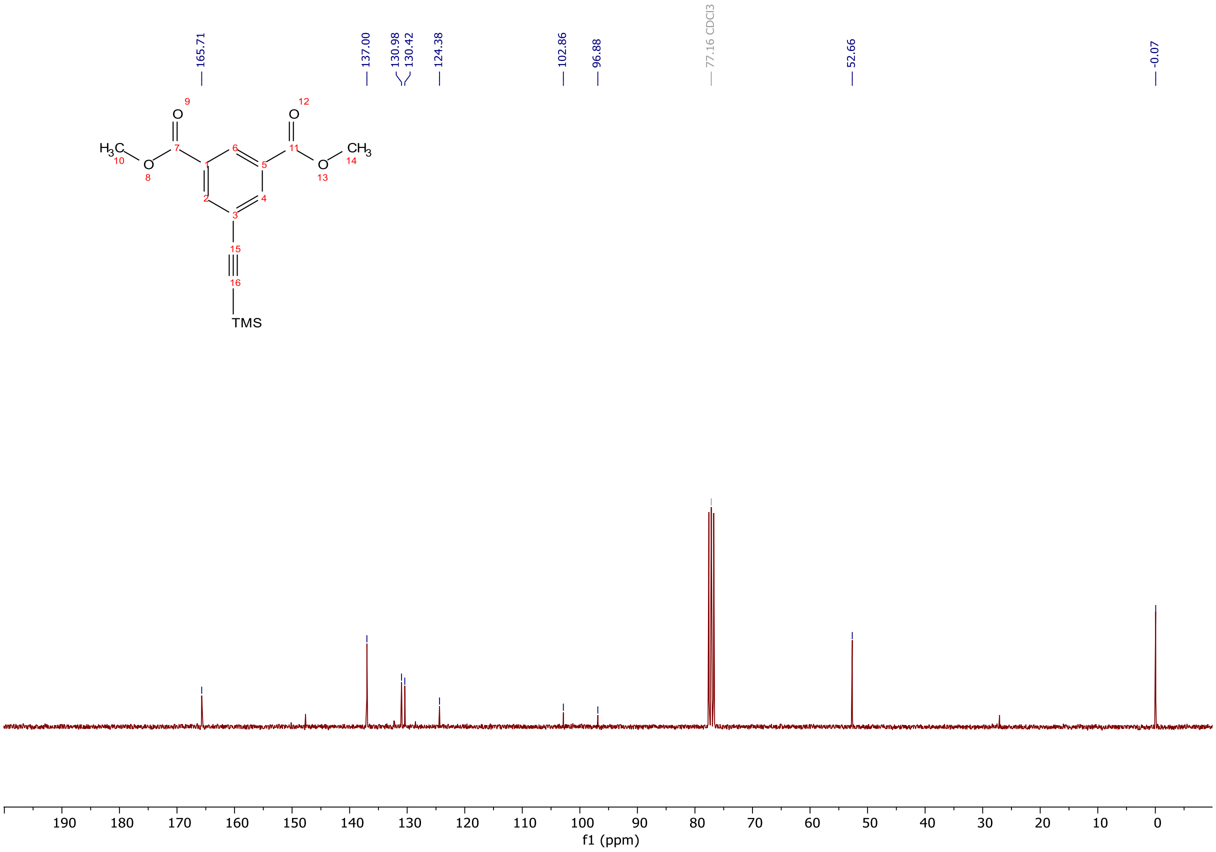


$^{13}C$ NMR spectra (75 MHz, $CDCl_3$) of dimethyl 5-((trimethylsilyl)ethynyl)isophthalate.

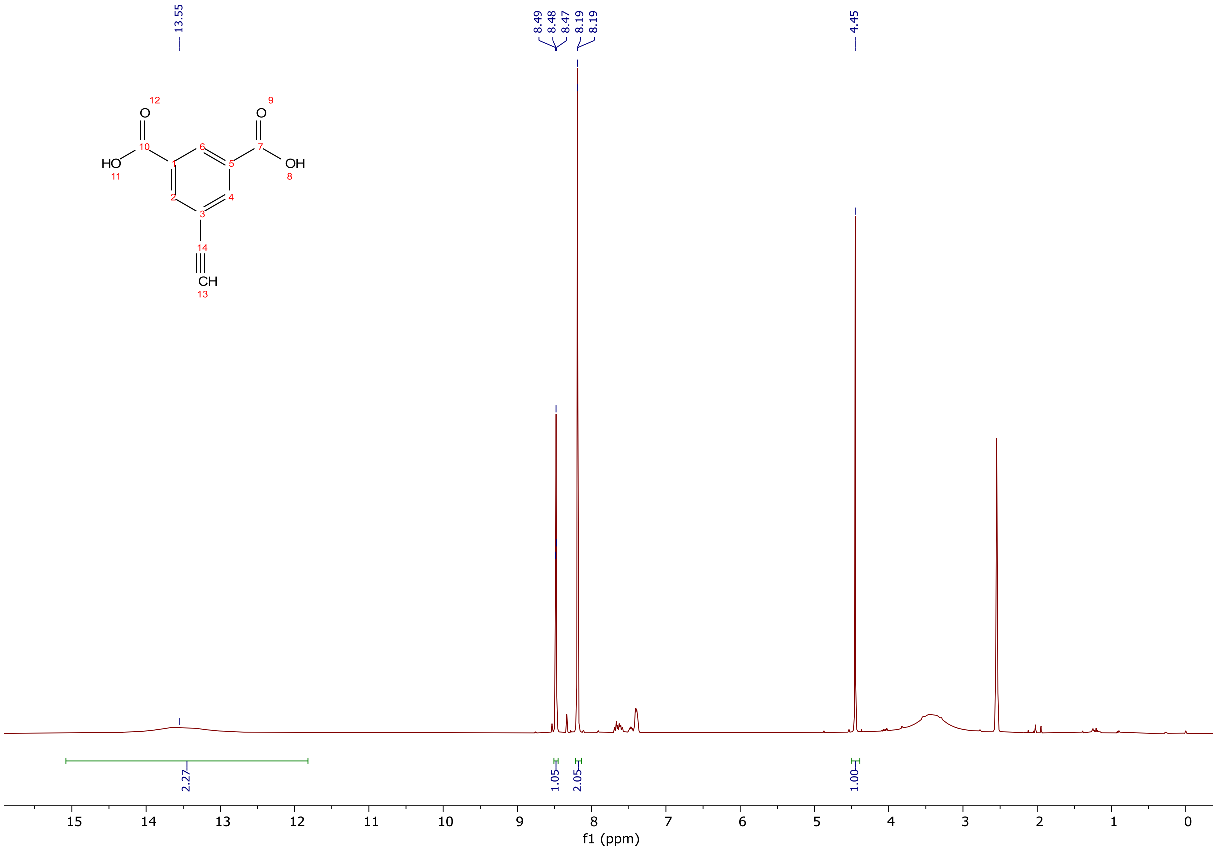


$^1$H NMR spectra (300 MHz, DMSO-$d_6$) of 5-ethynylisophthalic acid.

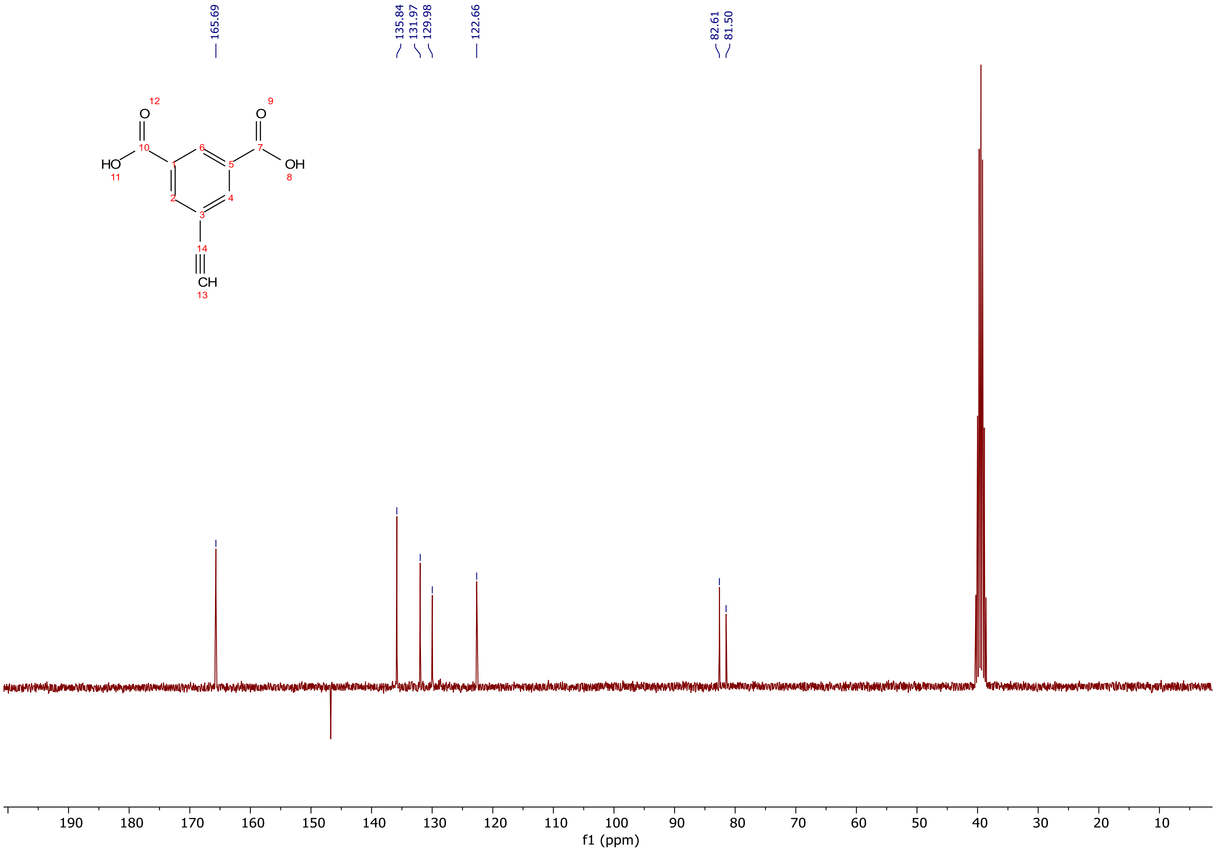


$^{13}$C NMR spectra (75 MHz, DMSO-$d_6$) of 5-ethynylisophthalic acid.

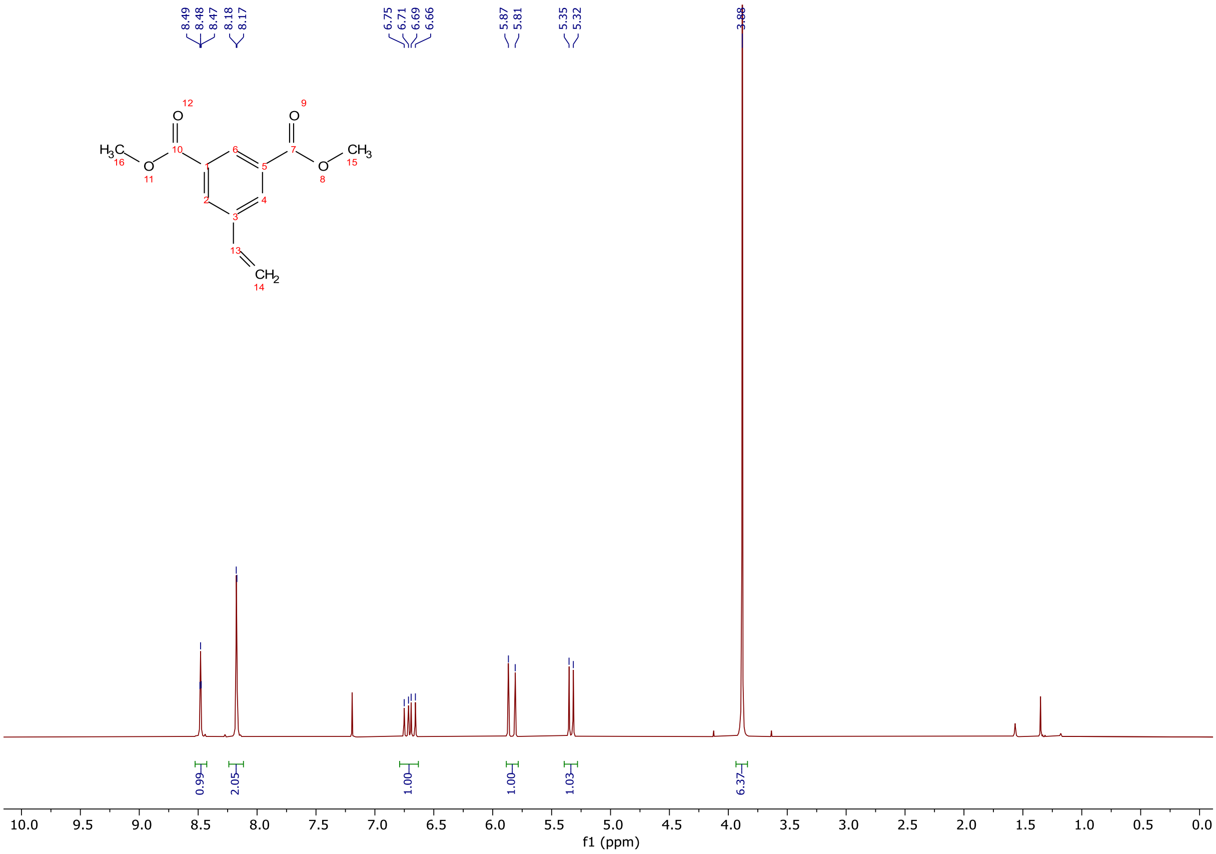


[1]H NMR spectra (300 MHz, $CDCl_3$) of dimethyl 5-vinylisophthalate.

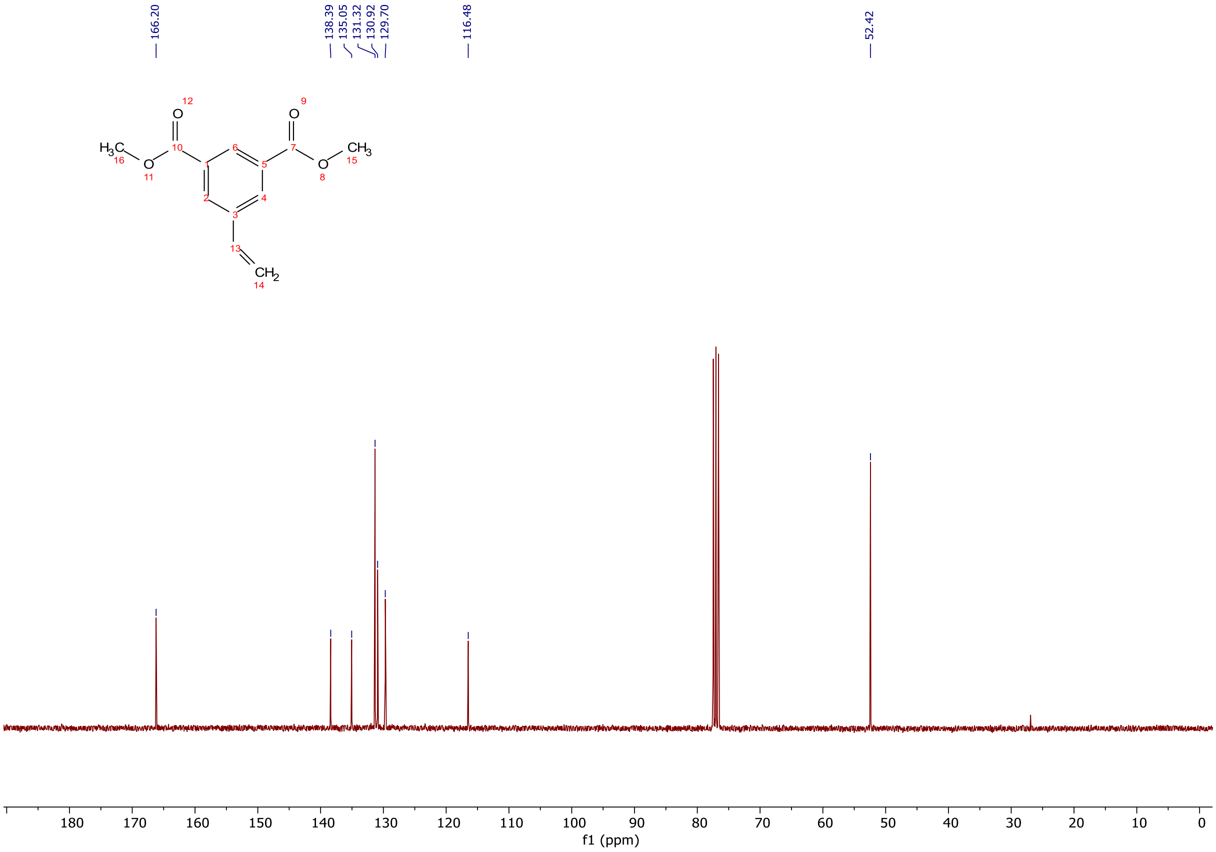


[13]C NMR spectra (75 MHz, $CDCl_3$) of dimethyl 5-vinylisophthalate.

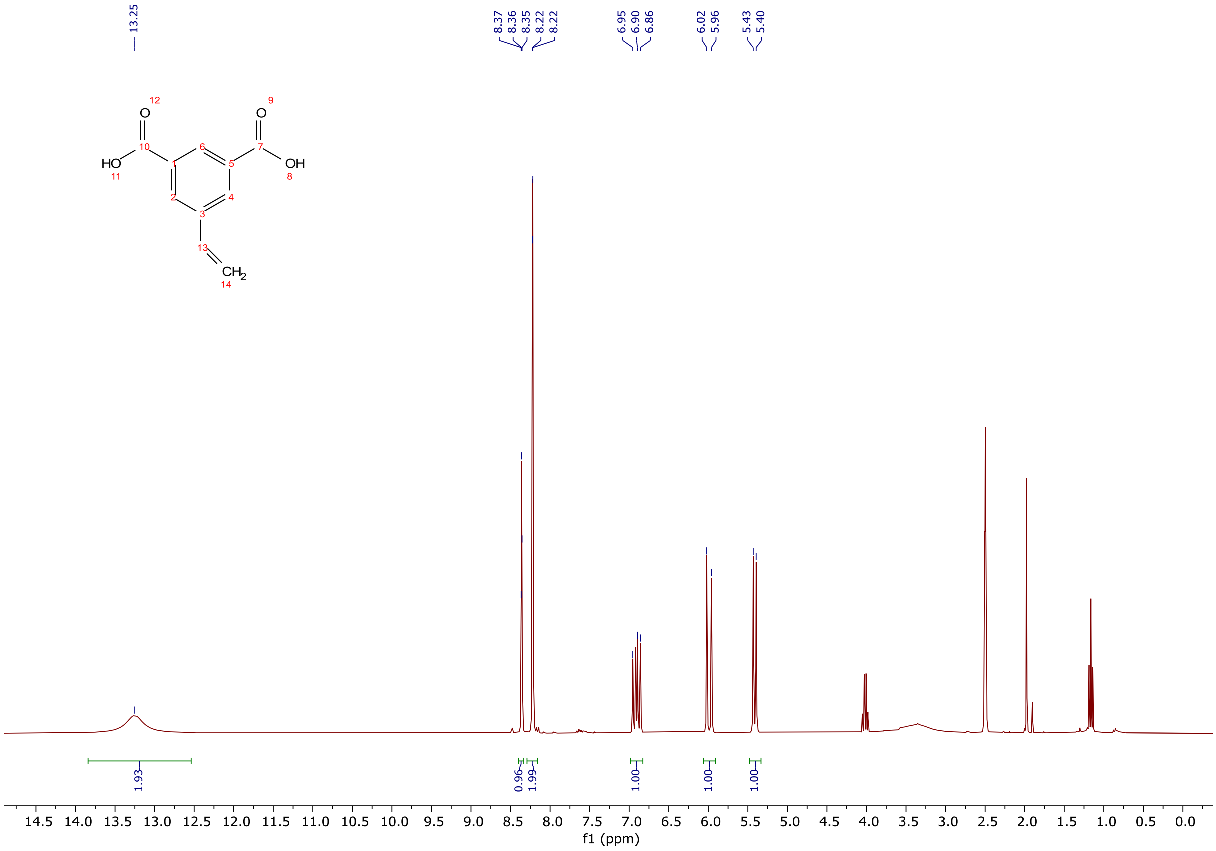


$^{1}$H NMR spectra (300 MHz, DMSO-d$_6$) of 5-vinylisophthalic acid.

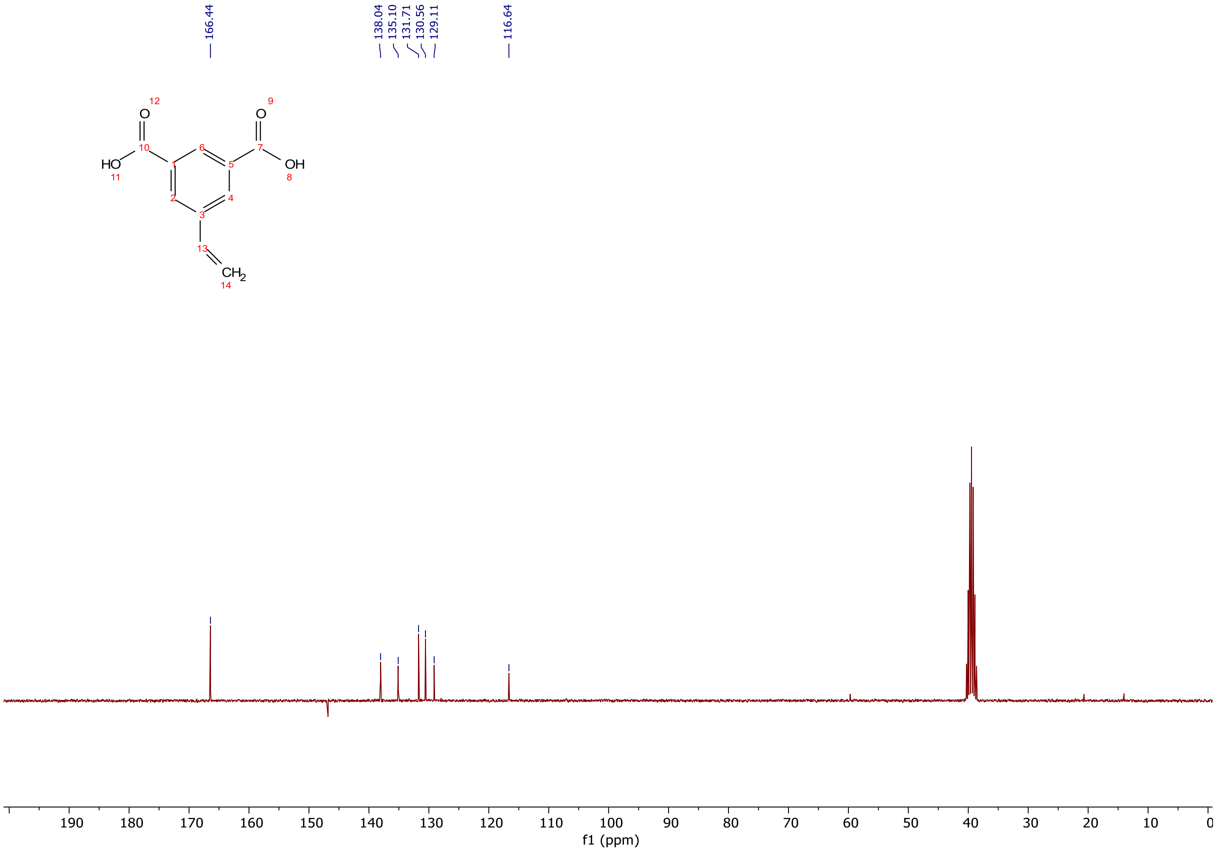


$^{13}$C NMR spectra (75 MHz, DMSO-d$_6$) of 5-vinylisophthalic acid.

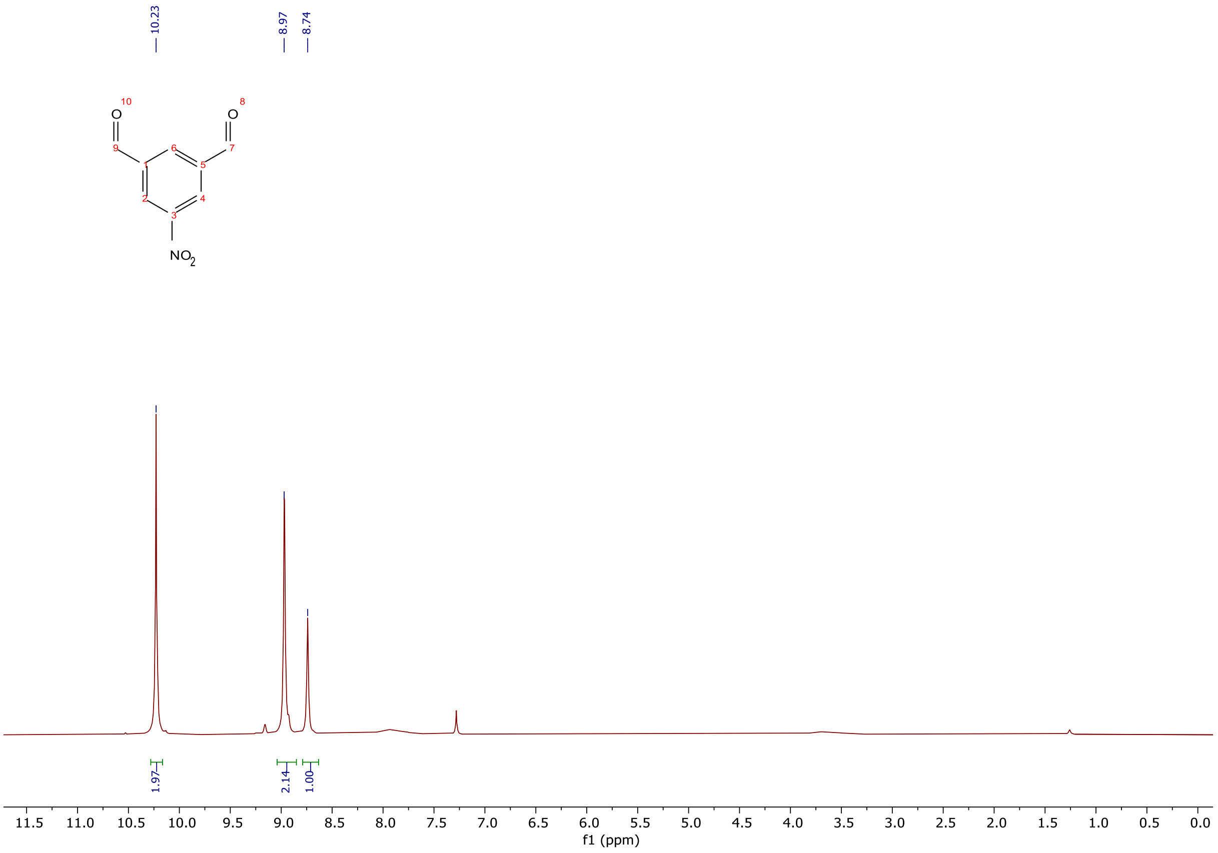


$^{1}$H NMR spectra (300 MHz, $CDCl_3$) of 5-nitroisophthalaldehyde.

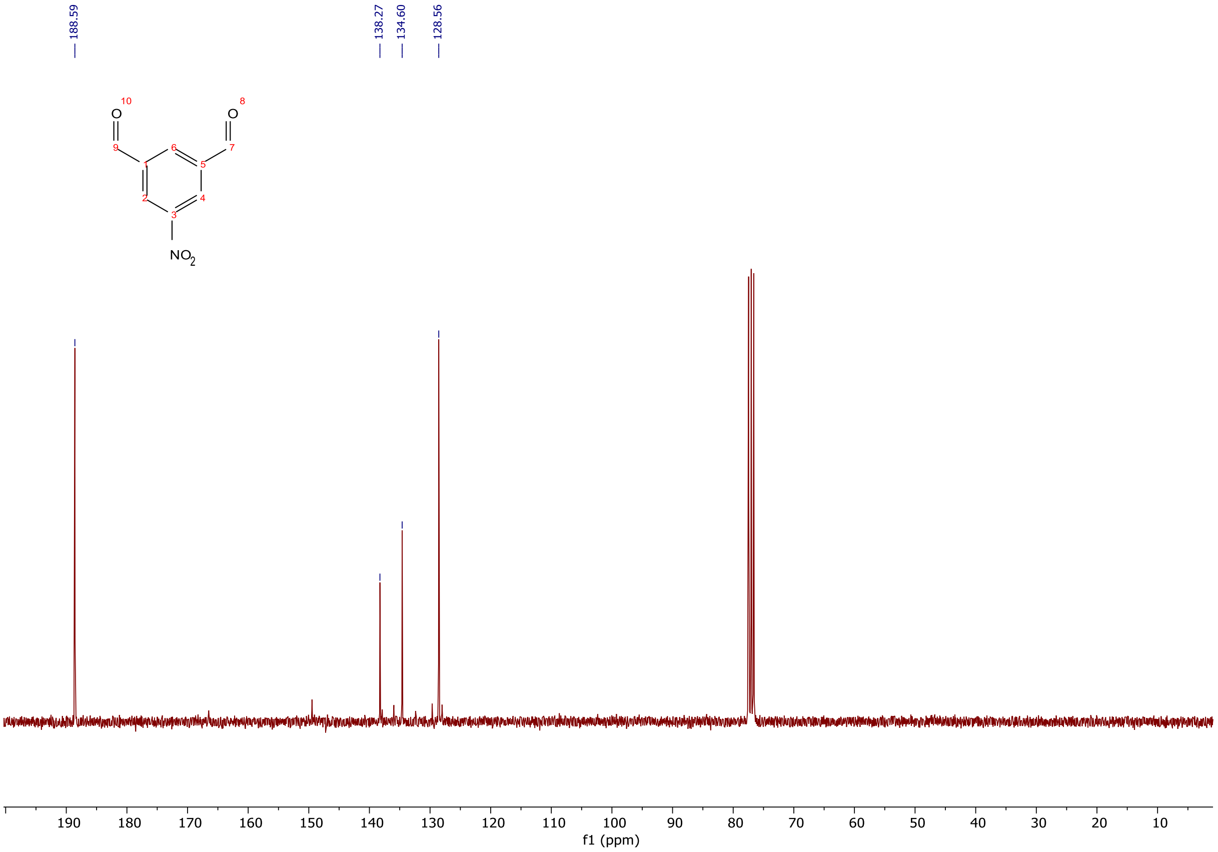


$^{13}$C NMR spectra (75 MHz, $CDCl_3$) of 5-nitroisophthalaldehyde.

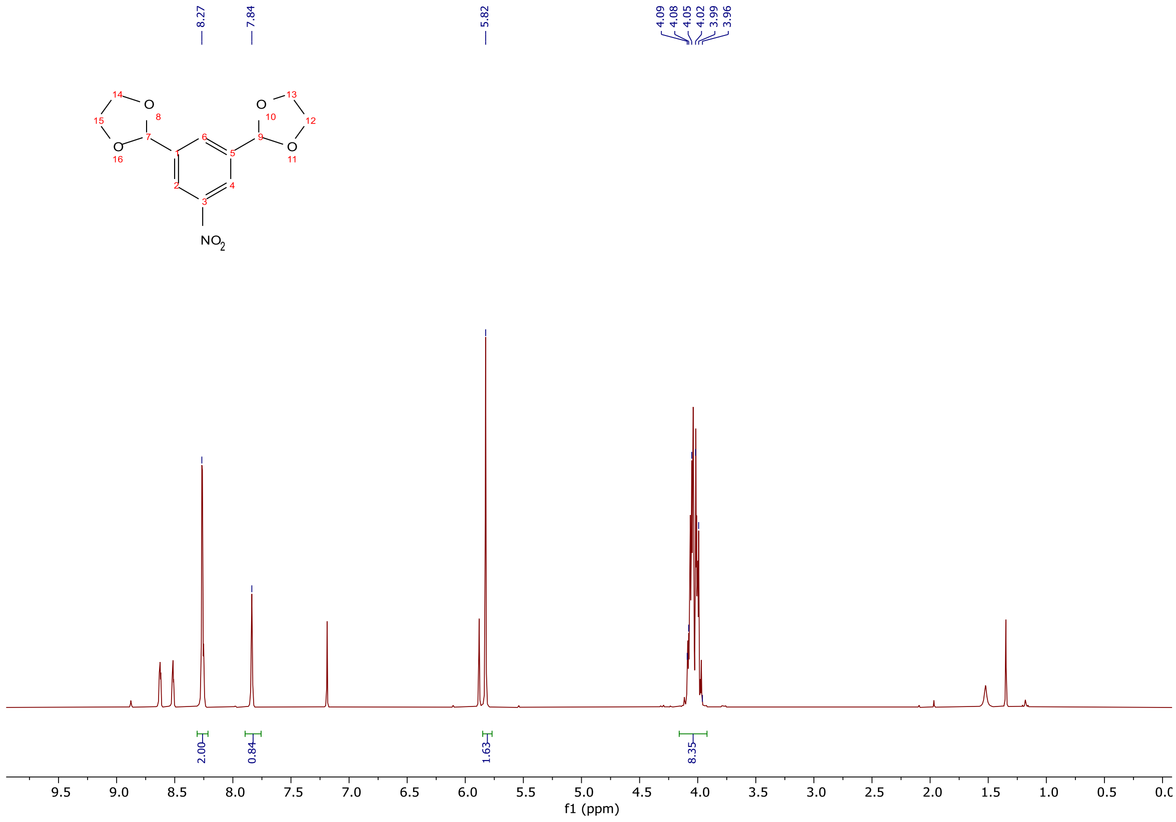


$^{1}H$ NMR spectra (300 MHz, $CDCl_3$) of 2,2'-(5-nitro-1,3-phenylene)bis(1,3-dioxolane).

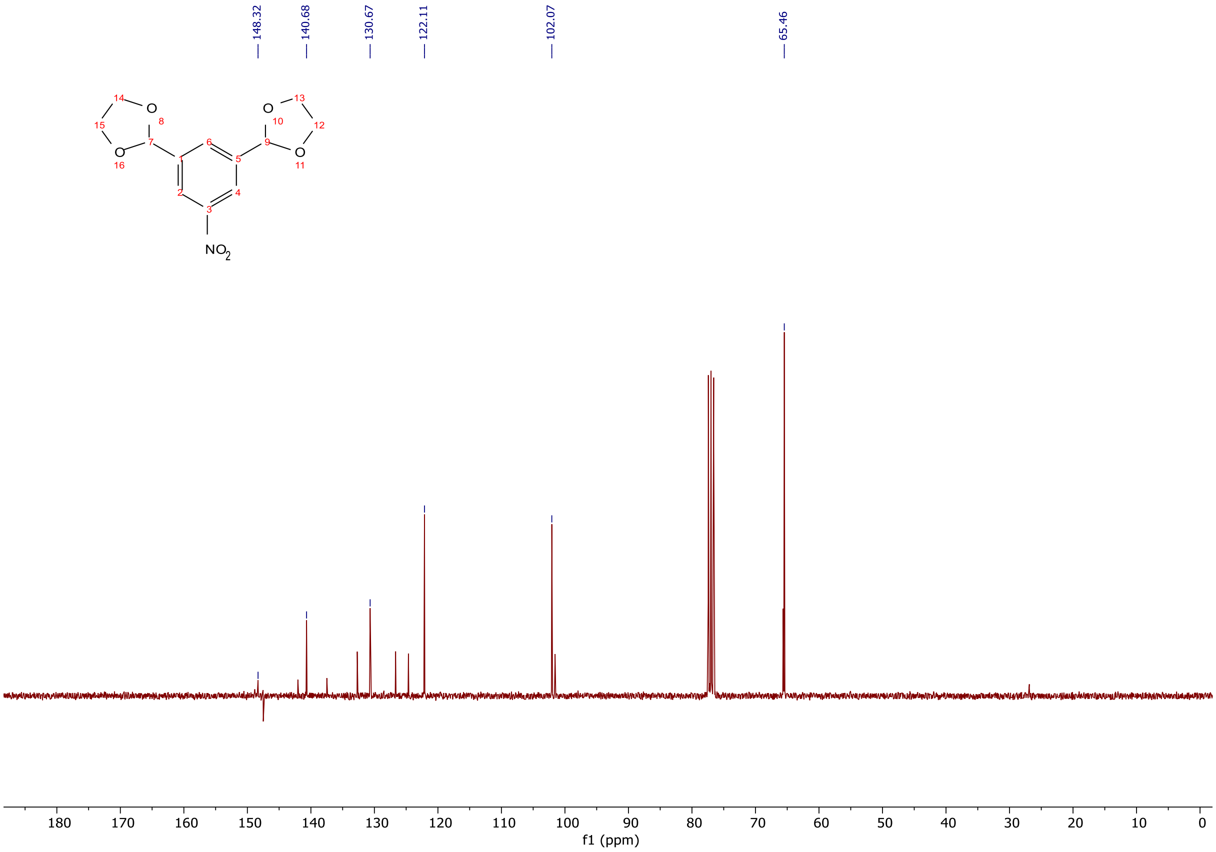


$^{13}C$ NMR spectra (75 MHz, $CDCl_3$) of 2,2'-(5-nitro-1,3-phenylene)bis(1,3-dioxolane).

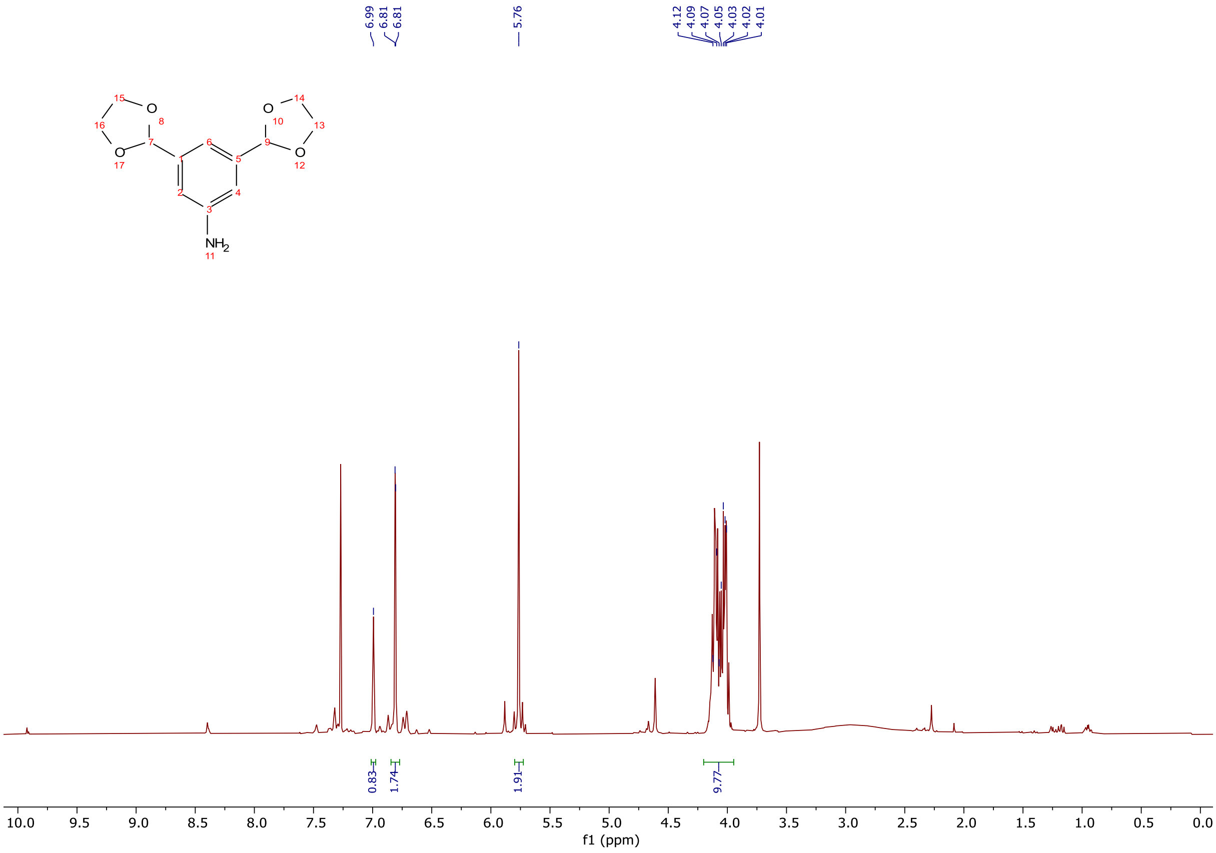

[1]H NMR spectra (300 MHz, $CDCl_3$) of 3,5-di(1,3-dioxolan-2-yl)aniline.

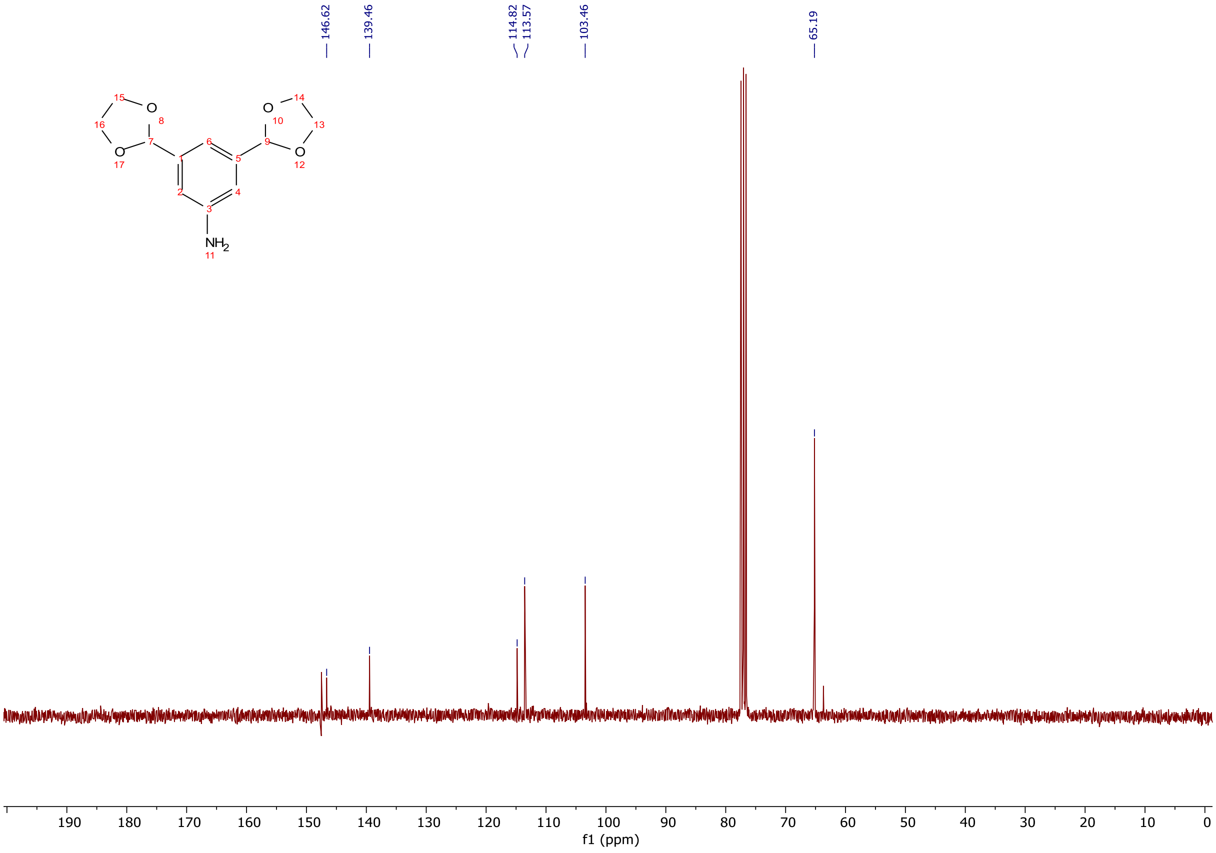

[13]C NMR spectra (75 MHz, $CDCl_3$) of 3,5-di(1,3-dioxolan-2-yl)aniline.

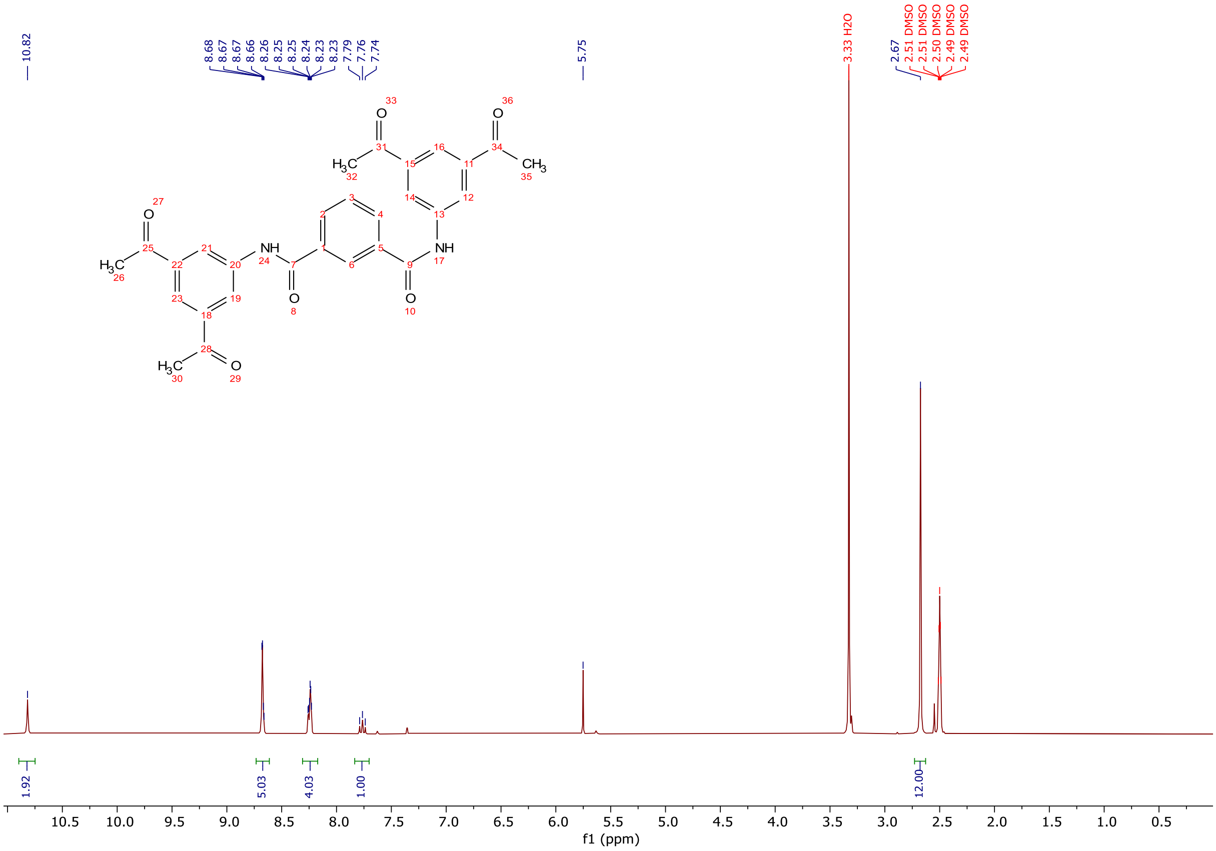


$^{1}$H NMR spectra (300 MHz, DMSO-d$_6$) of **1-1**.

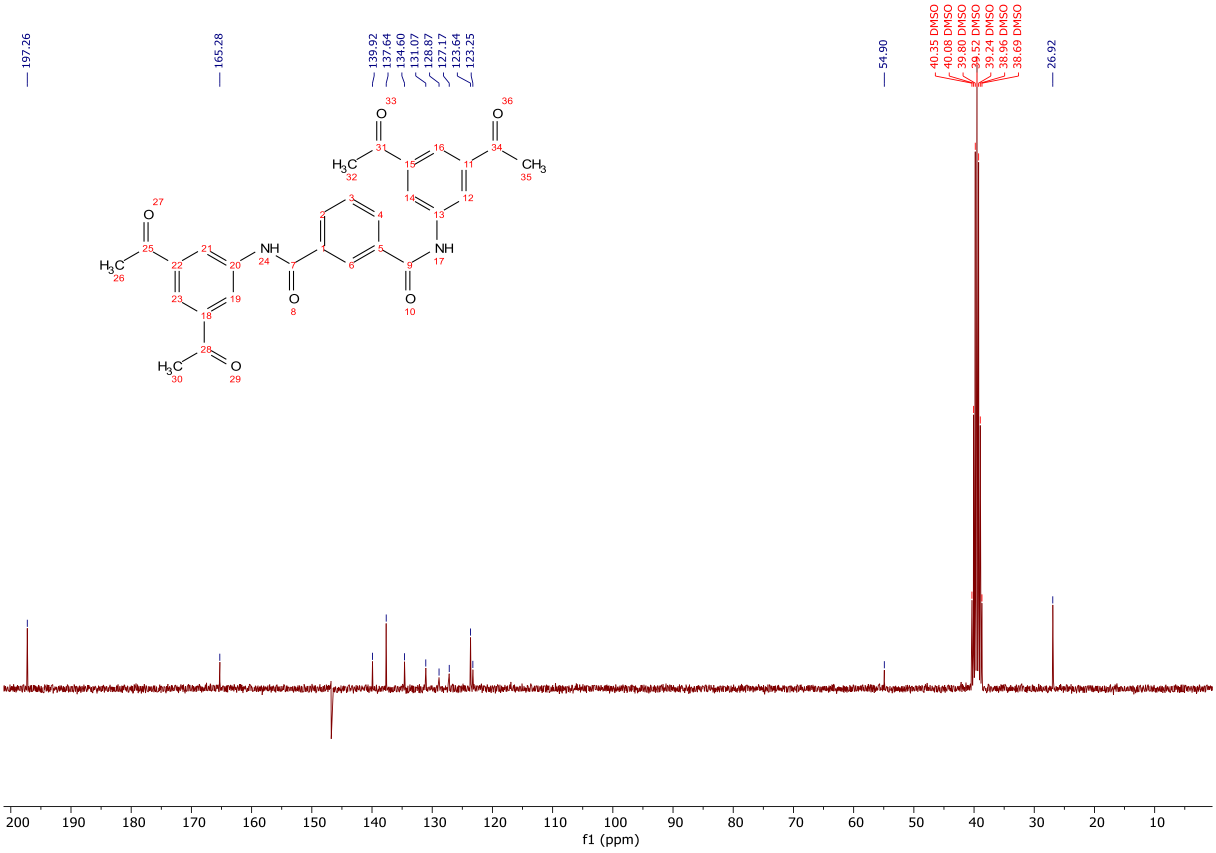


$^{13}$C NMR spectra (75 MHz, DMSO-d$_6$) of **1-1**.

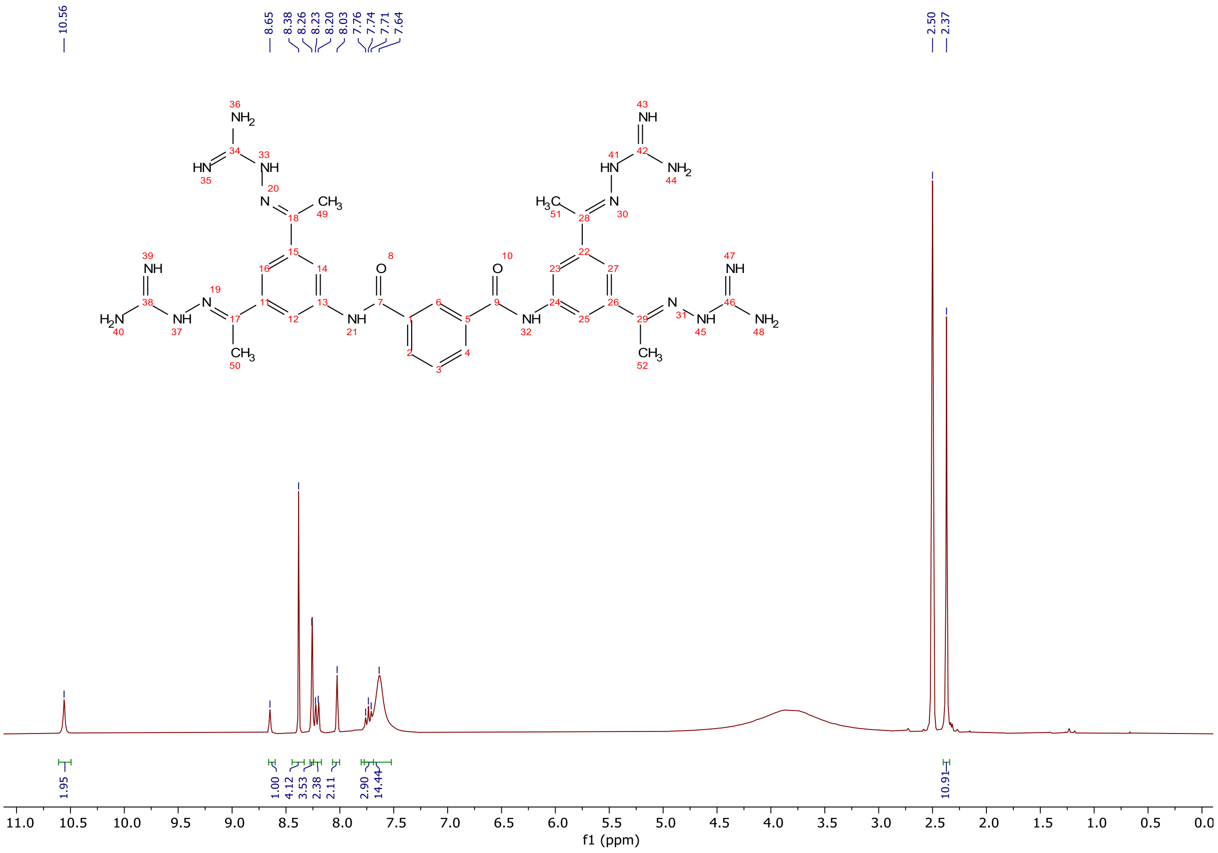

$^{1}H$ NMR spectra (300 MHz, DMSO-$d_6$) of **1** (TCB-32).

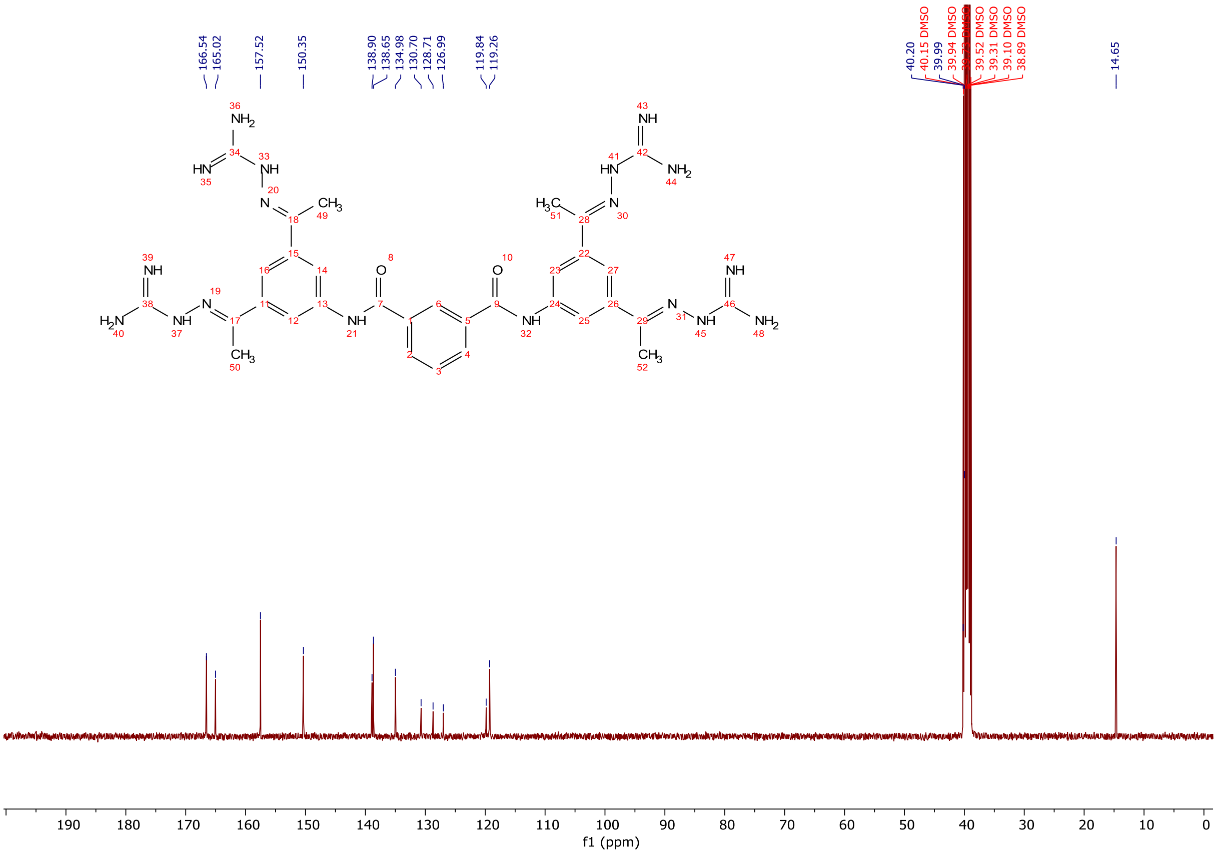

$^{13}C$ NMR spectra (300 MHz, DMSO-$d_6$) of (TCB-32).

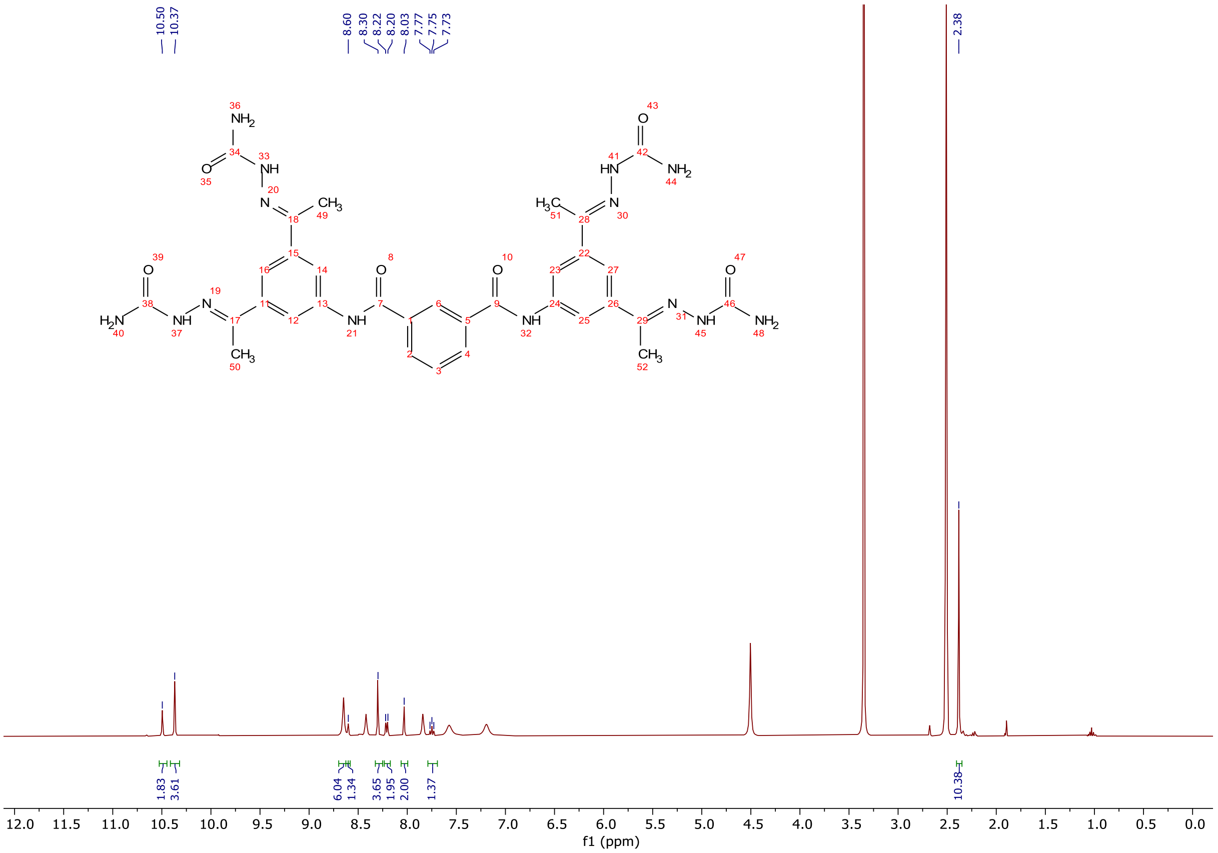

$^1$H NMR spectra (400 MHz, DMSO-d$_6$) of **2**.

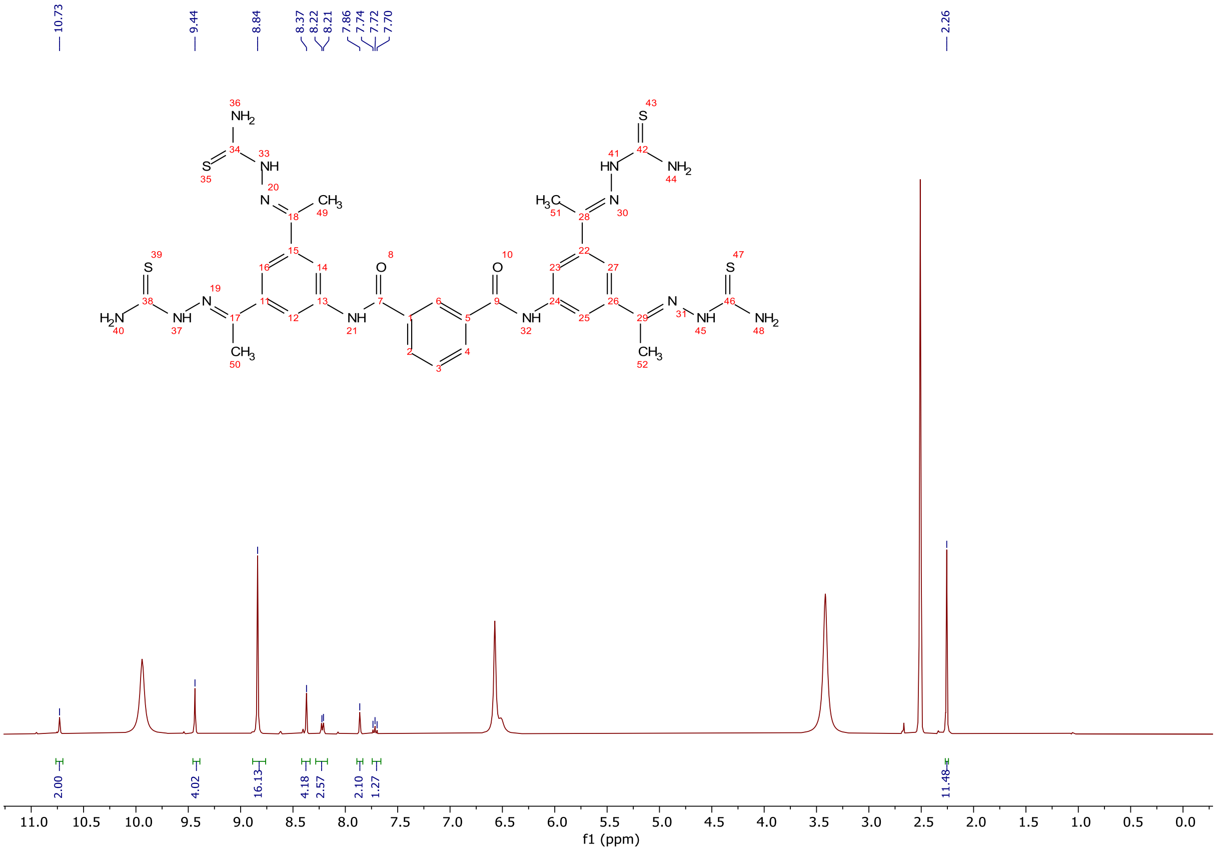

$^1$H NMR spectra (300 MHz, DMSO-d$_6$) of **3**.

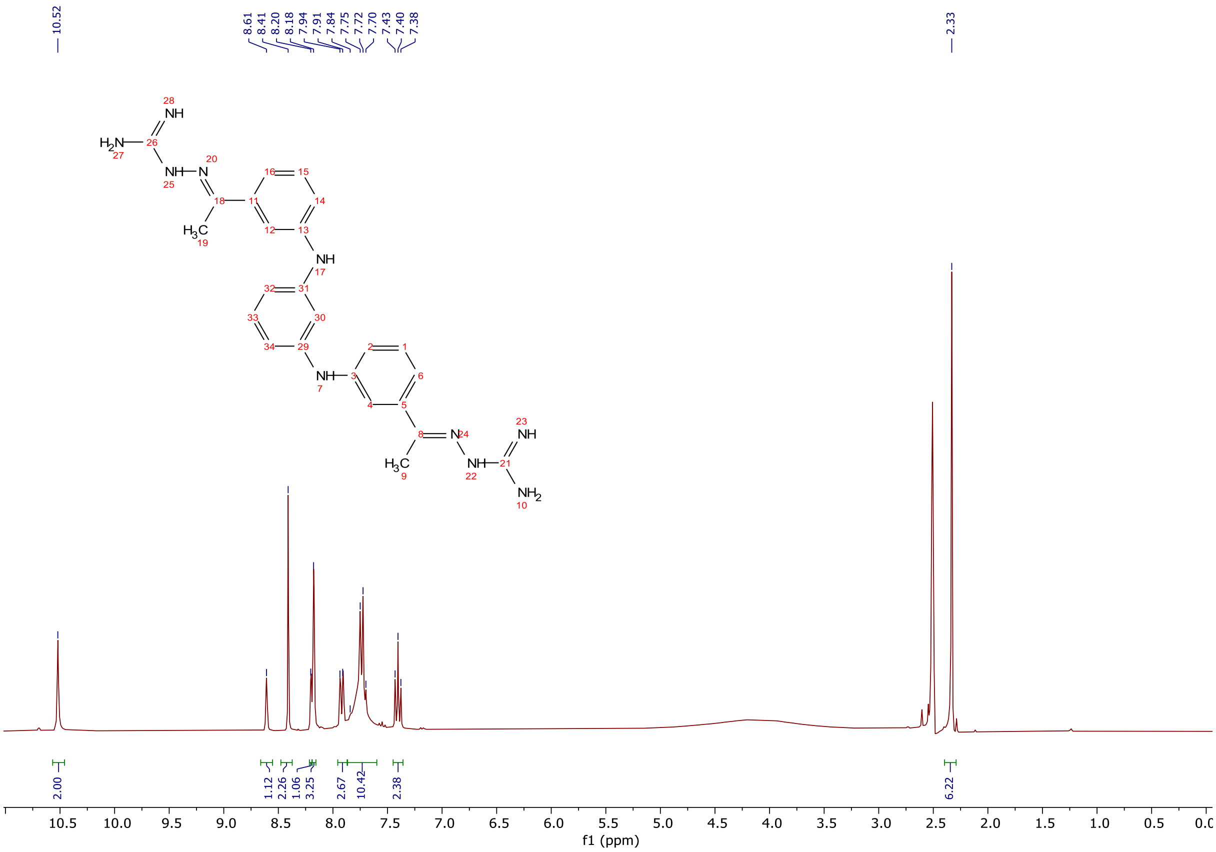

$^1$H NMR spectra (300 MHz, DMSO-$d_6$) of **4**.

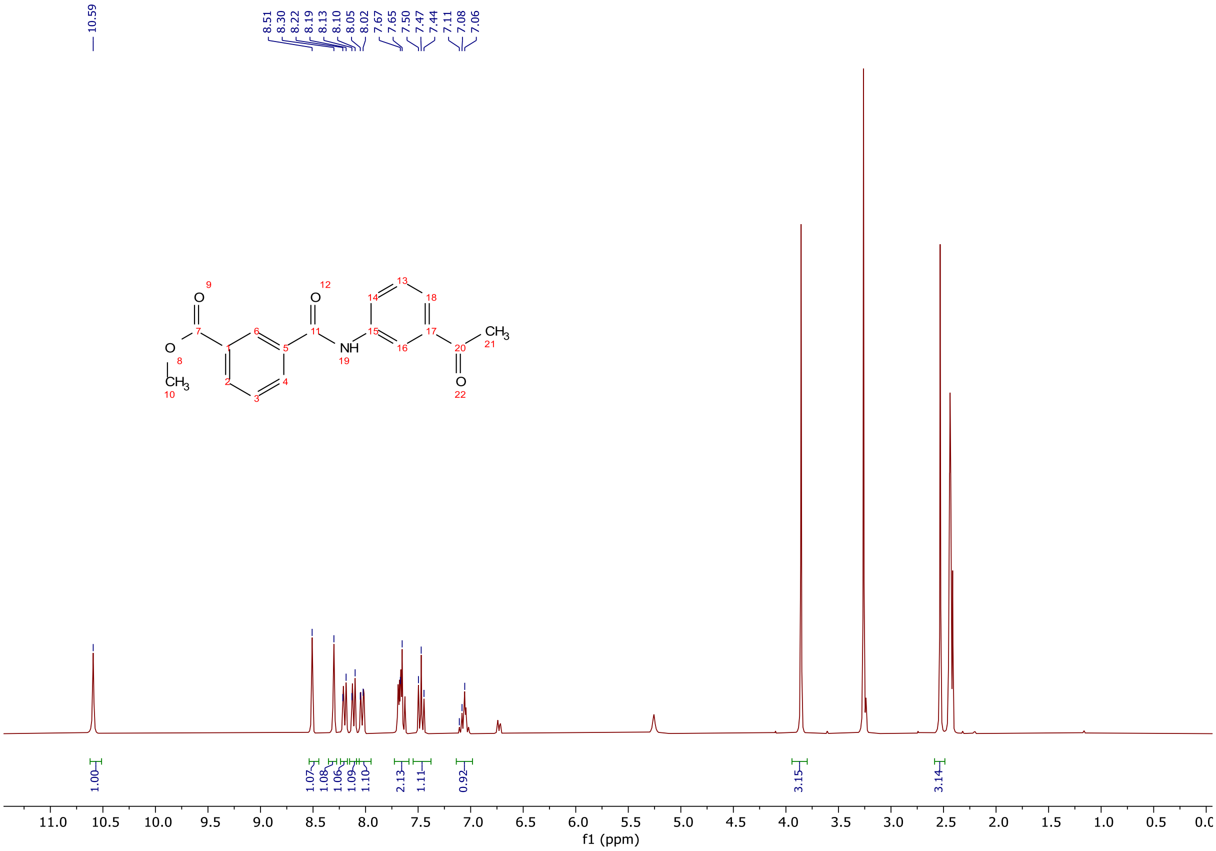

$^1$H NMR spectra (300 MHz, DMSO-$d_6$) of methyl 3-((3-acetylphenyl)carbamoyl) benzoate.

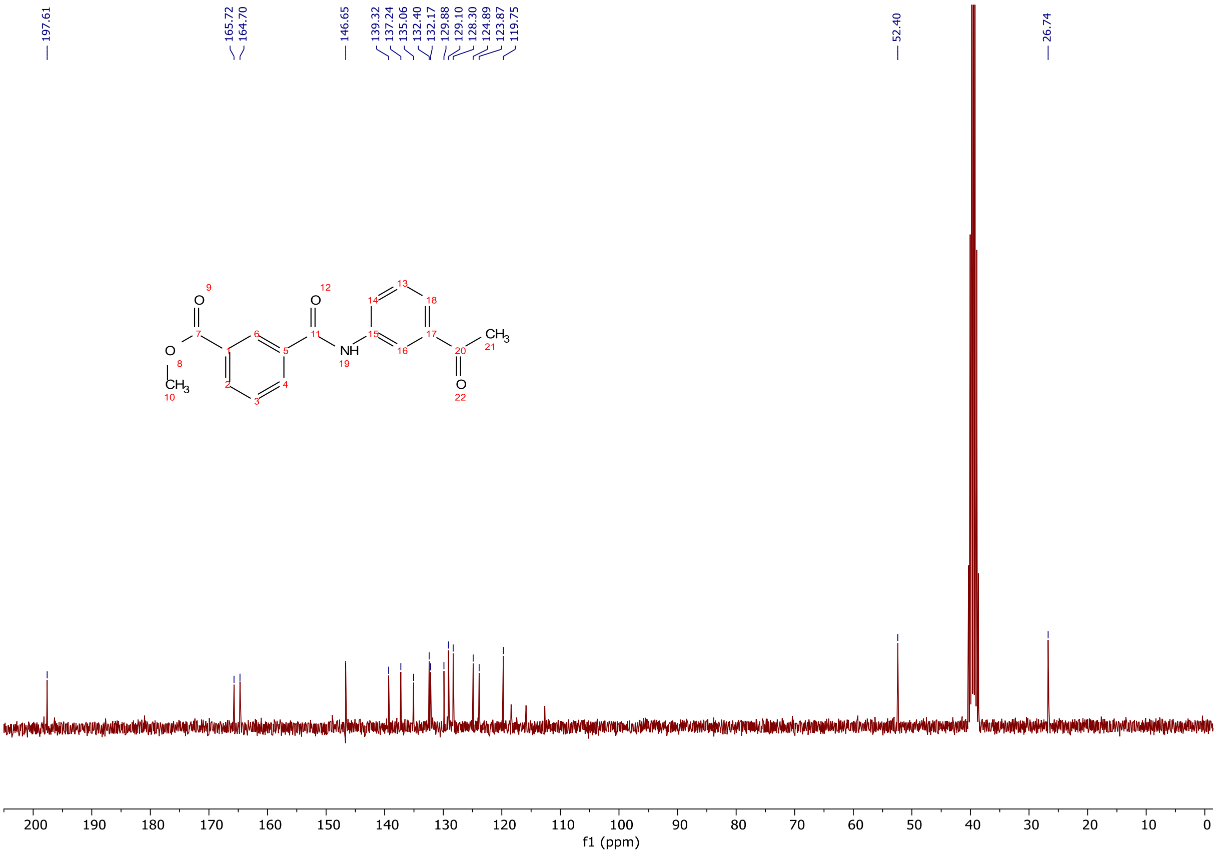

$^{13}C$ NMR spectra (75 MHz, DMSO-$d_6$) of methyl 3-((3-acetylphenyl)carbamoyl) benzoate.

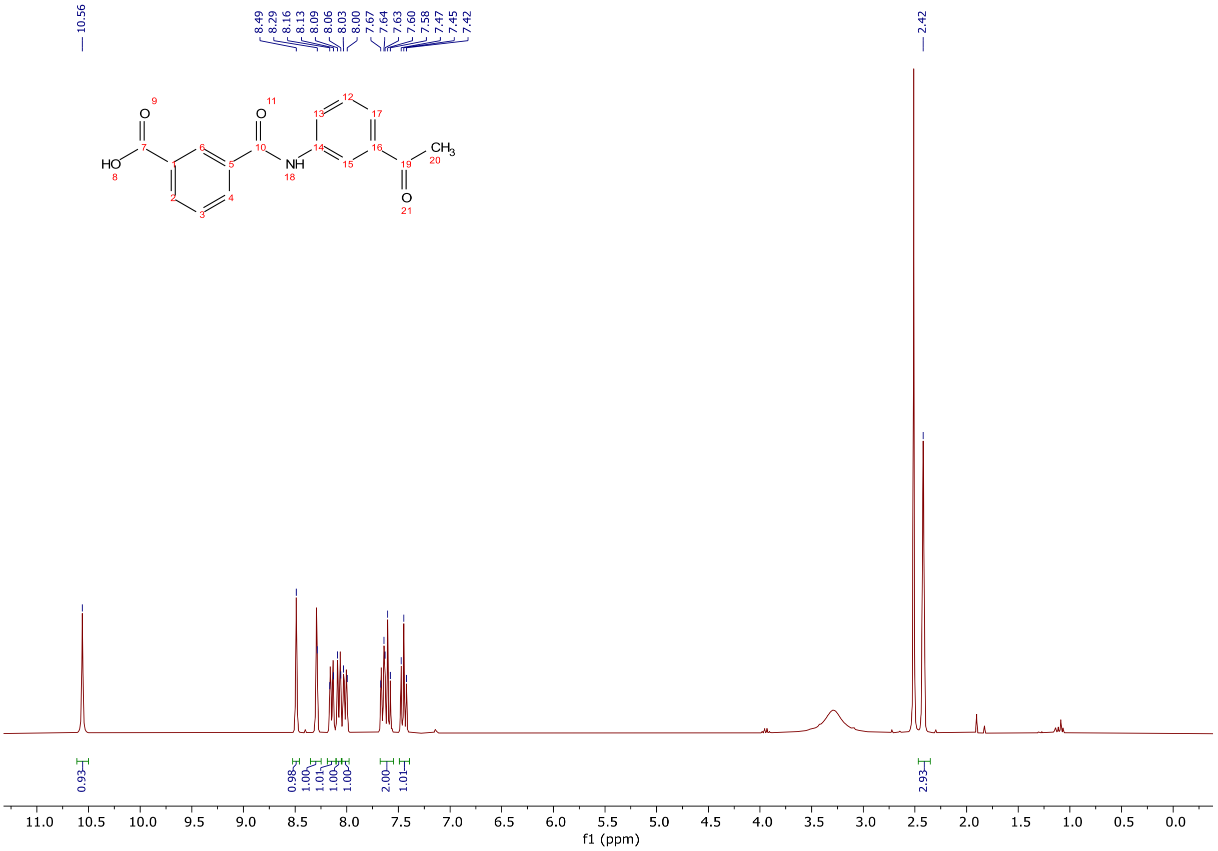

$^{1}H$ NMR spectra (300 MHz, DMSO-$d_6$) of 3-((3-acetylphenyl)carbamoyl)benzoic acid.

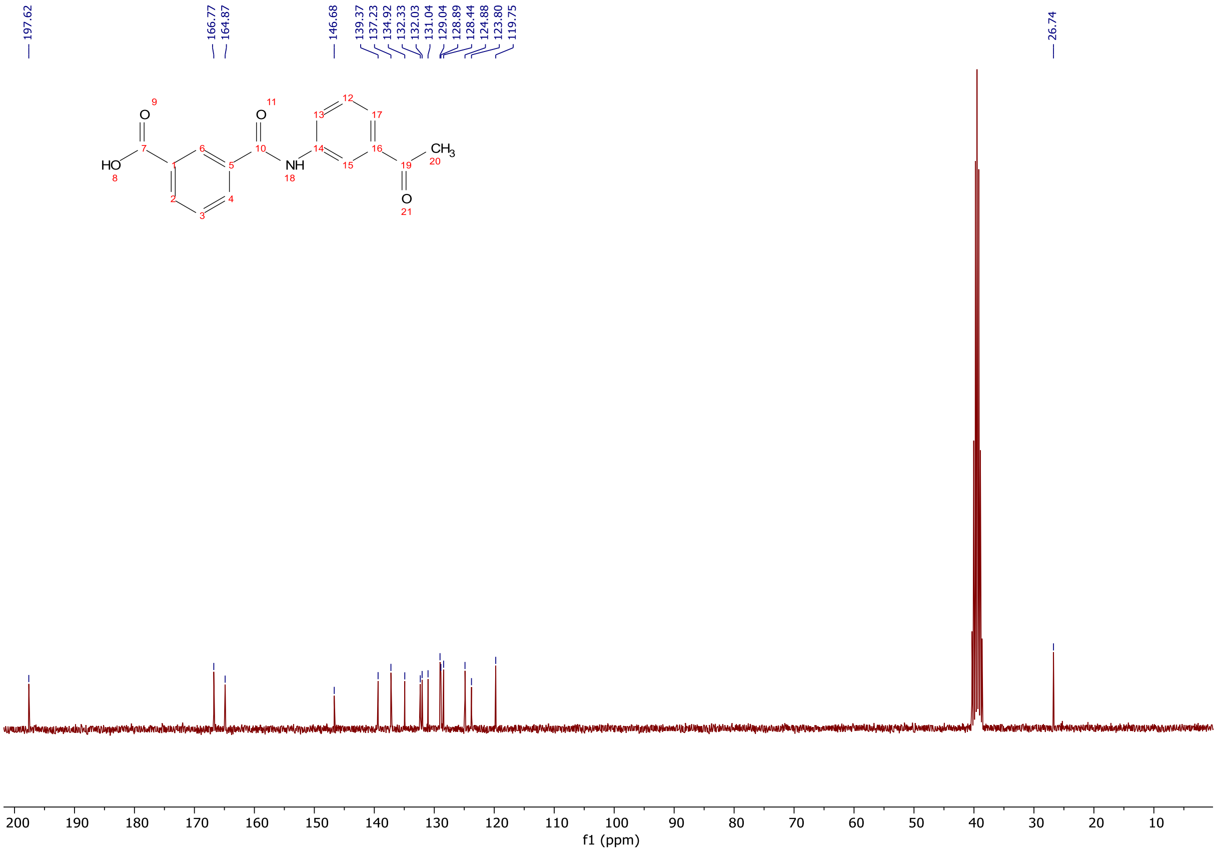


$^{13}C$ NMR spectra (75 MHz, DMSO-$d_6$) of 3-((3-acetylphenyl)carbamoyl)benzoic acid.

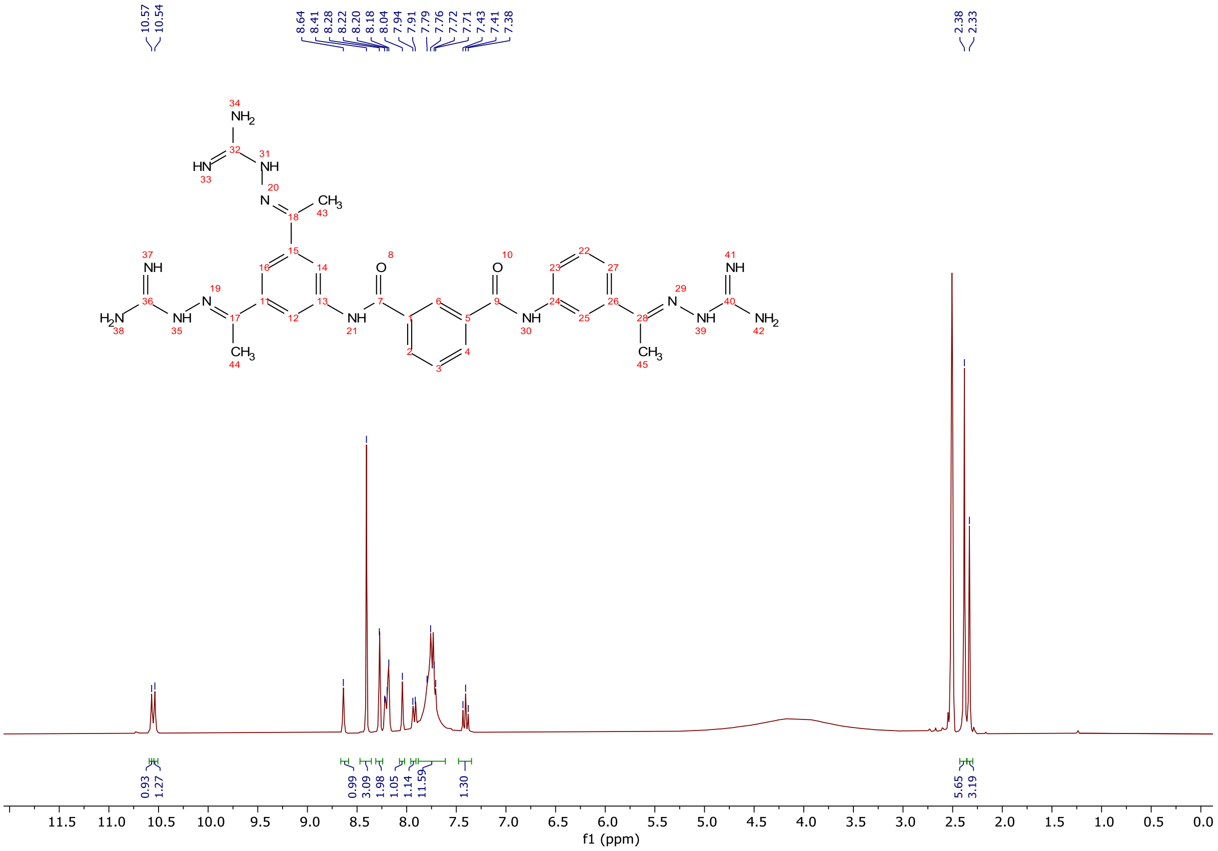


$^{1}H$ NMR spectra (300 MHz, DMSO-$d_6$) of **5**.

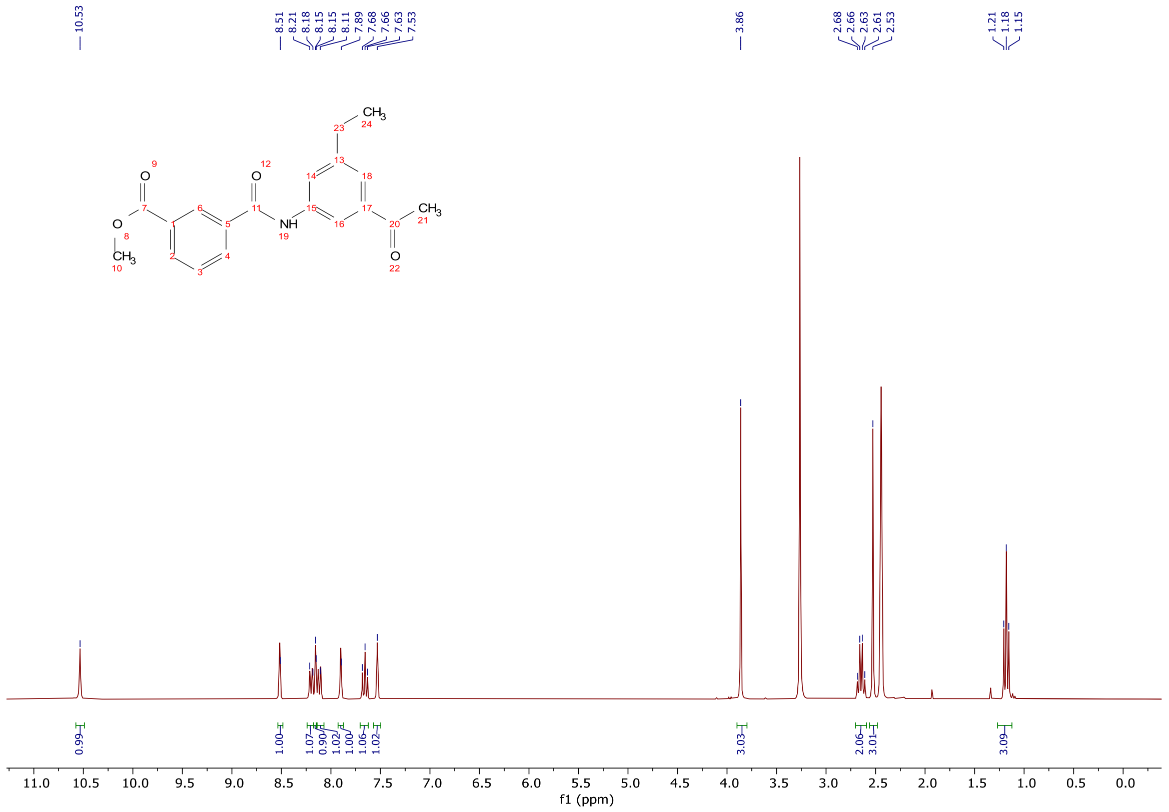

$^{1}H$ NMR spectra (300 MHz, DMSO-$d_6$) of methyl 3-((3-acetyl-5-ethylphenyl) carbamoyl)benzoate.

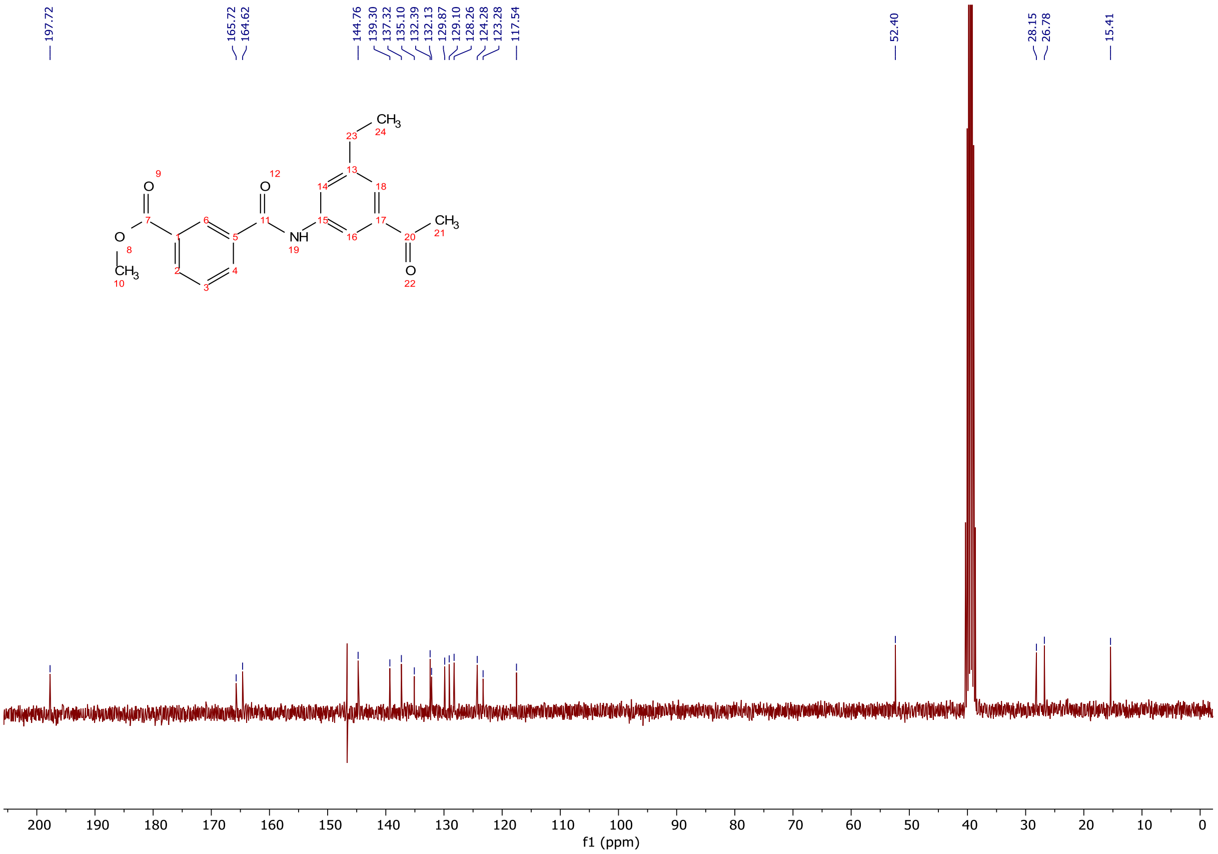

$^{13}C$ NMR spectra (75 MHz, DMSO-$d_6$) of methyl 3-((3-acetyl-5-ethylphenyl) carbamoyl)benzoate.

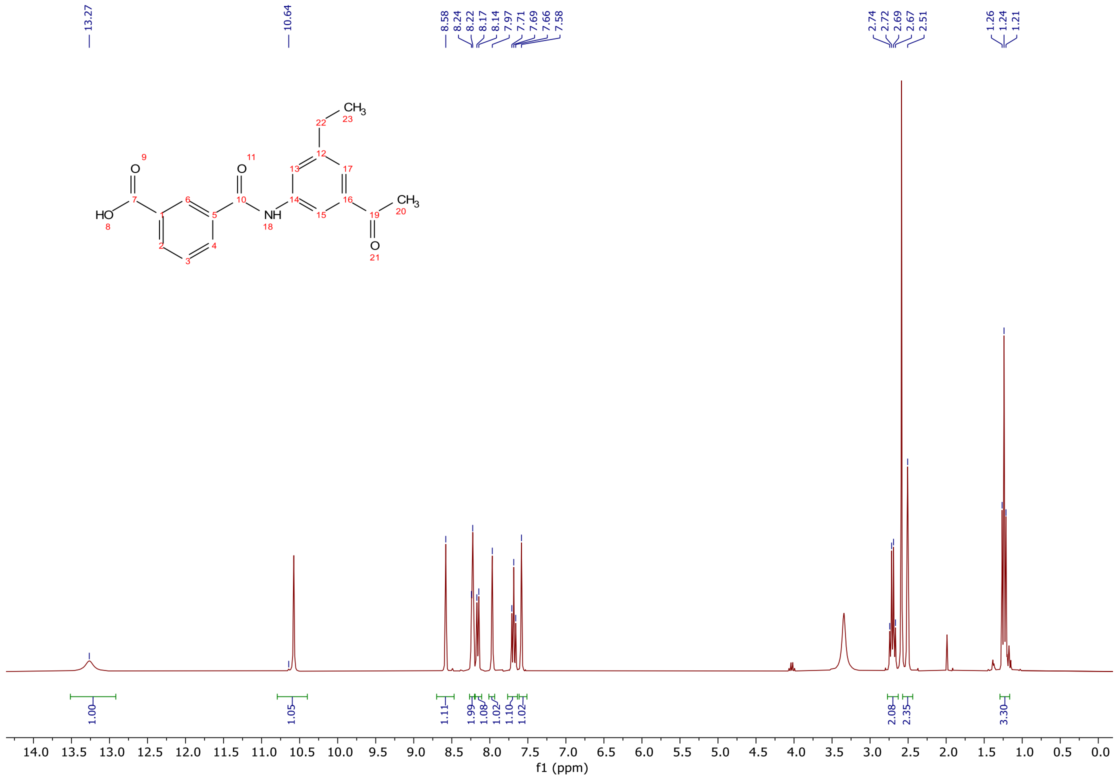

$^{1}H$ NMR spectra (300 MHz, DMSO-$d_6$) of 3-((3-acetyl-5-ethylphenyl)carbamoyl) benzoic acid.

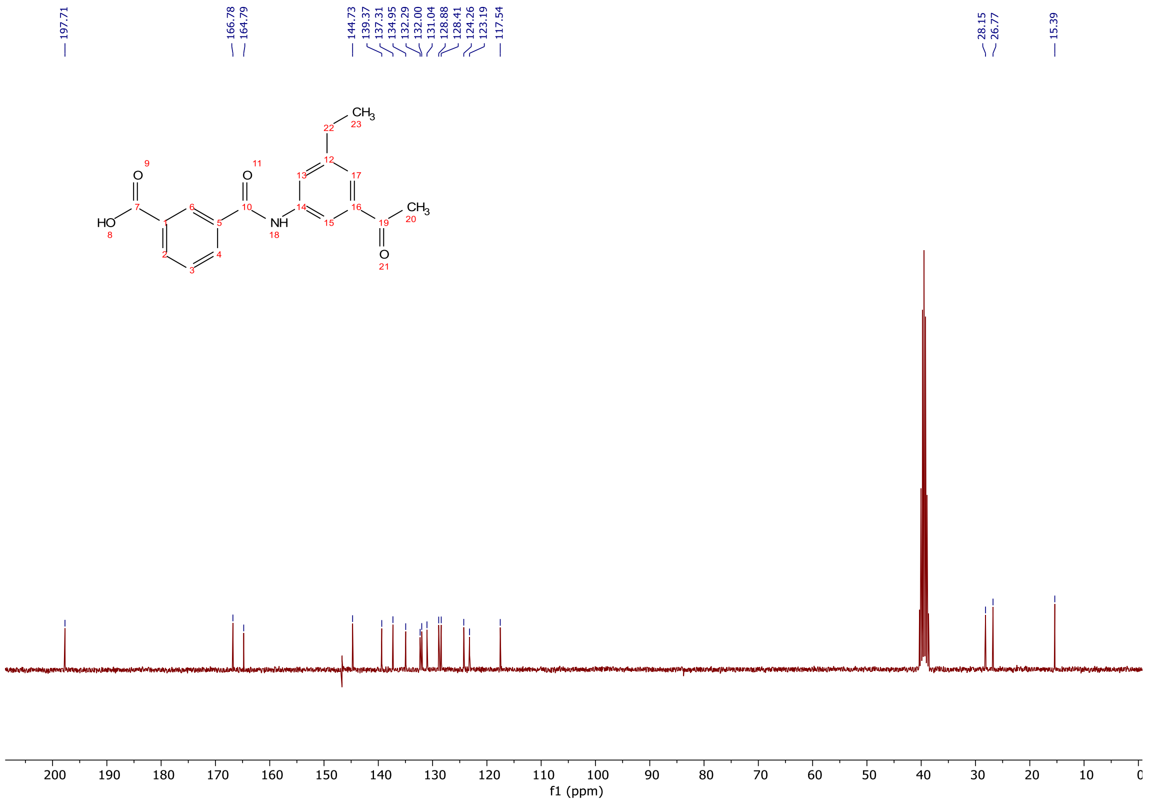

$^{13}C$ NMR spectra (75 MHz, DMSO-$d_6$) of 3-((3-acetyl-5-ethylphenyl)carbamoyl) benzoic acid.

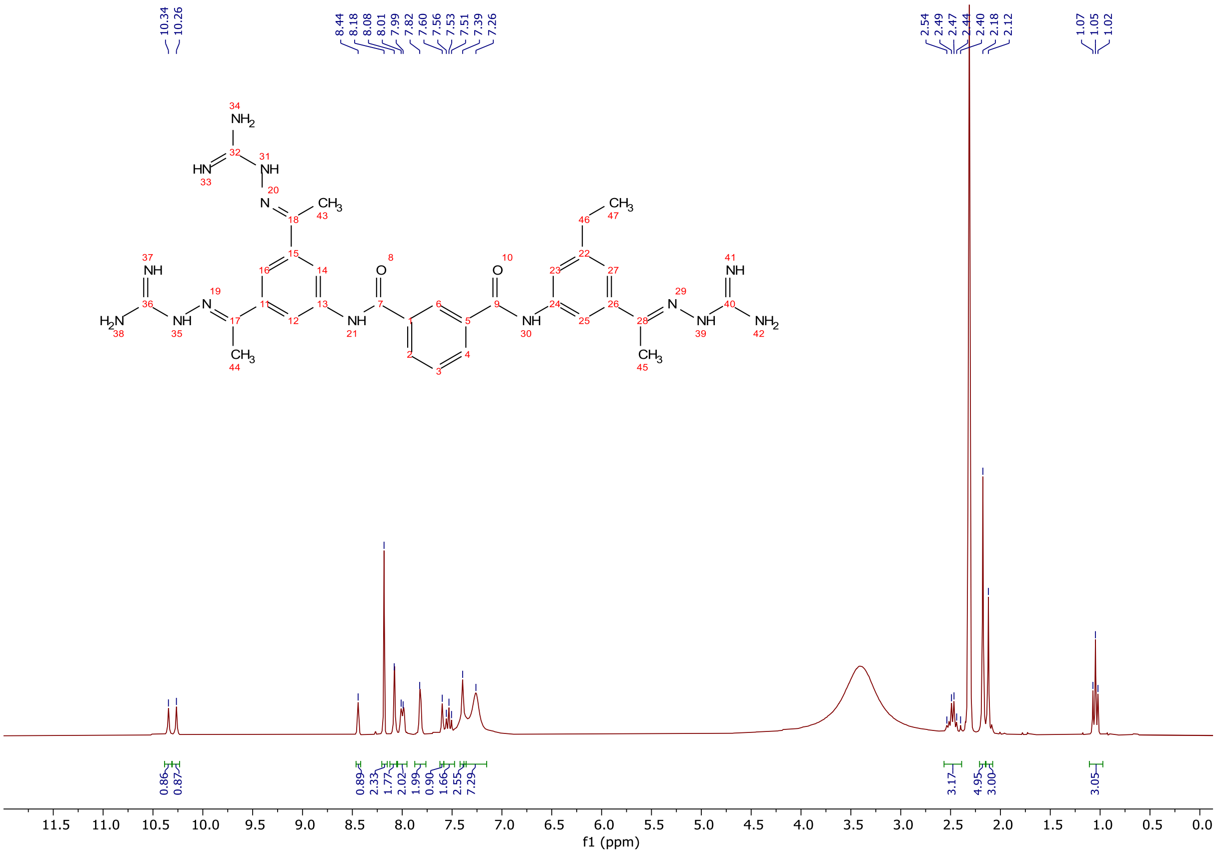

[1]H NMR spectra (300 MHz, DMSO-d$_6$) of **6**.

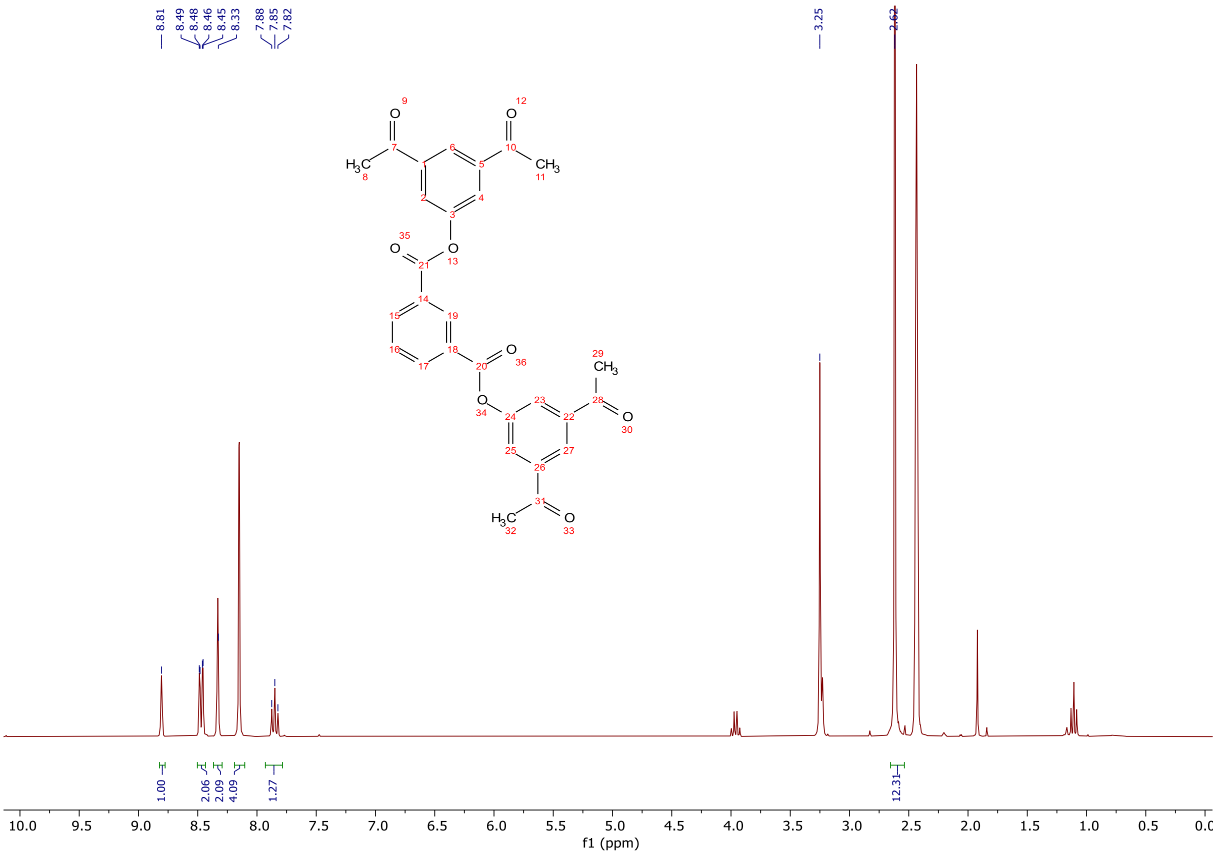

[1]H NMR spectra (300 MHz, DMSO-d$_6$) of **7-1**.

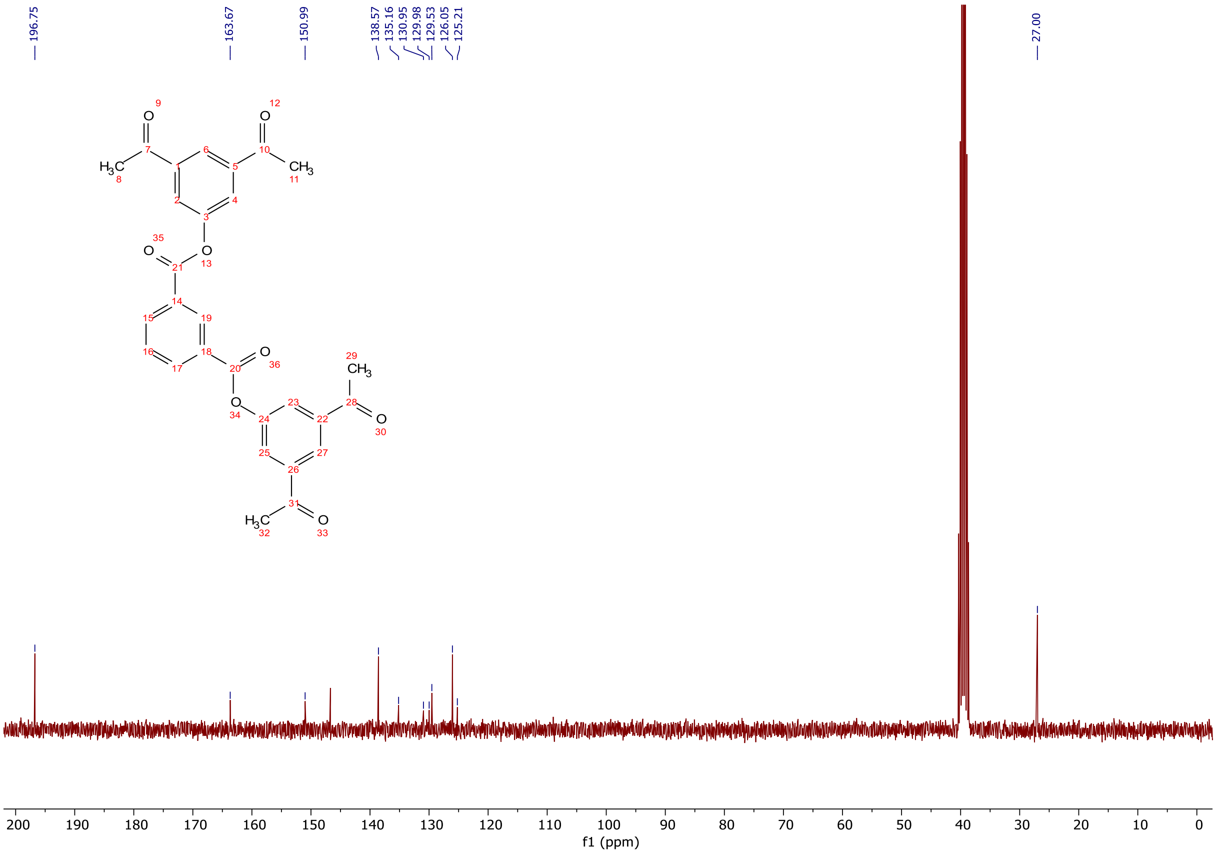

$^{13}C$ NMR spectra (75 MHz, DMSO-$d_6$) of **7-1**.

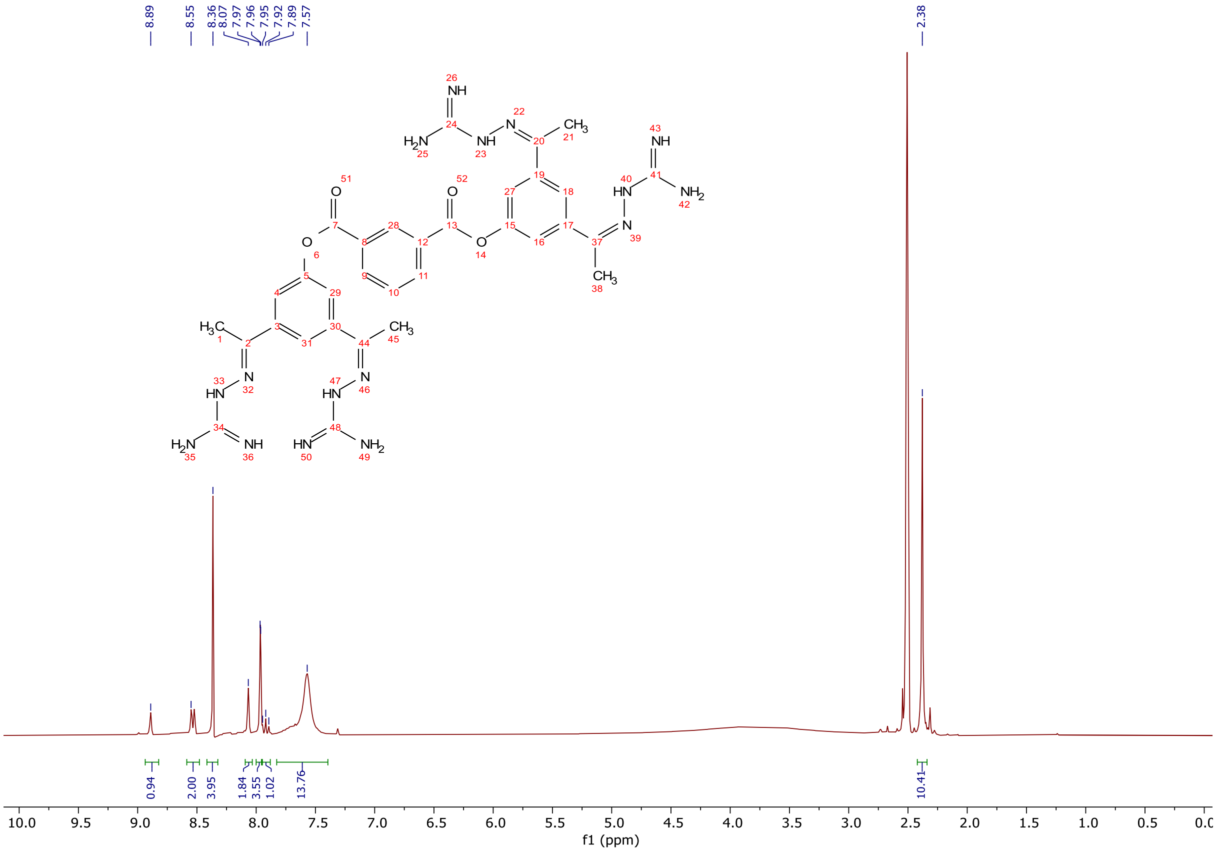

$^{1}H$ NMR spectra (300 MHz, DMSO-$d_6$) of **7**.

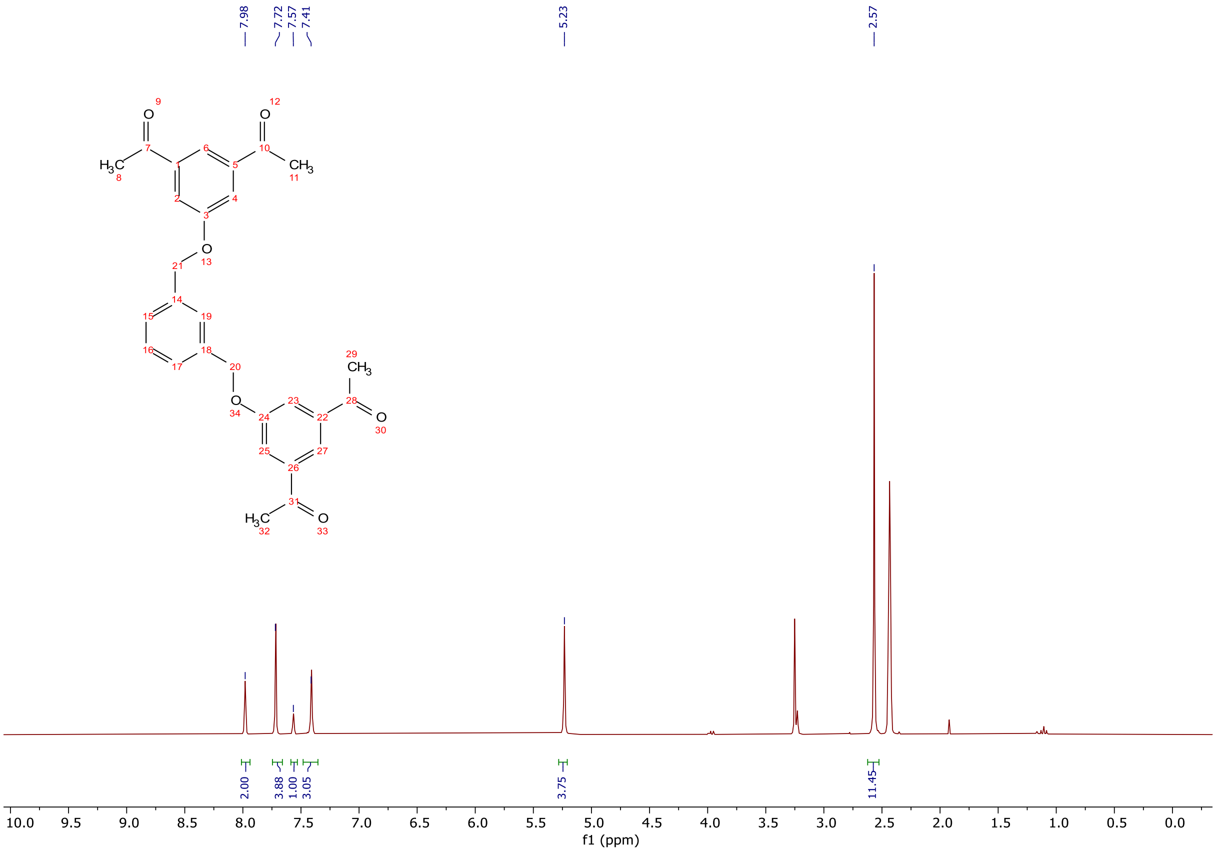

$^{1}$H NMR spectra (300 MHz, DMSO-$d_6$) of **8-1**.

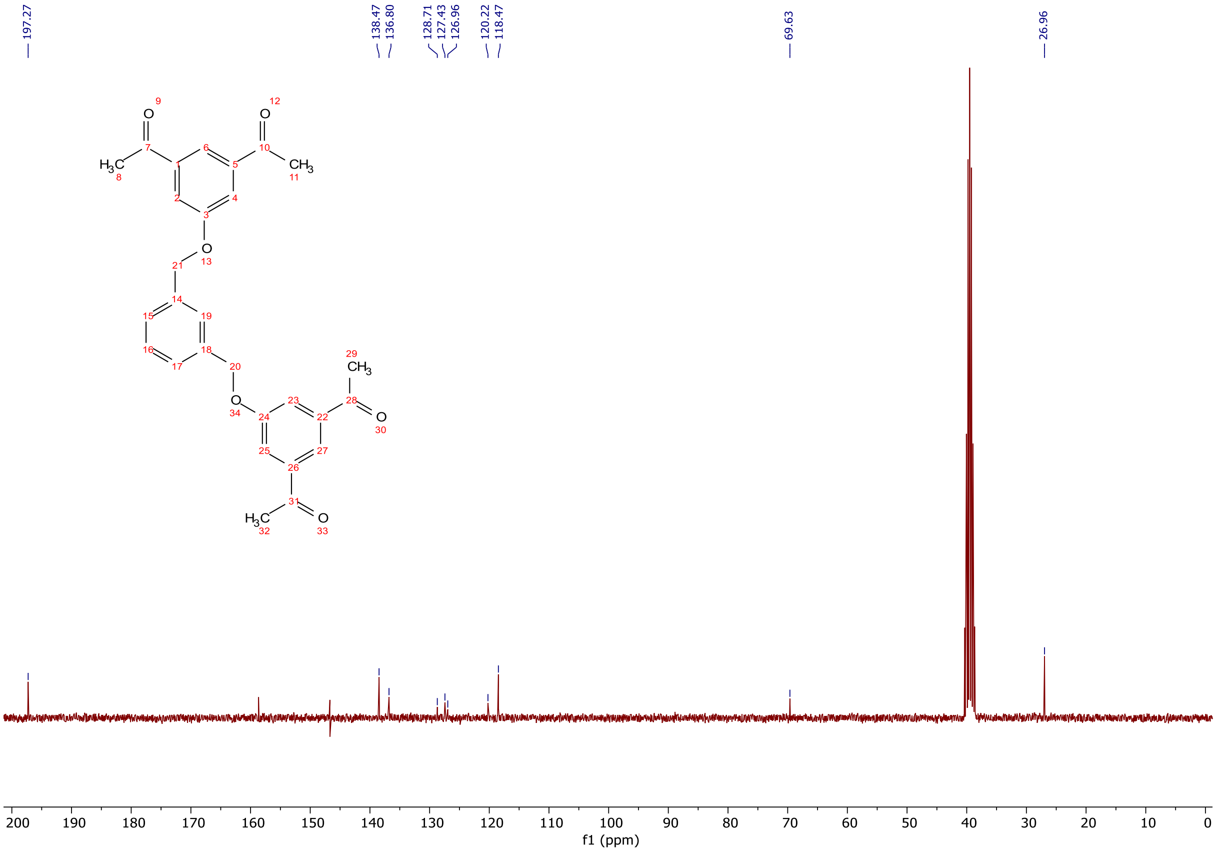

$^{13}$C NMR spectra (75 MHz, DMSO-$d_6$) of **8-1**.

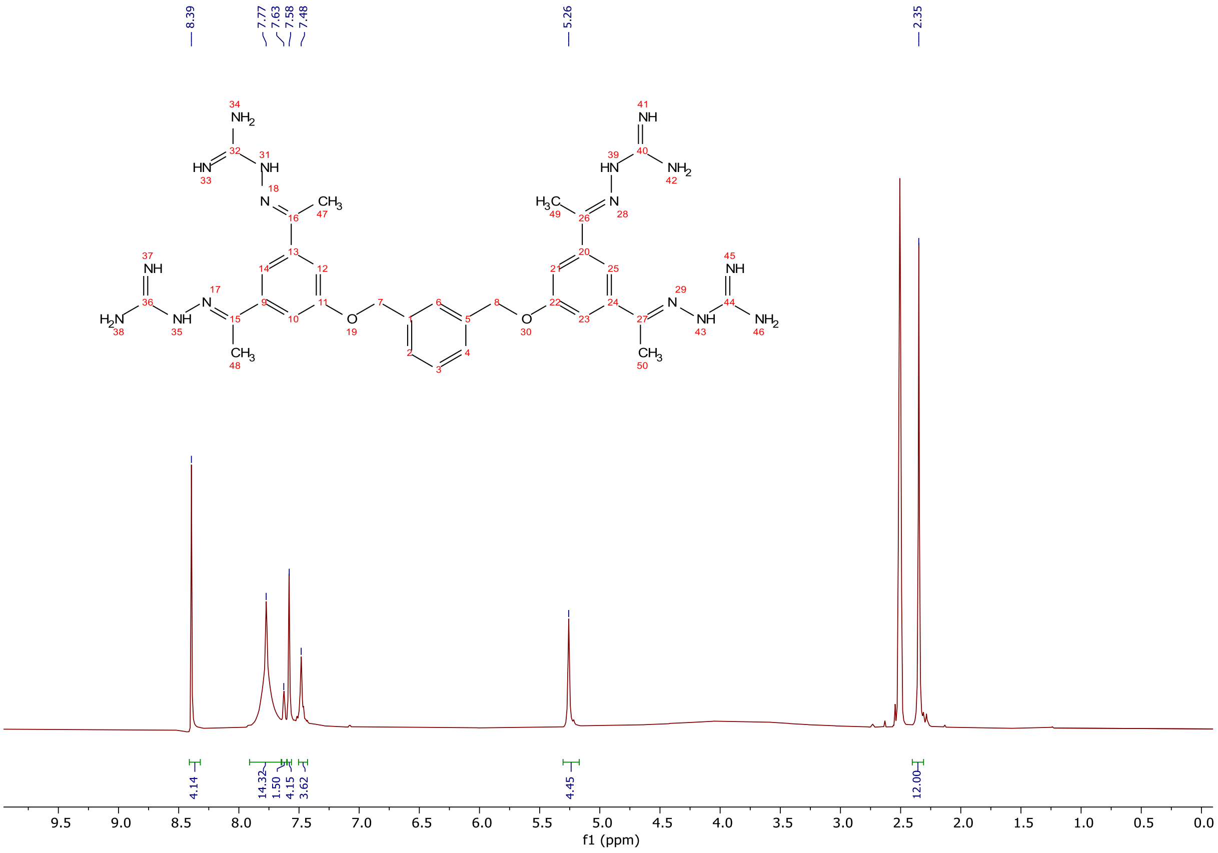


$^1$H NMR spectra (300 MHz, DMSO-d$_6$) of **8**.

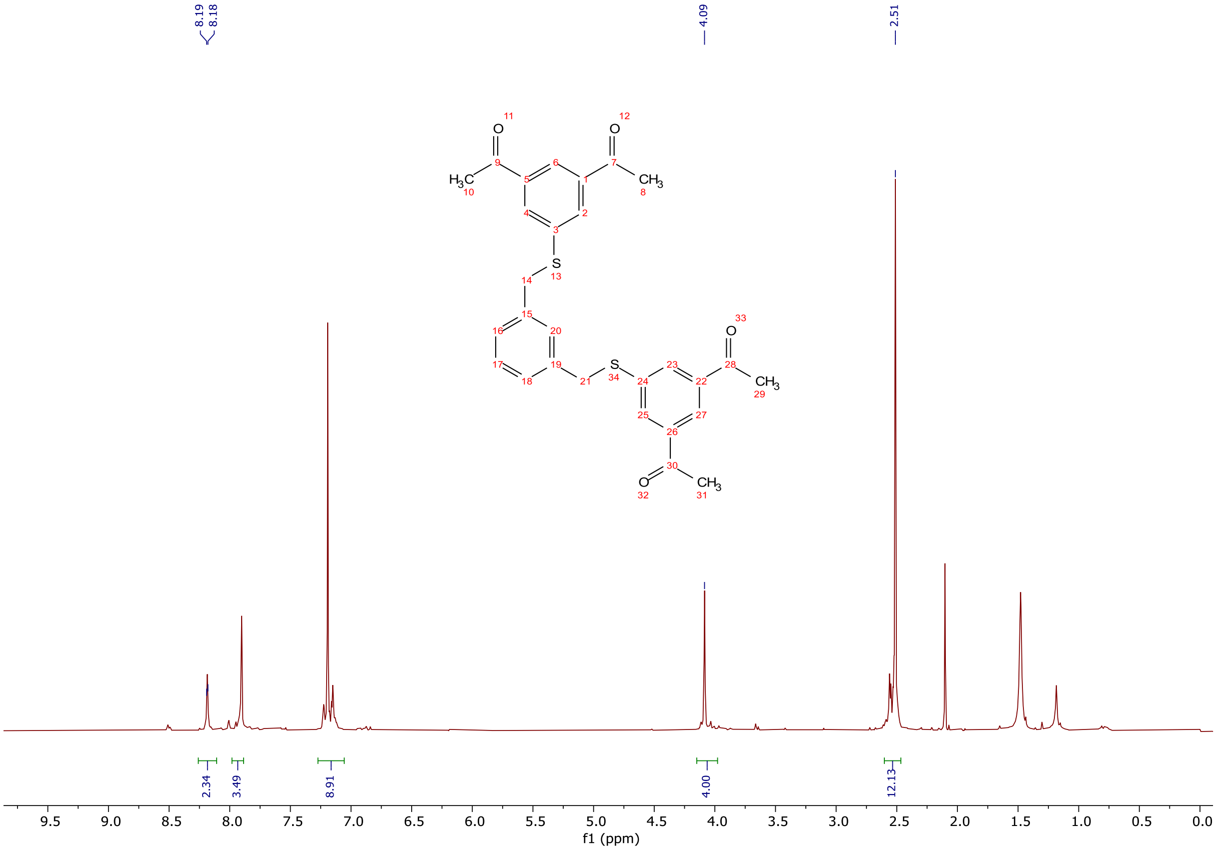


$^1$H NMR spectra (300 MHz, CDCl$_3$) of **9**.

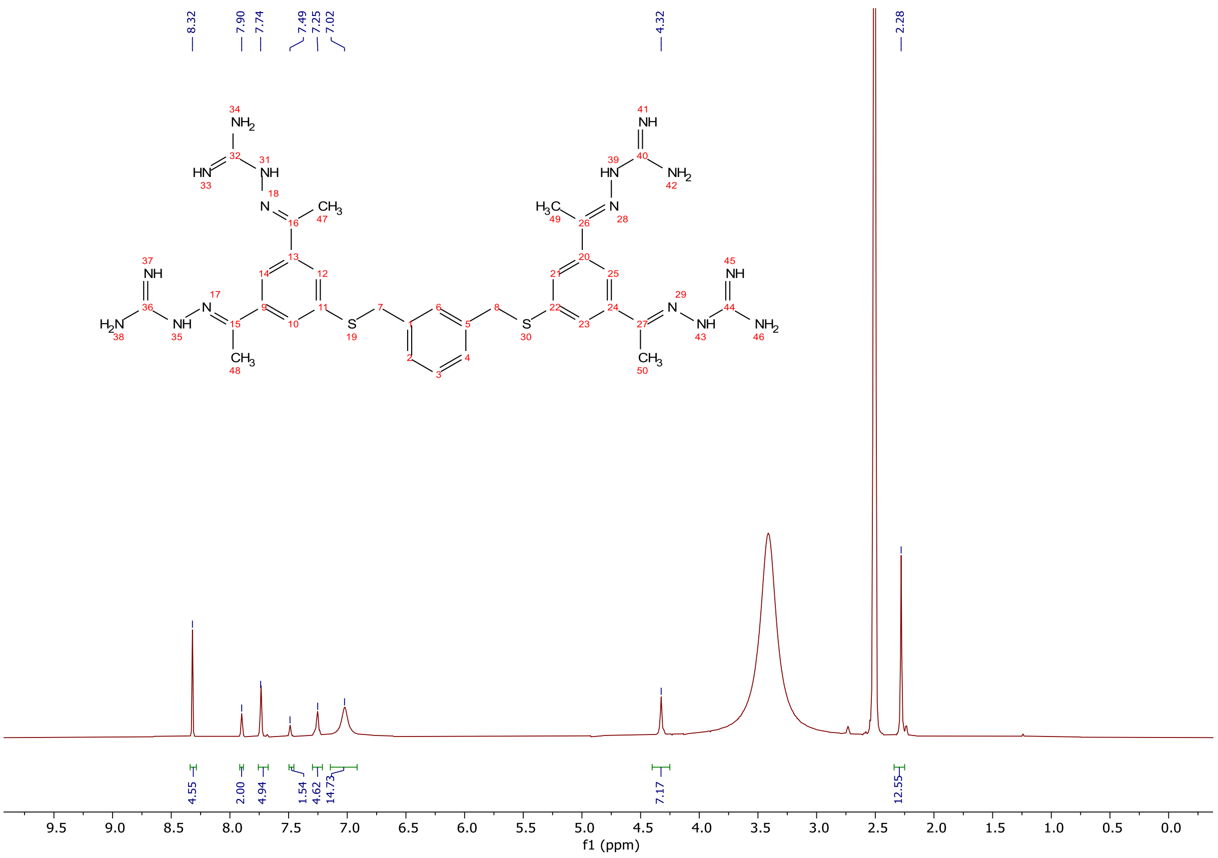

$^1$H NMR spectra (300 MHz, DMSO-d$_6$) of **9**.

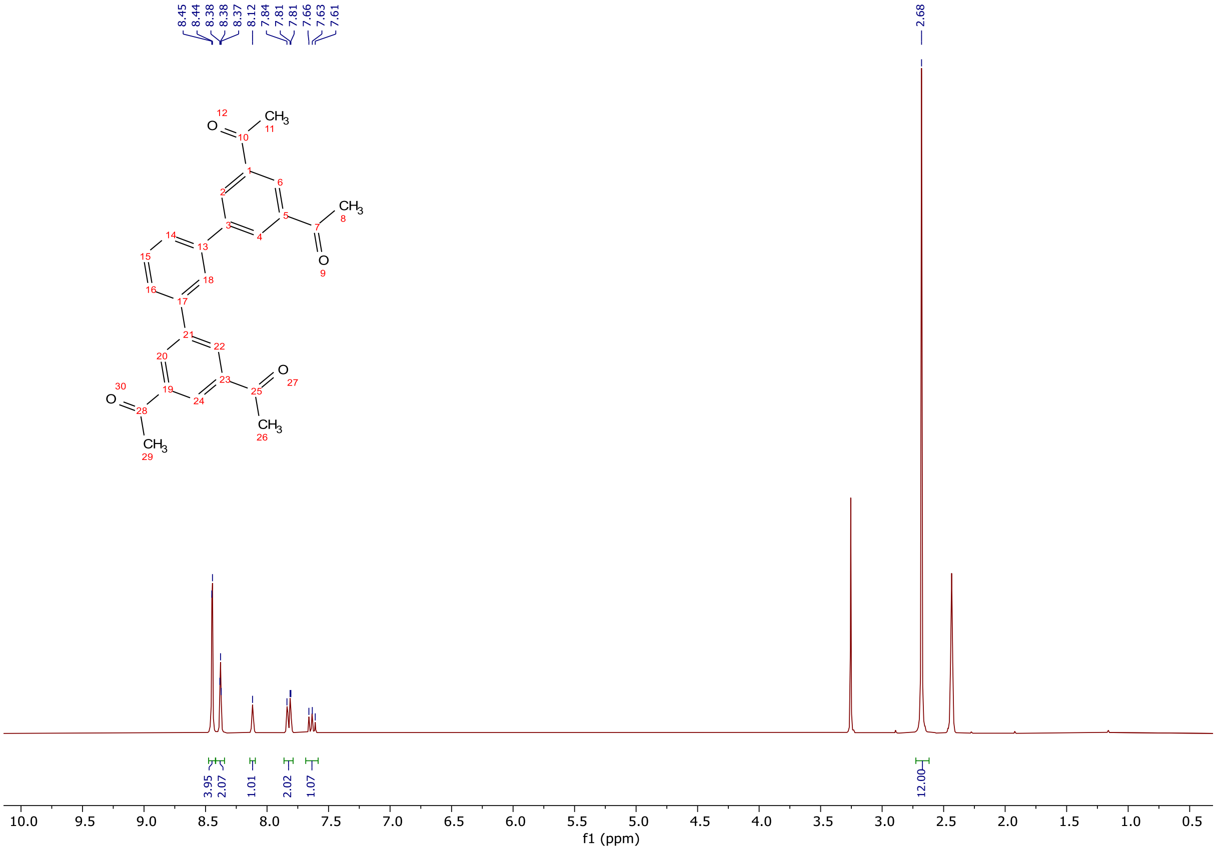

$^1$H NMR spectra (300 MHz, DMSO-d$_6$) of **10-1**.

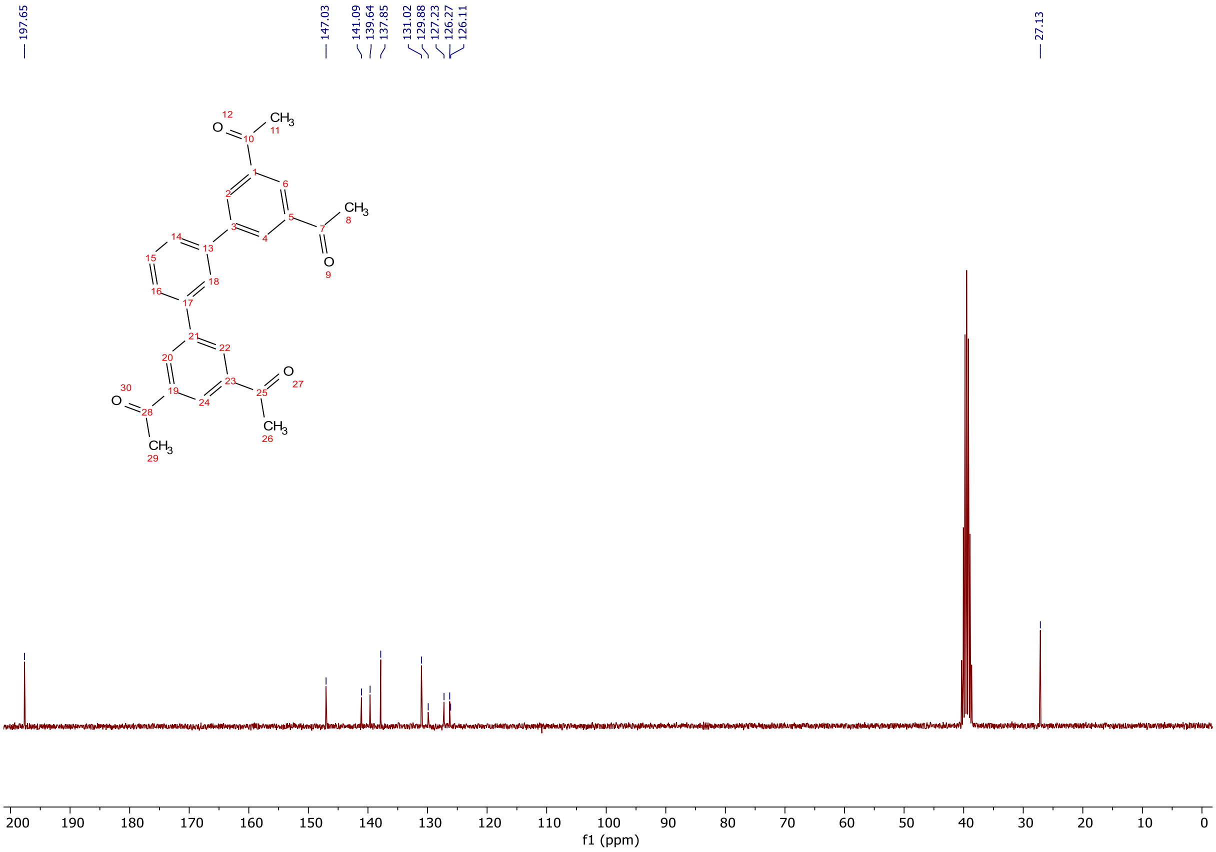

$^{13}C$ NMR spectra (75 MHz, DMSO-$d_6$) of **10-1**.

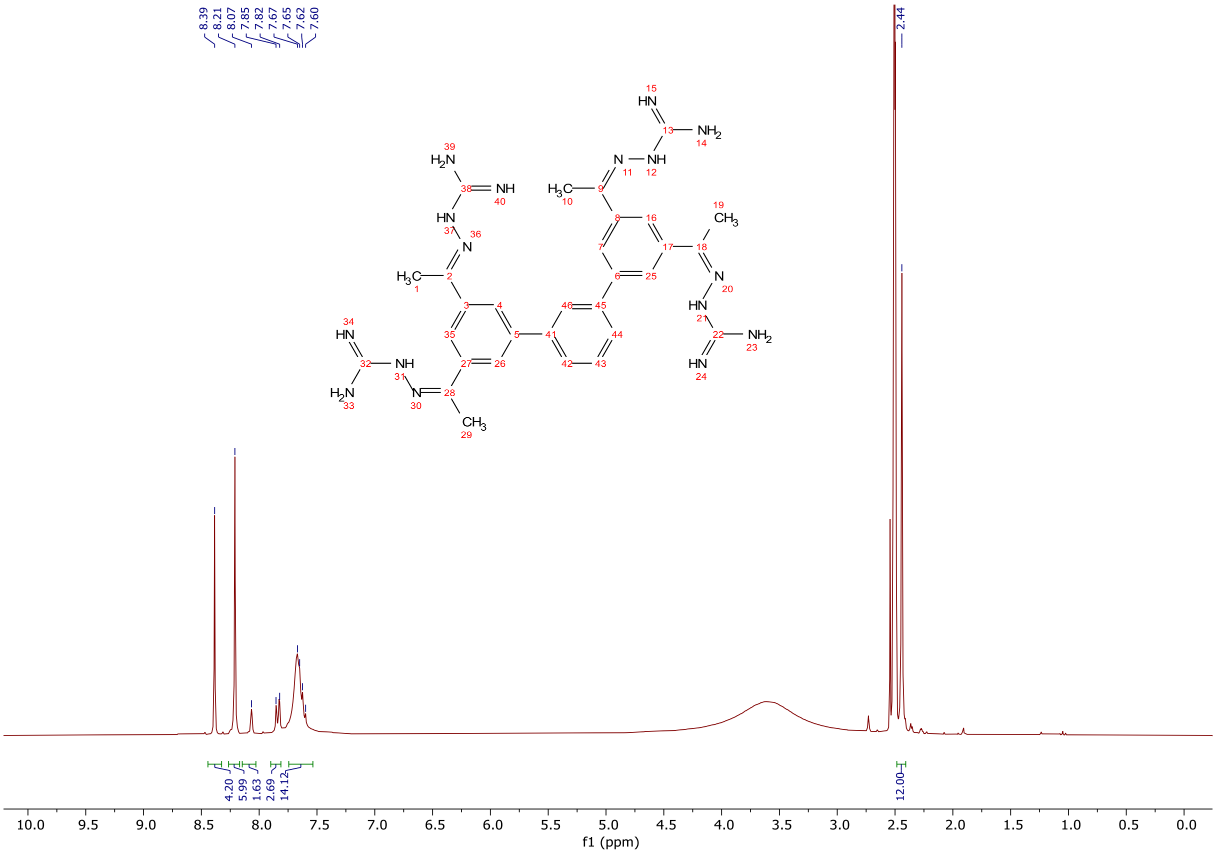

$^{1}H$ NMR spectra (300 MHz, DMSO-$d_6$) of **10**.

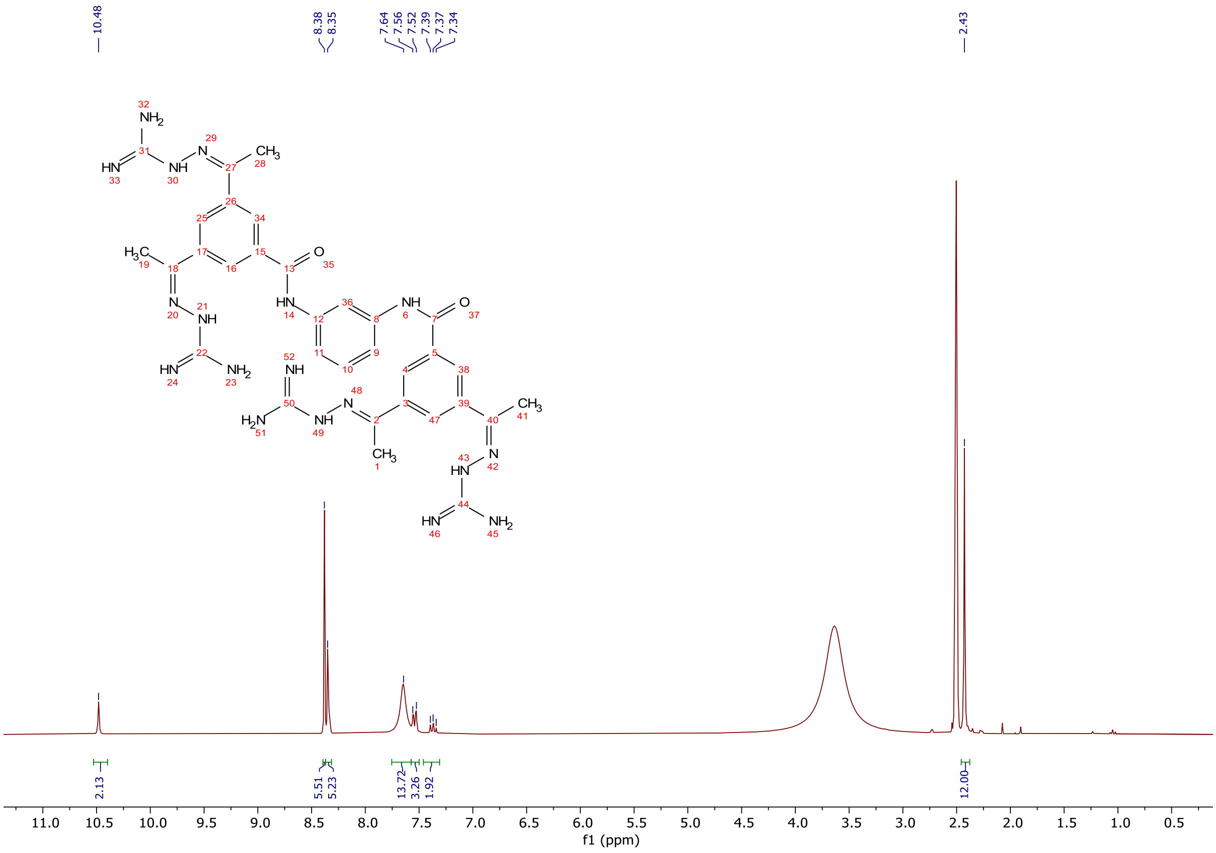

$^1$H NMR spectra (300 MHz, DMSO-$d_6$) of **11**.

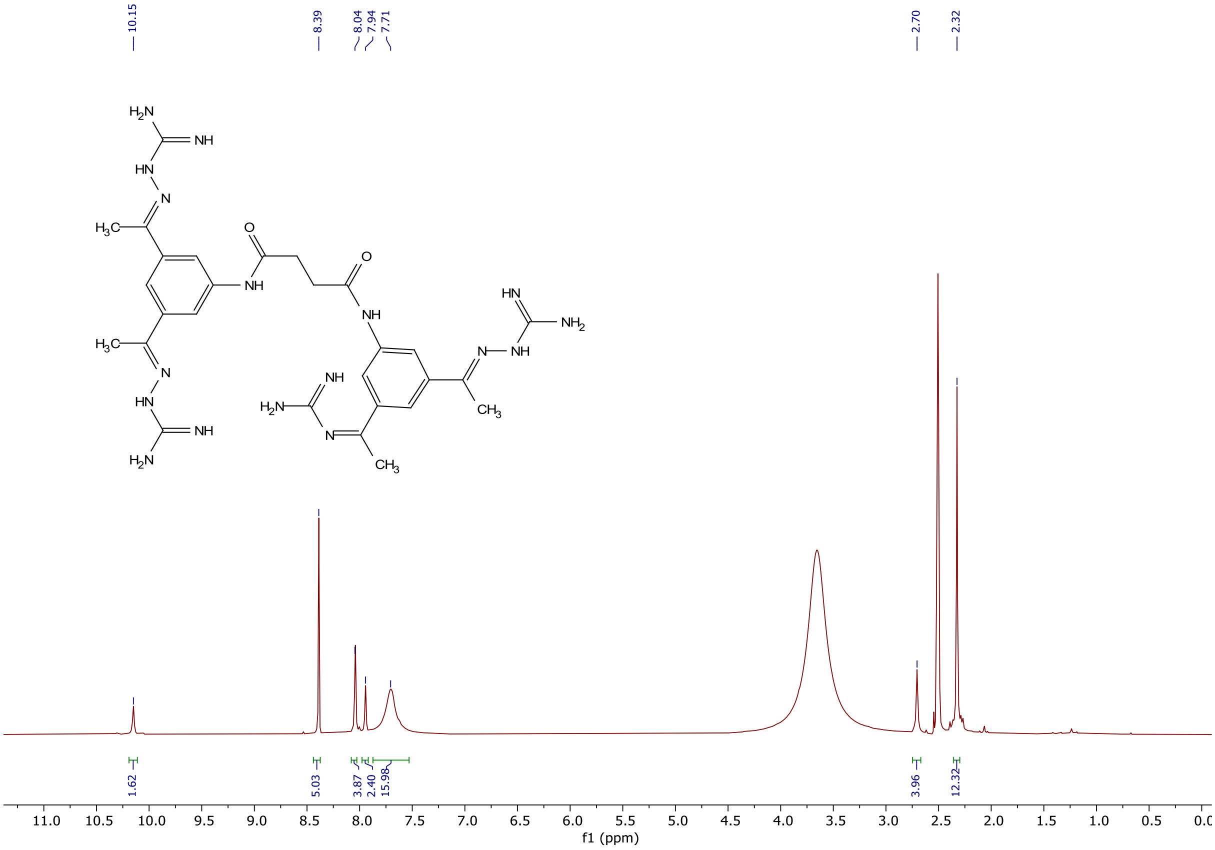

**Figure S**. $^1$H NMR spectra (300 MHz, DMSO-$d_6$) of **12**.

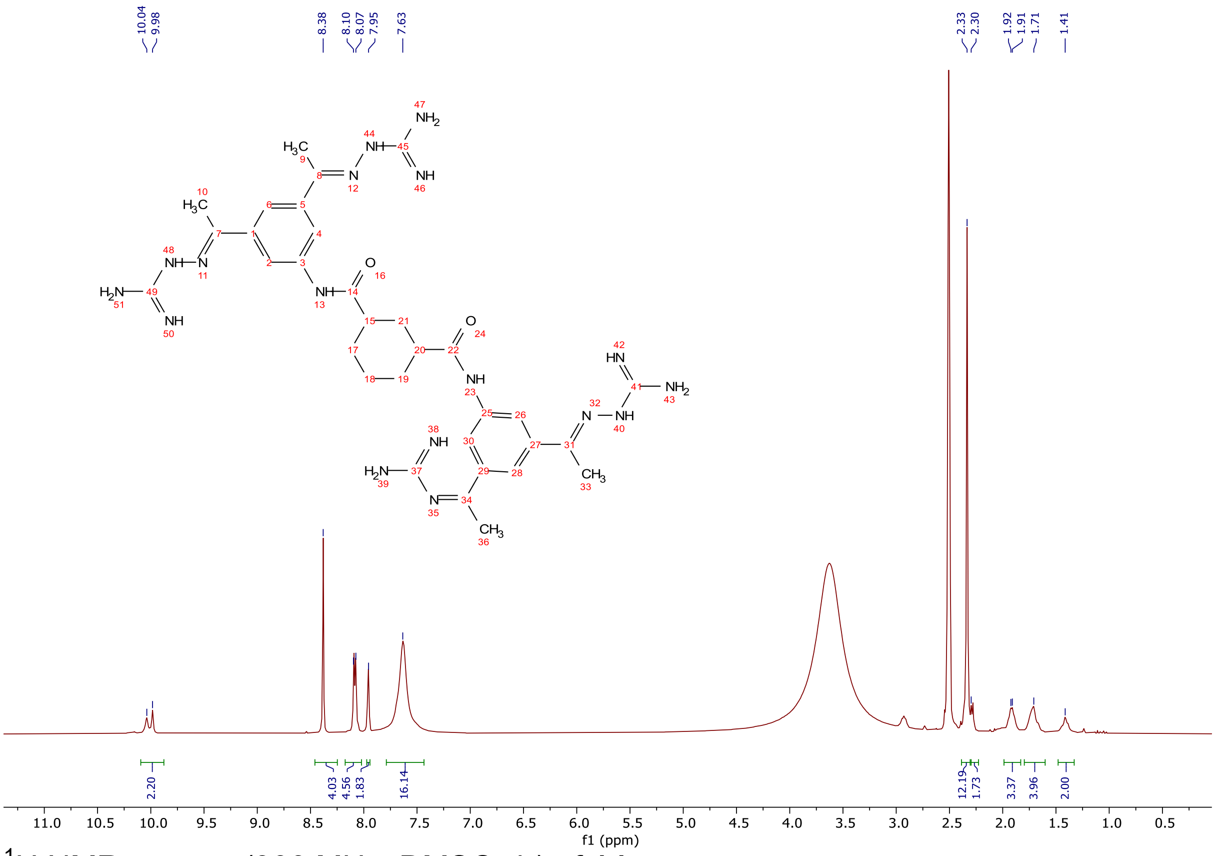


$^1$H NMR spectra (300 MHz, DMSO-$d_6$) of **14**.

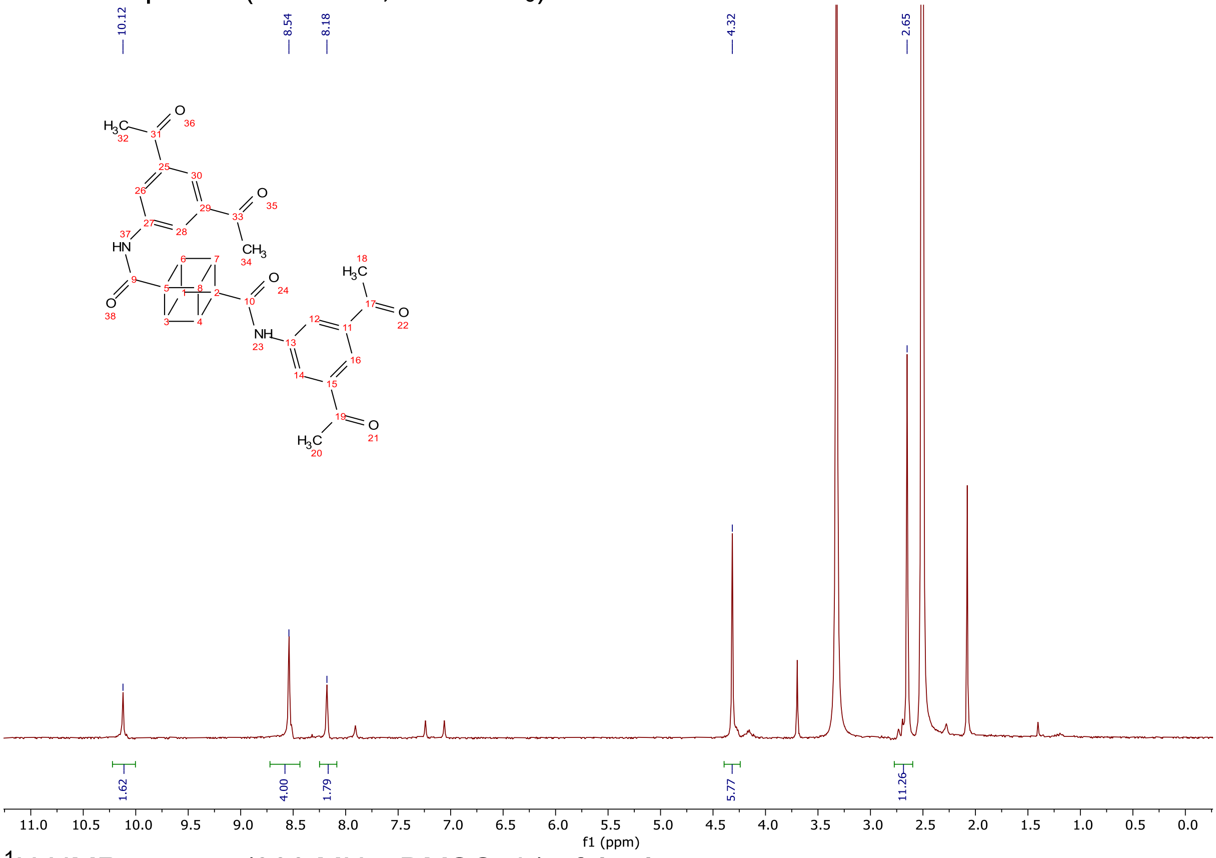


$^1$H NMR spectra (300 MHz, DMSO-$d_6$) of **15-1**.

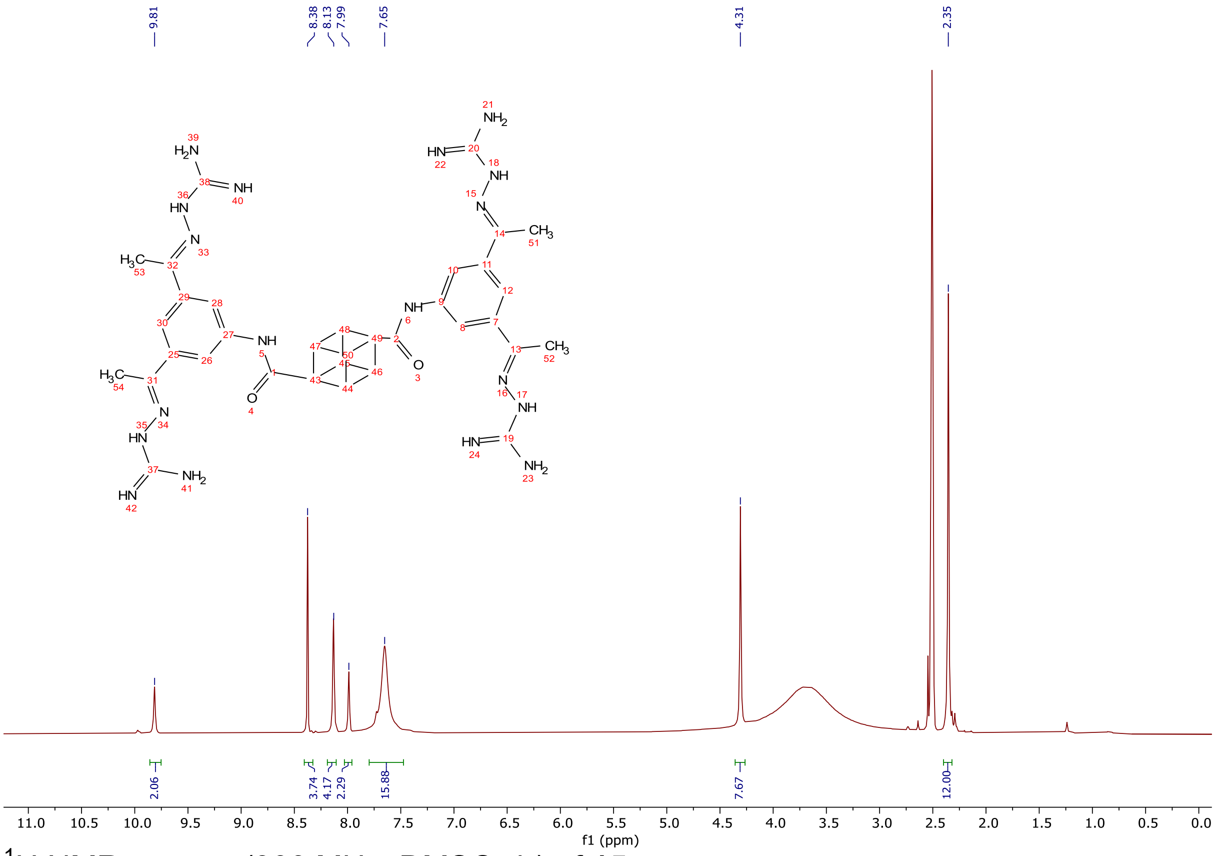


$^1$H NMR spectra (300 MHz, DMSO-d$_6$) of **15**.

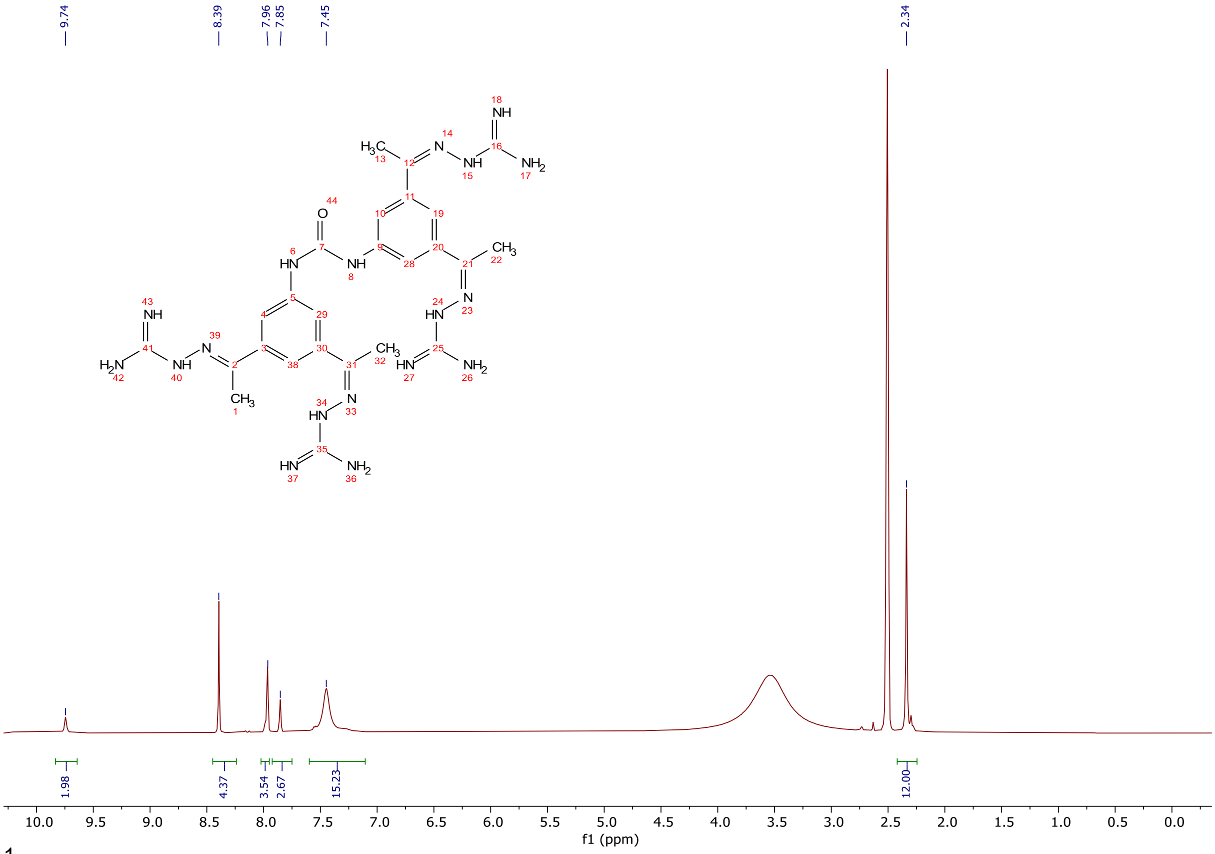


$^1$H NMR spectra (300 MHz, DMSO-d$_6$) of **16**.

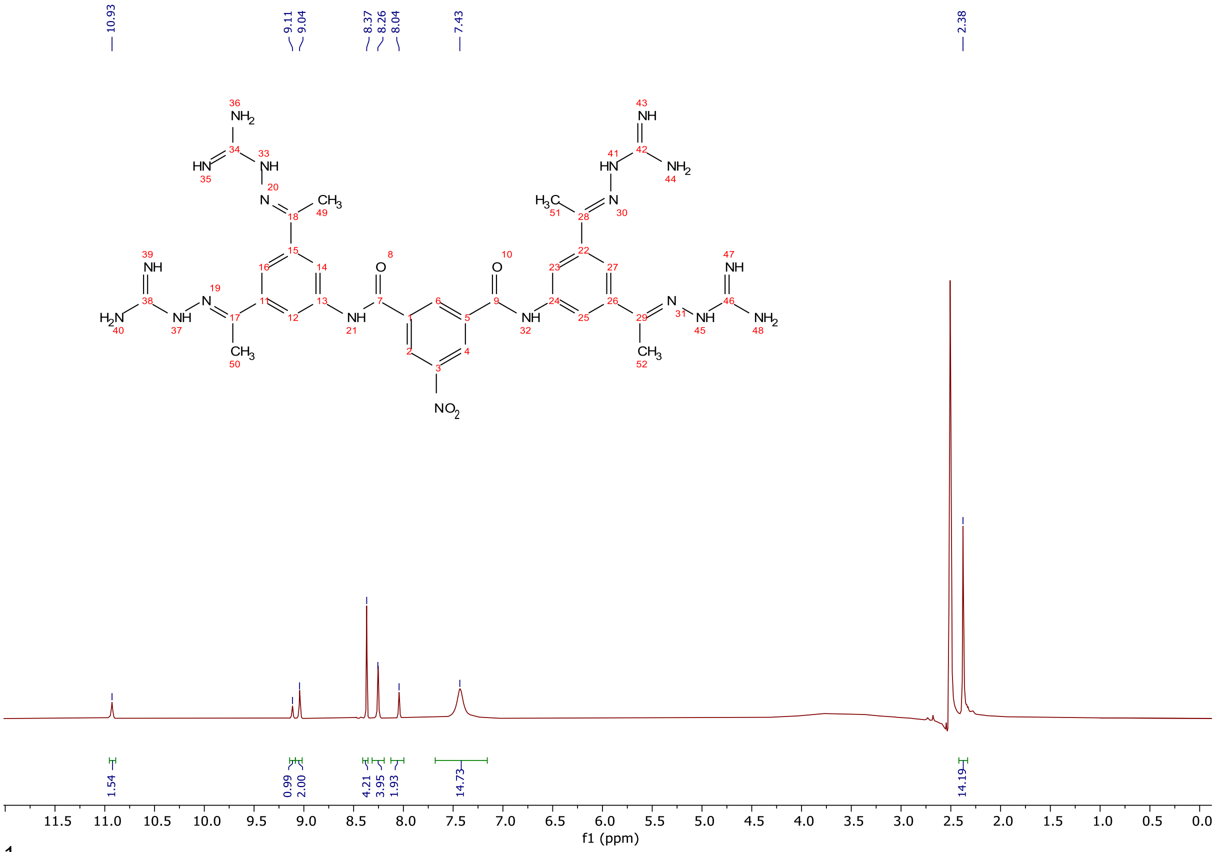

$^{1}H$ NMR spectra (300 MHz, DMSO-$d_6$) of **17** (TCB-494).

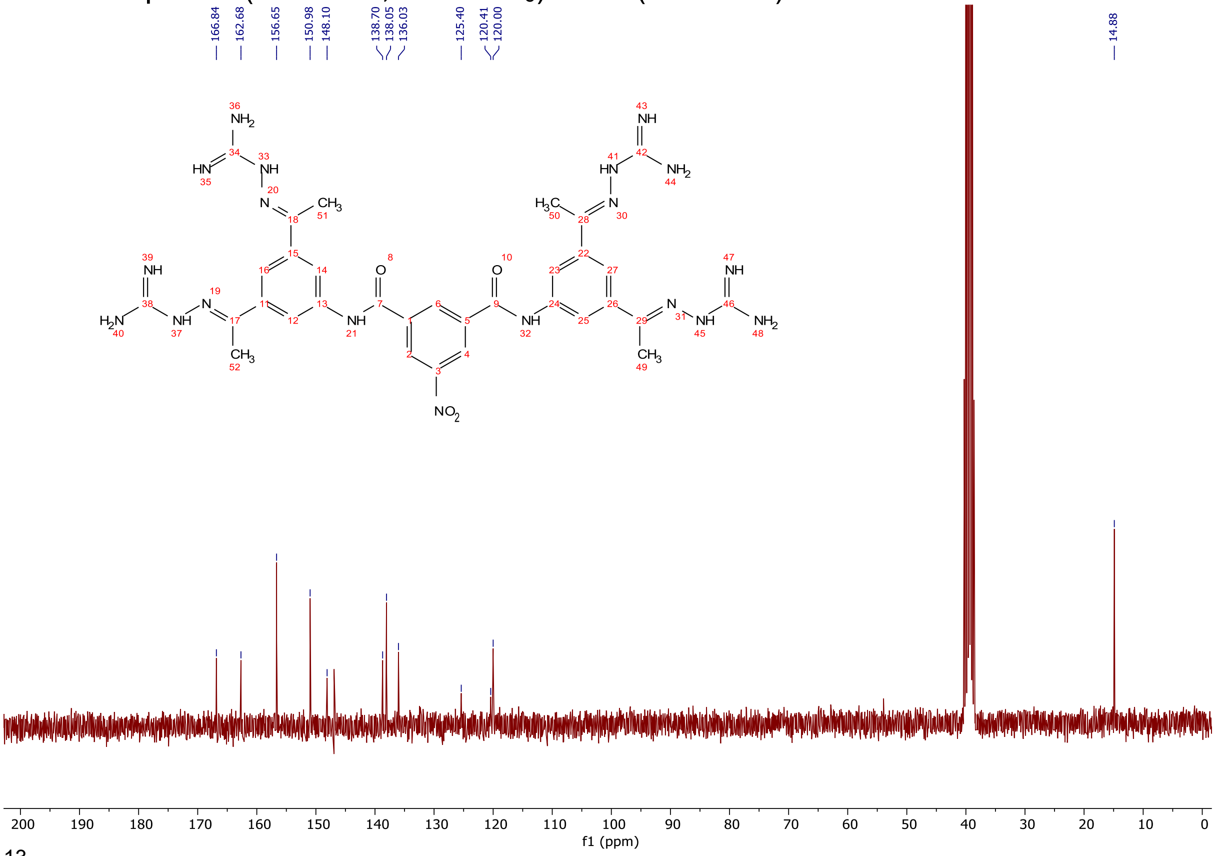

$^{13}C$ NMR spectra (75 MHz, DMSO-$d_6$) of **17** (TCB-494).

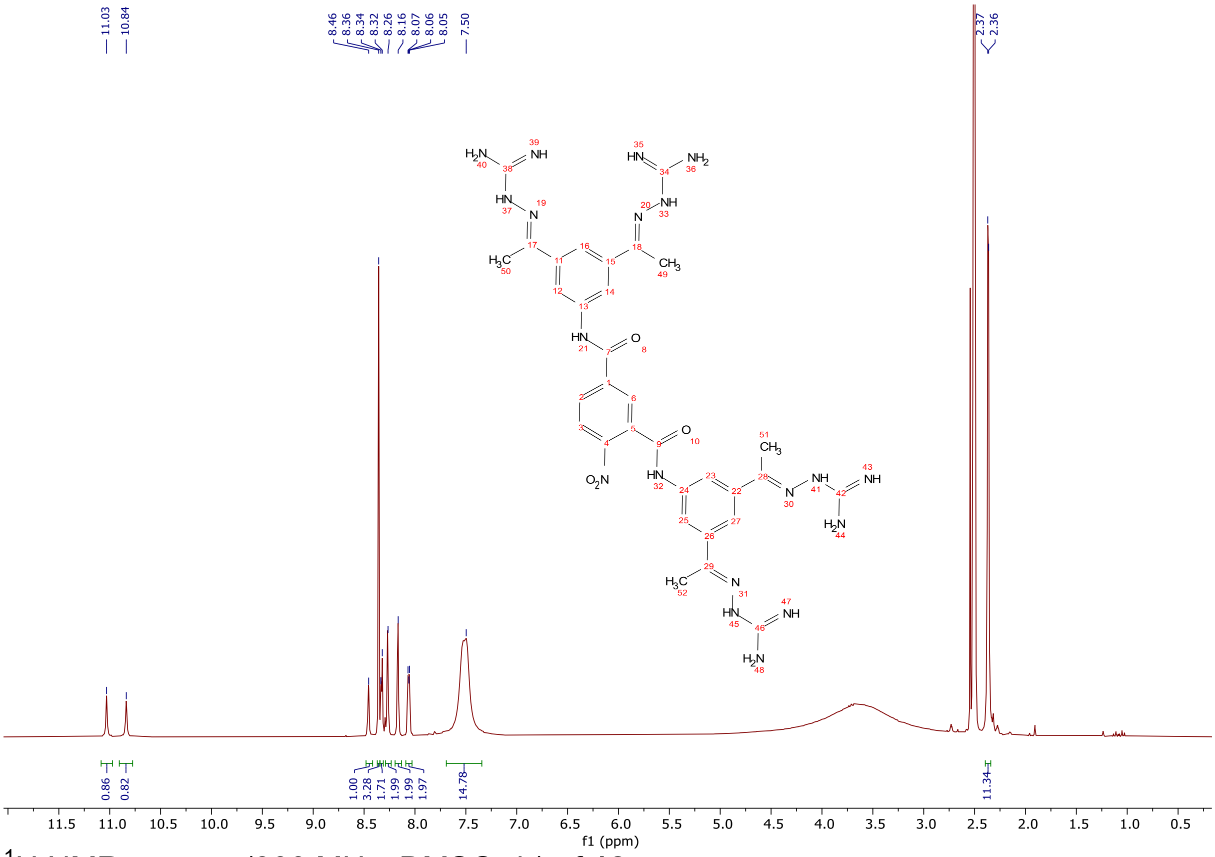

[1]H NMR spectra (300 MHz, DMSO-$d_6$) of **18**.

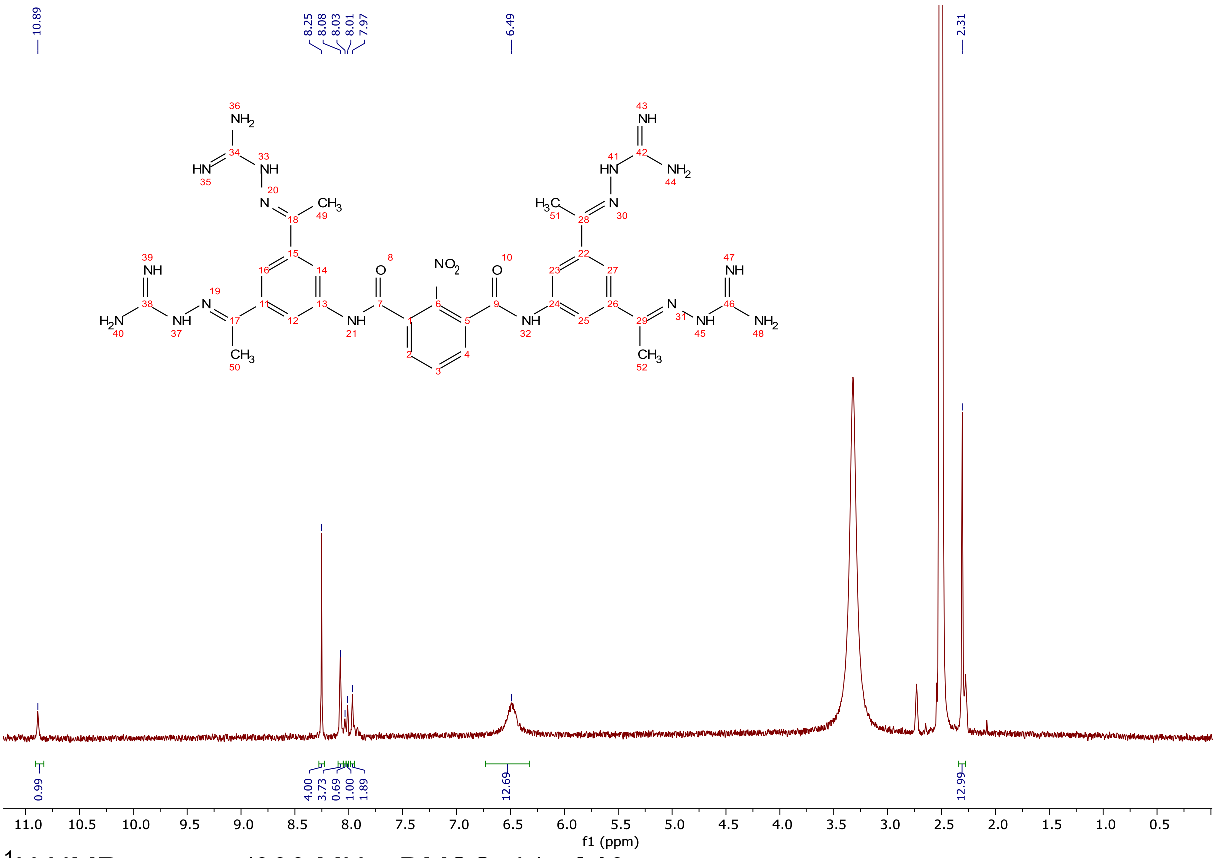

[1]H NMR spectra (300 MHz, DMSO-$d_6$) of **19**.

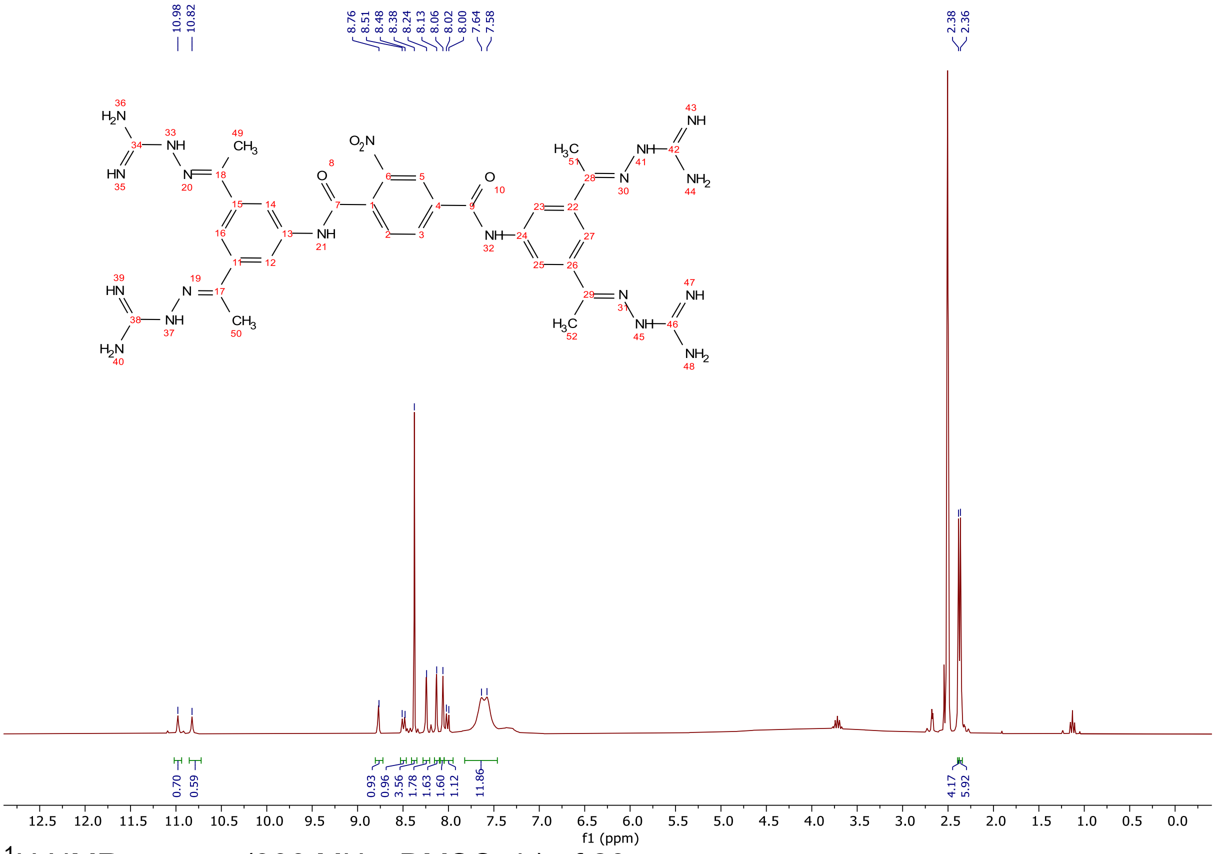

$^{1}$H NMR spectra (300 MHz, DMSO-$d_6$) of **20**.

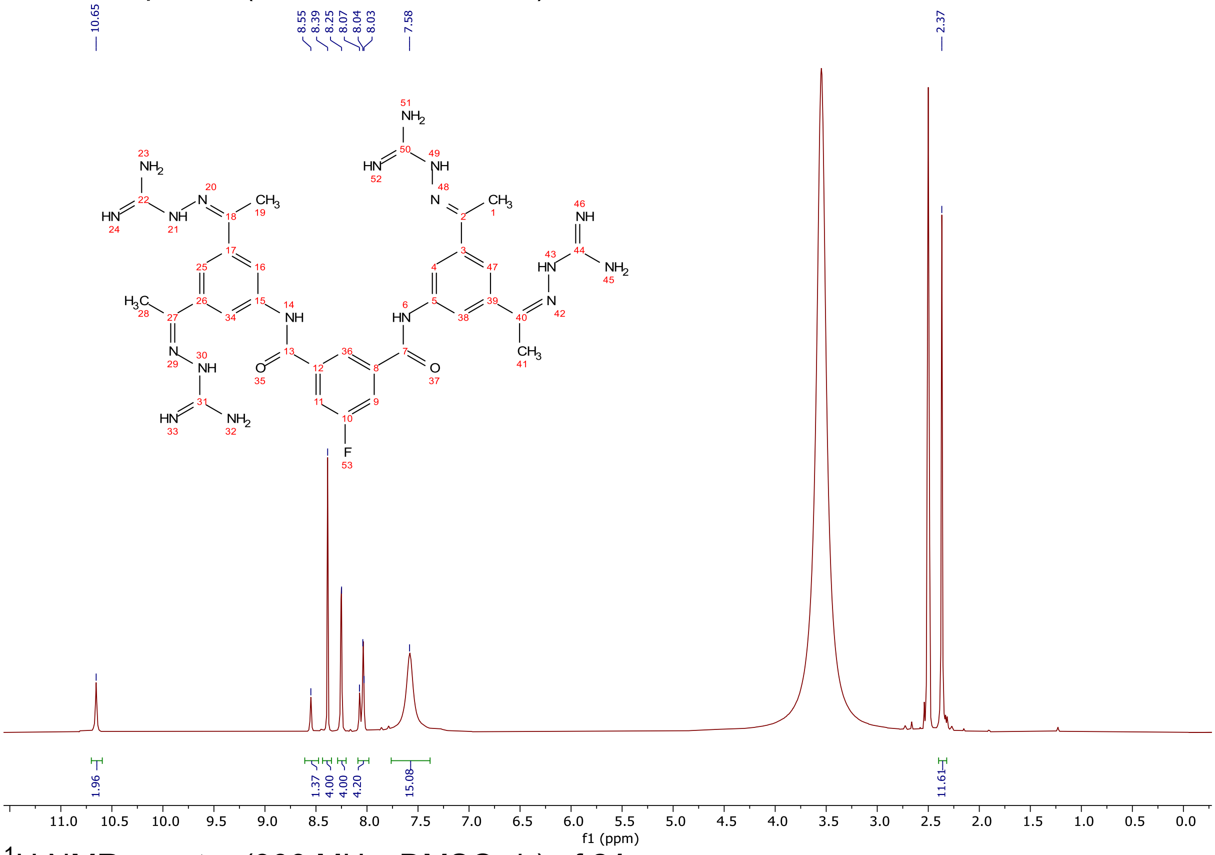

$^{1}$H NMR spectra (300 MHz, DMSO-$d_6$) of **21**.

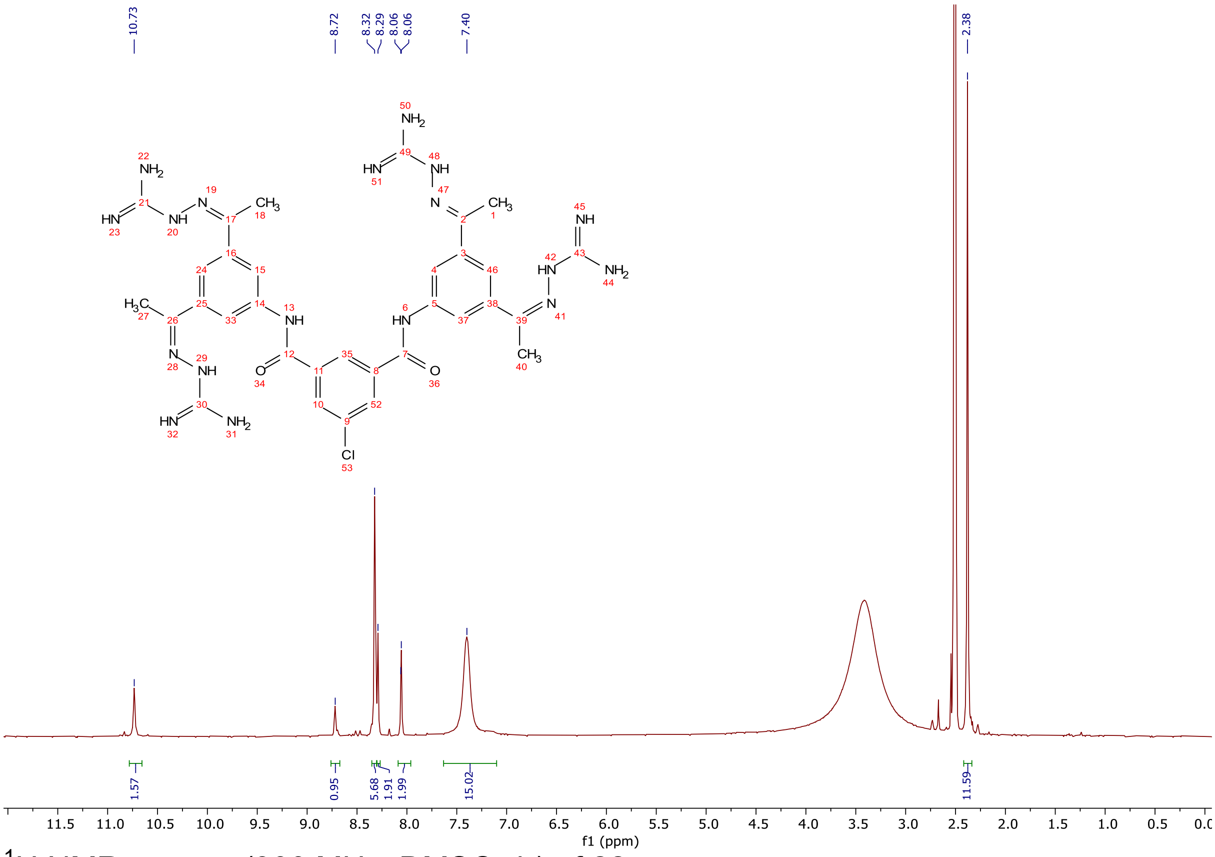

$^{1}H$ NMR spectra (300 MHz, DMSO-$d_6$) of **22.**

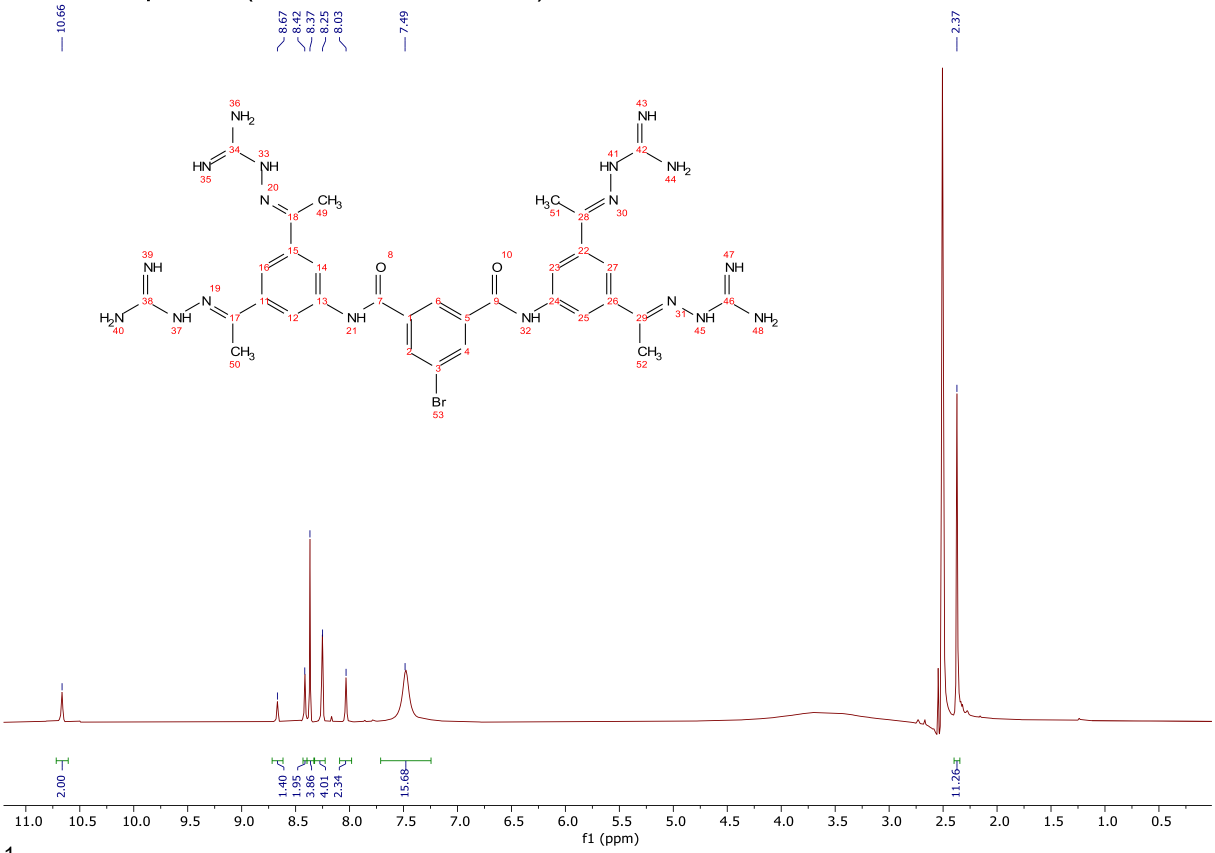

$^{1}H$ NMR spectra (300 MHz, DMSO-$d_6$) of **23**.

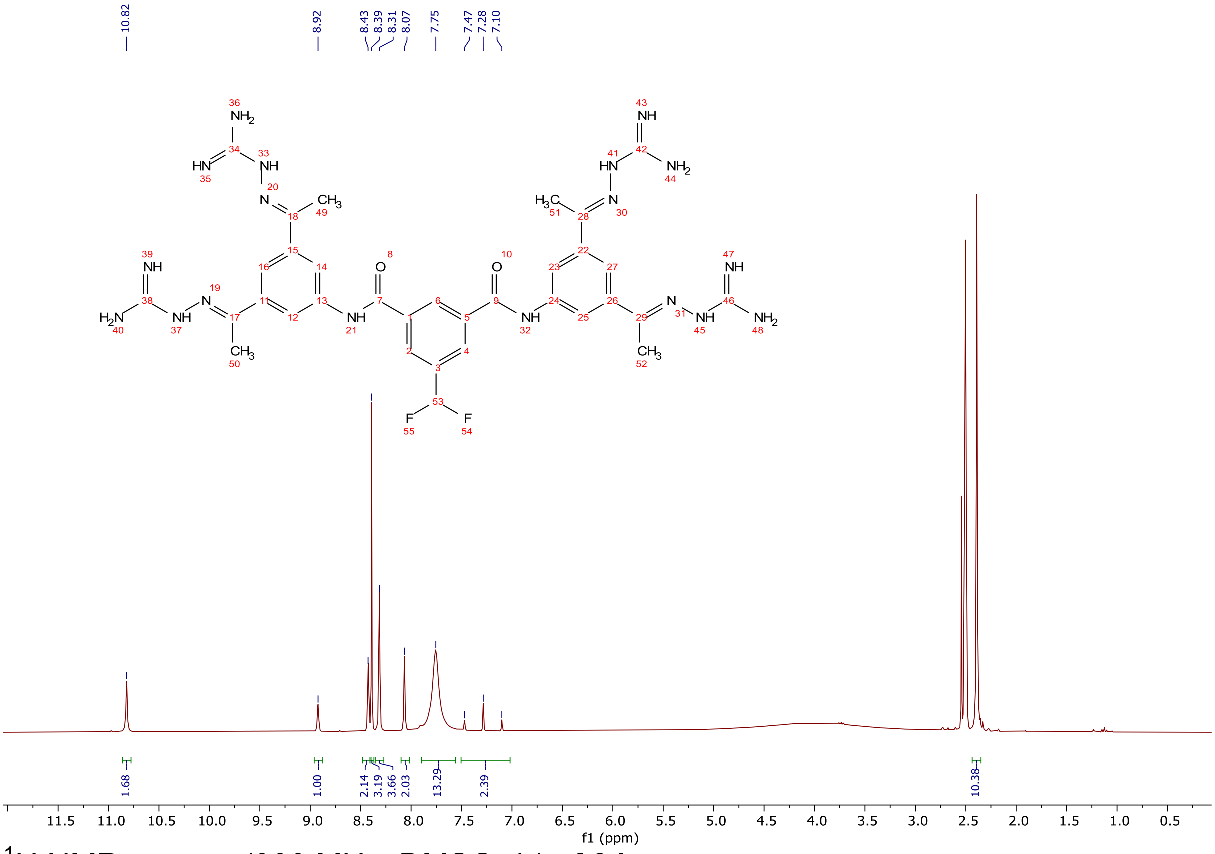


$^{1}H$ NMR spectra (300 MHz, DMSO-$d_6$) of **24**.

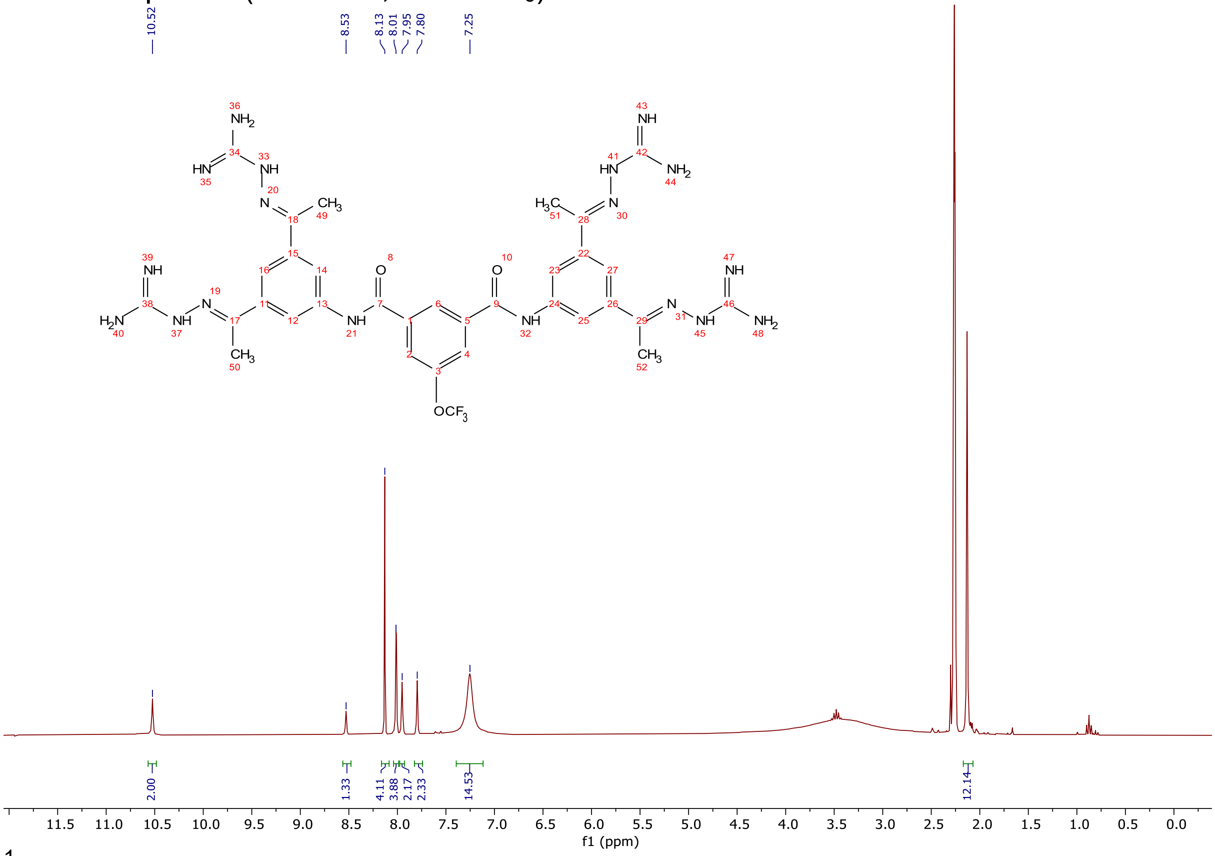


$^{1}H$ NMR spectra (300 MHz, DMSO-$d_6$) of **25**.

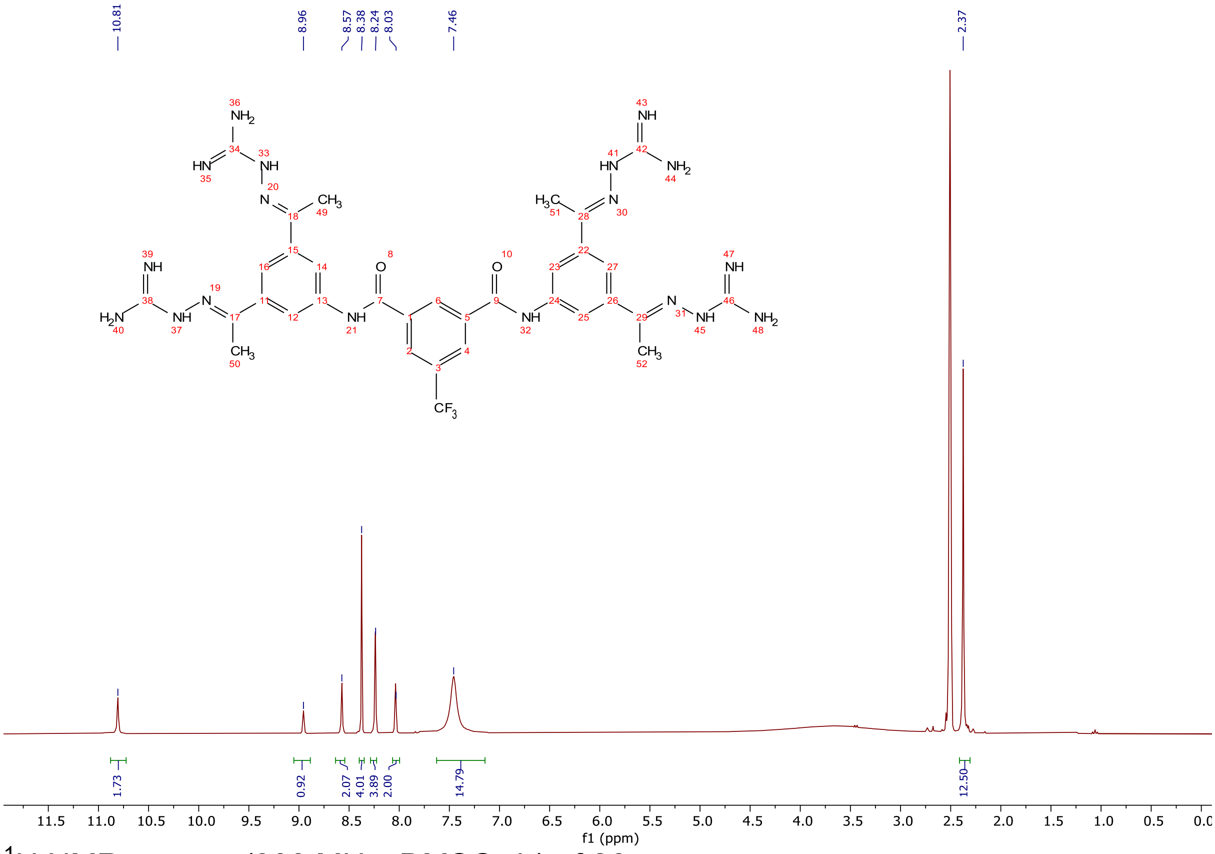


$^1$H NMR spectra (300 MHz, DMSO-d$_6$) of **26.**

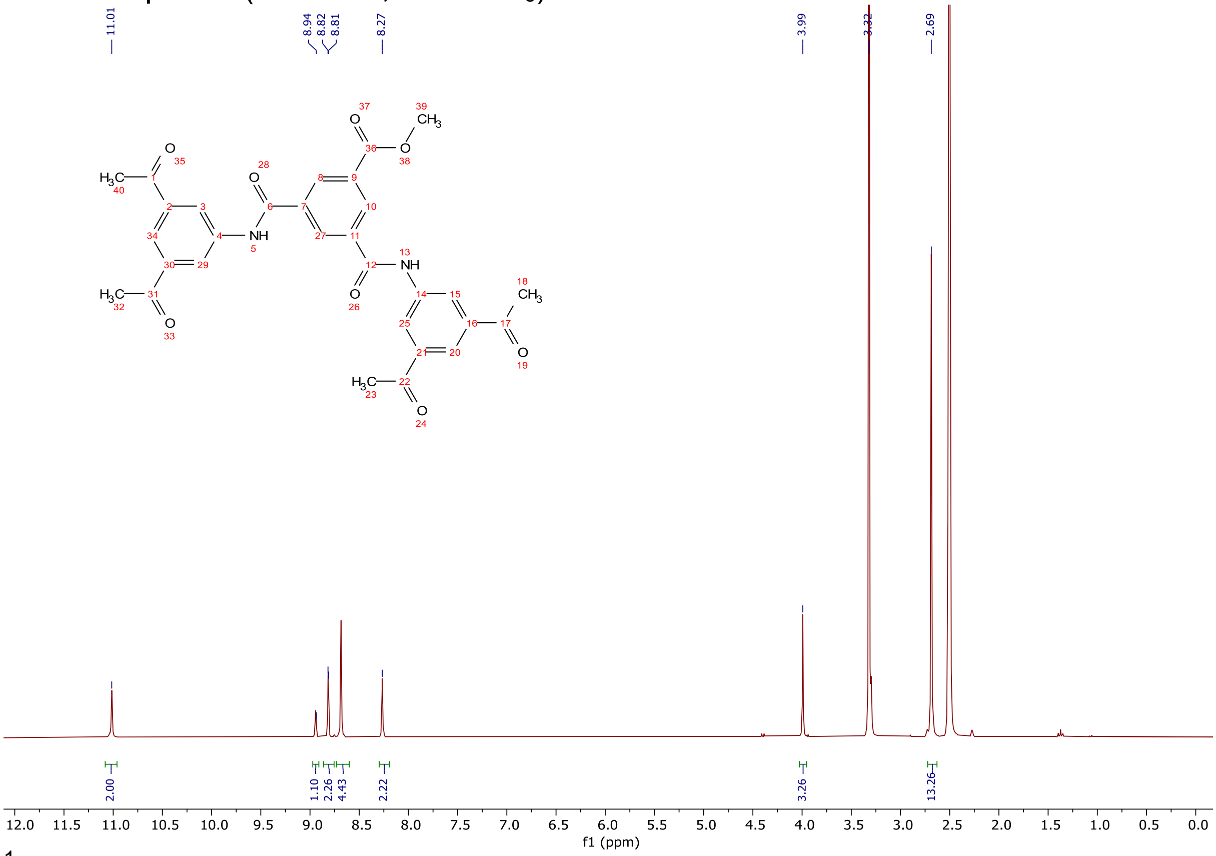


$^1$H NMR spectra (300 MHz, DMSO-d$_6$) of **27-1.**

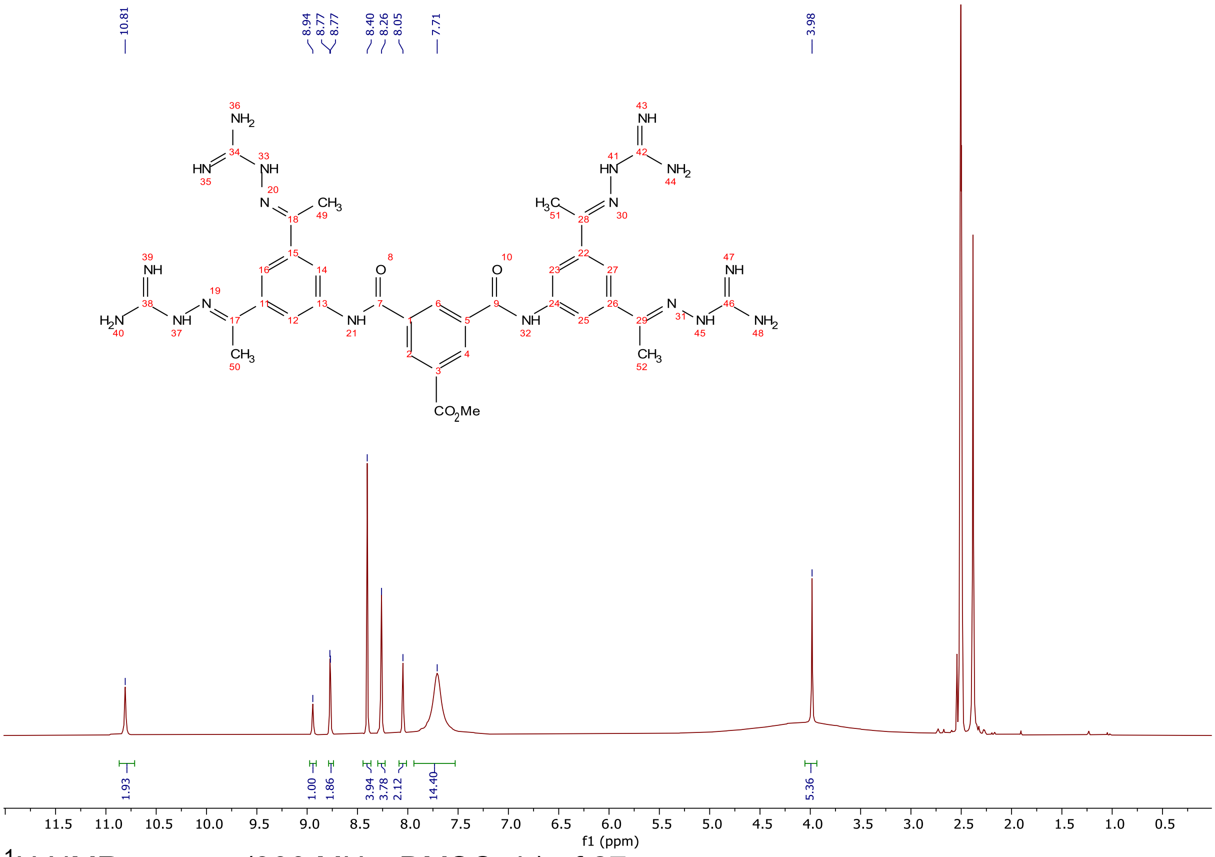

$^1$H NMR spectra (300 MHz, DMSO-$d_6$) of **27.**

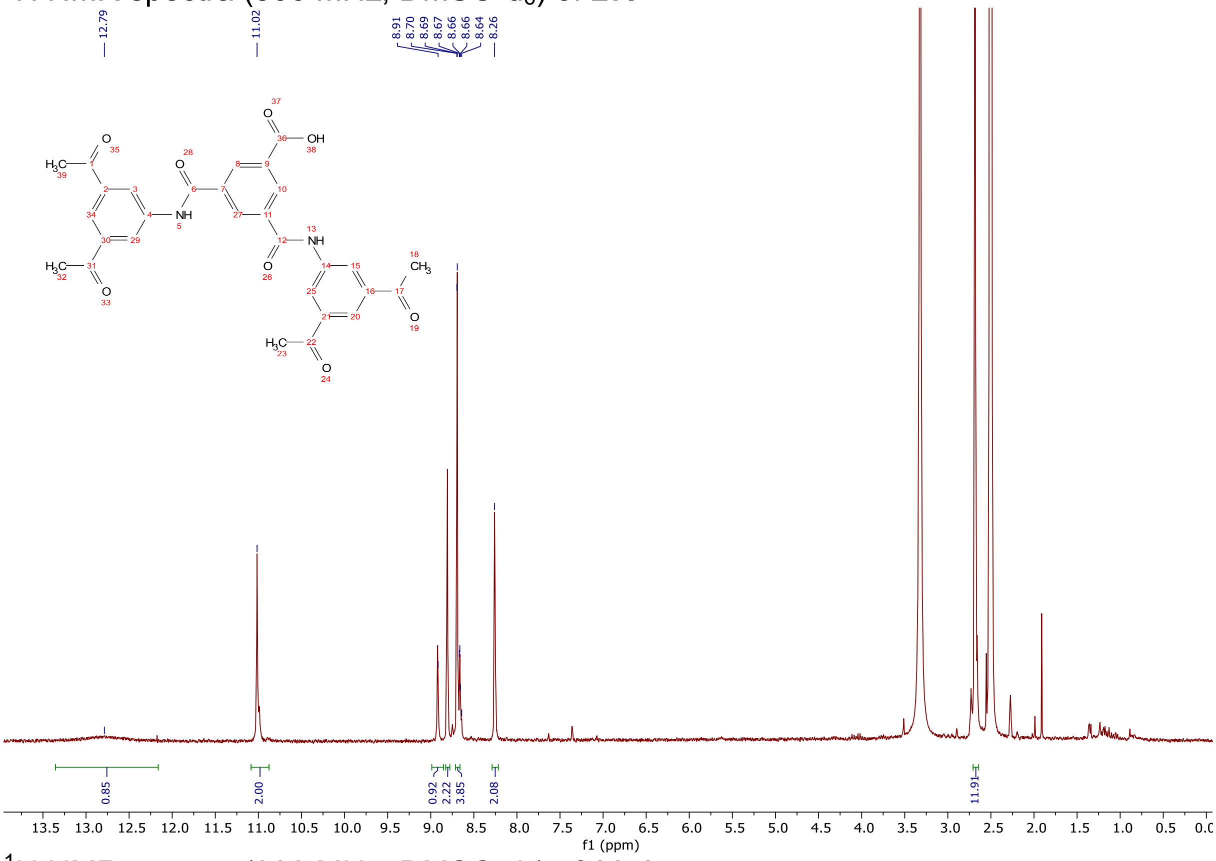

$^1$H NMR spectra (300 MHz, DMSO-$d_6$) of **28-1.**

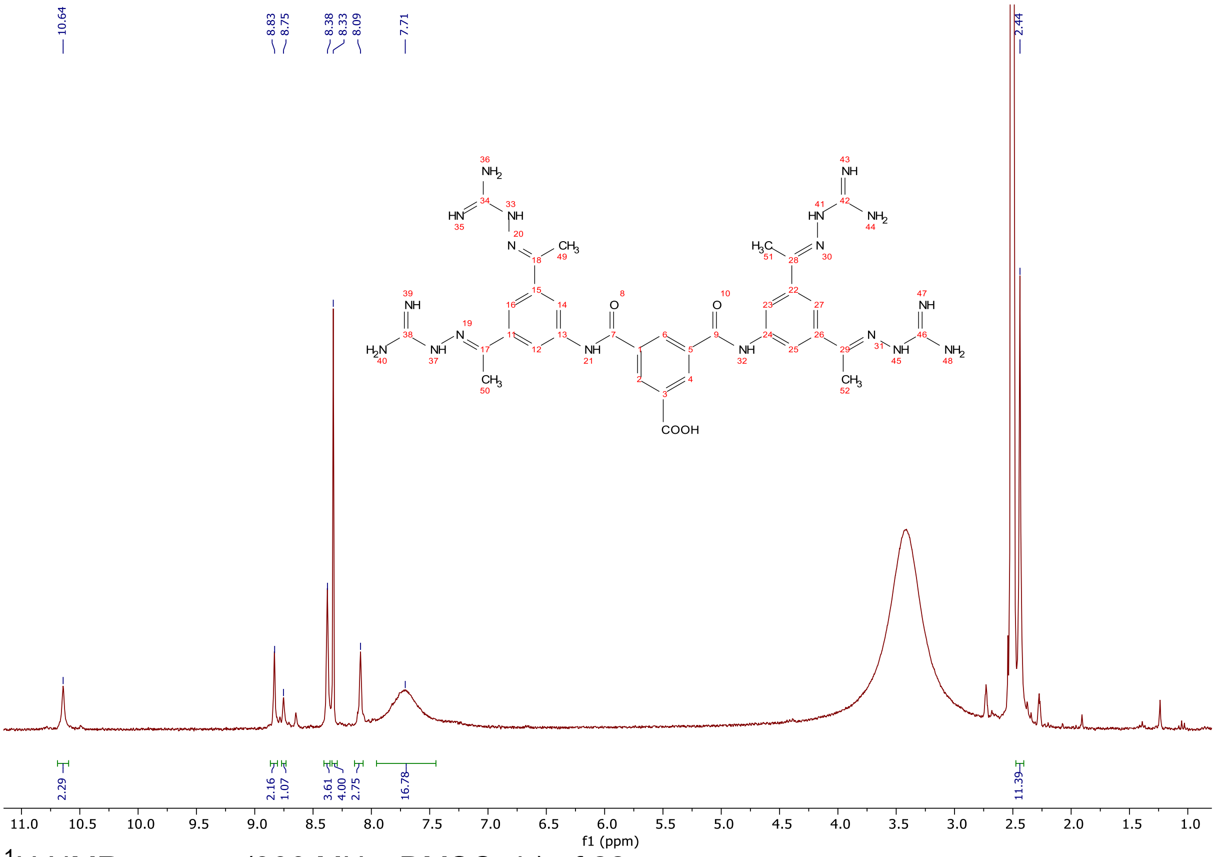

[1]H NMR spectra (300 MHz, DMSO-d$_6$) of **28.**

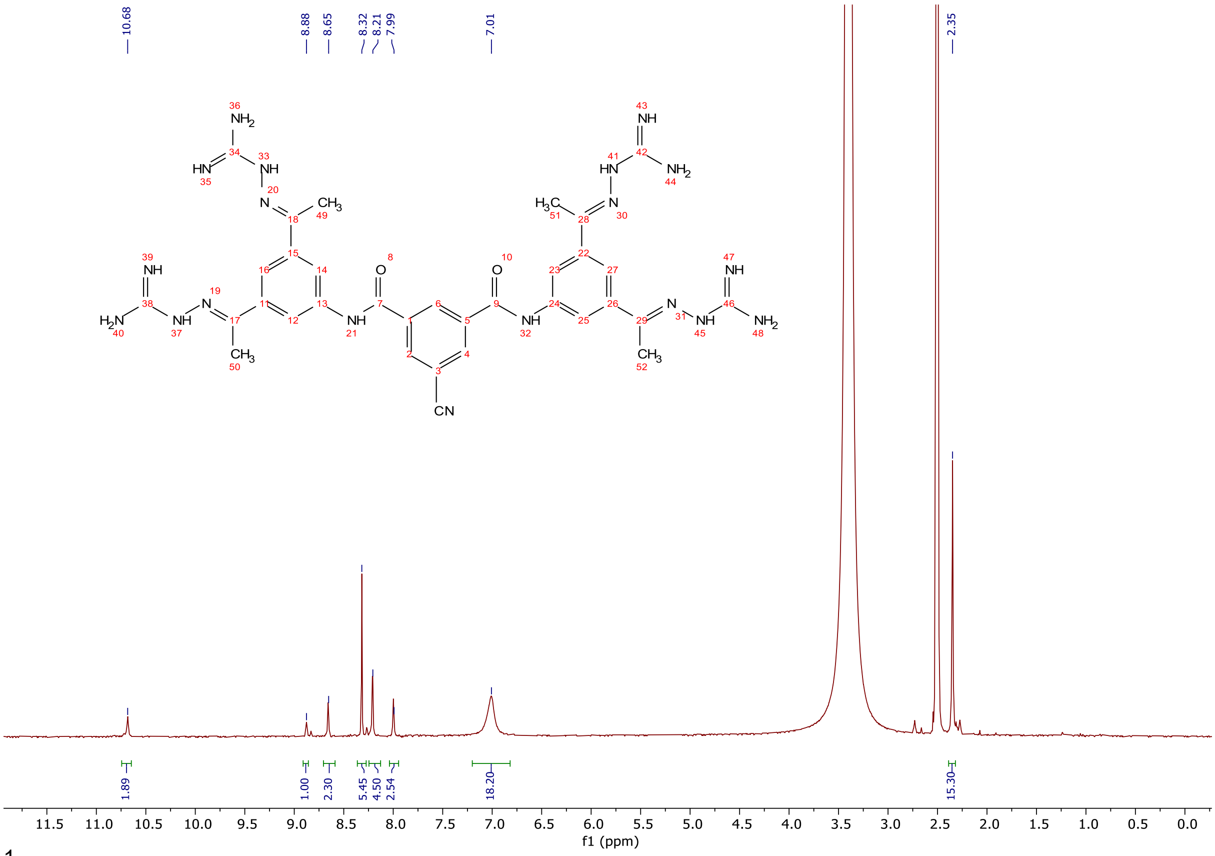

[1]H NMR spectra (300 MHz, DMSO-d$_6$) of **29.**

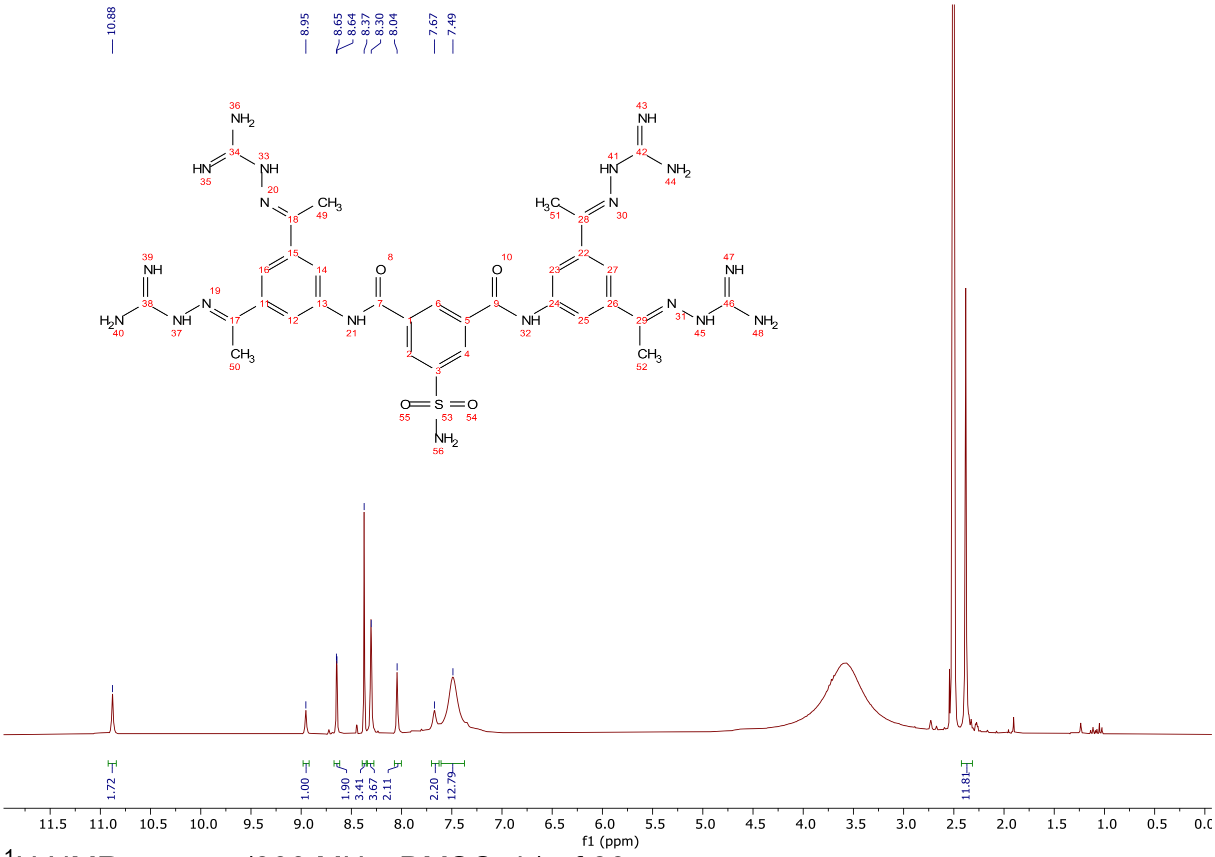

[1]H NMR spectra (300 MHz, DMSO-d$_6$) of **30.**

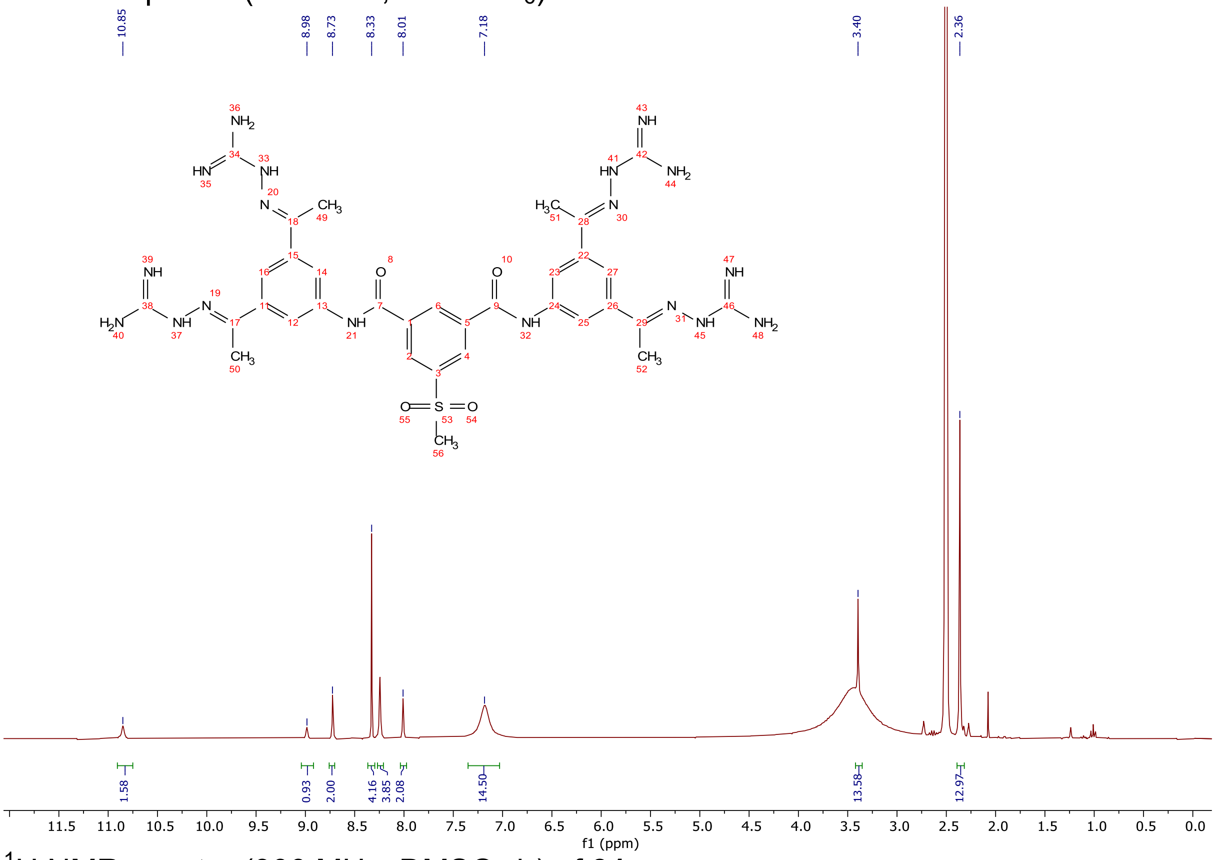

[1]H NMR spectra (300 MHz, DMSO-d$_6$) of **31.**

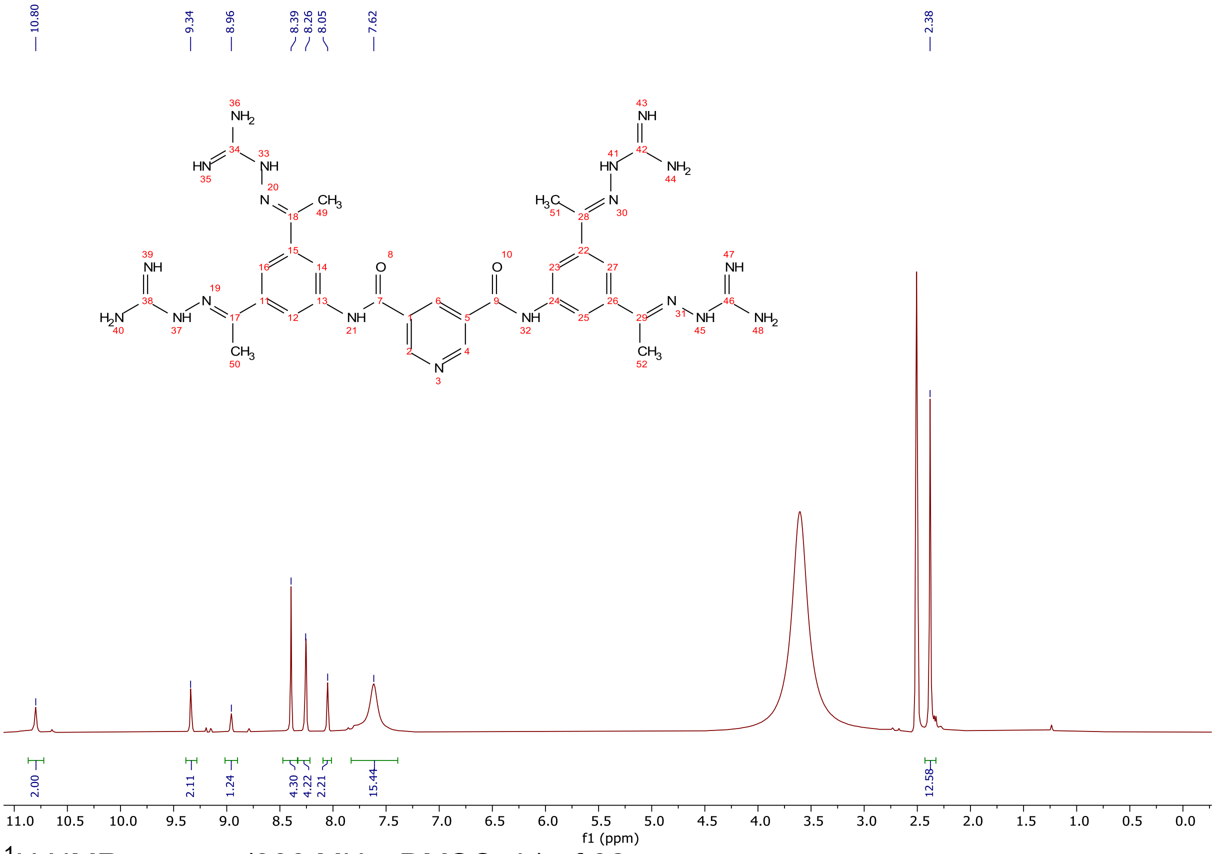


$^{1}$H NMR spectra (300 MHz, DMSO-d$_6$) of **32.**

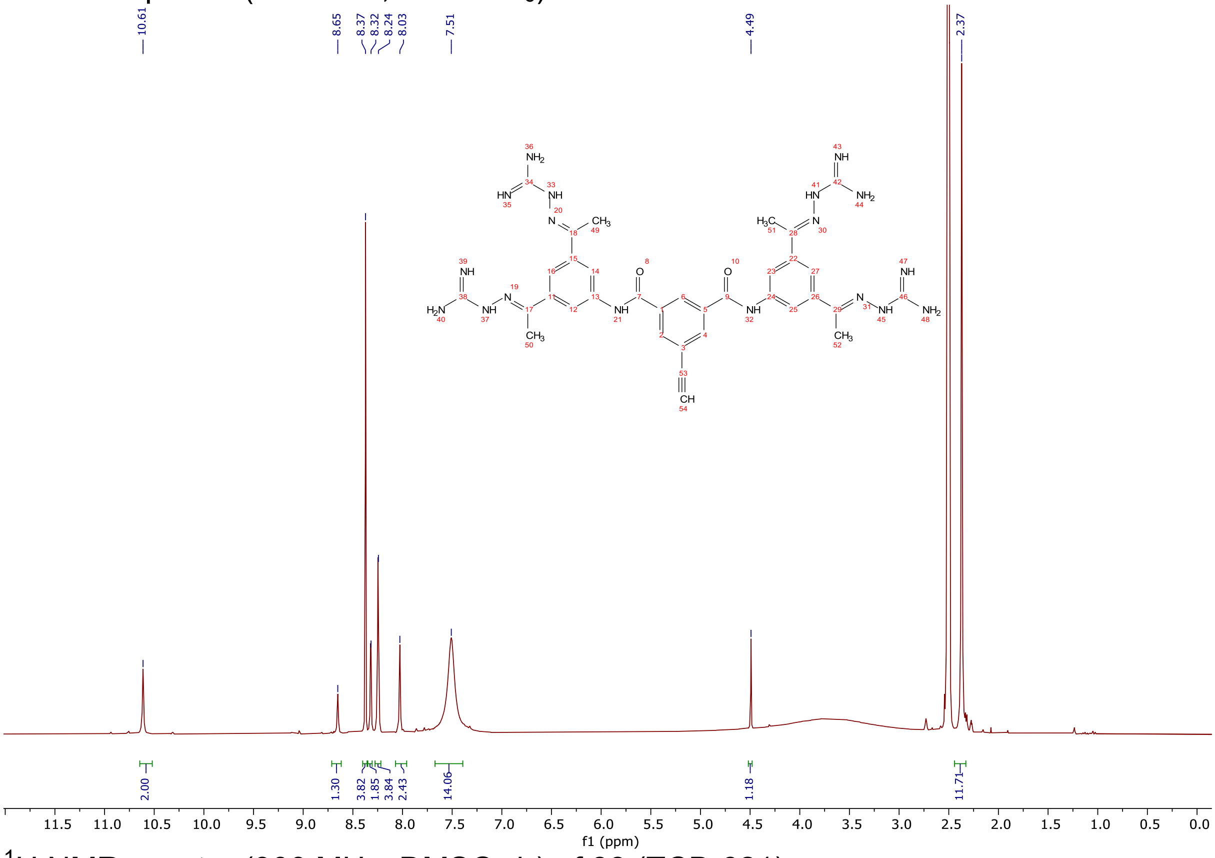


$^{1}$H NMR spectra (300 MHz, DMSO-d$_6$) of **33** (TCB-621)**.**

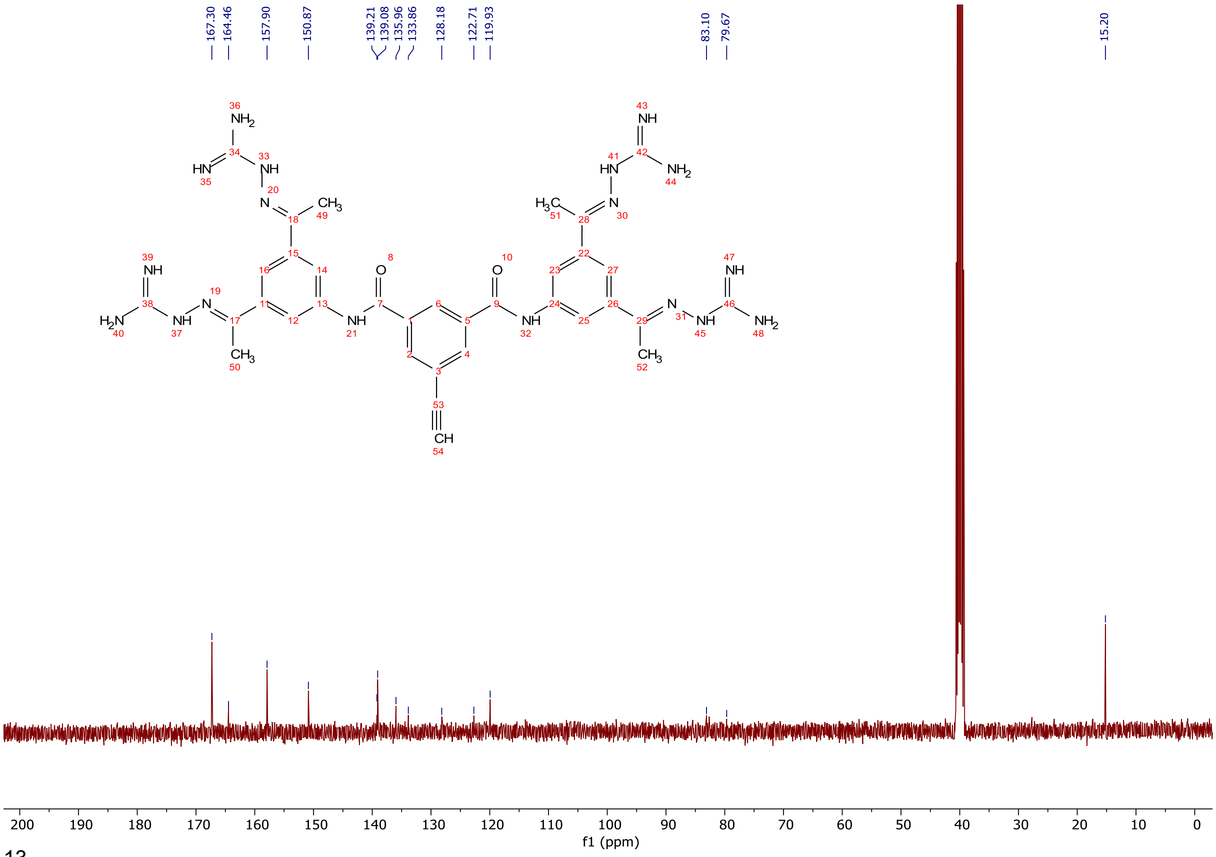

$^{13}$C NMR spectra (100 MHz, DMSO-d$_6$) of **33** (TCB-621).

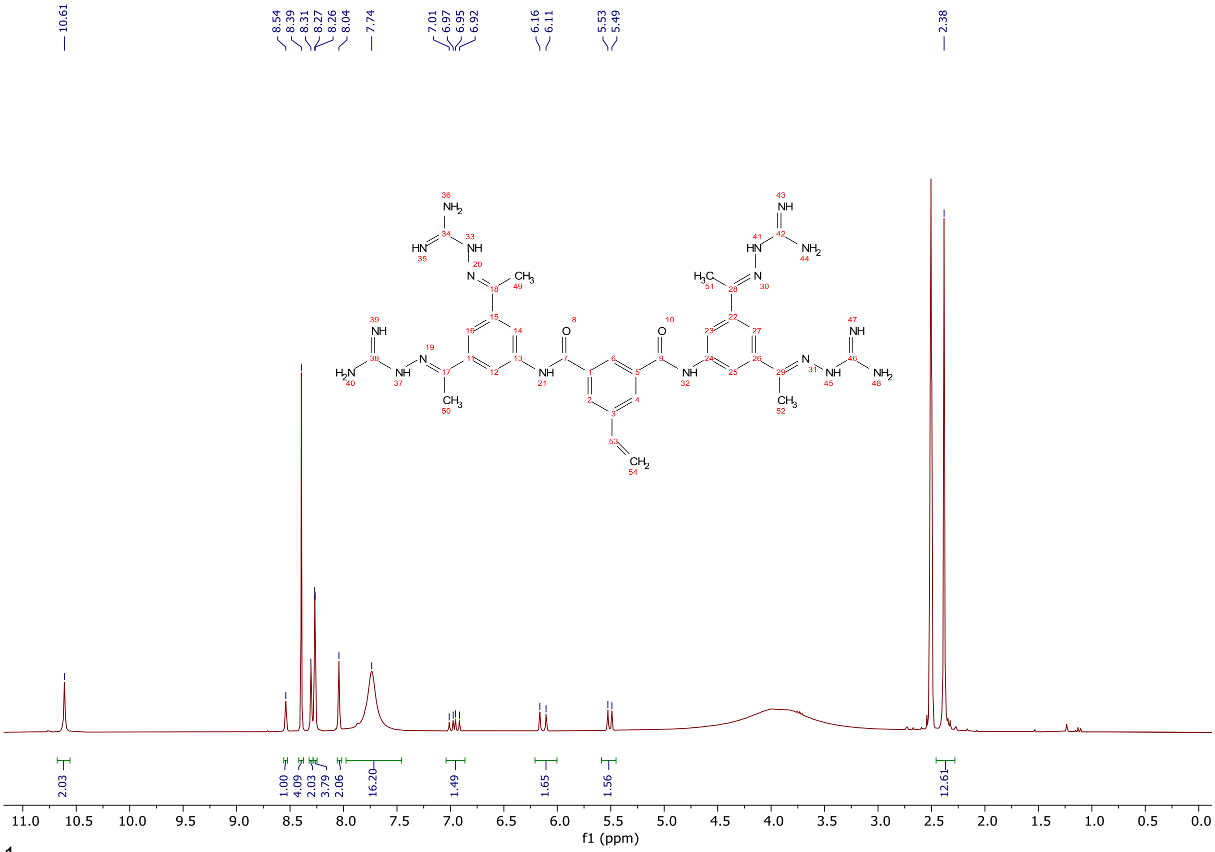

$^{1}$H NMR spectra (300 MHz, DMSO-d$_6$) of **34.**

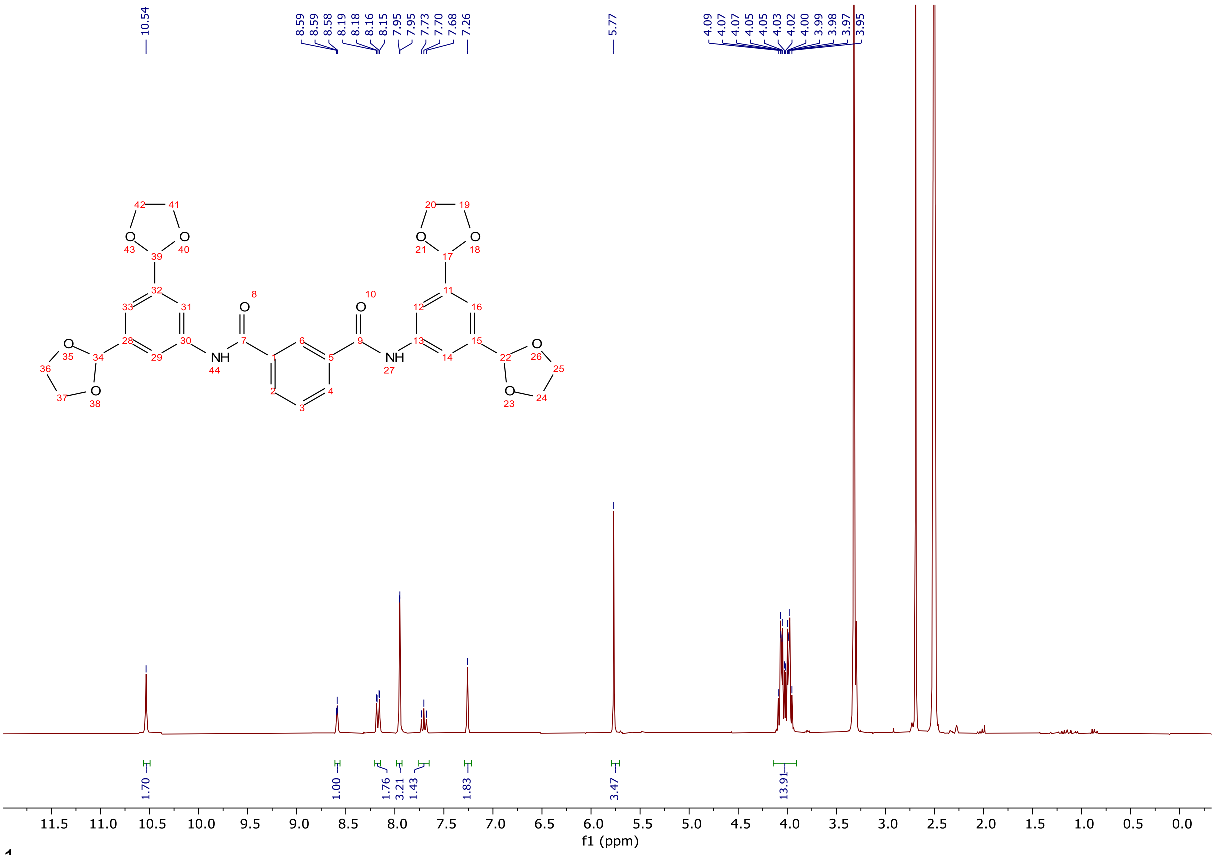

[1]H NMR spectra (300 MHz, DMSO-$d_6$) of **35-1.**

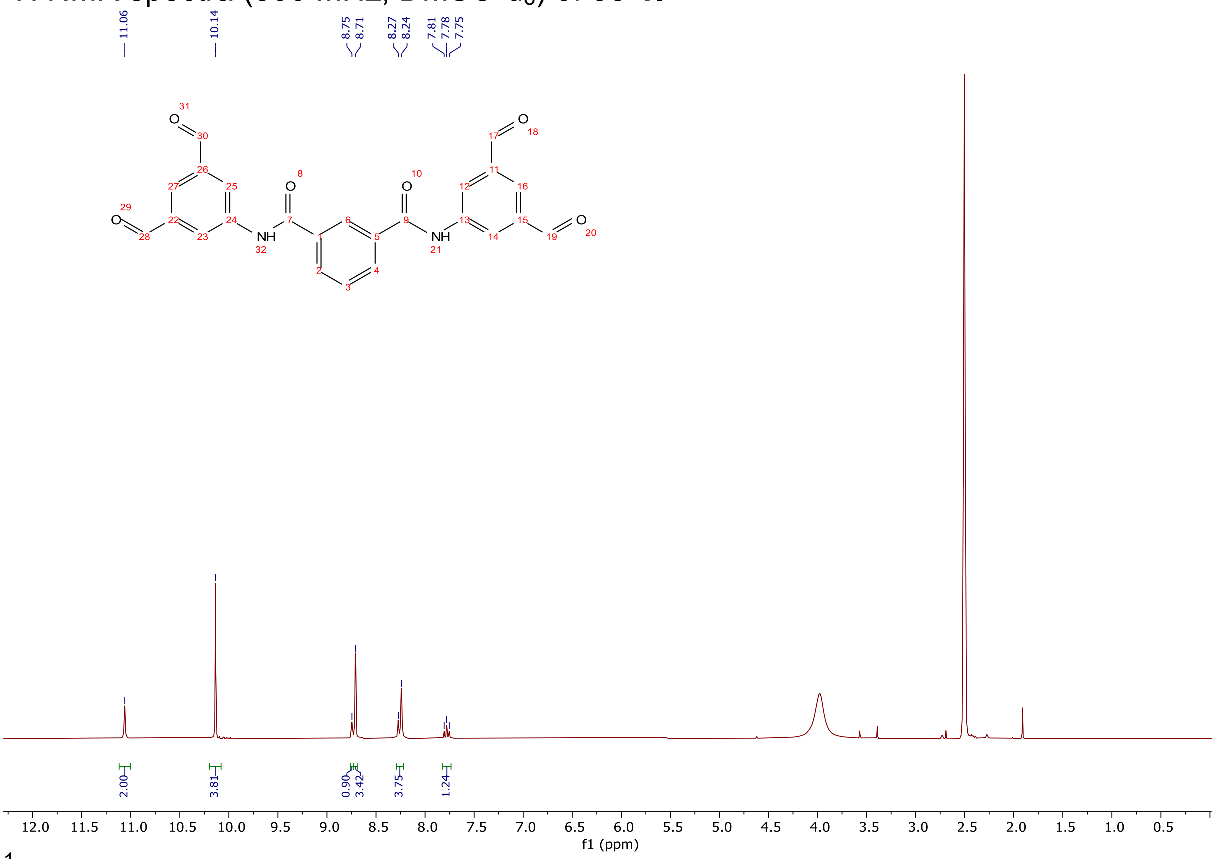

[1]H NMR spectra (300 MHz, DMSO-$d_6$) of **35-2.**

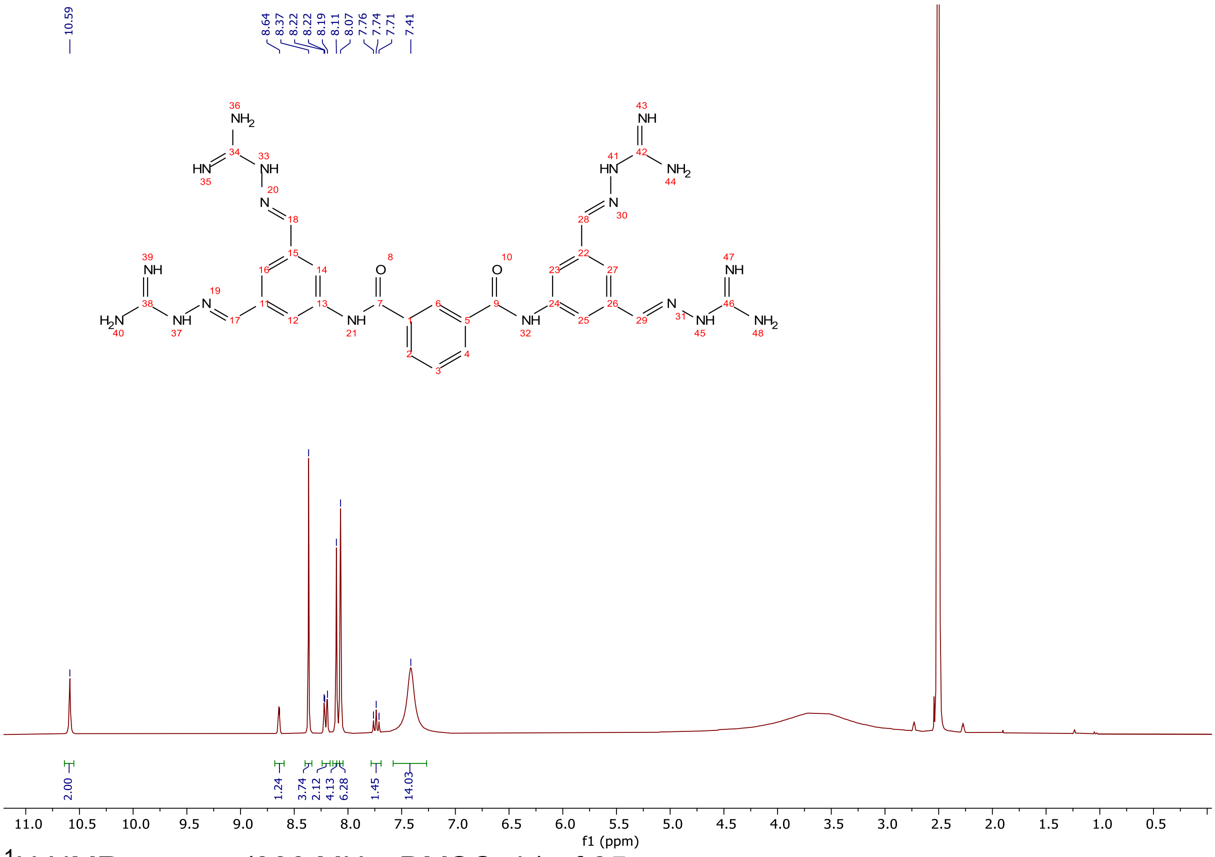


$^1$H NMR spectra (300 MHz, DMSO-$d_6$) of **35.**

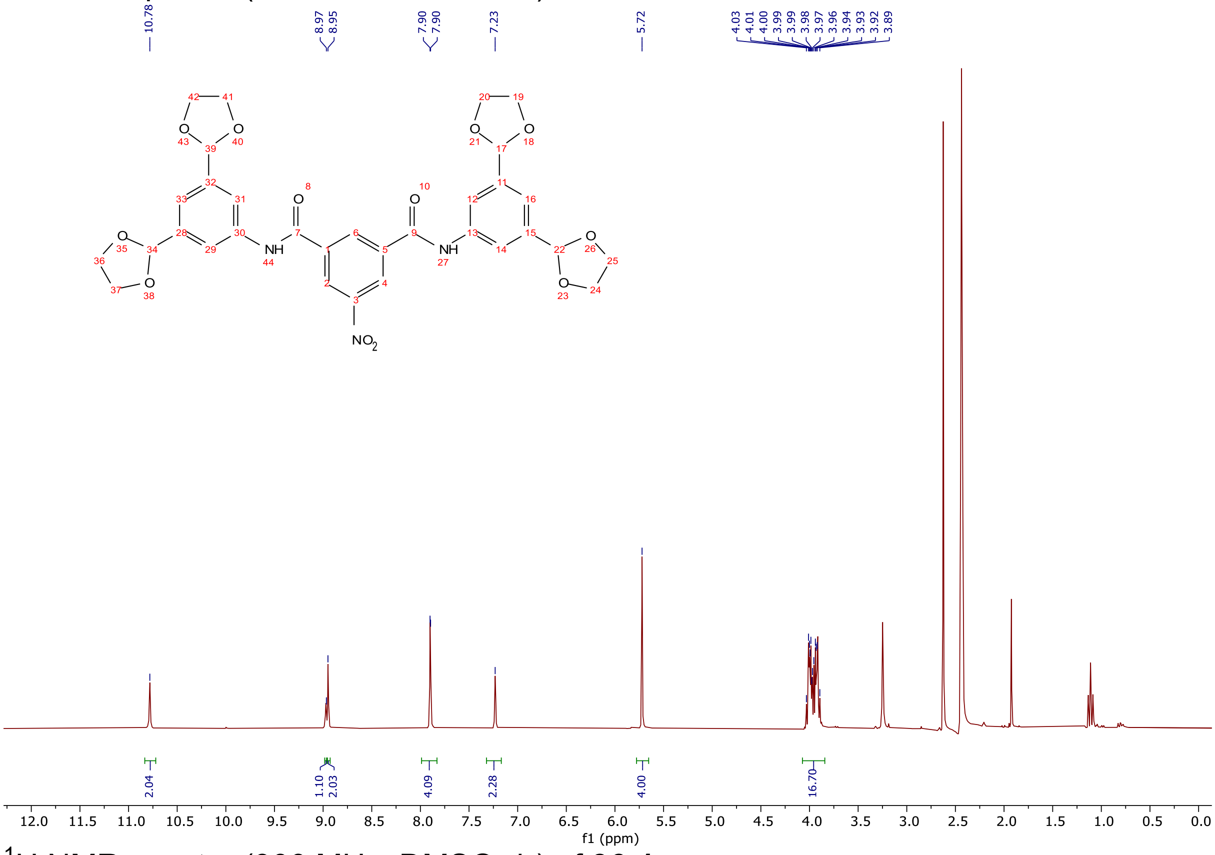


$^1$H NMR spectra (300 MHz, DMSO-$d_6$) of **36-1.**

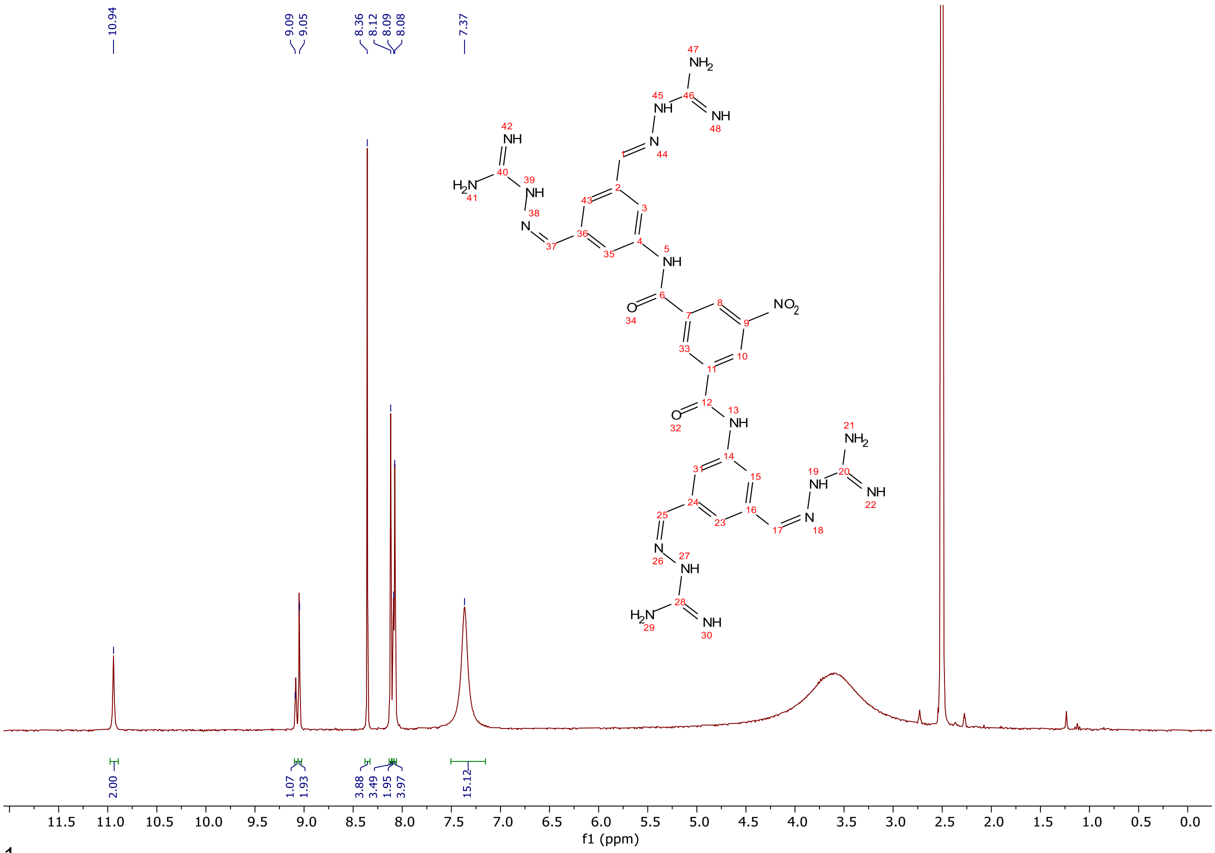


$^1H$ NMR spectra (300 MHz, DMSO-$d_6$) of **36.**

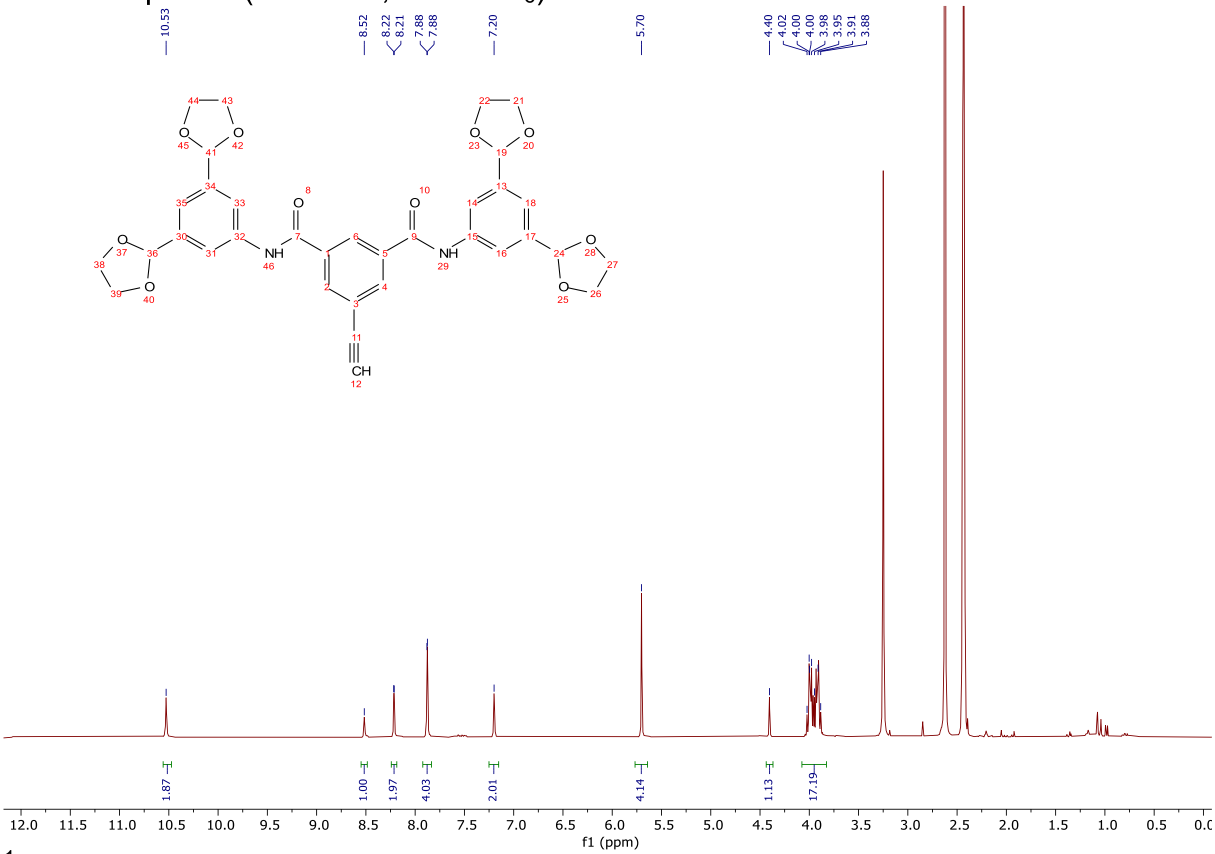


$^1H$ NMR spectra (300 MHz, DMSO-$d_6$) of **37-1.**

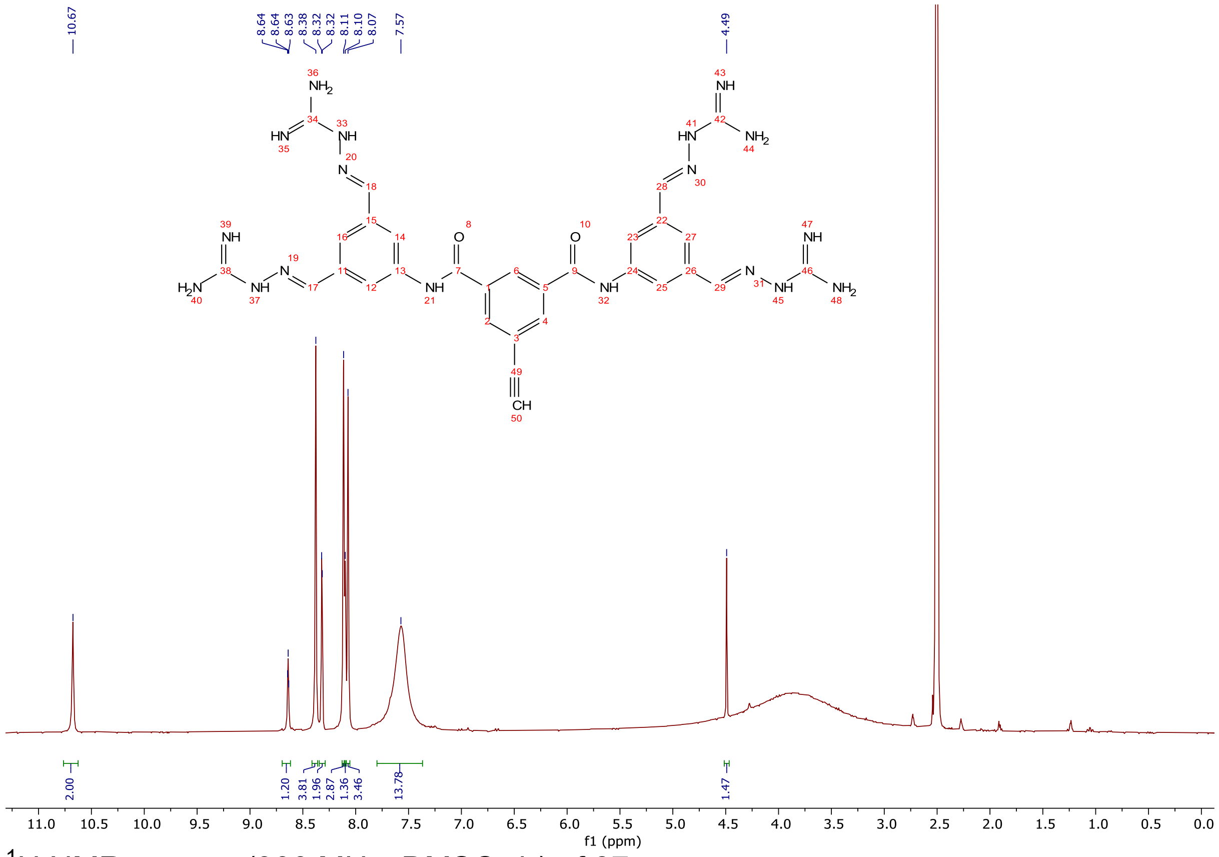


$^1$H NMR spectra (300 MHz, DMSO-d$_6$) of **37.**

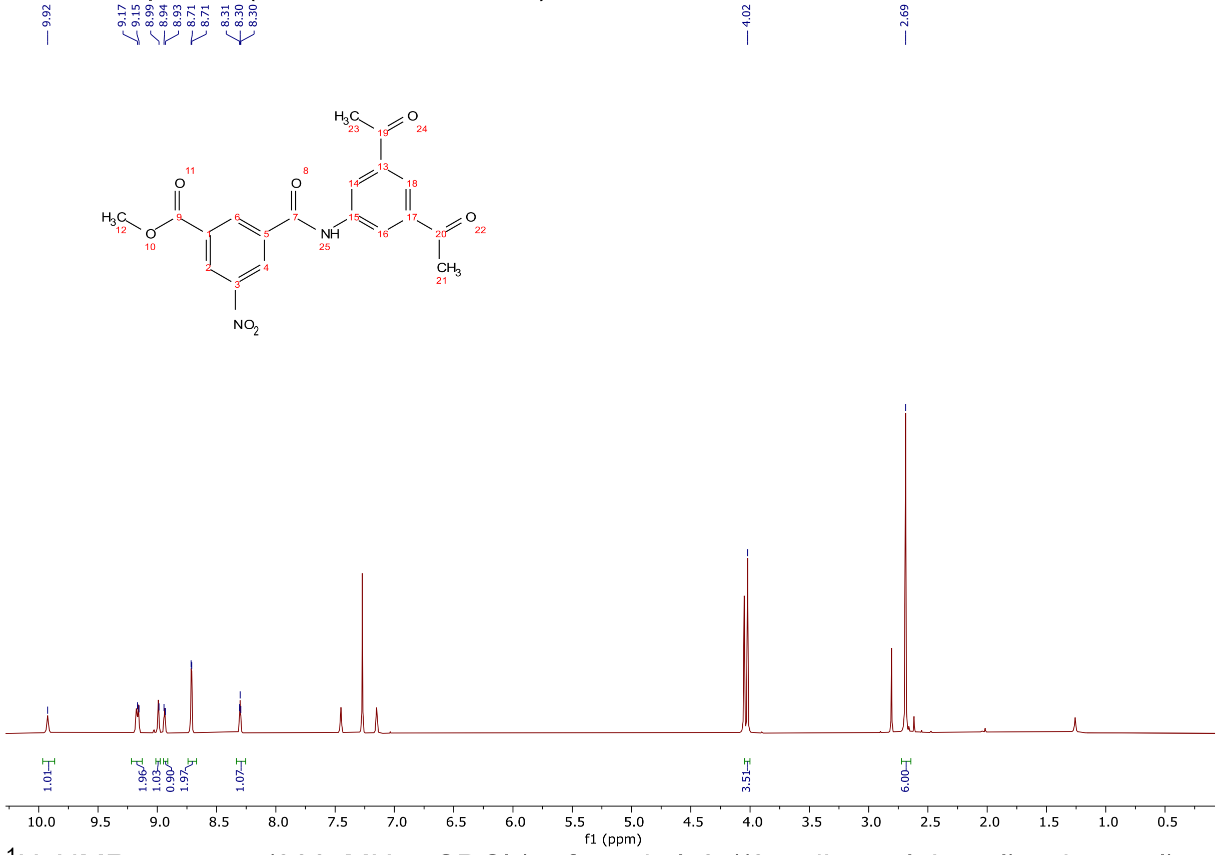


$^1$H NMR spectra (300 MHz, $CDCl_3$) of methyl 3-((3,5-diacetylphenyl)carbamoyl)-5-nitrobenzoate.

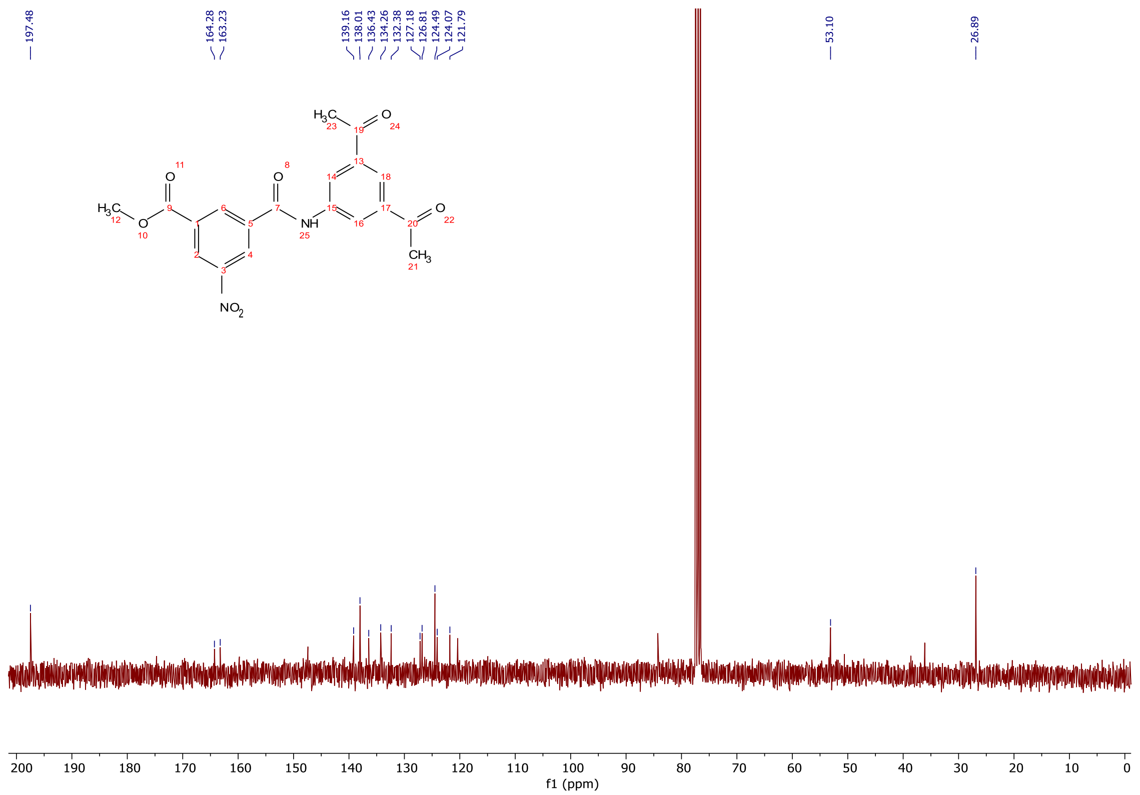


$^{13}C$ NMR spectra (75 MHz, $CDCl_3$) of methyl 3-((3,5-diacetylphenyl)carbamoyl)-5-nitrobenzoate.

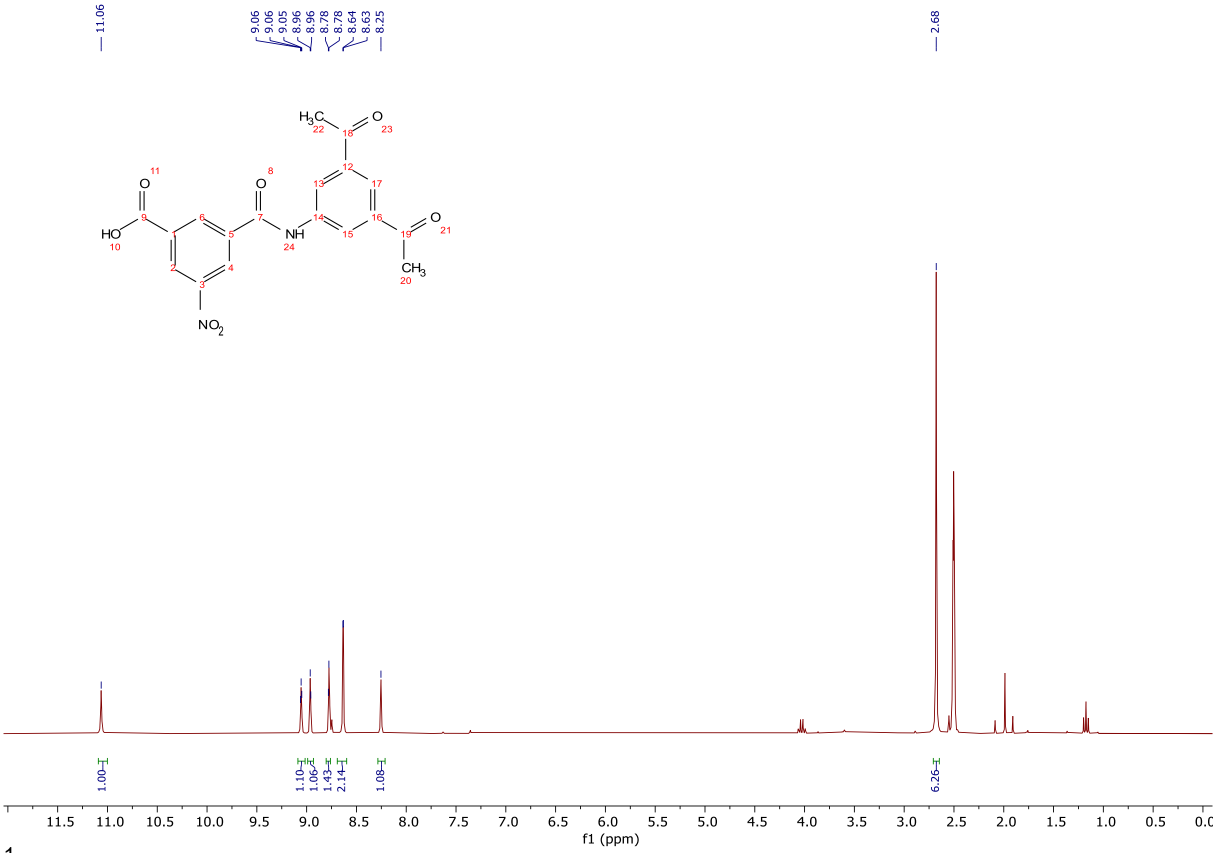


$^{1}H$ NMR spectra (300 MHz, DMSO-$d_6$) of 3-((3,5-diacetylphenyl)carbamoyl)-5-nitrobenzoic acid.

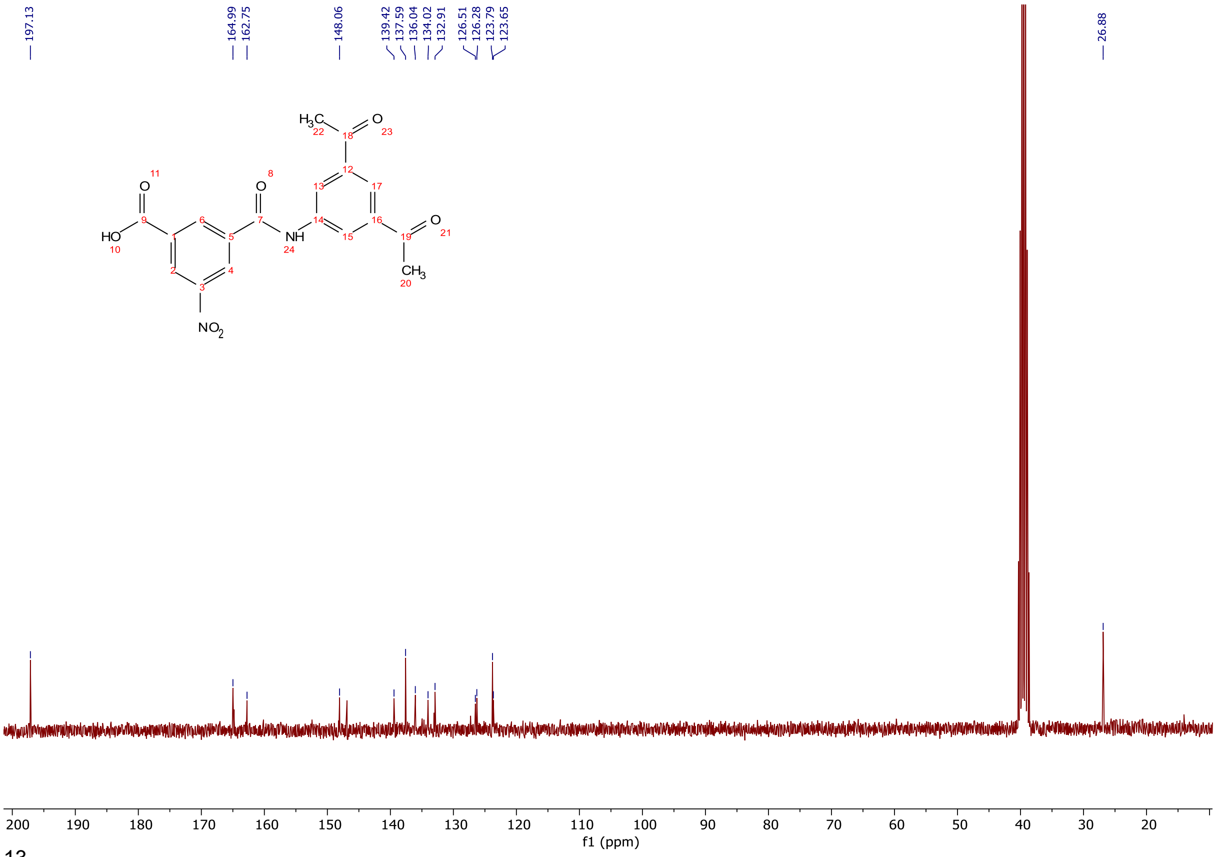

$^{13}C$ NMR spectra (75 MHz, DMSO-$d_6$) of 3-((3,5-diacetylphenyl)carbamoyl)-5-nitrobenzoic acid.

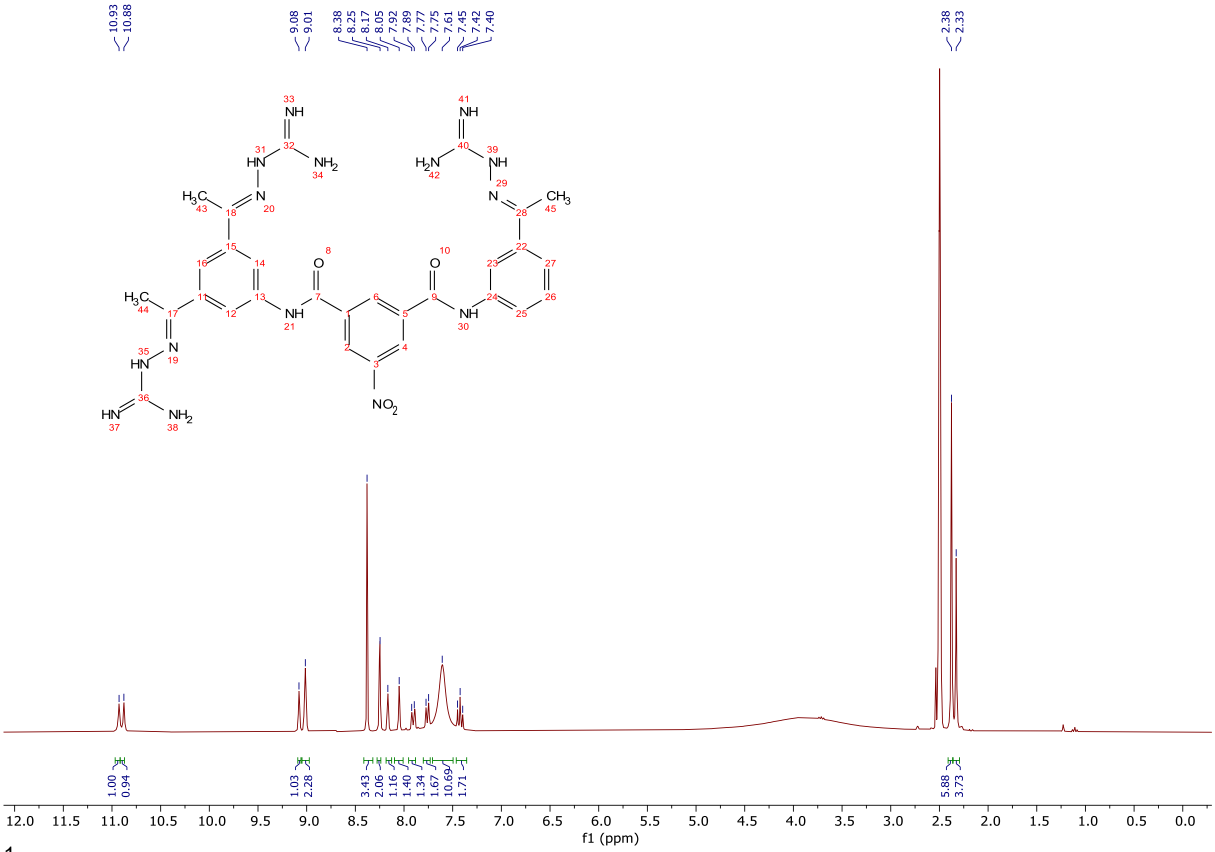

$^{1}H$ NMR spectra (300 MHz, DMSO-$d_6$) of **38.**

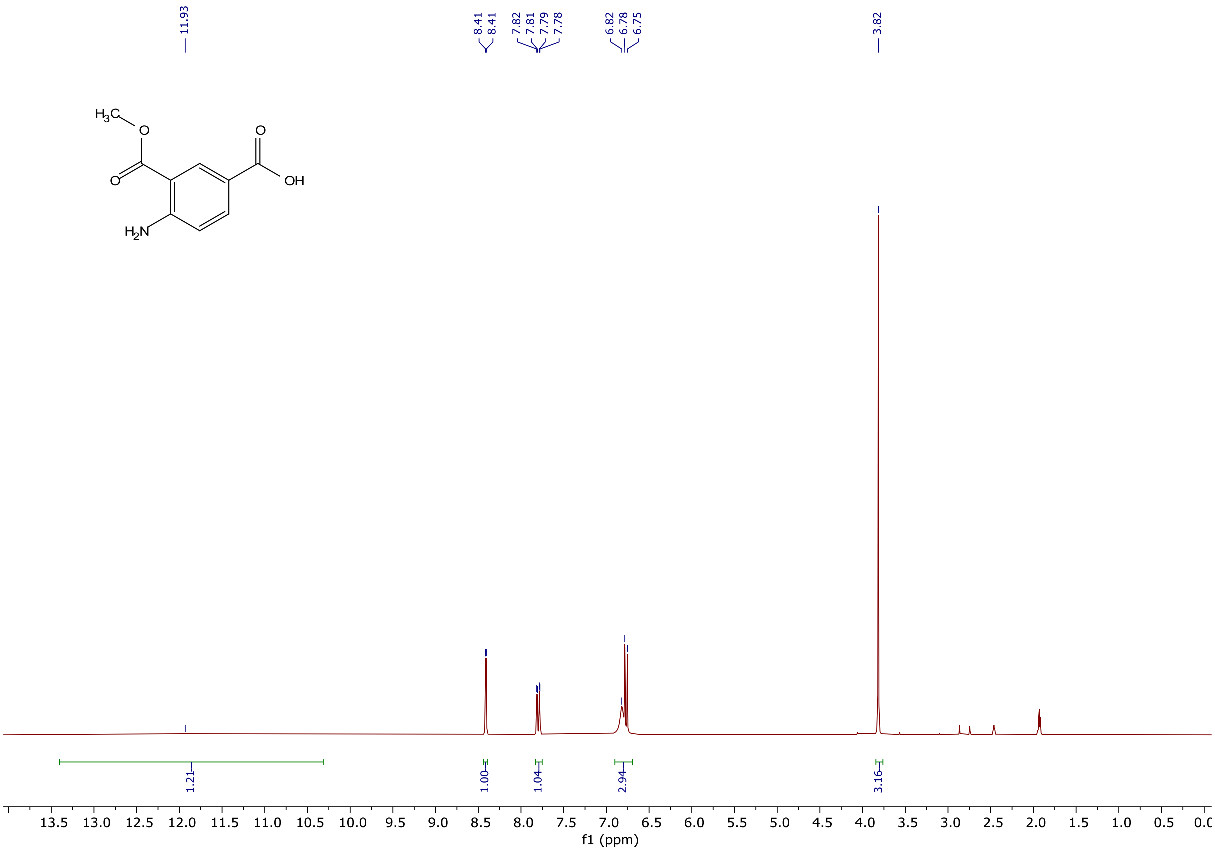


$^1$H NMR spectra (300 MHz, $CDCl_3$) of 4-amino-3-(methoxycarbonyl)benzoic acid.

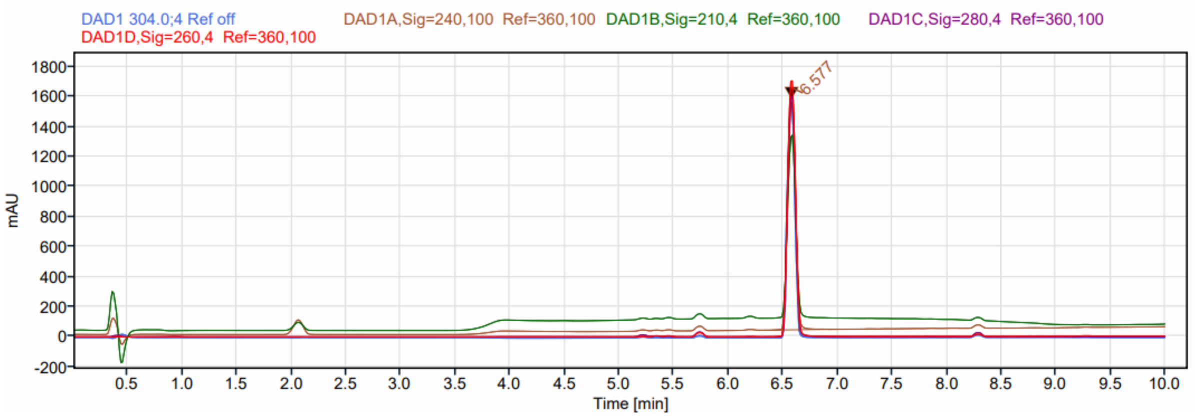


HPLC profile of 3-(methoxycarbonyl)-4-(2,2,2-trifluoroacetamido)benzoic acid.

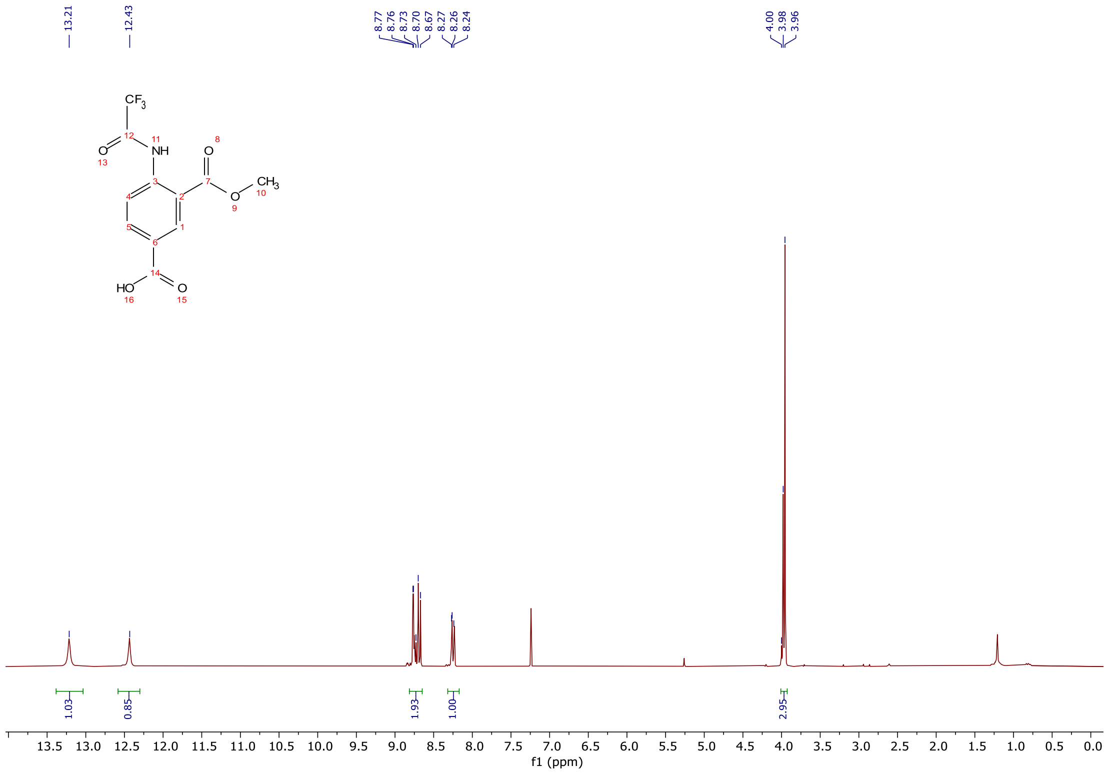


$^1H$ NMR spectra (300 MHz, $CDCl_3$) of 3-(methoxycarbonyl)-4-(2,2,2-trifluoroacetamido)benzoic acid.

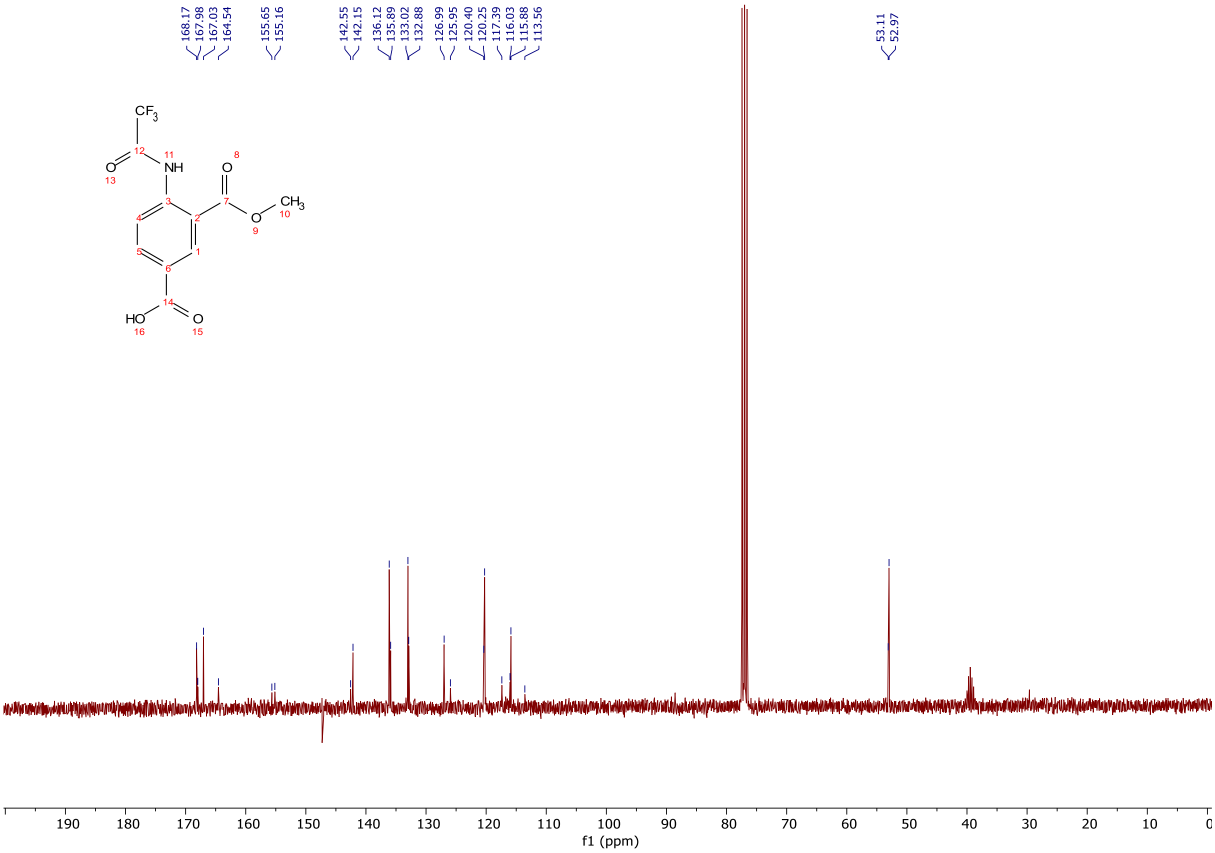


$^{13}C$ NMR spectra (75 MHz, $CDCl_3$) of 3-(methoxycarbonyl)-4-(2,2,2-trifluoroacetamido)benzoic acid.

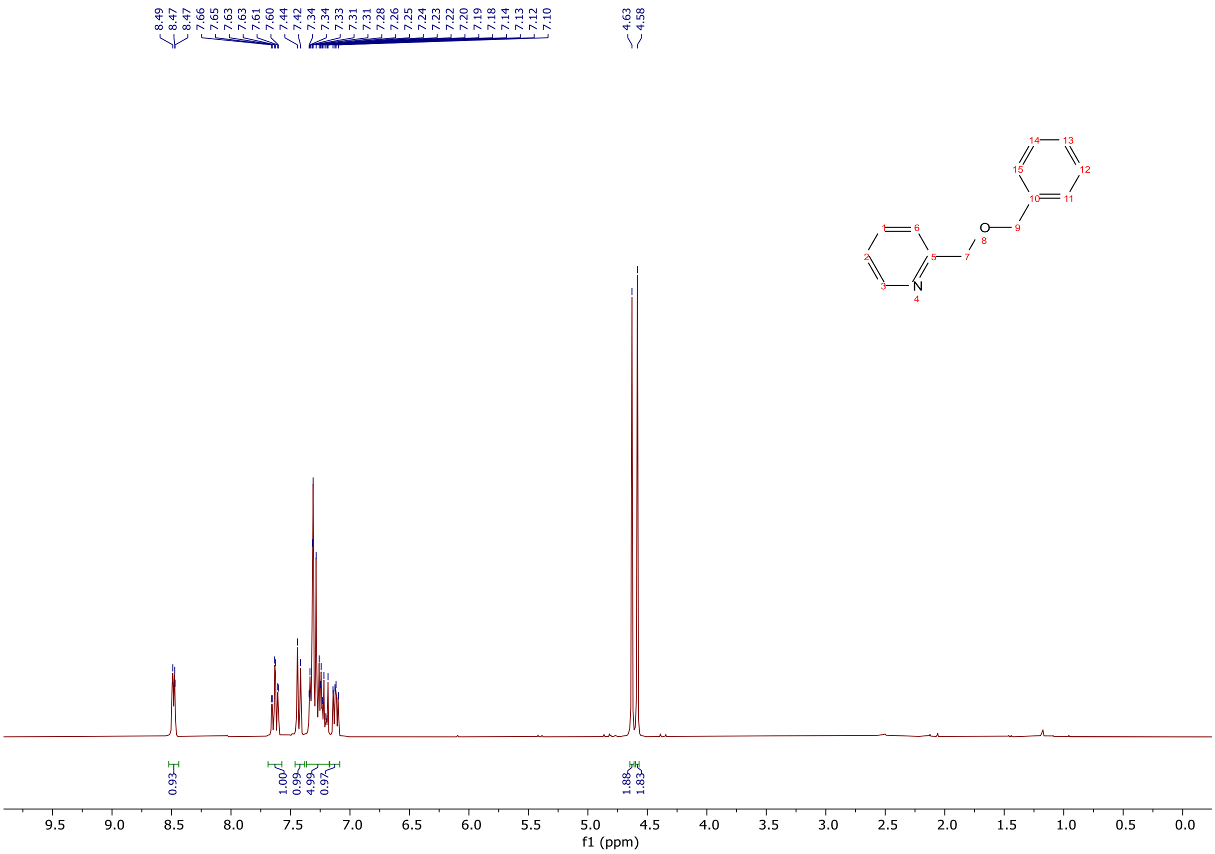

$^{1}H$ NMR spectra (300 MHz, $CDCl_3$) of 2-((benzyloxy)methyl)pyridine.

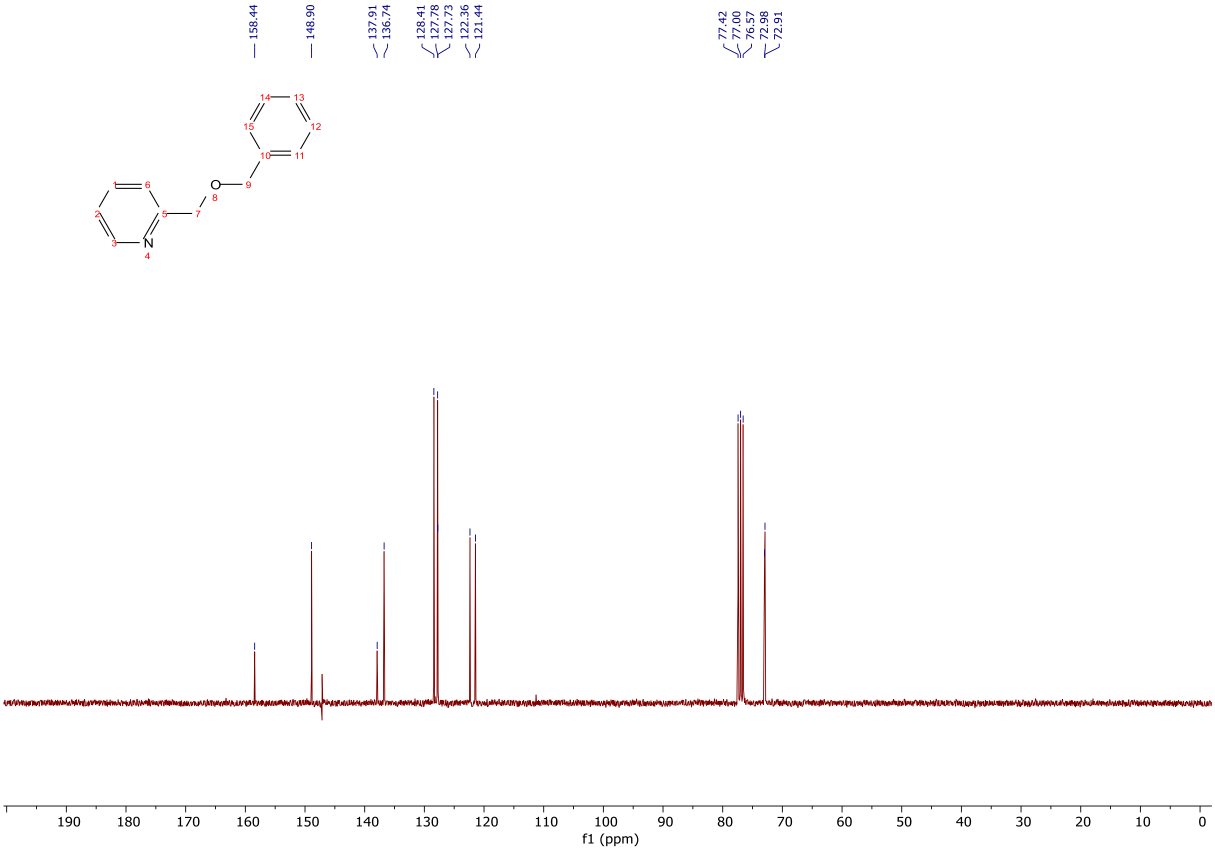

$^{13}C$ NMR spectra (300 MHz, $CDCl_3$) of 2-((benzyloxy)methyl)pyridine.

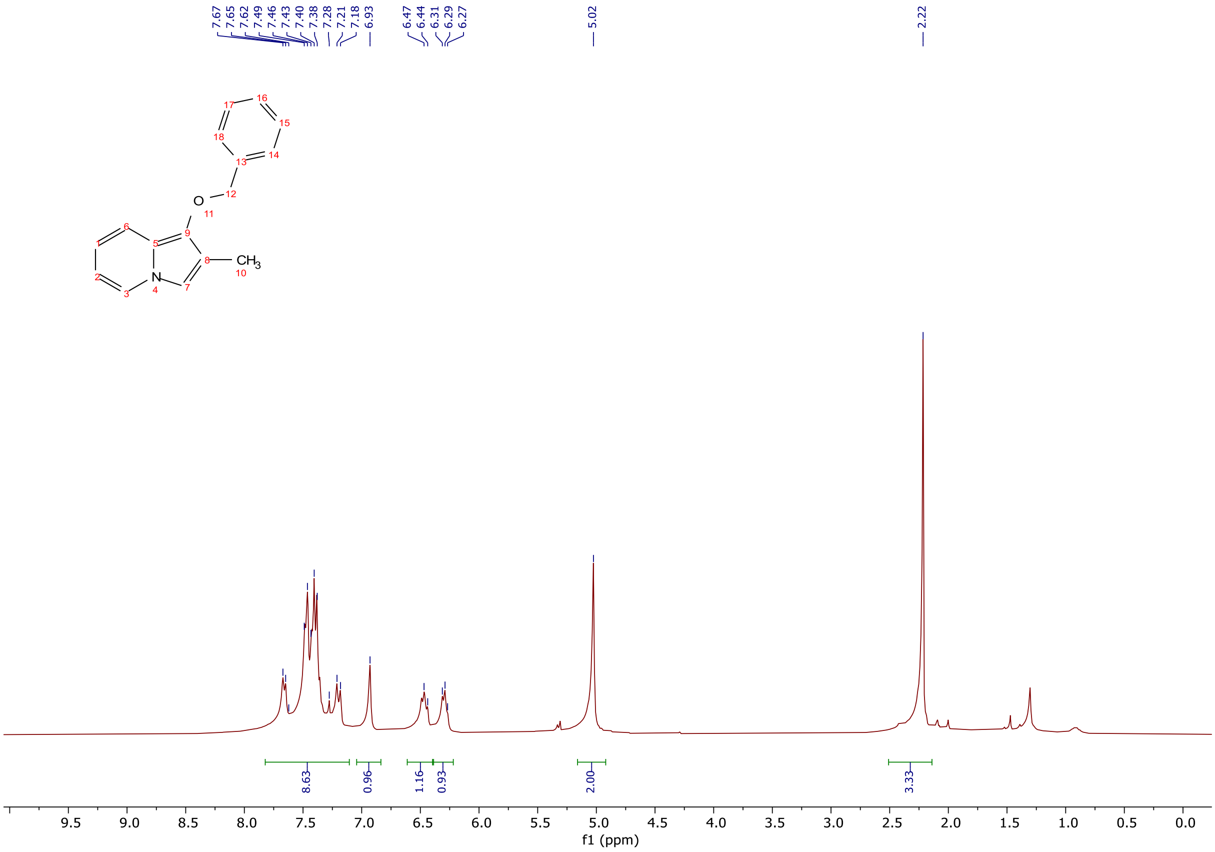

$^1$H NMR spectra (300 MHz, $CDCl_3$) of 1-(benzyloxy)-2-methylindolizine.

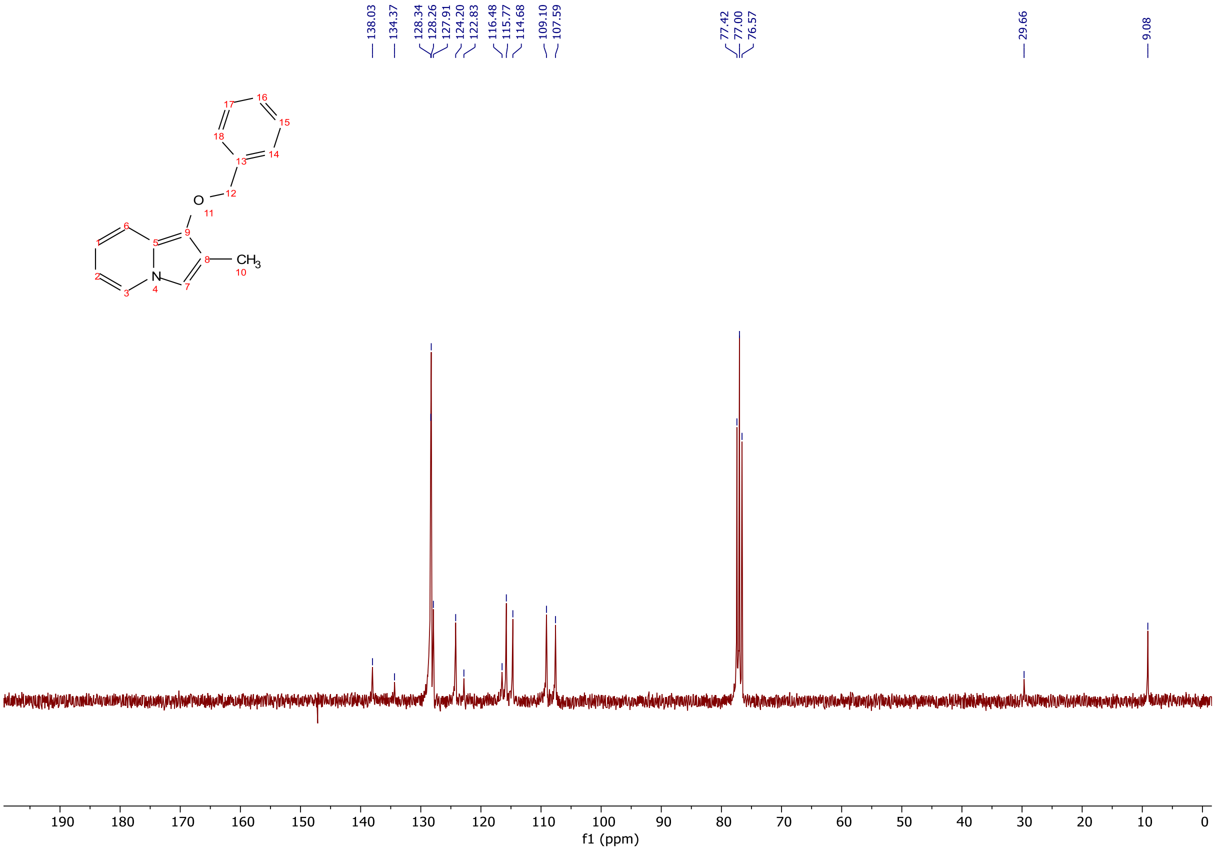

$^{13}$C NMR spectra (300 MHz, $CDCl_3$) of 1-(benzyloxy)-2-methylindolizine.

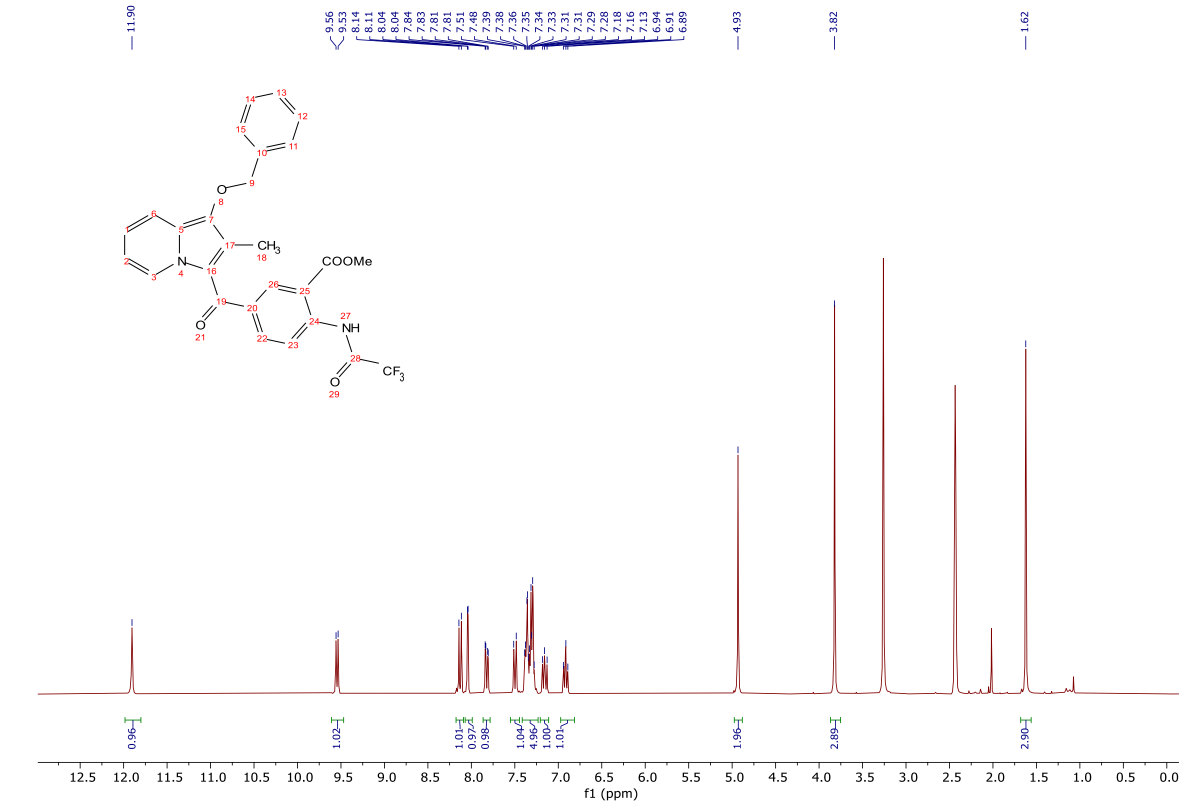


[1]H NMR spectra (300 MHz, DMSO-$d_6$) of methyl 5-(1-(benzyloxy)-2-methylindolizine-3-carbonyl)-2-(2,2,2-trifluoroacetamido)benzoate.

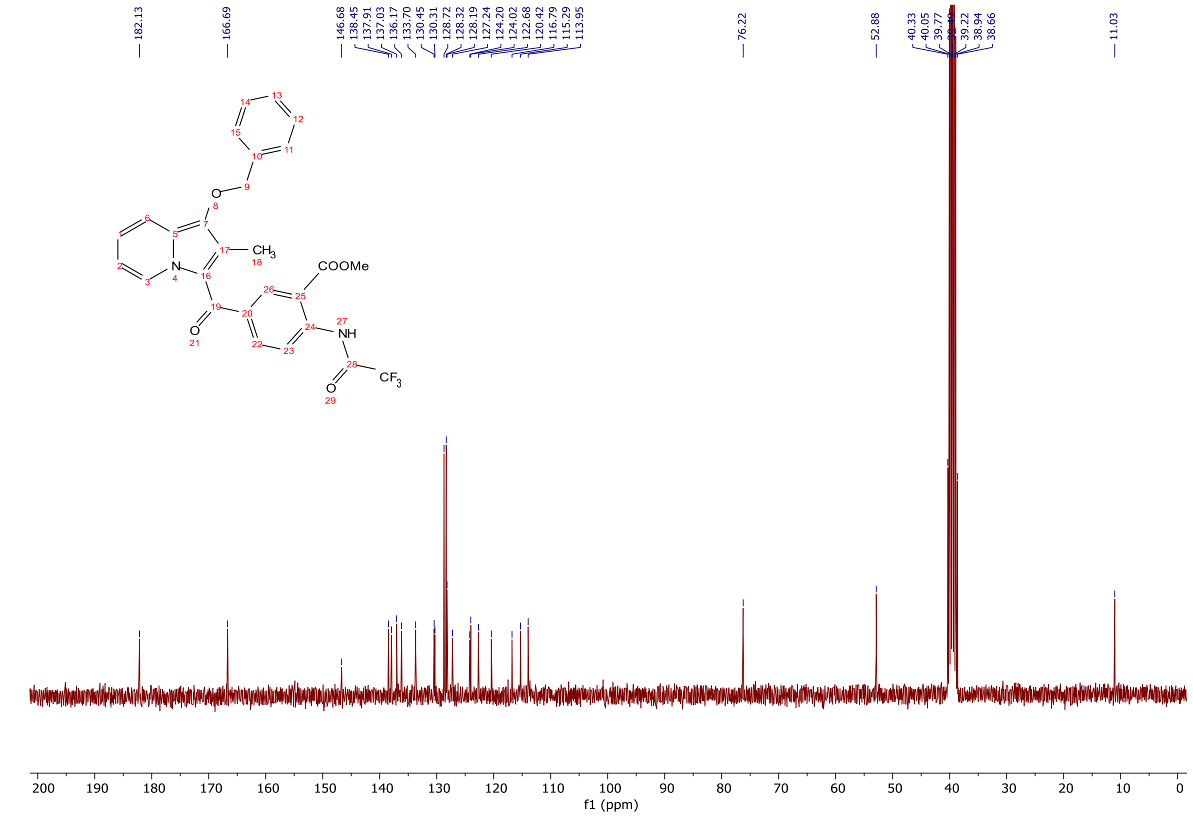


[13]C NMR spectra (75 MHz, DMSO-$d_6$) of methyl 5-(1-(benzyloxy)-2-methylindolizine-3-carbonyl)-2-(2,2,2-trifluoroacetamido)benzoate.

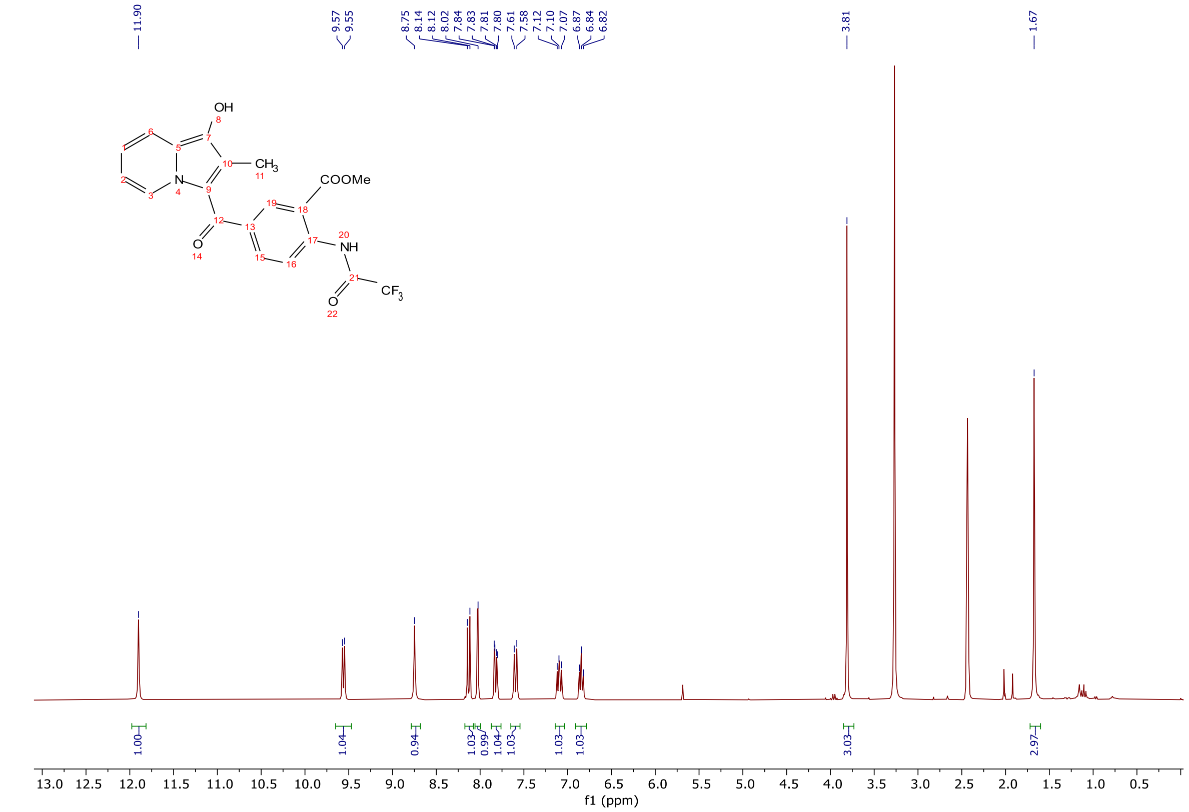


$^{1}H$ NMR spectra (300 MHz, DMSO-$d_6$) of methyl 5-(1-hydroxy-2-methylindolizine-3-carbonyl)-2-(2,2,2-trifluoroacetamido)benzoate.

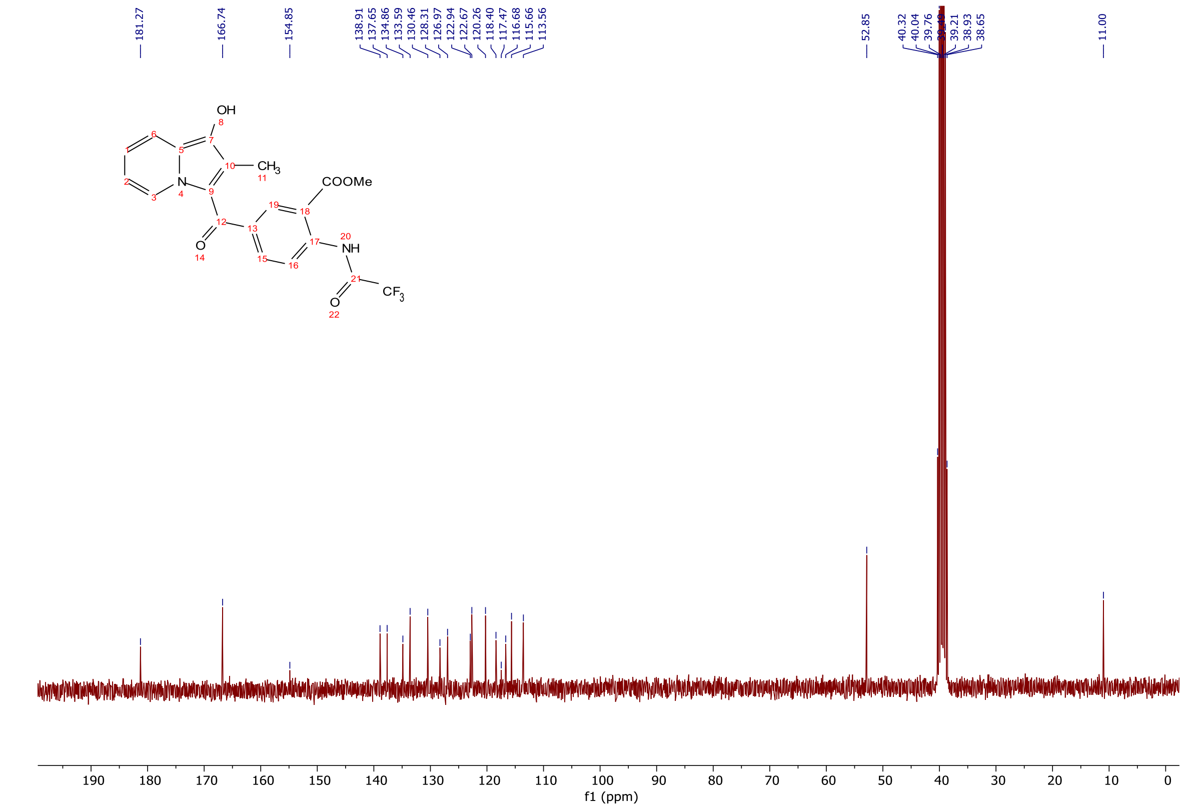


$^{13}C$ NMR spectra (75 MHz, DMSO-$d_6$) of methyl 5-(1-hydroxy-2-methylindolizine-3-carbonyl)-2-(2,2,2-trifluoroacetamido)benzoate.

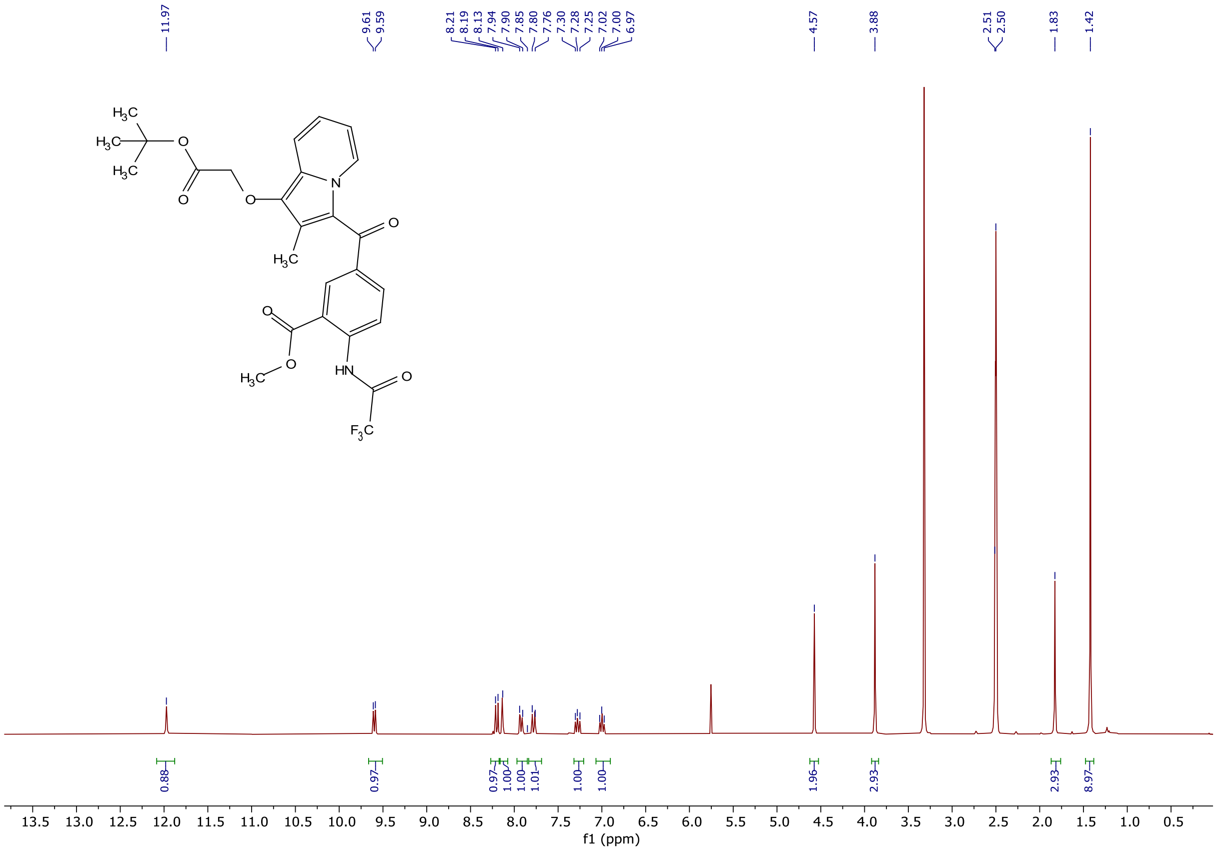

$^1$H NMR spectra (300 MHz, DMSO-$d_6$) of methyl 5-(1-(2-(tert-butoxy)-2-oxoethoxy)-2-methylindolizine-3-carbonyl)-2-(2,2,2-trifluoroacetamido)benzoate.

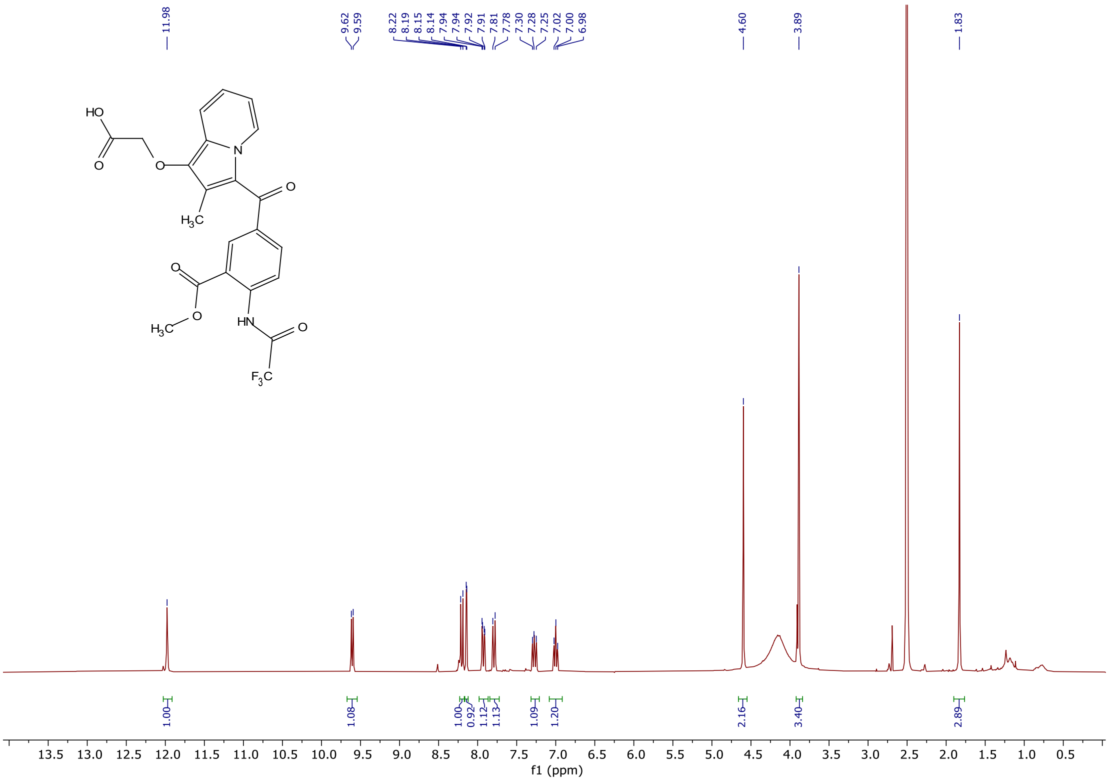

$^1$H NMR spectra (300 MHz, DMSO-$d_6$) of 2-((3-(3-(methoxycarbonyl)-4-(2,2,2-trifluoroacetamido)benzoyl)-2-methylindolizin-1-yl)oxy)acetic acid.

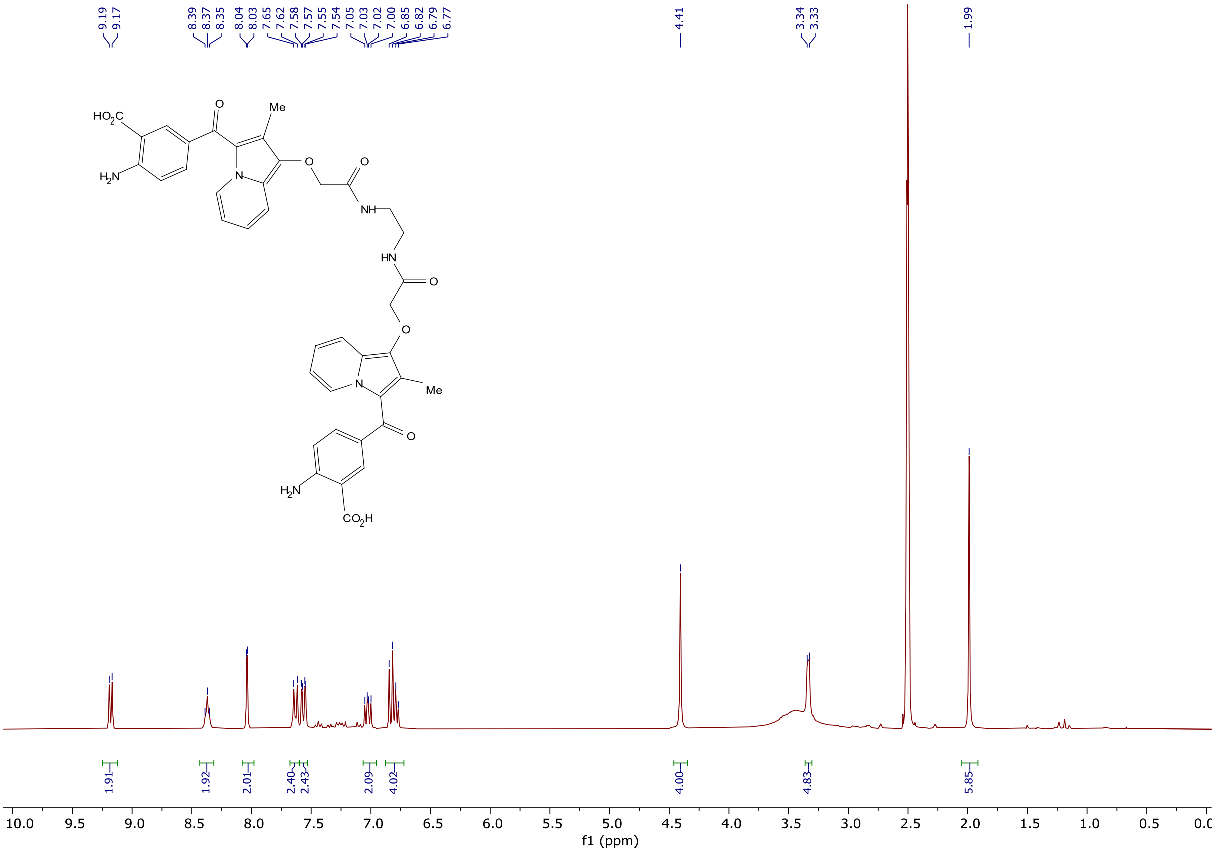

$^1H$ NMR spectra (300 MHz, DMSO-$d_6$) of **SI-1**.

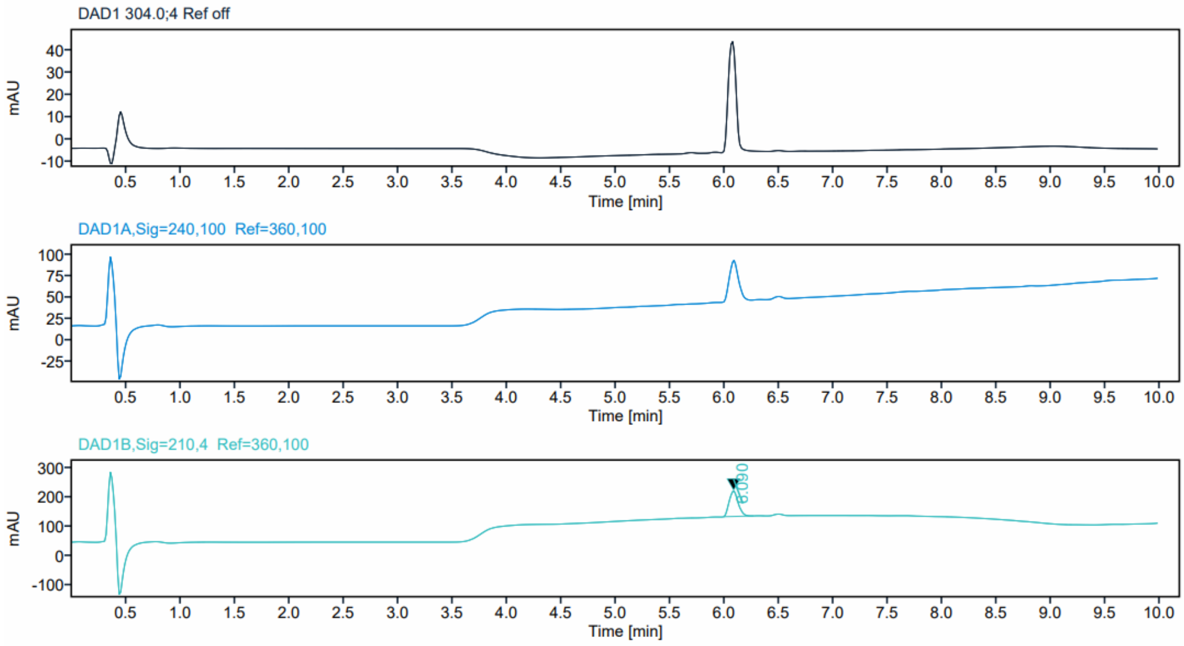

HPLC profiles of **SI-1**, detected at 304.4, 240.1 and 210.4 nm.

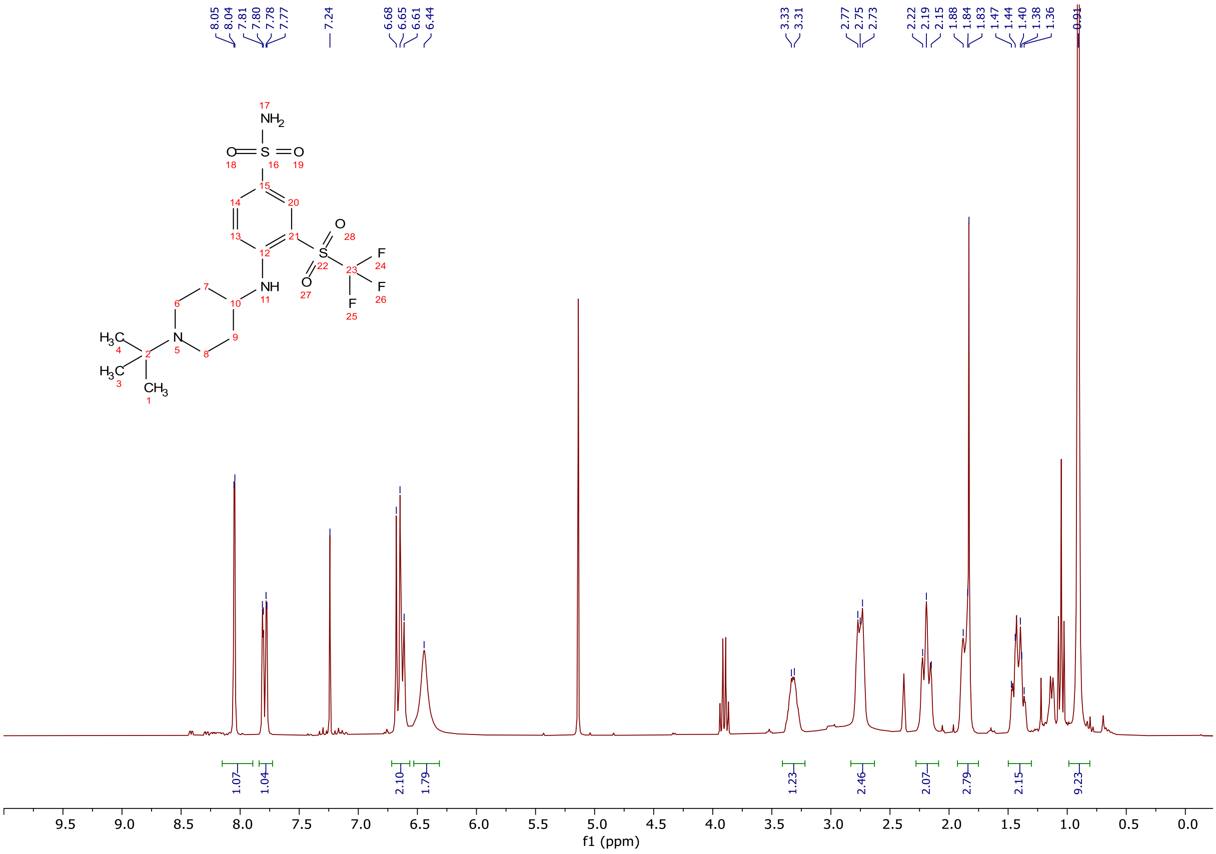


$^{1}$H NMR spectra (300 MHz, $CDCl_3$) of **VS-1-1**.

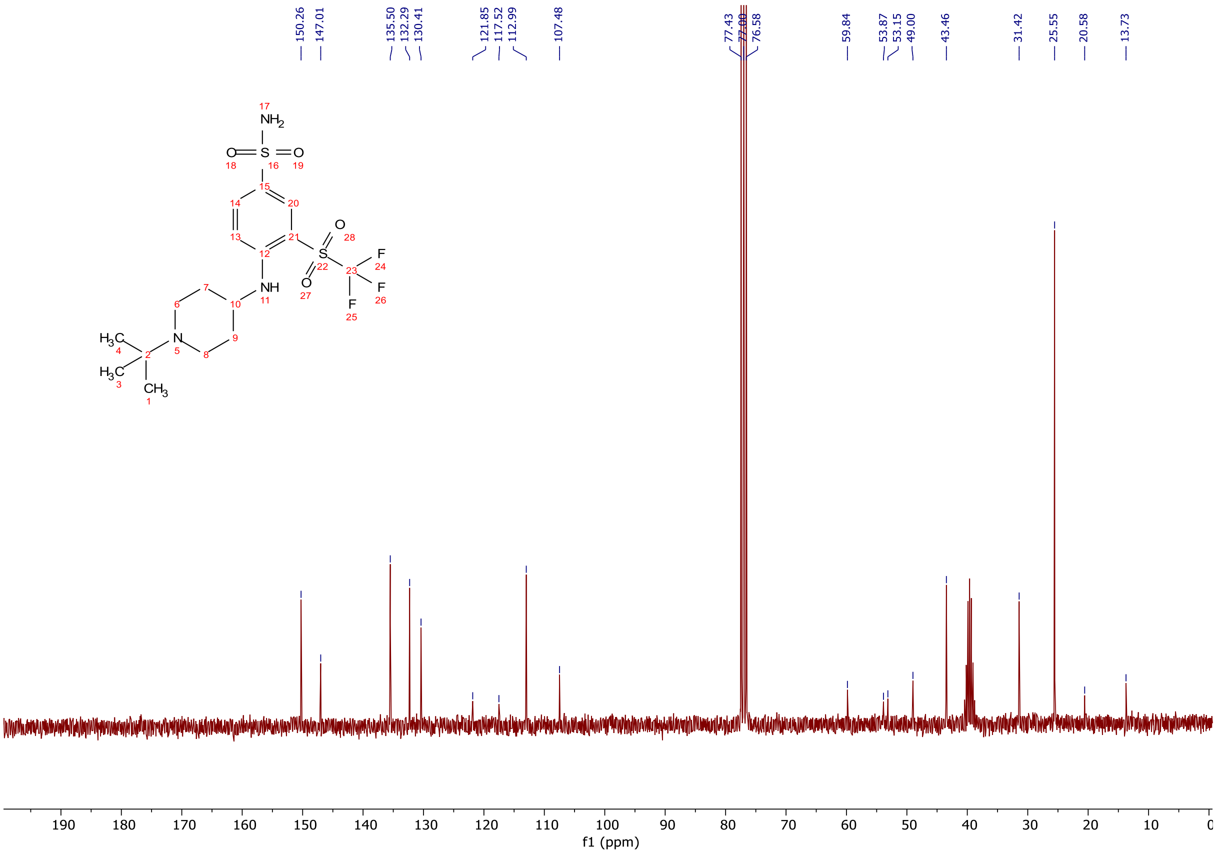


$^{13}$C NMR spectra (75 MHz, $CDCl_3$) of **VS-1-1**.

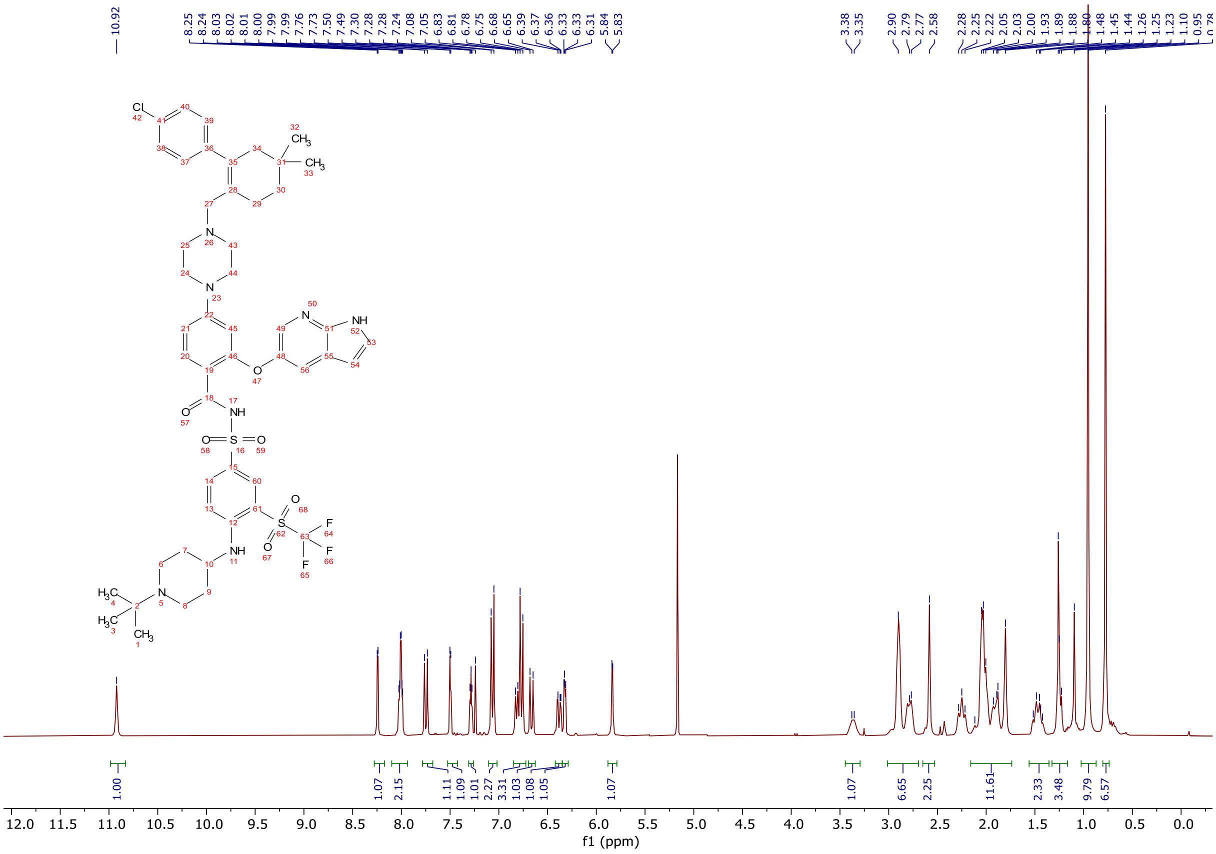

$^{1}H$ NMR spectra (300 MHz, $CDCl_3$) of **VS-1**.

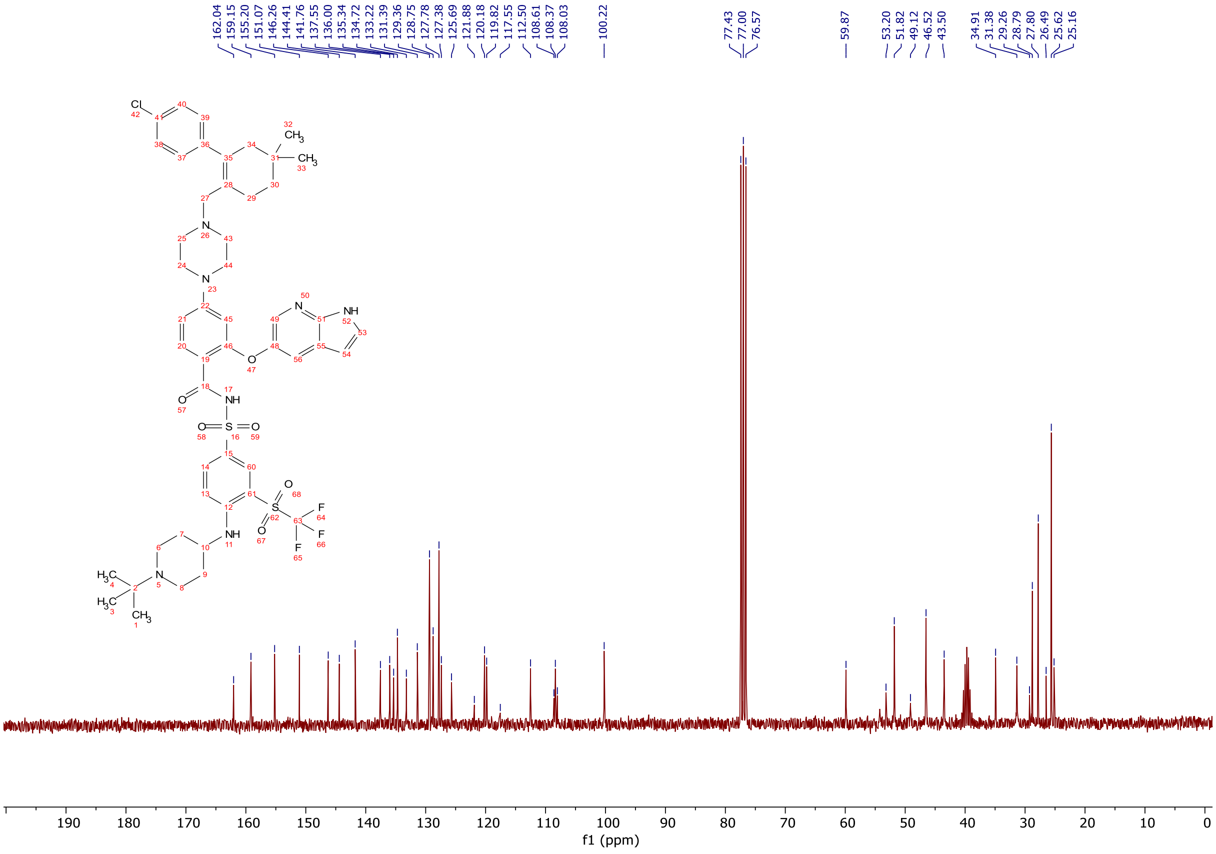

$^{13}C$ NMR spectra (75 MHz, $CDCl_3$) of **VS-1**.

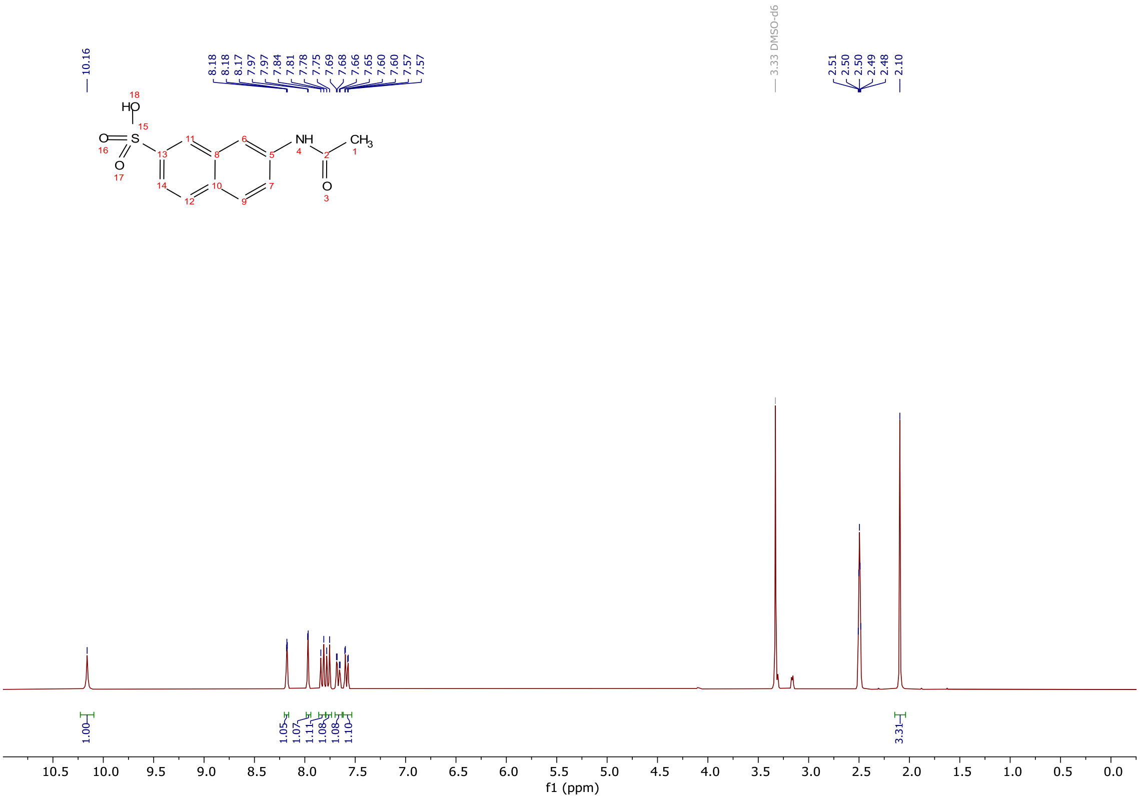


$^1$H NMR spectra (300 MHz, DMSO-d$_6$) of sodium 7-acetamidonaphthalene-2-sulfonate.

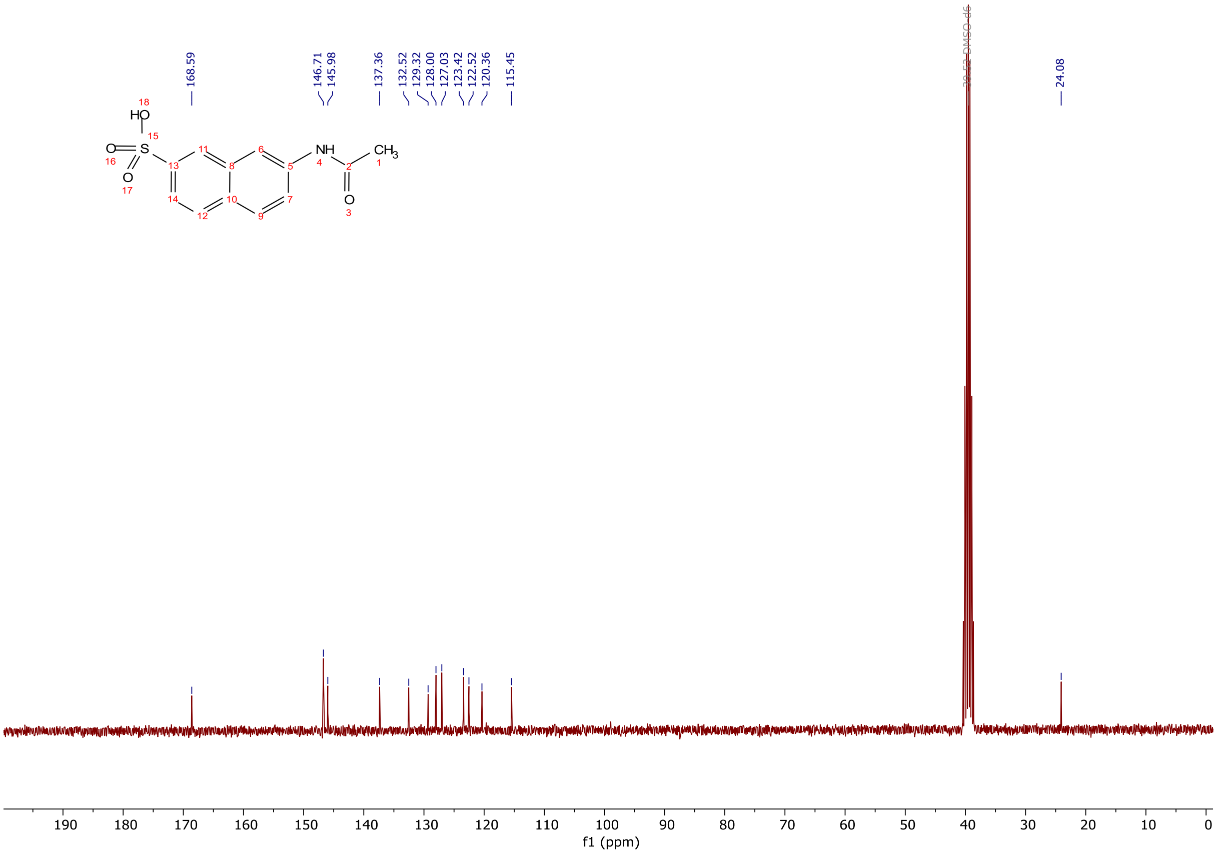


$^{13}$C NMR spectra (75 MHz, DMSO-d$_6$) of sodium 7-acetamidonaphthalene-2-sulfonate.

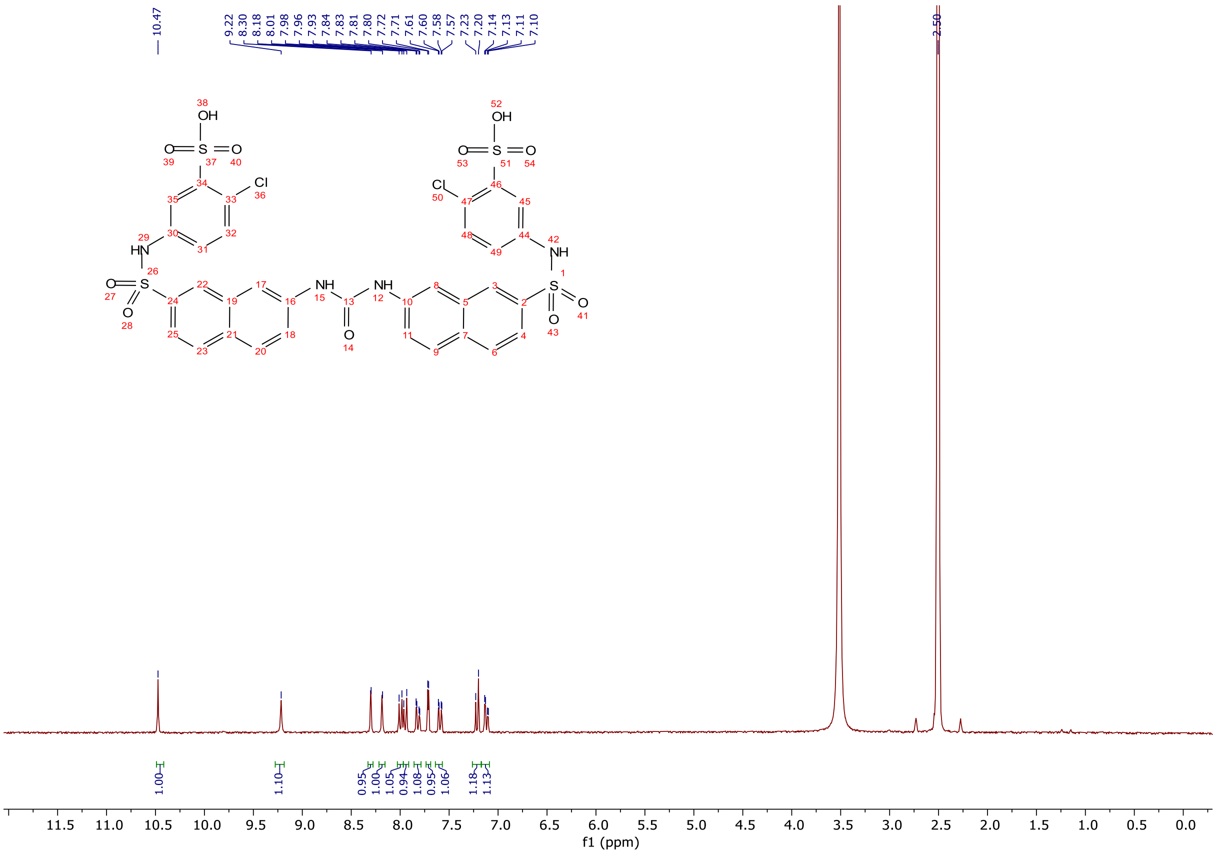


$^{1}$H NMR spectra (300 MHz, DMSO-$d_6$) of **VS-2**.

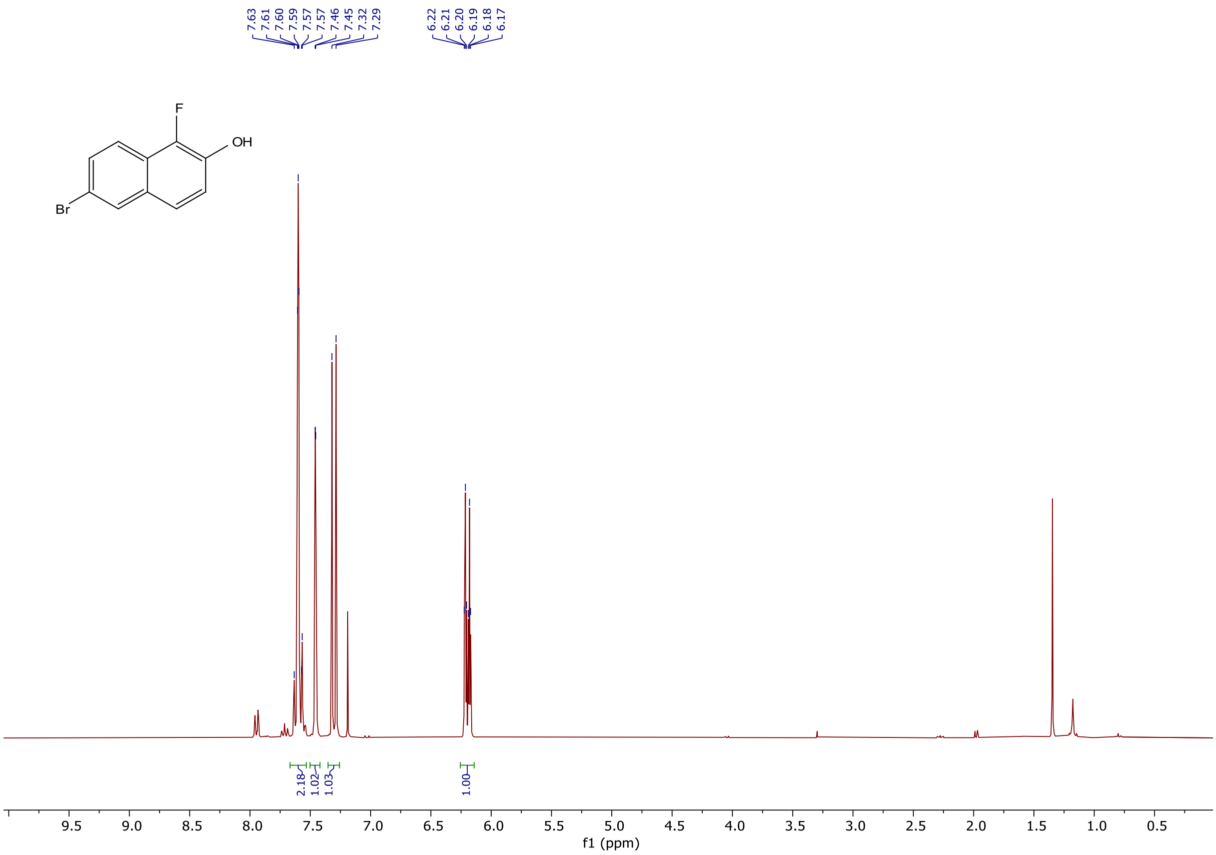


$^{1}$H NMR spectra (300 MHz, $CDCl_3$) of 6-bromo-1-fluoronaphthalen-2-ol.

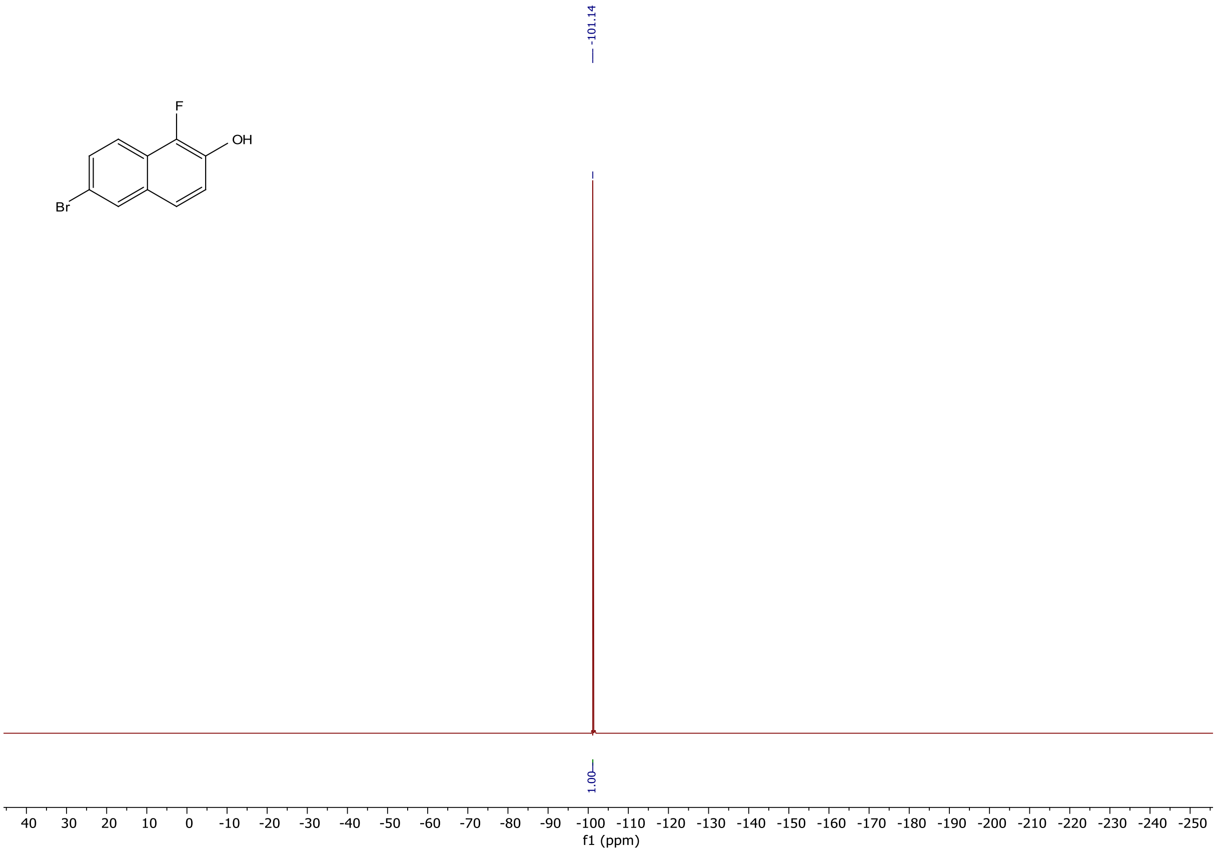


$^{19}F$ NMR spectra (283 MHz, $CDCl_3$) of 6-bromo-1-fluoronaphthalen-2-ol.

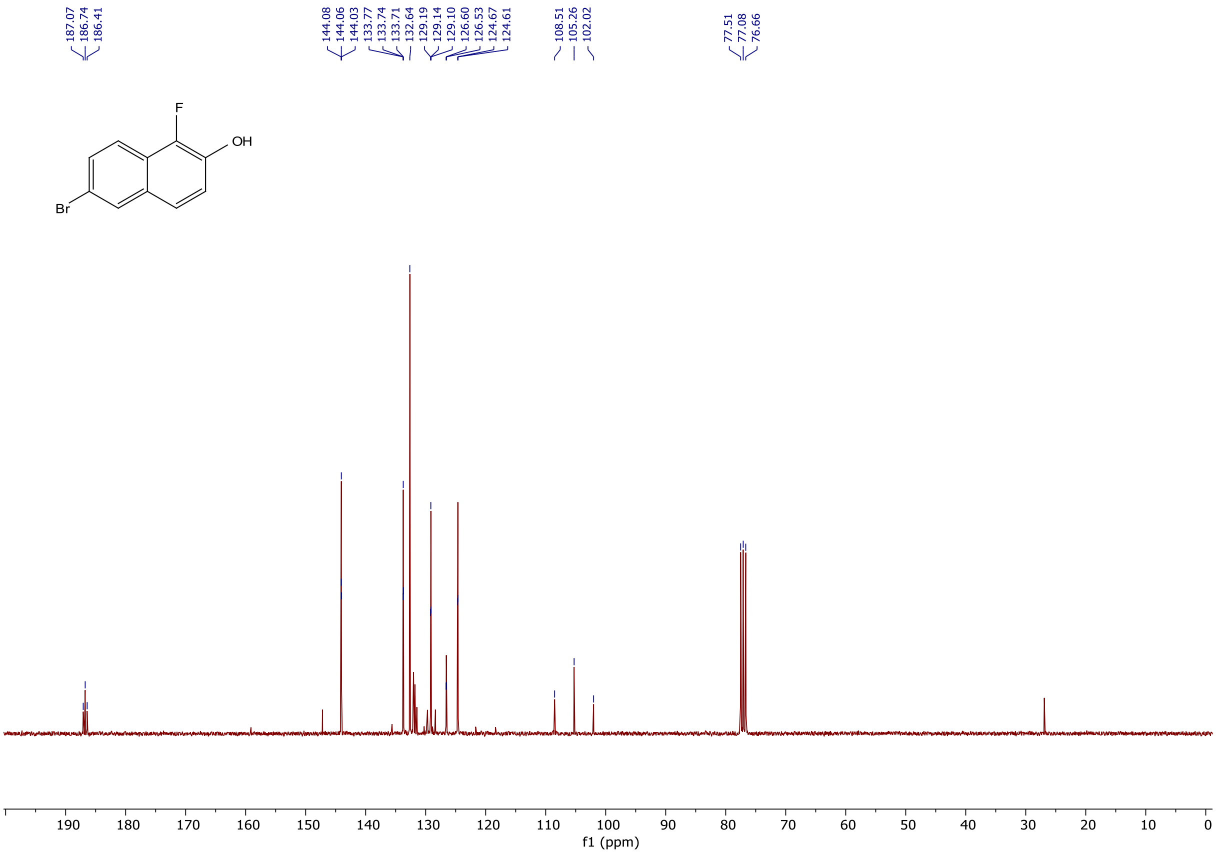


$^{13}C$ NMR spectra (75 MHz, $CDCl_3$) of 6-bromo-1-fluoronaphthalen-2-ol.

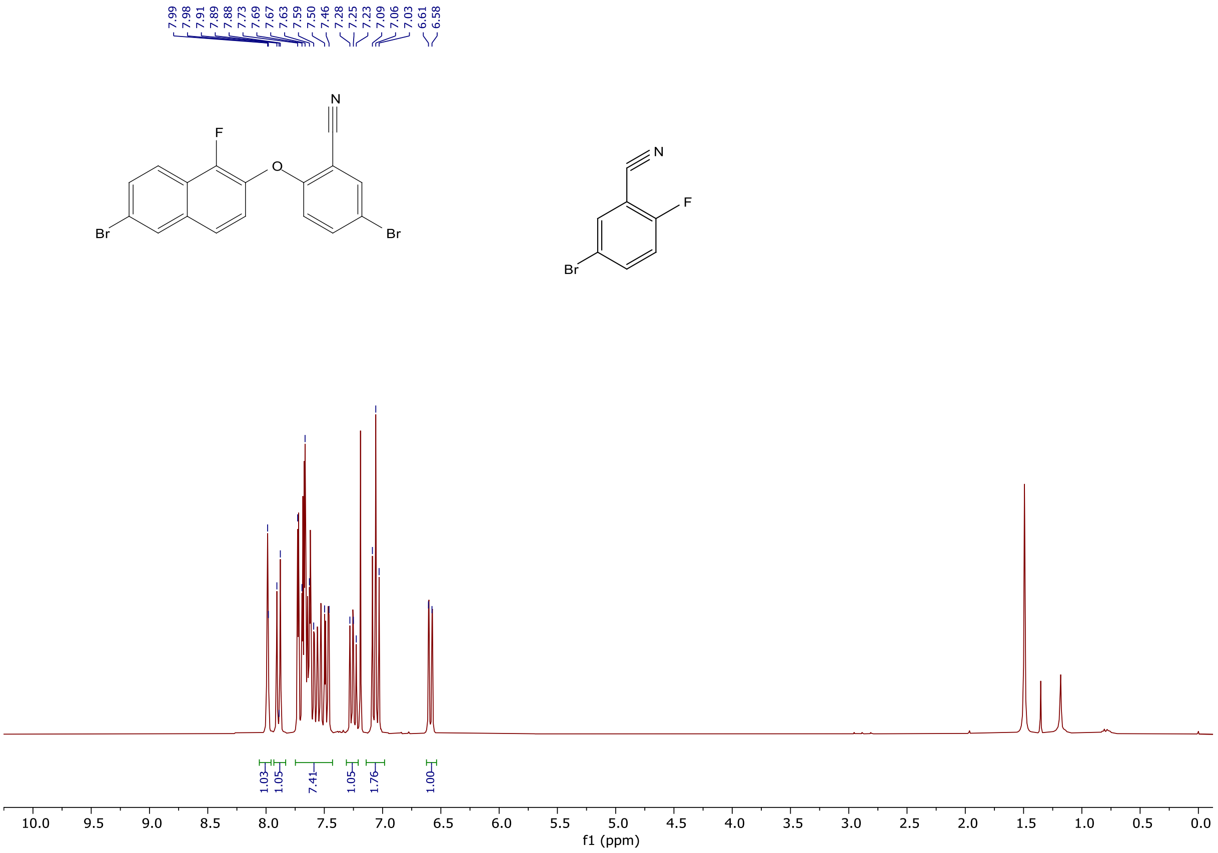


$^1$H NMR spectra (300 MHz, $CDCl_3$) of 4-bromo-2-((6-bromo-1-fluoronaphthalen-2-yl)oxy)benzonitrile.

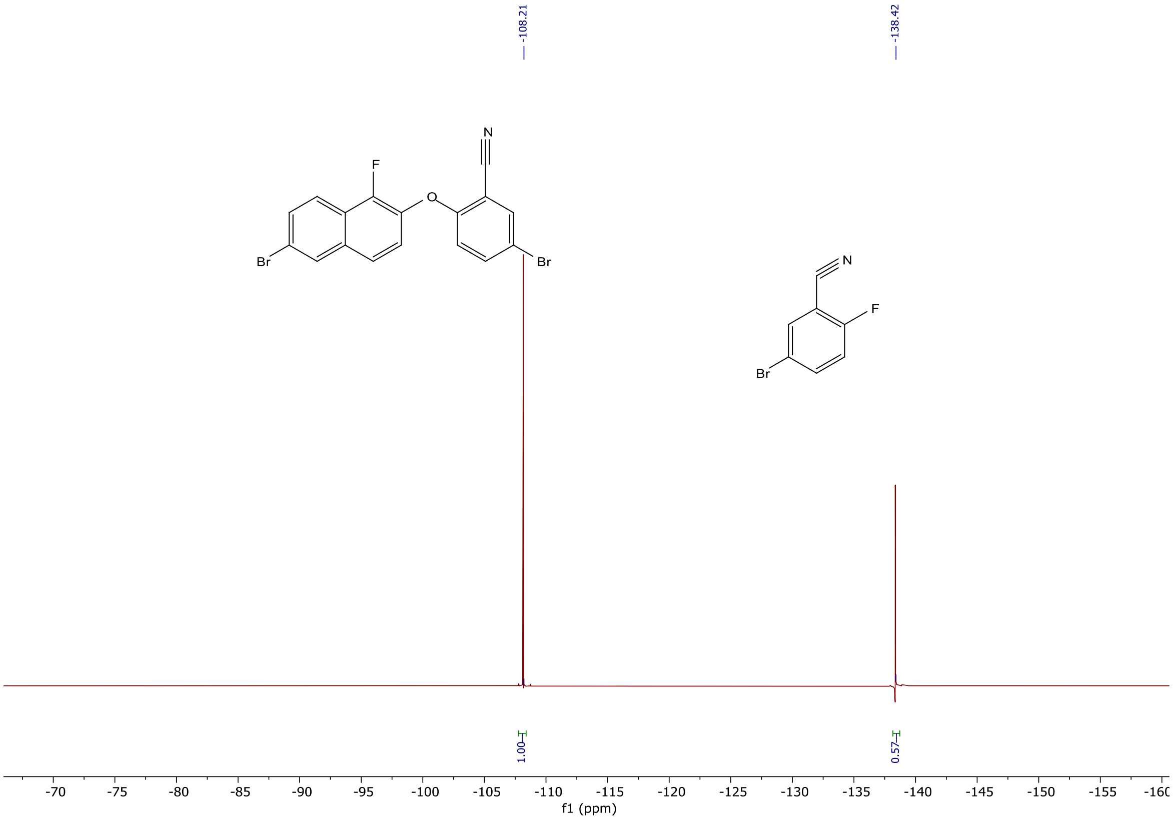


$^{19}$F NMR spectra (300 MHz, $CDCl_3$) of 4-bromo-2-((6-bromo-1-fluoronaphthalen-2-yl)oxy)benzonitrile.

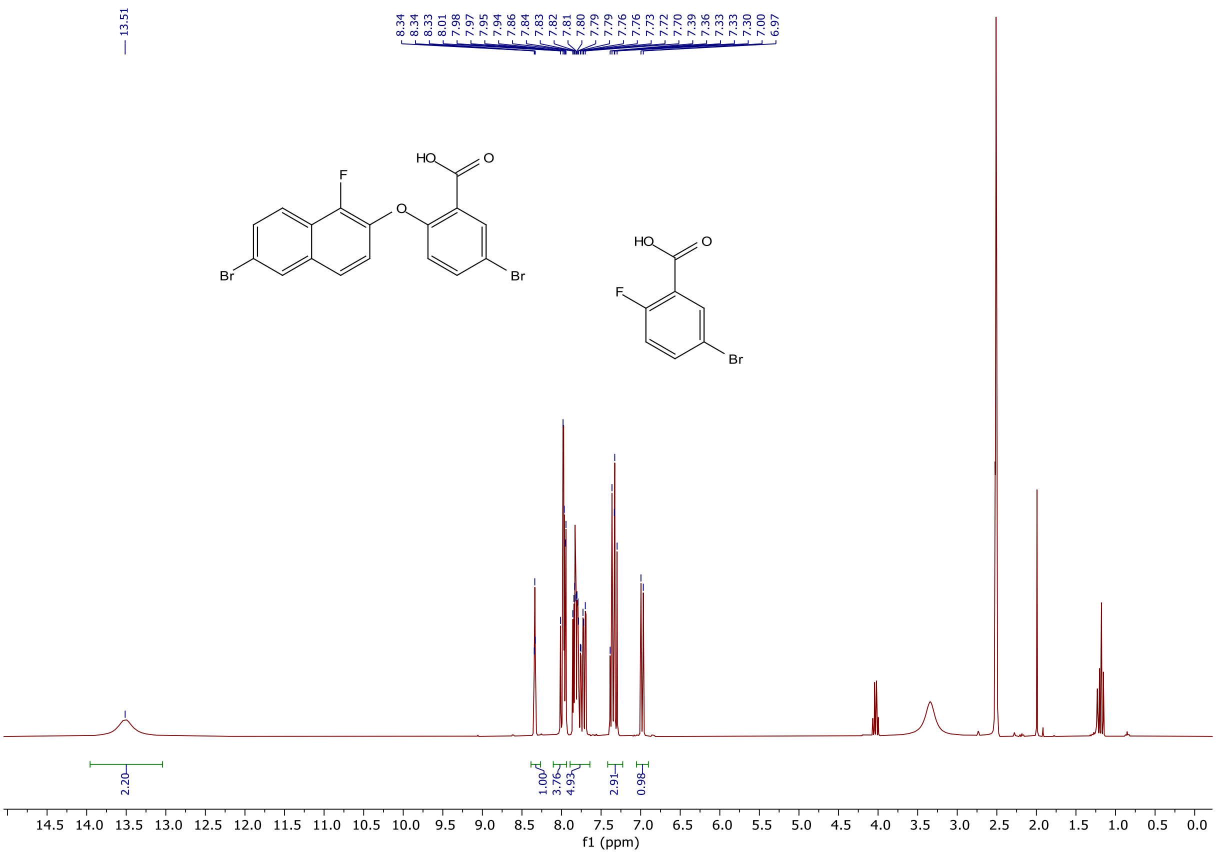

[1]H NMR spectra (300 MHz, DMSO-$d_6$) of 4-bromo-2-((6-bromo-1-fluoronaphthalen-2-yl)oxy)benzoic acid.

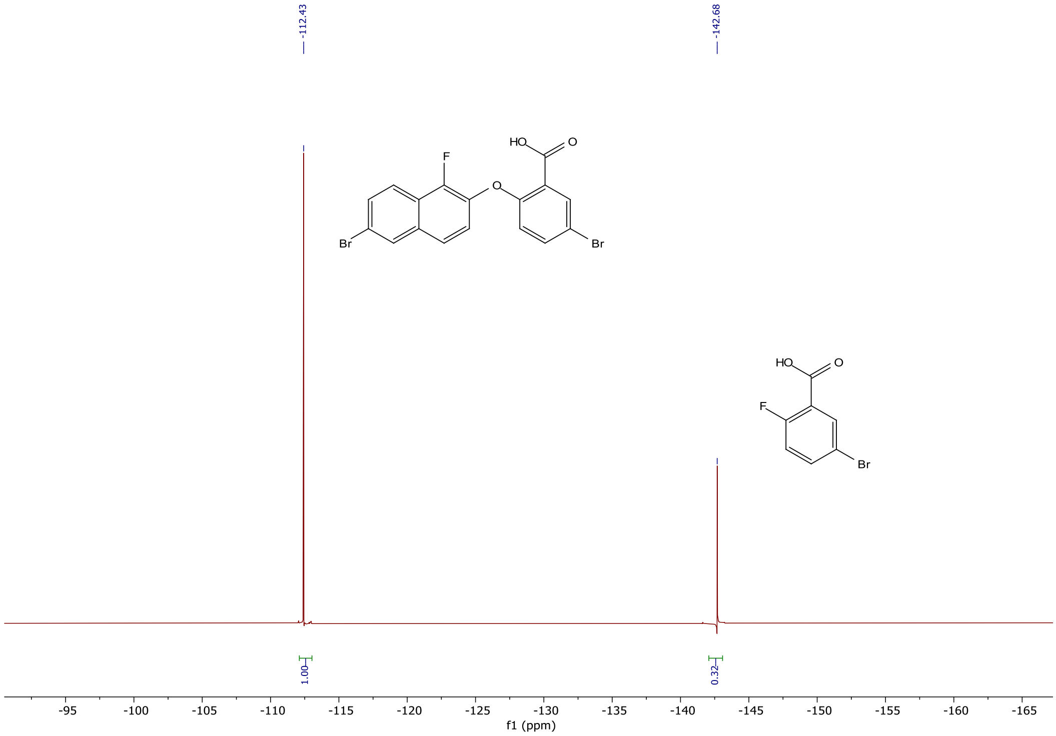

[19]F NMR spectra (283 MHz, DMSO-$d_6$) of 4-bromo-2-((6-bromo-1-fluoronaphthalen-2-yl)oxy)benzoic acid.

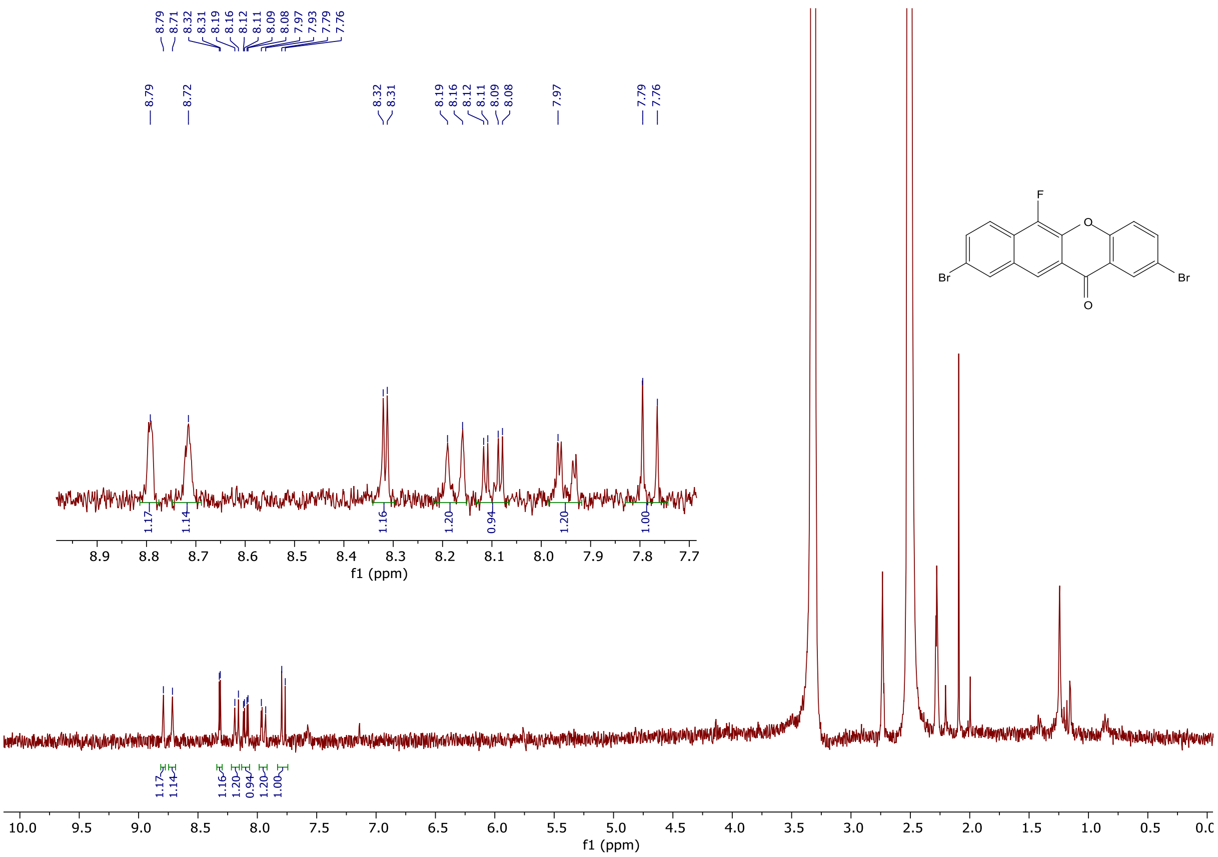

[1]H NMR spectra (301 MHz, DMSO-$d_6$) of 2,9-dibromo-6-fluoro-12H-benzo[b]xanthen-12-one.

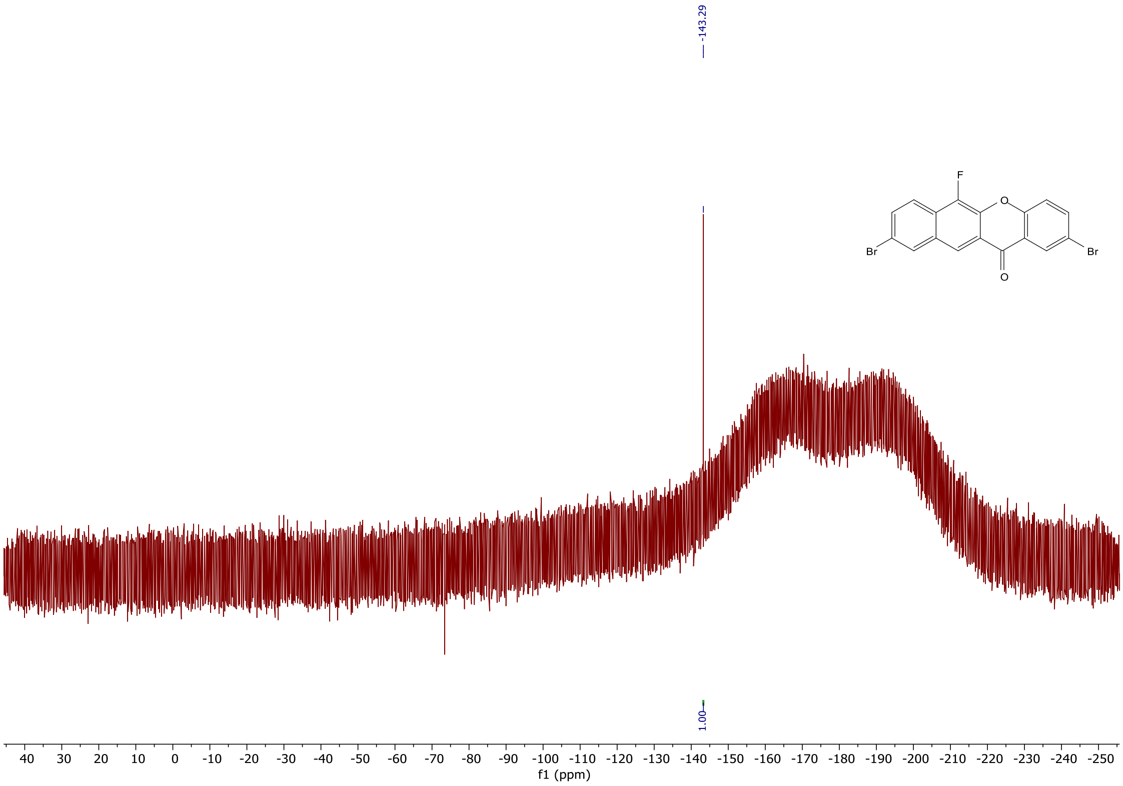

[19]F NMR spectra (283 MHz, DMSO-$d_6$) of 2,9-dibromo-6-fluoro-12H-benzo[b]xanthen-12-one.

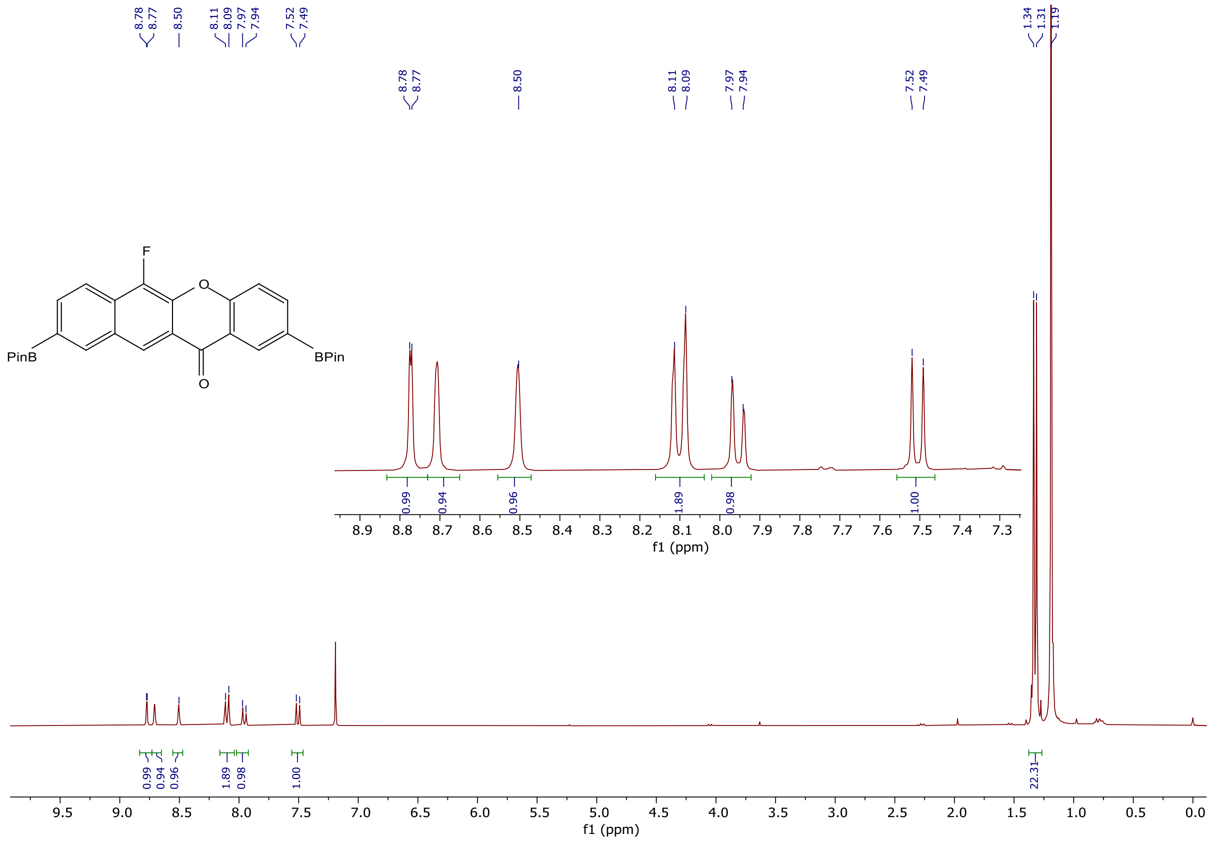


$^1$H NMR spectra (300 MHz, $CDCl_3$) of 6-fluoro-2,9-bis(4,4,5,5-tetramethyl-1,3,2-dioxaborolan-2-yl)-12H-benzo[b]xanthen-12-one.

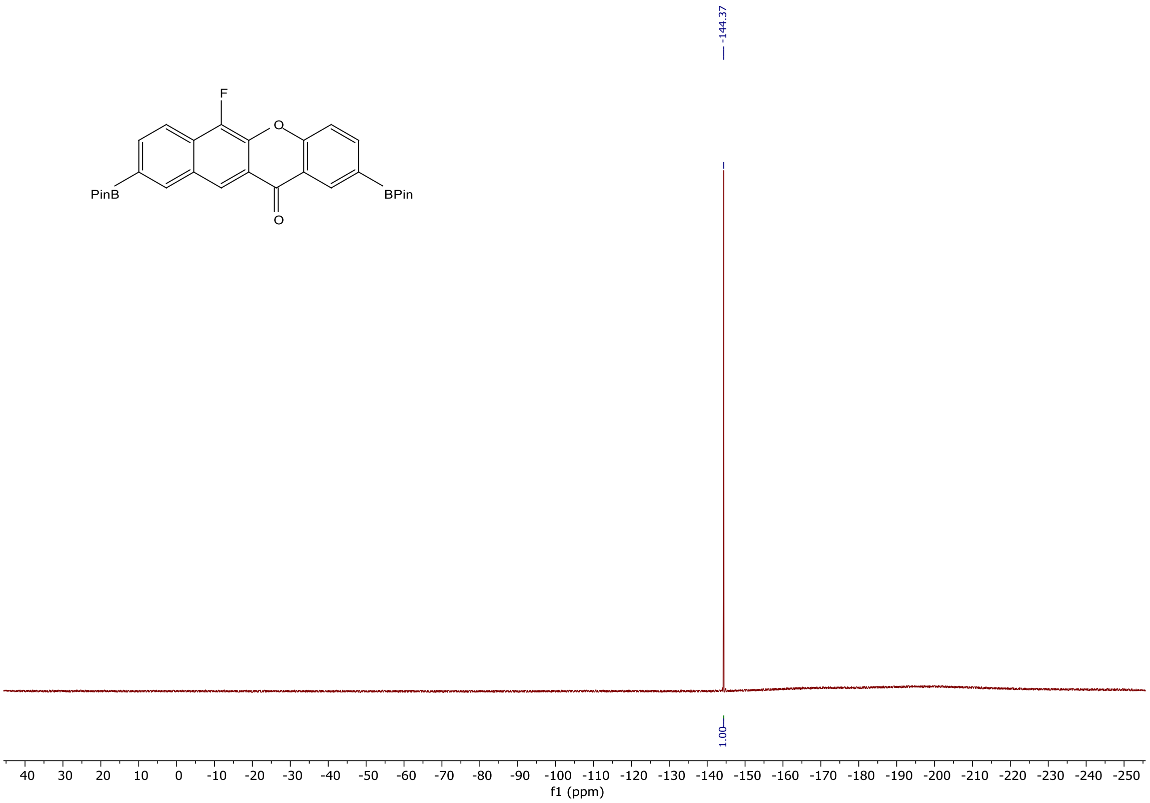


$^{19}$F NMR spectra (283 MHz, $CDCl_3$) of 6-fluoro-2,9-bis(4,4,5,5-tetramethyl-1,3,2-dioxaborolan-2-yl)-12H-benzo[b]xanthen-12-one.

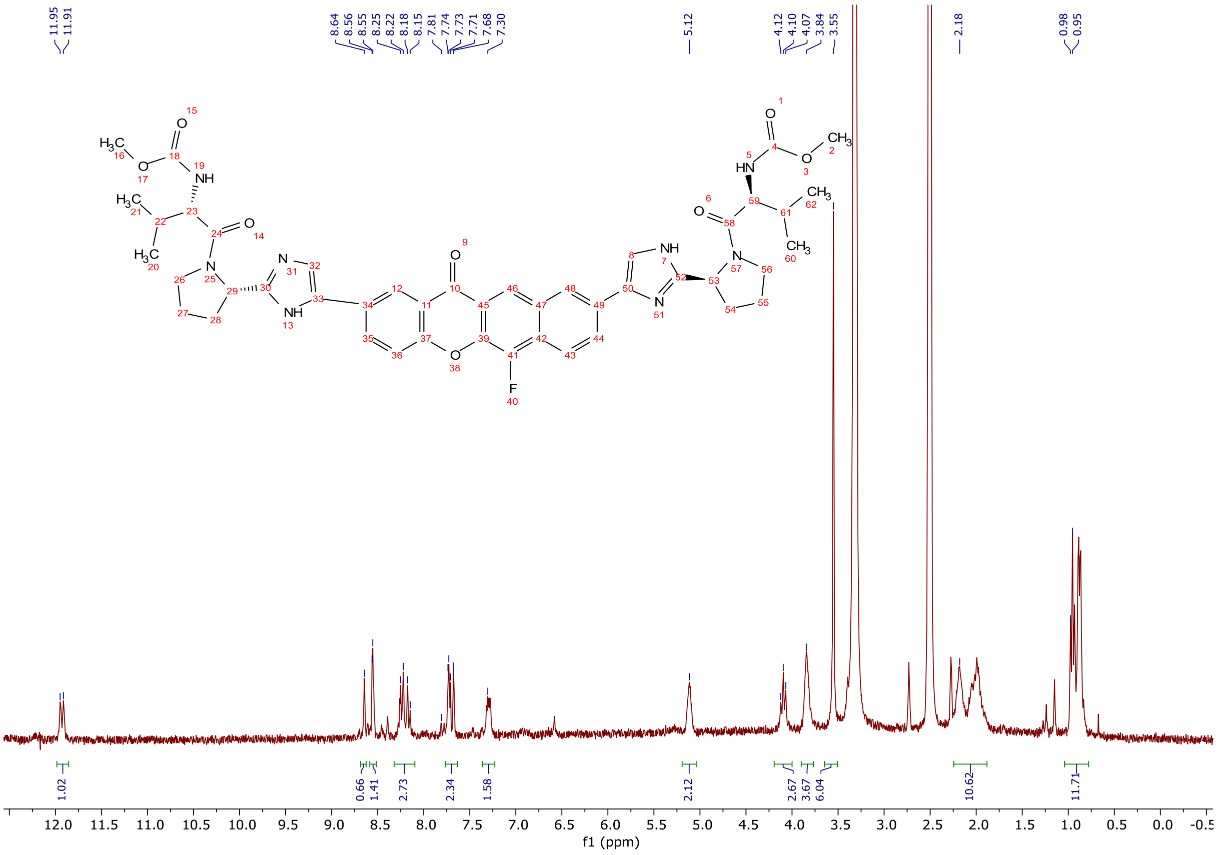

[1]H NMR spectra (300 MHz, DMSO-d$_6$) of **VS-3**.

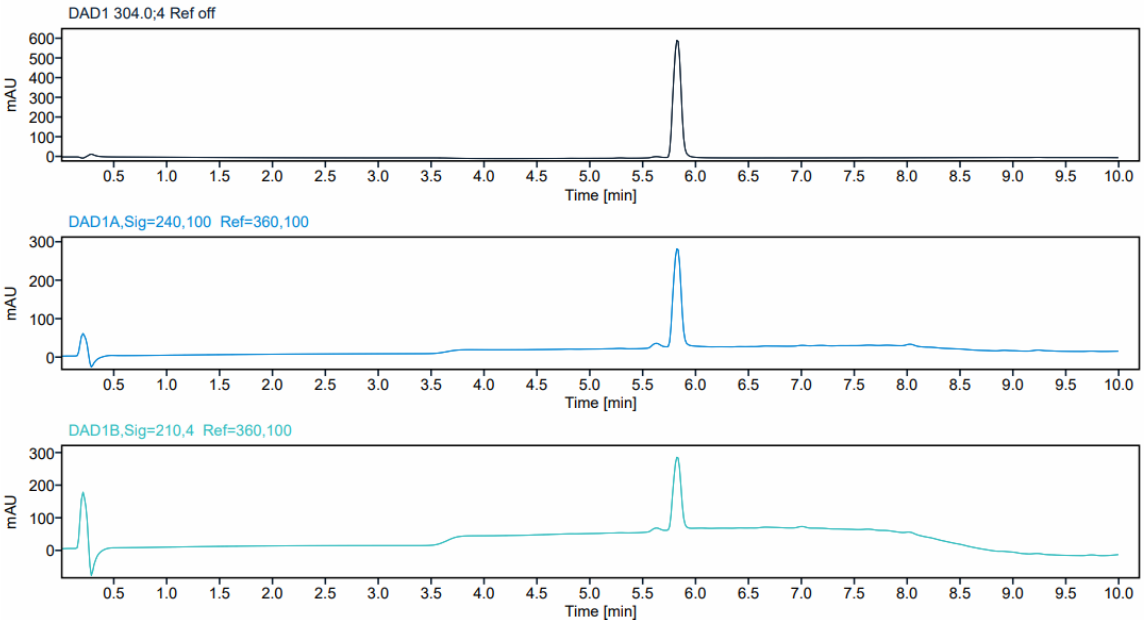


HPLC profiles of **VS-3**, detected at 304.4, 240.1 and 210.4 nm.
